# Unlocking Mitochondrial Dysfunction-Associated Senescence (MiDAS) with NAD^+^ – a Boolean Model of Mitochondrial Dynamics and Cell Cycle Control

**DOI:** 10.1101/2023.12.18.572194

**Authors:** Herbert Sizek, Dávid Deritei, Katherine Fleig, Marlayna Harris, Peter L. Regan, Kimberly Glass, Erzsébet Ravasz Regan

## Abstract

The steady accumulation of senescent cells with aging creates tissue environments that aid cancer evolution. Aging cell states are highly heterogeneous. ‘Deep senescent’ cells rely on healthy mitochondria to fuel a strong proinflammatory secretome, including cytokines, growth and transforming signals. Yet, the physiological triggers of senescence such as the reactive oxygen species (ROS) can also trigger mitochondrial dysfunction, and sufficient energy deficit to alter their secretome and cause chronic oxidative stress – a state termed Mitochondrial Dysfunction-Associated Senescence (MiDAS). Here, we offer a mechanistic hypothesis for the molecular processes leading to MiDAS, along with testable predictions. To do this we have built a Boolean regulatory network model that qualitatively captures key aspects of mitochondrial dynamics during cell cycle progression (hyper-fusion at the G1/S boundary, fission in mitosis), apoptosis (fission and dysfunction) and glucose starvation (reversible hyper-fusion), as well as MiDAS in response to *SIRT3* knockdown or oxidative stress. Our model reaffirms the protective role of NAD^+^ and external pyruvate. We offer testable predictions about the growth factor- and glucose-dependence of MiDAS and its reversibility at different stages of reactive oxygen species (ROS)-induced senescence. Our model provides mechanistic insights into the distinct stages of DNA-damage induced senescence, the relationship between senescence and epithelial-to-mesenchymal transition in cancer and offers a foundation for building multiscale models of tissue aging.

**Highlights:**

- Boolean regulatory network model reproduces <u>mitochondrial dynamics</u> during cell cycle progression, apoptosis, and glucose starvation.
- Model offers a mechanistic explanation for the positive feedback loop that locks in <u>Mitochondrial Dysfunction-Associated Senescence</u> (MiDAS), involving autophagy-resistant, hyperfused, dysfunctional mitochondria.
- Model reproduces <u>ROS-mediated mitochondrial dysfunction</u> and suggests that MiDAS is part of the early phase of damage-induced senescence.
- Model <u>predicts</u> that cancer-driving mutations that bypass the G1/S checkpoint generally increase the incidence of MiDAS, except for p53 loss.

## Introduction

Aging is a major risk factor for cancer [1]. One likely culprit rendering aged tissues hospitable to tumor progression is the prevalence of senescent cells [2–5]. As senescence involves permanent cell cycle arrest, it was thought to be primarily tumor suppressive [6]. The inability of senescent cells to become cancerous themselves is, unfortunately, counterbalanced by their effects on their neighbors. Senescent cells secrete a complex cocktail of signals termed the senescence-associated secretory phenotype (SASP), which drives proliferation, migration, and epithelial-to-mesenchymal transition (EMT) in their neighbors [6,7]. In brief localized doses, SASP can aid wound healing – as long as senescent cells are subsequently cleared by the immune system [8,9]. In contrast, chronic SASP amplified in aging tissues by high senescent cell density also causes oxidative damage and senescence-induced senescence [10]. Mixed with proliferative and EMT-promoting signals, this SASP microenvironment is favorable for chronic inflammation and cancer evolution [6,7].

An aspect of tissue aging associated with oxidative damage is mitochondrial dysfunction, which has been documented to increase with age in both post-mitotic cells and mitotically active tissues [11,12]. The hallmark of mitochondrial dysfunction is a low-efficiency electron transport chain (ETC) and reduced ATP production, accompanied by low cellular and mitochondrial NAD^+^/NADH ratios and increased Reactive Oxygen Species (ROS) production (**Fig. 1A**, *left vs. middle*) [11]. Mitochondrial dysfunction has been shown both to trigger senescence and be a feature of senescent cells [11,13,14]. Several plausible, potentially redundant mechanisms were proposed for the former, including ROS-induced DNA damage [15] and p53 activation by the energy sensor AMPK (**Fig. 1A**, *middle*) [16]. An in-depth comparison of senescent cells generated by direct mitochondrial damage versus irradiation in proliferating human and mouse fibroblasts revealed that mitochondrial dysfunction-associated senescence (MiDAS) involved low NAD^+^/NADH ratio and elevated AMPK/p53, compared to irradiated cells [17]. In MiDAS, low NAD^+^ served as both driver and consequence of increased mitochondrial ROS production (main results summarized in **Table 1**).

**Figure 1.**
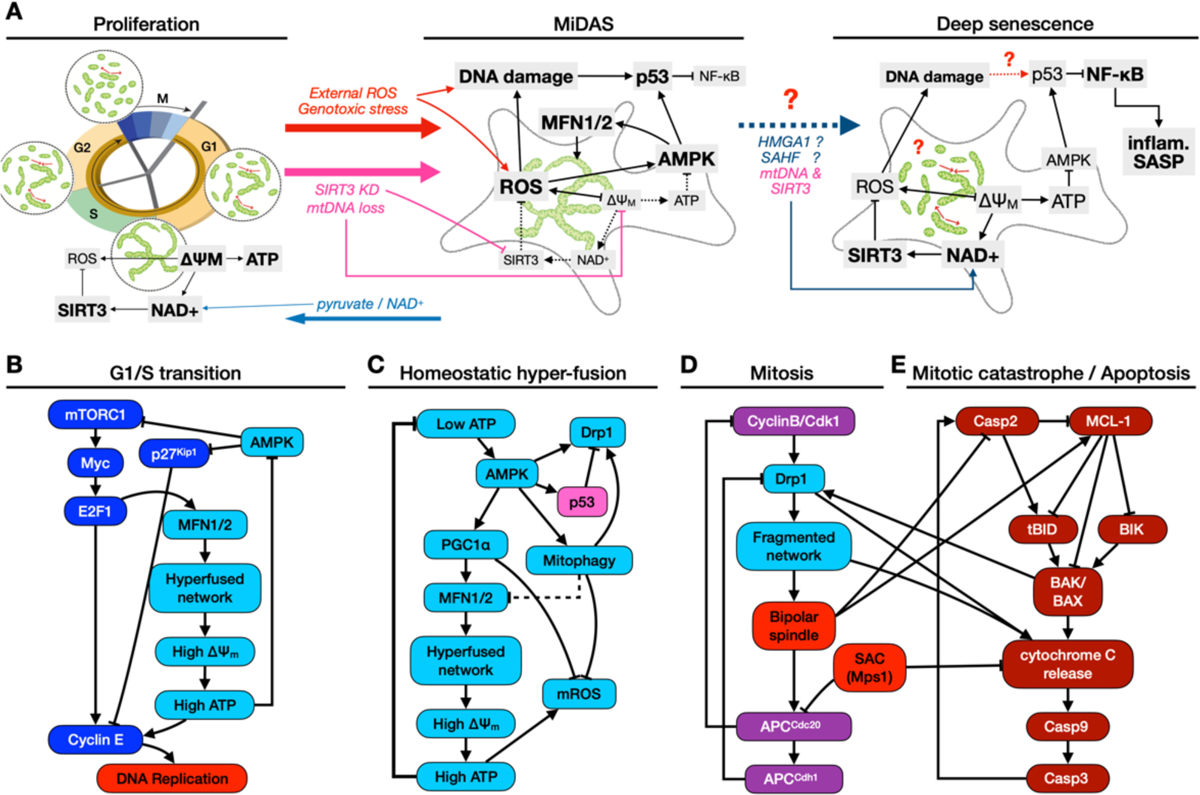
Control of mitochondrial morphology by cell cycle, energy stress and apoptosis. **A)** *Proliferation:* mitochondria in cycling cells hyperfuse before the G1/S transition, increasing their ΔΨ_M_ to boost ATP production for DNA synthesis. Increased ROS generation by the Electron Transport Chain (ETC) is counteracted in large part by NAD^+^ dependent SIRT3 activity. *MiDAS:* perturbations that increase internal ROS directly / via DNA damage (*red arrows*), compromise healthy ETC function and ΔΨ_M_ via the loss of *SIRT3* or mitochondrial DNA (*pink arrows*) lead to chronic mitochondrial dysfunction marked by AMPK and p53 activation, and subsequently MiDAS. We hypothesize that this state involves hyperfused mitochondrial networks with low ΔΨ_M_. MiDAS can be prevented (and potentially reversed) by access to pyruvate or a sustained boost to NAD^+^ (*blue arrows*). *Deep senescence*: in cells with intact SIRT3 and mtDNA, deepening of senescence over ∼6 days following DNA damage appears to rescue mitochondrial dysfunction by HMGA1-mediated NAMPT transcription and NAD^+^ generation (*blue arrows*). This restores normal ATP production, lowers AMPK and p53, and results in NF-κB-dependent inflammatory SASP. **B)** At the G1/S boundary, E2F1 induces microfusin (MFN2) to generate a hyperfused mitochondrial network that can supply the increased ATP demands of DNA synthesis. **C)** A drop in the ATP/AMP ratio activates AMPK, which is part of an incoherent feed-forward loop: its immediate effect on mitochondria is rapid fragmentation through Drp1 [26], subsequently reversed by AMPK-induced p53 [18] and transcriptional Drp1 repression. In addition, AMPK upregulates MFN1/2 transcription via PGC1α [27,28]. The resulting hyperfused network restores ATP, closing a homeostatic negative feedback loop. Increased ATP generation also increases mitochondrial ROS, but both PGC1α’s target genes and AMPK-induced mitophagy work to counteract it. **D)** At the start of mitosis, Cdk1 activates Drp1 to trigger fission, which is required for proper spindle formation and Spindle Assembly Checkpoint (SAC) passage. APC^Cdh1^ then lowers Drp1 levels in anaphase. **E)** During mitotic catastrophe, failure to assemble a bipolar spindle and pass SAC activates Caspase 2, which in turn truncates BID to trigger apoptosis. BAK/BAX recruit Drp1 to mitochondria to fragment them, aiding Cytochrome C release and downstream executioner caspase activation. References for each link in **SM Table 1**.

**Table 1.**
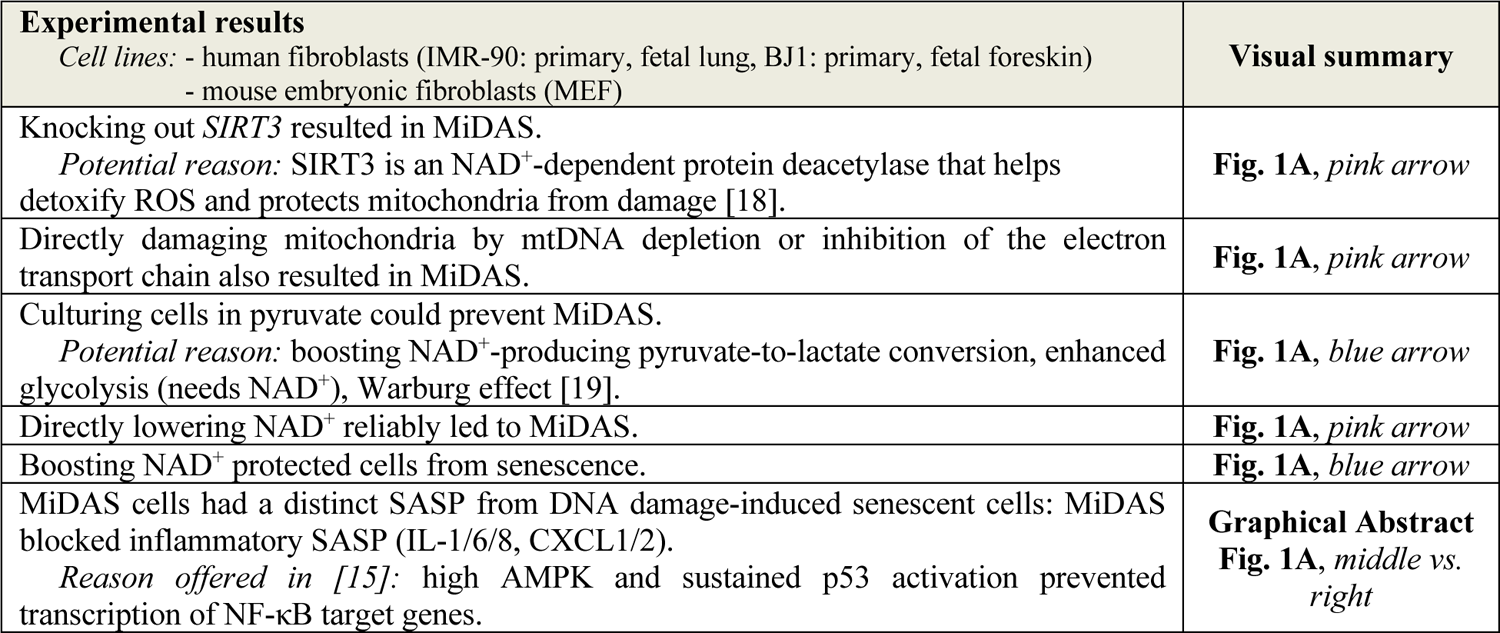
Experimental results in Wiley et. al. [17], detailing the role of SIRT3 and NAD^+^ in preventing MiDAS.

A paradoxical aspect of the relationship between mitochondrial dysfunction and DNA damage-induced senescence is that, on one hand, NF-κB-mediated SASP requires *functional* mitochondria with low AMPK and p53 [17,20,21]. On the other hand, mitochondria with low membrane potential (ΔΨ_M_) and a hyperfused morphology are nevertheless observed during DNA damage-induced senescence [22]. A potential solution to the paradox is the observation that SASP appears late in senescence (after ∼ 6 days) [23], the time HMGA1 activity is increased [24]. HMGA1 was shown to induce nicotinamide phosphoribosyl-transferase (NAMPT) [20], a rate-limiting enzyme of the NAD^+^ salvage pathway responsible for most NAD^+^ production in mammals [25]. This chain culminates in a reversal of mitochondrial dysfunction, increasing the translation of SASP proteins. In parallel, downregulation of p53 in deep senescence [26] reactivates NF-κB, which drives the pro-inflammatory SASP (Fig. 1A, *right*). The intriguing implication is that MiDAS may not be an inherently distinct cell fate from DNA damage-induced deep senescence (Fig. 1A, *blue dashed arrow*). Rather, MiDAS may be an early stage that cells with intact mitochondrial DNA and protective enzymes such as SIRT3 recover from as they progress to deep senescence. In contrast, cells in which MiDAS is triggered by unrepairable or ongoing mitochondrial damage (e.g., loss of mtDNA or SIRT3) can never progress past the MiDAS stage and maintain a distinct, NF-κB-independent SASP [17].

Though hyperfusion of mitochondria occurs during senescence, it also occurs in healthy cells. Hyperfusion in proliferating cells is associated with *increased* ΔΨ_M_ and ATP generation [27], fueling DNA synthesis (Fig. 1B) [28]. A similar ΔΨ_M_ increase due to hyperfusion is induced by AMPK during ATP shortage (Fig. 1C); ATP production is increased, lowering AMPK and subsequently reducing hyperfusion in a negative feedback loop. In both cases, the hyperfused state is short-lived and protected from ROS. In damaged-induced senescence, hyperfused mitochondria are typically dysfunctional; ATP levels do not recover, leading to chronic AMPK activation [13,29,30]. Just as mitochondrial hyperfusion and ΔΨ_M_ increase in DNA synthesis, mitochondrial shape and ΔΨ_M_ are regulated in mitosis and apoptosis. Mitochondria are fragmented during mitosis without significant ΔΨ_M_ loss (Fig. 1D) [28,31], while apoptotic fragmentation aids outer membrane permeabilization and crashes the ΔΨ_M_ (Fig. 1E) [32]. None of these processes directly explain how the homeostatic fusion/fission dynamics are disrupted in senescence, or why the hyperfused network gets stuck in a dysfunctional state.

Published computational models of mitochondrial function and/or dysfunction generally address a single aspect of the multifaceted processes described above [33]. One group of models focuses on ROS production [34–37], ROS propagation [38], or DNA damage-induced NAD^+^ loss [39], but do not account for the role of mitochondrial shape dynamics. Another group of models captures the homeostasis of mitochondrial morphology in the absence of acute energy stress and/or increased ROS [40–42]. The two exceptions we are aware of are the Pezze et. al. model [29], the first to center mitochondrial dysfunction as a driver of damage-induced senescence and the original inspiration of this study. The other model by Hoffman et. al. [43], while specific to *C. elegans*, contains a key feedback loop between mitochondrial NAD^+^ generation and SIRT activation, leading to antioxidant gene transcription. This feedback has a similar role to that of NAD^+^ in SIRT3-mediated protection from mitochondrial ROS [17]. While both models link mitochondrial damage to senescence, they do not connect them to changes in mitochondrial shape, and thus cannot reproduce the context-dependent relationship between ΔΨ_M_, ATP production and mitochondrial morphology in proliferating versus senescent cells.

Here we offer a mechanistic regulatory network model that brings together mitochondrial energy production and its failure, involving dynamic changes in mitochondrial morphology leading to MiDAS. We focus on the interdependence of mitochondrial morphology, renewal by mitophagy, and ROS generation. We *hypothesize* that when mitochondrial ROS increases and NAD^+^ levels drop due to ETC damage, this system can lock itself into MiDAS via the following positive feedback: the hyperfused network no longer restores ATP, AMPK sustains hyperfusion, which in turn continues to produce ROS [44] – unprotected by SIRT3 due to low NAD^+^ and exempt from mitophagy [44] – further compromising ATP production (**Fig 1A**, *middle*). Leveraging this hypothesis and key insights from Pezze et. al. [29], we propose a regulatory network model which integrates the role of NAD^+^ in preventing MiDAS stabilization. We combine this novel mechanism with a previously published Boolean network model of cell cycle regulation [45] into a mechanistic model of *SIRT3* deletion- or ROS-driven MiDAS.

Boolean network modeling focuses on the combinatorial logic by which regulatory influences converge to control molecules, pathways, or cellular processes [46,47]. In these models, the activity of each regulatory molecule is approximated as ON (expressed and active) or OFF (not expressed or inactive), and each node is updated in time via a logic gate or table that specifies its response to every combination of its input states. Thus, Boolean modeling sacrifices precision in the concentration- and time-dependence of regulatory processes in favor of scalability. This tradeoff is both necessary and justified. The kinetic parameters driving the interactions of large regulatory networks (> 100 nodes) are typically unknown. Moreover, a common observation of kinetic studies on small circuits is their remarkable lack of sensitivity to internal parameters and/or modeling details [48–52]; justifying a Boolean approximation [53,54].

The principal predictive feature – and means of validation – of Boolean models is the equilibrium dynamics emergent from the network’s logical regulatory rules [47]. All Boolean models have a set of steady states and/or limit cycles, also referred to as attractors. Every dynamically evolving system eventually converges into one of these states (or sets of states) and, unperturbed, will stay there indefinitely. The canonical expectation is that the attractors of Boolean regulatory network models represent biologically meaningful phenotypes [55]. Furthermore, we expect that both changes in the model cell’s environmental inputs and perturbations of its internal nodes result in biologically meaningful signal transduction and phenotype transitions, in accordance with experimental data. Indeed, the Boolean network model of MiDAS presented here reproduces the connection between mitochondrial dynamics and cell cycle progression, apoptosis, glucose starvation and finally, *SIRT3* deletion- or ROS-driven MiDAS along with its rescue by external pyruvate. We then use this model to generate testable predictions about MiDAS in healthy versus cancer-related mutant cells.

### Computational Methods

#### I. Boolean model building

To build the Boolean model we extended our previously published cell cycle and apoptosis network [45] with pathways that control glycolysis, mitochondrial ATP production, mitochondrial morphology, and DNA damage response (a circuit focused on p53/p21-mediated cell cycle arrest). The model synthesizes qualitative experimental data from 466 papers into a 134-node Boolean network in which all links and regulatory functions are experimentally justified (**SM Table 1**). For details of our model construction approach, the use of synchronous vs. asynchronous update, automatic module isolation from the larger network, storing our model in *Dynamically Modular Model Specification* (.*dmms*) format and exporting it to SBML-qual or *BooleanNet* formats (**SM Files 1-3**), see *Supplementary Methods* (*Suppl. Mat. 1*) and STAR Methods in [56].

#### II. Model availability

The Boolean model presented here was deposited in the BioModels online repository (MODEL2312140001). In addition, model files in *SBML* format (BioModels; used by *GinSim* [57] and *The Cell Collective* [58]), *dmms* format (used by our own discrete-state modeling software *dynmod*), *BooleanNet* format (used by the BooleanNet Python library [59]), and an editable network visualization in *gml* format (read by yED [60]) are included as **SM Files 1-4**.

#### III. Dynmod Boolean modeling software

Boolean simulations and analysis were performed with the discrete-state modeling software *dynmod*, developed in Haskell by the Regan lab (justification for using in-house software detailed in *Suppl. Mat. 1a*). All original code is available on GitHub (https://github.com/Ravasz-Regan-Group/dynmod); instructions to install Haskell and compile *dynmod* are included in *Suppl. Mat. 1b*. A guide to reproduce our modeling results is included in *Suppl. Mat. 1c-f*, aided by **SM Files 5-12, 15**. Additional information to run the simulations is available upon request from E.R.R; technical help with installing, using and/or extending *dynmod* is available upon request from P.L.R. To reproduce our main findings outside of *dynmod,* Jupyter Notebook using BooleanNet [59]: https://github.com/deriteidavid/midas_boolean_model).

Rather than replicating existing functionality of widely used Boolean modeling software (e.g., graphical user interface to create, document and simulate models, extensive update options, efficient attractor detection/ visualization, network control) [57–59,61,62], *dynmod* focuses on *automating the detection, evaluation and analysis of complex biological phenotype-combinations* represented by the attractors and time-series of our models (*Suppl. Mat. 1e-f*). Briefly, our code can: **a)** use user-defined signatures attached to regulatory switches to automatically map each attractor to a combinatorial phenotype profile (e.g., quiescent, alive, MiDAS); **b)** visualize and filter attractors of interest via their phenotype profiles, organizing them within a coordinate system of independent environmental input-combinations; **c)** set up simulations by specifying the initial environment and phenotype-combination (rather than each node state); **d)** collect phenotype statistics on large cell ensembles (independent simulation runs) in non-saturating environments and/or non-saturating perturbations (e.g., 10% Trail, 50% AMPK inhibition); and **e)** use metadata from *dmms* files to generate a formatted table with all biological documentation (**SM Table 1**).

#### IV. Attractor detection

To test whether all model attractors represent biologically relevant, distinct cell states and are robust to Boolean update, we perform both synchronous and asynchronous attractor search using *dynmod* and *AEON* [63], respectively.

a. *Synchronous attractors. Dynmod* uses synchronous update to find stable phenotypes and/or oscillations (attractors) via a stochastic sampling procedure [64] detailed in [65,45,66,56]. We find attractors by running noisy time-courses of length *T* with noise *p* from *N* different random initial conditions for each unique combination of environmental inputs. For each observed state/step along these time-courses, the synchronous attractor basin is determined. Finally, we test for the convergence of this sampling process by repeating it with increasing N and T, collating all detected attractors (*Suppl. Mat. 1d*).
b. *Asynchronous complex attractors. AEON* (https://github.com/sybila/biodivine-aeon-py) [63] uses general asynchronous update, but in place of random sampling it uses symbolic computation to detect all bottom strongly connected components (BSCCs) in the model’s state transition graph [67]. These subspaces trap the model’s dynamics and thus represent asynchronous attractors. *AEON* detected 78 fixed points (matching *dynmod*) and 16 complex attractors corresponding to the 16 non-cell cycle synchronous oscillations involving PI3K (*Suppl. Mat. 7;* **SM Files 7,8**; run AEON on our model: https://github.com/deriteidavid/midas_boolean_model).

#### V. Running simulations with dynmod

To run simulations, *dynmod* parses a user-generated experiment file (*.vex* format). Precise use of each command is described in *MiDAS Main_Figures.vex* (**SM File 10**) and *MiDAS SM_Figures.vex* (**SM File 11**), helping readers reproduce nearly all data figures (exceptions: automatic module isolation, modeling network errors and complex attractors detected by AEON). Run with the **-e** command-line tag: dynmod MiDAS_Cell_Cycle_Arrests_Apoptosis.dmms -e MiDAS Main_Figures.vex dynmod MiDAS_Cell_Cycle_Arrests_Apoptosis.dmms -e MiDAS SM_Figures.vex

These files include instructions to simulate and visualize: **a)** synchronous time courses from a subset of cell states in a given initial environment, exposed to reversible changes in a single environmental signal; **b)** non-saturating environments where an input is stochastically tuned between 0 and 1; **c)** partial or full knockdown/hyper-activation of arbitrary sets of nodes [65,45,66,56] (justification & limitations in [56]); **d)** combine these in an arbitrary sequence of perturbations and environments defined in distinct time windows; **e)** average activity of all nodes and/or all module phenotypes in an ensemble of independent cells; **d)** bar charts of the activity of user-specified nodes and/or module phenotypes, averaged over an ensemble of independent rns and across each distinct time window of an experiment (*Suppl. Mat. 1e-g*).

#### VI. Texting the robustness of model behavior under random-order asynchronous update and random errors in node/link/Boolean gate activity

*Suppl. Mat. 7* includes variants of our Results figures generated using the biased asynchronous update described in [45], in which a small subset of nodes are updated at the start/end of an otherwise random update step. This data indicates that our findings are robust under asynchronous update (**SM Figs. 18-22**; reproduce with **SM File 14**). In addition, *Suppl. Mat. 8* includes variants of our main results generated with ensamples of mutant models. To generate these ensembles: a) *n*_Node_ ∈ {1,2,3} nodes were randomly locked on/off; b) *n*_Link_ ∈ {5,10,15} links were removed, or c) *n*_Gate_ ∈ {5,10,15} Boolean gates were randomly perturbed [56]. Mutant ensemble results indicate that our findings are robust with respect to minor errors or random perturbations in model architecture (**SM** Fig. 23).

#### VII. Previously introduced regulatory modules

Detailed descriptions of Growth factor signaling, Replication origin licensing, Restriction switch, Mitotic phase switch, Apoptotic switch and Cell cycle processes are detailed in [45,65].

## Results

### 1. Extending a Mitogen Signaling-Driven Cell Cycle and Apoptosis Model to Account for the Regulation of Mitochondrial Morphology

To examine the way MiDAS arises from the dynamics of the regulatory processes that govern mitochondrial energy production and the dynamics of fusion and fission, we used our previously published model of growth signaling, cell cycle, and apoptosis as a starting point [45]. This Boolean regulatory network model was shown to reproduce *PI3K* oscillations linked to cell cycle progression, capture the cell cycle-phase dependent roles of Plk1, and reproduce cell cycle errors in perturbed cells (e.g., mitotic catastrophe, aneuploidy, endoreduplication). Here we extend this model to include three new regulatory modules, capturing changes in metabolism, DNA damage signaling, and a mitochondrial module capturing energy production linked to the dynamics of mitochondrial morphology.

First, the ***Warburg*** module accounts for glycolysis and the Warburg effect – the increase in glucose uptake and lactate production characteristic of dividing cells (Fig. 2A). The module focuses on the regulatory logic behind the shift from moderate glycolysis producing pyruvate for the TCA cycle to increased glycolysis coupled with fermentation (pyruvate to lactate conversion) [19]. HIF-1α-mediated fermentation can support ATP production in the absence of oxidative phosphorylation, but its other critical function is to regenerate the cytosolic pool of NAD^+^ required for and depleted by glycolysis (Fig. 2A; *Glycolysis_H* ⊣ *NADp_c*; overridden by *Fermentation* → *NADp_c*). The Warburg effect is induced by HIF-1α and/or Myc-mediated up-regulation of glucose transporters and glycolytic enzymes; blocked by p53 [68].

**Figure 2.**
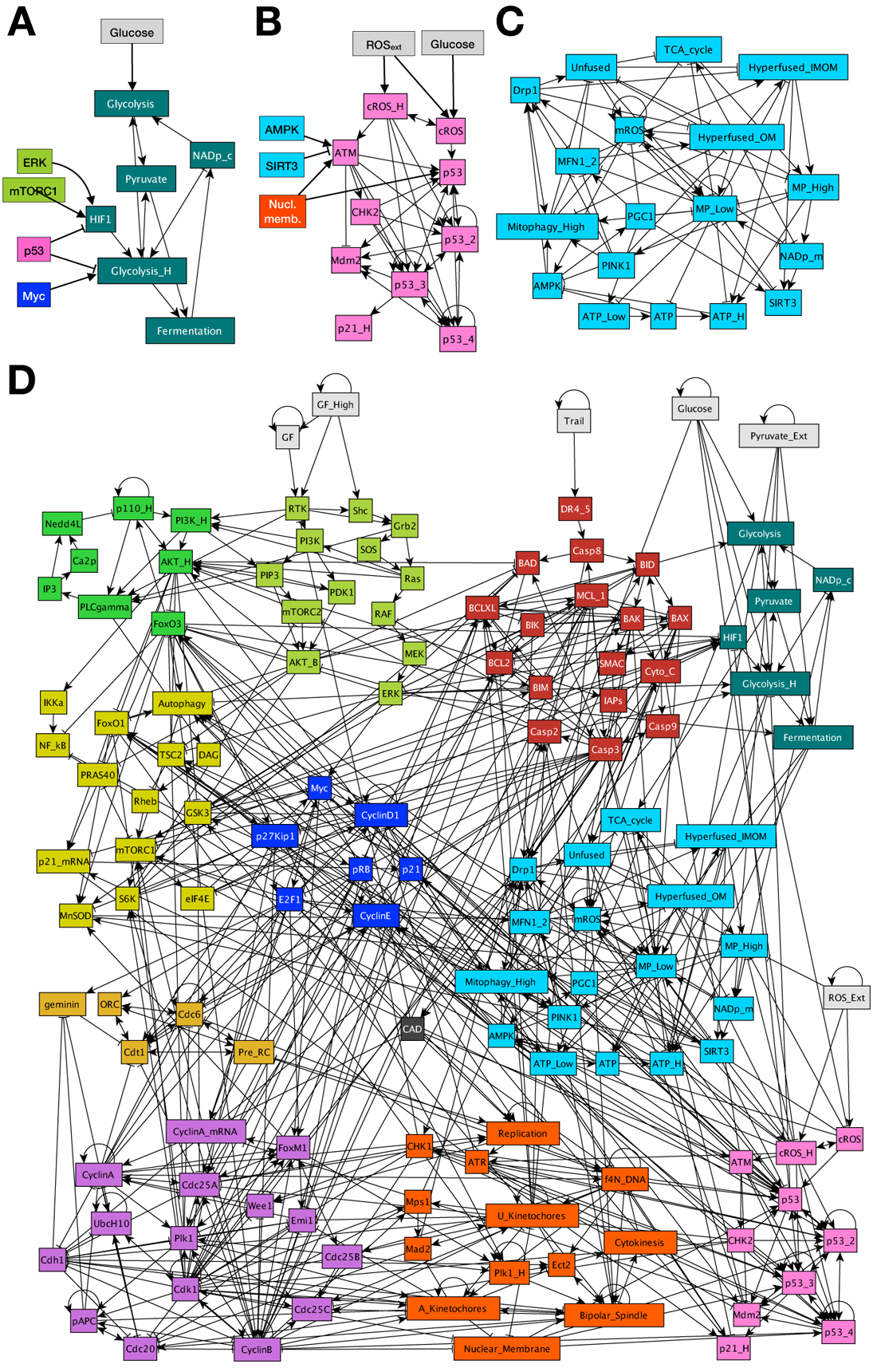
Boolean regulatory network model of cell cycle-linked mitochondrial energy production and shape dynamics. **A)** Warburg effect module tracking NAD^+^ use and regeneration by glycolysis and fermentation, with key influences from cell cycle control (ERK, mTROC1, Myc) and p53. **B)** DNA damage module showing ROS or ATM/Chk2-mediated p53 activation, its 4-node persistence tracker, negative feedback via Mdm2, downstream p21 induction, and key influences from the mitochondrial module (AMPK, SIRT3) and mitosis. **C)** Mitochondrial regulatory switch controlling energy production, mitochondrial NAD^+^ homeostasis, morphology, and ROS production. **D)** Modular network representation of our Boolean model. *Gray:* inputs representing environmental factors; *green*: Growth Signaling (*lime green*: basal AKT & MAPK, *bright green:* PI3K/AKT oscillations, *mustard*: NF-κB & autophagy (new), mTORC1, GSK3, FoxO1); *dark red:* Apoptotic Switch; *black:* Caspase-activated DNAse (CAD); *light brown:* Origin of Replication Licensing; *blue:* Restriction Switch; *purple:* Phase Switch; *dark orange:* cell cycle processes; *dark green:* Warburg effect; *teal:* mitochondrial switch; *pink*: DNA damage response; →: activation; ⊣: inhibition.

Second, the ***DNA Damage Signaling*** module is a representation of ROS-mediated p53 and p21 activation (Fig. 2B) [69]. The rules for p53 activation include phosphorylation by AMPK, ATM and ATR [16,70], as well as the lack of p53 transcriptional activity in mitosis [71]. Some of p53’s effects, such as *Cyclin B* repression, only occur after sustained or repeated p53 activation [72,73]. We account for this via four nodes that track increasing/sustained p53 activity under ongoing damage, and only link the 4th level (*p53_4* node) to these targets. In contrast, the effect of p53 on metabolic and cell cycle targets (*HIF1α, Myc, FoxO1/3, p21*) are immediate [74].

Third, the ***Mitochondria*** module contains a detailed map of all the relevant interactions that govern mitochondrial energy production coupled to the dynamics of their morphology (Fig. 2C) [75]. Briefly, the TCA cycle uses (and requires) pyruvate and mitochondrial NAD^+^, to generate NADH. The electron transport chain (ETC) utilizes the high energy electrons of NADH and regenerates NAD^+^ while generating the mitochondrial membrane potential (ΔΨM). Thus, through feeding the ETC the TCA cycle helps maintain normal mitochondrial membrane potential (ΔΨM), which is utilized by ATP synthase to synthesize ATP, and keep AMPK inactive. To capture the four qualitatively different regimes of ATP/AMP ratio in cells (deadly, survivable but with activated AMPK, normal, and high), we use three Boolean nodes for ATP (bottom row on Fig. 2C). Upon AMPK activation, the module captures the increase in mitophagy as well as *PGC1α* activation leading to increased mitochondrial biogenesis, ROS reduction, and increased transcription of the mitofusins MFN1/2, which start mitochondrial fusion at the outer membrane (*Hyperfused_OM* node) [76]. In mitochondria with normal ΔΨM, inner membranes also fuse (*Hyperfused_IMOM* node) to increase ATP production and lower AMPK. This negative feedback is homeostatic; hyperfusion only persists as long as ATP is in demand. Here, SIRT3 is a critical protector of ΔΨM as it lowers mitochondrial ROS and protects its ATP-generating potential [18]. In contrast, activation of the fission protein Drp1 results in a highly fragmented network seen in mitotic or apoptotic cells [31,77]. The mitochondrial module interacts with cell cycle control and apoptosis as summarized in Fig. 1. The complete regulatory network is shown on Fig. 1D, with detailed node and link justifications described in **SM Table 1** (2D layout in **SM File 4**; *gml* format).

### 2. Extended Model Reproduces Known Cell Cycle-Dependence of Mitochondrial Morphology as Well as G1 Arrest with Mitochondrial Fusion in Glucose-Starved Cells

First, we validated our extended model by comparing its dynamical states and responses to experimental literature. In the simulations discussed below we use synchronous update, in which all nodes are updated simultaneously in each time-step. Thus, these results are deterministic, allowing us to examine the effects of precisely timed perturbations; especially relevant in triggering responses known to change with cell cycle progression. Robustness of our results to asynchronous update is examined in *Suppl. Mat 7*, verifying our deterministic results. Robustness to errors in construction and/or minor mutations is included in *Suppl. Mat 8*.

As expected, our extended model has a quiescent attractor in the presence of limited mitogens, similar to its predecessor model [45]. Metabolically, this state is characterized by moderate glycolysis and pyruvate flux to the TCA cycle, normal ΔΨ_M_, and nominal relative ATP as well as mitochondrial NAD^+^ (Fig. 3A, *left*). Upon mitogen stimulation the model mimics cell cycle entry (Fig. 3A, *middle*), then cell cycle-exit upon a drop in mitogens to survival-sustaining levels (Fig. 3A, *middle*). Furthermore, our new model captures the cell cycle-dependent dynamics of mitochondrial morphology. These include mitochondrial hyperfusion at the G1/S transition (Fig. 3A, *red box & arrow, left*) [28], reset to a G0-like morphology in G2, and an unfused (fragmented) state in mitosis (Fig. 3A, *red box & arrow, right*) [31]. Note the enhanced protective SIRT3 activity during the reversible G1/S hyperfusion, *predicted* by the model as a requirement for keeping the hyperfused network healthy (*left-side red box / arrow*). As further validation, we reproduced experimentally documented behaviors inherited from the Sizek et. al. model [45], as well as five experiments that perturb mitochondrial morphology during cell cycle progression (**SM Tables 2-3** in *Suppl. Mat. 2* include a point-by-point comparison of model vs. experimental data [17,27,28,31,78–99], referencing results on **SM Figs. 1-6**).

**Figure 3.**
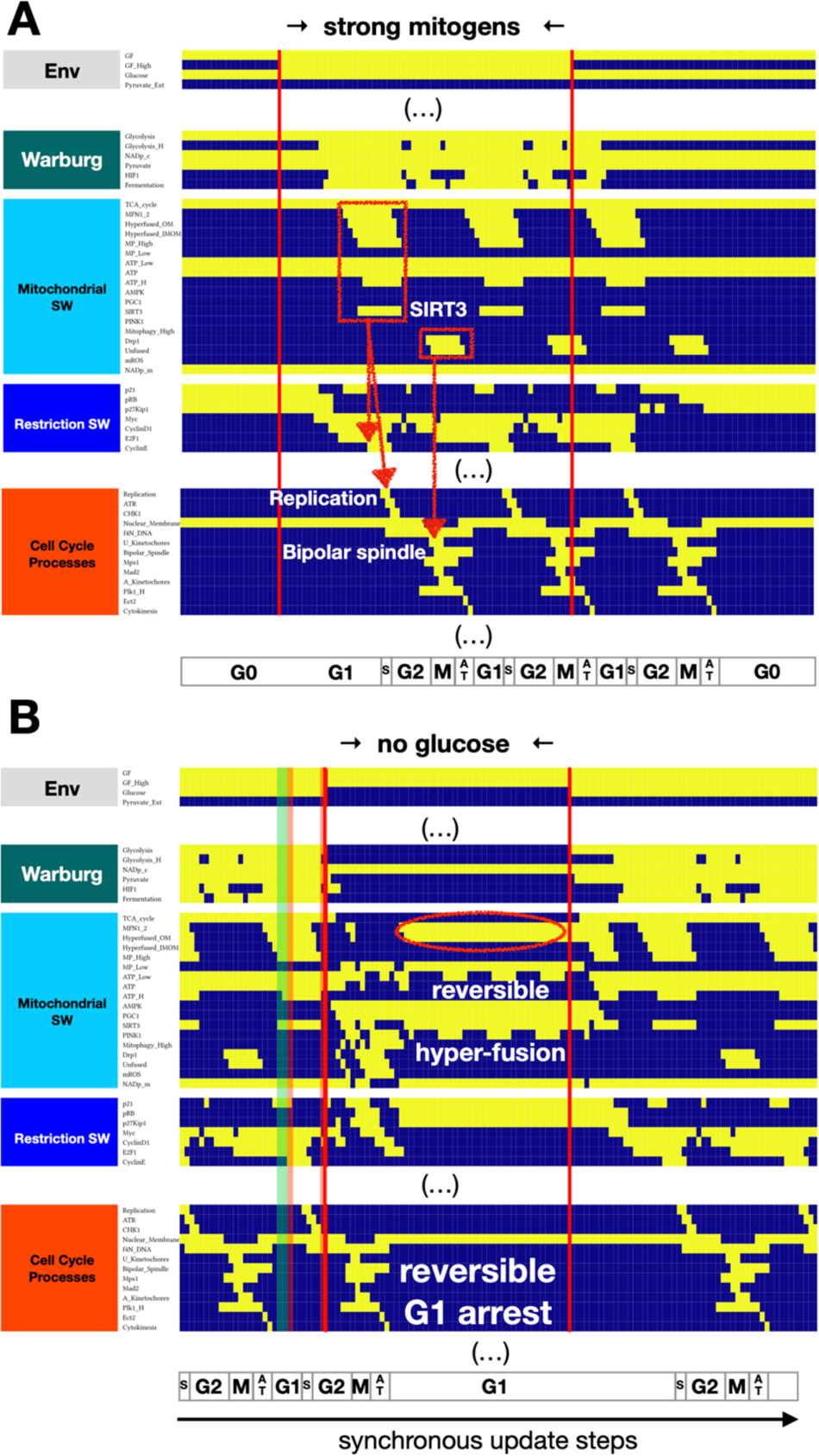
Model reproduces cell cycle-linked mitochondrial dynamics (hyperfusion at G1/S, fission in mitosis) and G1 arrest in response to glucose withdrawal. **A)** Dynamics of relevant regulatory molecule expression/activity during exposure of a quiescent cell to a saturating mitogenic signal for 60 update steps (full network dynamics in **SM** Fig. 1A). **B)** Dynamics of relevant regulatory molecule expression/ activity in a dividing cell responding to glucose withdrawal in early G1 for 50 update steps (full network dynamics in **SM** Fig. 1B). *X-axis:* time-steps annotated by cell cycle phase (*G0*: quiescence; *G1*: start of cell cycle entry; *S*: DNA synthesis; *G2*: growth phase 2; *M*: metaphase; *A T*: anaphase, telophase, cytokinesis); *y-axis:* nodes organized by regulatory modules; *yellow/dark blue:* ON/OFF; *vertical red lines:* start/end of signal; *red boxes with arrows:* morphological changes to mitochondria linked to cell cycle processes; *red oval*: reversibly hyperfused mitochondrial network; *shaded green band:* glucose withdrawal resulting in endore-duplication (**SM** Fig. 6A); *shaded red bands:* glucose withdrawal resulting in apoptosis (**SM** Fig. 6B**-C**).

Another physiological signal with a reversible effect on mitochondrial morphology is non-lethal glucose starvation, shown to induce a reversible G1 arrest [100]. This cell state is characterized by strong AMPK activation, increased mitochondrial fusion and decreased mitophagy [101]. Our model reproduces these observations, including a reversible G1 arrest accompanied by AMPK-driven PGC1α-activation, increased MFN1/2 expression, mitochondrial hyperfusion, and delayed cell cycle entry upon re-exposure to glucose (Fig. 3B). Based on the molecular mechanisms driving our model’s dynamics, we *predict* that these mitochondrial networks mainly fuse their outer membranes (*Hyperfused_OM* = ON; *Hyperfused_IMOM* = OFF), as inner membrane fusion requires normal ΔΨ_M_ [102]. Moreover, we do not expect a drastic drop in mitochondrial (or cytosolic) NAD^+^. Both primary processes consuming NAD^+^, the TCA cycle and glycolysis, are paused alongside reduced NAD^+^ generation by the ETC. If glucose withdrawal in our model occurs in late G2 or mitosis, we observe a G1 arrest (Fig. 3B, *non-shaded areas*). Outside this window, however, we *predict* two alternative cell fates shown in **SM** Fig. 6: G2 arrest followed by endo-reduplication in cells that experience glucose withdrawal in a G1 state already committed to DNA replication, and mitotic catastrophe whenever mitochondrial hyperfusion due to glucose withdrawal persists into mitosis (predicted mechanisms detailed in *Suppl. Mat. 3*, **SM Table 4**).

### 3. Mitochondrial Dysfunction-Induced Senescence (MiDAS) is locked in by positive feedback, maintaining mitochondrial hyperfusion with low ΔΨ_M_ and high ROS

To test whether our model can reproduce *irreversible* MiDAS, we first examined the steady states of the isolated Mitochondrial Module (automatic module isolation method in [56,65]). As Figure 4A indicates, this module acts as a three-state switch (has three distinct steady states). First, the ***healthy*** mitochondrial state (*top*) has no hyperfusion or excess fission. It has a healthy TCA cycle maintaining ΔΨ_M_, ATP, and mitochondrial NAD^+^ within their nominal range, keeping AMPK off, and mitophagy in its basal range (*Mitophagy_High* node is OFF). Second, mitochondria can get stuck in a ***fragmented*** state with low ΔΨ_M_ and high mitochondrial ROS, leading to low ATP and active AMPK. As AMPK can phosphorylate and activate Drp1 [103], this state is stabilized by a combination of Drp1-induced fission and a dynamic balance between high mitophagy and PGC-1α-induced mitochondrial biogenesis [76]. While PGC-1α induces MFN1/2, their ability to counterbalance fission and restore a healthy state is blocked by the molecular machinery of mitophagy [104,105].

**Figure 4.**
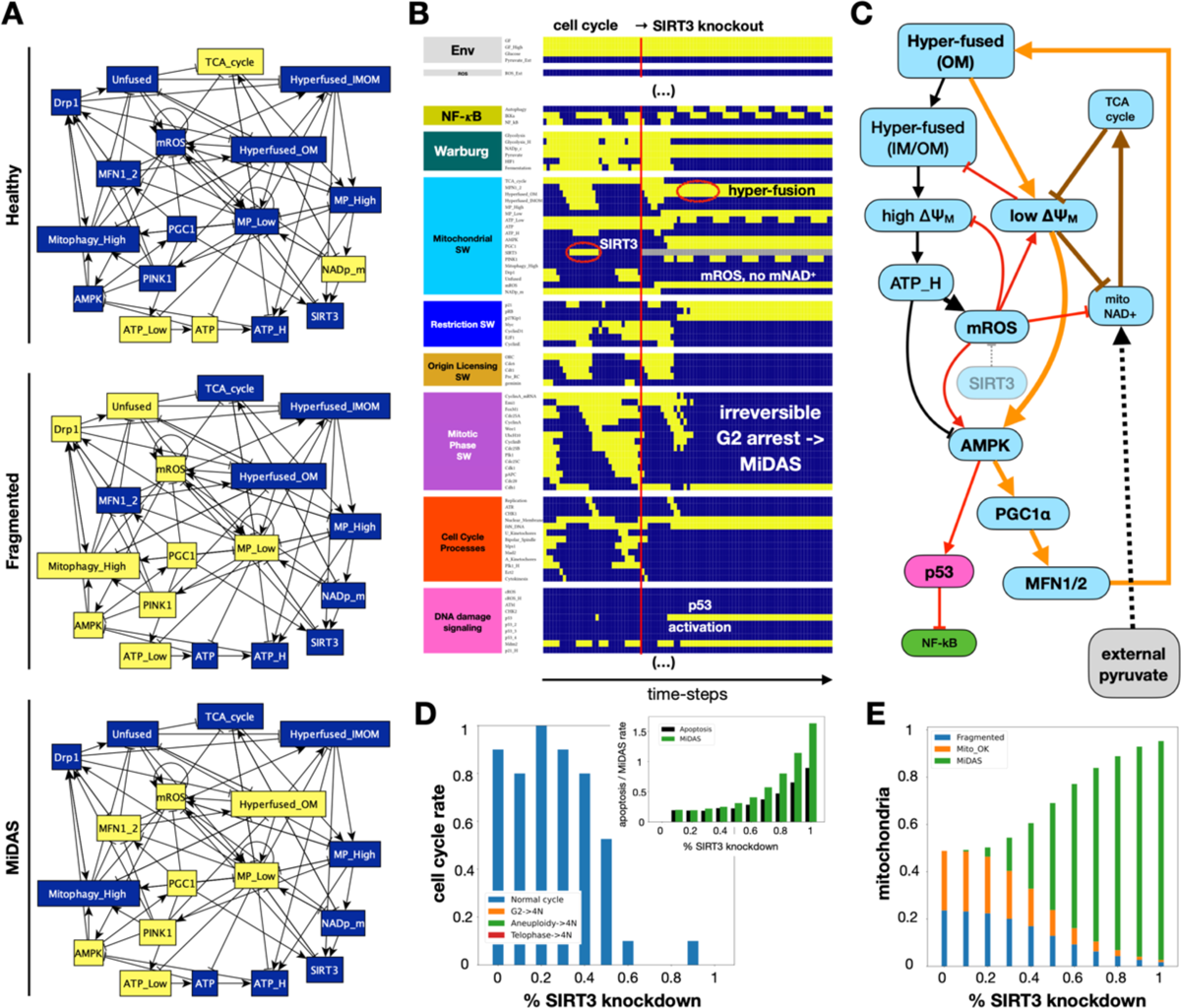
Model reproduces MiDAS in response to SIRT3 knockdown. **A)** Three stable states of the isolated mitochondrial module. *Yellow/dark blue background:* ON/OFF state of nodes in each stable state. **B)** Dynamics of relevant regulatory molecule expression/activity in a dividing cell responding to full *SIRT3* knockout in early G1 (full network dynamics in **SM** Fig. 7A). *X-axis:* time-steps; *y-axis:* nodes organized by regulatory modules; *yellow/dark blue:* ON/OFF; *vertical red line:* start of *SIRT3* knockout; *red ovals*: SIRT3 expression during cell cycle (*left*) / hyperfused mitochondrial network in MiDAS (*right*); *labels:* relevant molecular or phenotypic changes in response to *SIRT3* knockout. **C)** Circuit responsible for locking in MiDAS following S-phase in *SIRT3*-null cells. *Light blue/pink/green nodes*: molecules from the mitochondrial/DNA damage/growth signaling modules. *Black links:* triggering event of MiDAS is a healthy but hyperfused network of mitochondria, responsible for high ATP generation during S phase but also mitochondrial-derived ROS (normally mitigated by SIRT3). *Red links:* in the absence of SIRT3, excess mitochondrial ROS damages the ETC, lowers ΔΨ_M_ and consequently the mitochondrial NAD^+^/NADH ratio, and helps activate AMPK. At this point, two positive feedback loops engage to keep the mitochondrial network dysfunctional. *Orange links:* first positive feedback loop involving AMPK-mediated hyperfusion of the outer membrane (AMPK → PGC1α → MFN1/2), no longer paired with inner membrane fusion and efficient ATP generation but still blocking its own disassembly (mitophagy) and subsequent renewal, which in turn keeps ATP low and AMPK active. *Brown links:* second positive feedback loop involving mitochondrial ROS-mediated damage to ETC proteins and mDNA, which keeps mitochondrial NAD^+^ low, compromises the TCA cycle, and further lowers ΔΨ_M_. *Black dashed link:* excess pyruvate can interrupt these feedback loops by restoring NAD^+^ levels. **D-E)** Response of cells dividing in 95% saturating growth stimuli to increasing levels of *SIRT3* knockdown. **D)** rate of normal cell cycle completion (*blue*) vs. G2 → G1 reset (*orange*, not observed), aberrant mitosis (*green*, not observed), or failed cytokinesis followed by genome duplication (*red*, not observed), relative to wild-type cell cycle (25 steps), shown as stacked bar charts. *Inset:* rate of apoptosis (*black*) or MiDAS (green) relative to wild-type cell cycle (25 steps) in ensembles where individual cell simulations are terminated at apoptosis, MiDAS entry, or 250 update steps. ***E*)** Fraction of time cells in an ensemble display normal mitochondria (*orange*), MiDAS (*green*), or fragmented mitochondria (*blue*) in simulations terminated at apoptosis or 250 update steps. *Initial state for sampling:* cycling cell in high glucose and no external pyruvate, ROS, or Trail; *sample size:* ≥ 2000 cells; *stop at:* apoptosis or MiDAS (D), or at apoptosis (E); *maximum length of single-cell tracks:* 250 update steps (10 wild-type cell cycle lengths); *total sampled cell time:* 500,000 steps; *update:* synchronous.

Third is a stable state that matches the experimentally observed signatures of ***MiDAS*** (*bottom*) [17]. It is characterized by low ΔΨ_M_, low ATP and active AMPK, which in turn keeps PGC-1α active. Downstream of PGC-1α, increased MFN1/2 keeps the outer mitochondrial membranes fused in a misplaced attempt to boost ATP levels via hyperfusion (as seen in G1/S). Here, however, low ΔΨ_M_ keeps Opa1 from fusing the inner membranes [106], rendering the hyperfused network dysfunctional. This positive feedback from low ΔΨ_M_ to AMPK to dysfunctional hyperfusion (and thus low ΔΨ_M_) is stabilized by the fact that hyperfused mitochondria are refractory to mitophagy [101], compromising their usual renewal. Unlike the hyperfused state depicted in Fig. 3B, the model’s MiDAS state is irreversible. A key difference between these two scenarios is the absence of SIRT3 activity to mitigate mitochondrial ROS. While PGC1α can induce *SIRT3* expression [18], its enzyme activity depends on NAD^+^, which in turn requires a functioning ETC. Our model assumes that the combination of low ΔΨ_M_ and mitochondrial ROS – which damages ETC components and mDNA – hinder the regeneration of the mitochondrial NAD^+^ pool [107,108].

When incorporated into the rest of the network, the *Mitochondrial Module* must work in concert with the other modules in a way that preserves its phenotypic states and reproduces the experimentally observed commitment to MiDAS. To test this, we first knocked down *SIRT3* in a model cell undergoing rapid proliferation, a scenario shown to induce MiDAS [17]. Indeed, Figure 4B shows permanent cell cycle arrest following *SIRT3* knockout, along with low mitochondrial NAD^+^/NADH ratio, active AMPK and p53, and no NF-κB activity – the reported hallmarks of MiDAS [17]. In addition to reproducing observed features, our model offers a mechanistic explanation for the transition (Fig. 4C), along with several testable predictions related to *SIRT3* knockdown-induced MiDAS (**SM** Fig. 7). Namely, we *predict* that: **a)** *SIRT3*-null MiDAS cells have a hyperfused, non-functional mitochondrial network with low ΔΨ_M_ and excessive ROS production; a phenotype also observed in damage-induced senescence [22]. **b)** p53 dynamics in MiDAS is not oscillatory, as AMPK is not expected to be sensitive to inhibition by Wip1 [109] (thus not explicitly modeled). **c)** SIRT3 knockout cells arrest in MiDAS from early G2 with 4N DNA content, and **d)** G0 cells with access to glucose are protected from *SIRT3* knockout-induced MiDAS. **e)** In contrast, cells held in G0 by the lack of glucose with reversibly hyperfused mitochondria do enter MiDAS in response to *SIRT3* knockdown, as their ROS production increases and NAD^+^ levels plummet.

Further validation detailed in *Suppl. Mat. 4* shows that our model can reproduce MiDAS triggered by ETC inhibitors (a response prevented by pyruvate) [15] and *predicts* that pyruvate can rescue wild-type cells from MiDAS (early senescence only). Furthermore, we have found that perturbations disrupting the cell cycle at specific points can also trigger MiDAS (**SM Figs. 8-10**). The *predicted* cause of MiDAS is a sub-lethal mitochondrial membrane permeabilization that lowers the ΔΨ_M_ without activating executioner caspases (which would trigger apoptosis) – at a time when the mitochondria are hyperfused. Given these conditions, cells react to low ΔΨ_M_ with AMPK activation and lock in MiDAS.

### 4. External ROS exposure causes MiDAS, preventable by excess pyruvate

Given the central role of ROS in locking in MiDAS in *SIRT3*-null or ETC-deficient cells, we next modeled the effects of extracellular ROS (**SM** Fig. 11). To this end we linked an *ROS_Ext* environmental input to high intracellular ROS and added a direct inhibitory effect on high ΔΨ_M_. High internal ROS, in turn, activates ATM/ATR, AMPK, p53 and autophagy, but also promotes mitochondrial dysfunction by interfering with the TCA cycle, lowering ΔΨ_M_, and increasing mitochondrial ROS. Exposing our dividing cells to prolonged external ROS reliably triggered MiDAS (Fig. 5, full version on **SM 12**). Cells exposed to ROS in G1 arrested with 2N DNA content (Fig. 5A), while G2 cells lost their G2/M associated cyclin expression following sustained (non-oscillatory) p53 accumulation and underwent irreversible G2 arrest into MiDAS (Fig. 5B). Probing the relationship between ROS exposure, proliferation, and the roles of NAD^+^, pyruvate and SIRT3 (Fig. 5C**-D**; *Suppl. Mat. 5*; **SM Figs. 12-14**), we predicted that quiescent but not glucose-starved cells were protected from ROS-induced MiDAS, and that pyruvate or SIRT3 hyper-activation could prevent and even reverse MiDAS in cells capable of restoring ETC function. A similar result could be achieved by boosting NAD^+^ levels **–** previously shown to reverse MiDAS [17] and support pro-inflammatory SASP by restoring mitochondrial function in deep senescence [20,21].

**Figure 5.**
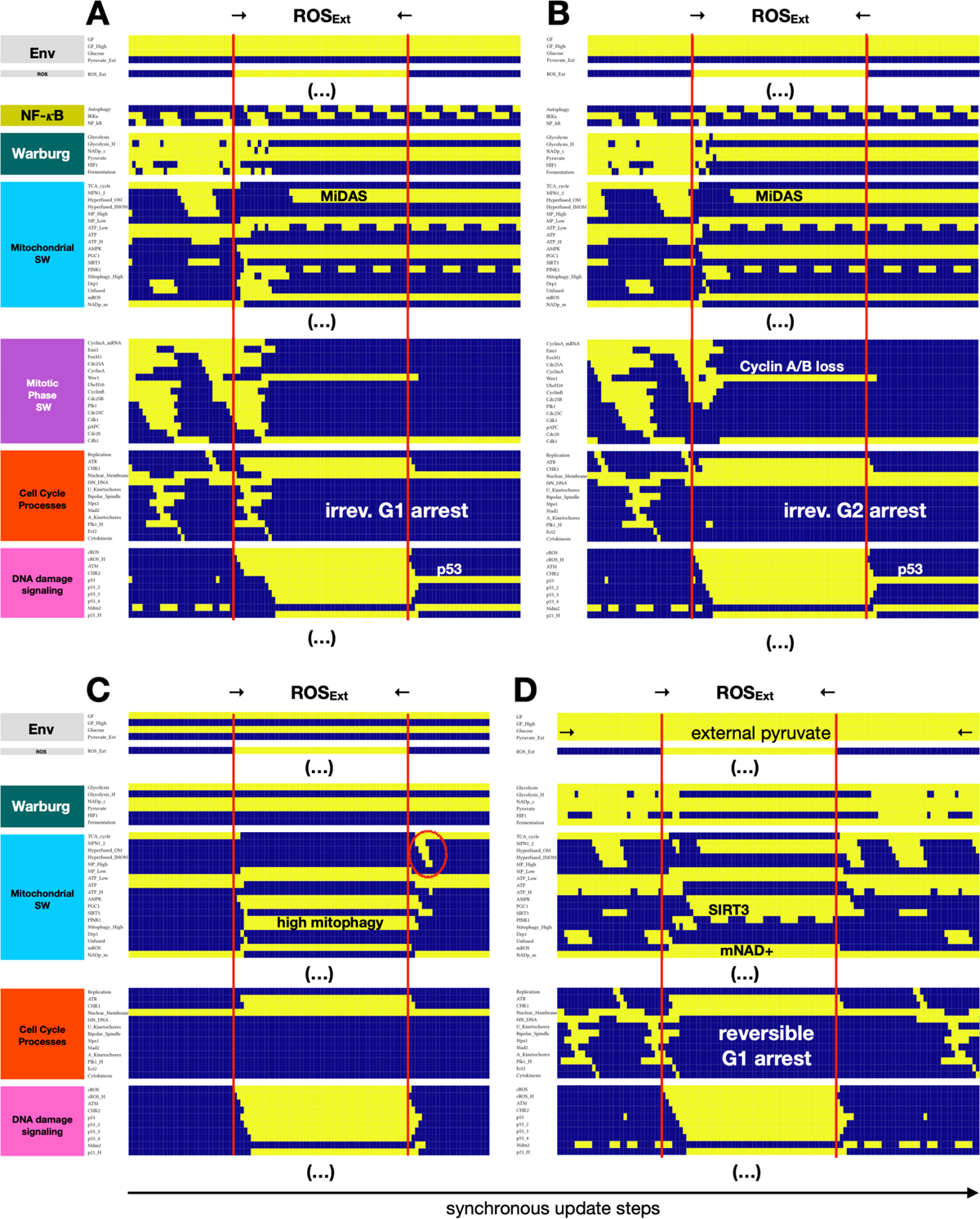
Model reproduces ROS-induced MiDAS in cycling cells and predicts protection from MiDAS in external pyruvate. **A-B)** Dynamics of relevant regulatory molecule expression/activity during exposure of a cycling cell to external ROS for 50 update steps in (A) prometaphase, leading to cytokinesis followed by MiDAS (2N DNA), vs. (B) early G2, leading to irreversible G2 arrest and MiDAS with 4N DNA. **C)** Dynamics of relevant regulatory molecule expression/activity during exposure of a quiescent cell to external ROS for 50 update steps, leading to high mitophagy and short-term hyperfusion to restore ATP levels. **D)** Dynamics of relevant regulatory molecule expression/activity during exposure of a cycling cell exposed to saturating levels of external pyruvate to external ROS near the SAC, for 50 update steps, leading to reversible G1 arrest (cell cycle entry after external ROS is removed). *X-axis:* time-steps; *y-axis:* nodes organized by regulatory modules; *yellow/dark blue:* ON/OFF; *vertical red lines:* start/end of ROS exposure; *red oval on (C):* brief, homeostatic hyperfusion involving both outer and inner mitochondrial membranes; *white/black labels:* relevant molecular changes or outcomes; *full network dynamics:* **SM** Fig. 12.

### 5. The stability of MiDAS depends on the combinatorial influence of mitogens, glucose, ROS, and pyruvate

To examine the stability of MiDAS to perturbation in the cell’s microenvironment, we surveyed our model’s stable phenotypes (attractors) in every combination of its inputs. Here we focus on a subset of attractors; namely we visualize those representing live diploid cells exposed to glucose, survival signals (GF = ON) and no Trail (Fig. 6A). These cell states are organized on a coordinate system of the other remaining inputs: low vs. high growth factor (*x* axis), ROS (*y* axis) and external pyruvate (*left/right*), each attractor represented by a barcode and a small visual indicating mitochondrial shape. For example, a quiescent cell exposed to low growth signals, no external ROS, and no pyruvate (*left panel, bottom left pair*) is marked by a barcode showing the state of its key molecular switches, including normal mitochondria, a cell cycle machinery consistent with quiescence, and inactive apoptotic machinery. The *green-outlined cell* to the right of this barcode summarizes the following phenotype: a healthy quiescent cell with dynamic mitochondria. In contrast, the cell cycle attractor (*left panel, bottom right, red outline*) has a wide barcode showing oscillatory PI3K (*green*), dynamic mitochondria (*light blue*), a cyclic toggle of the Phase Switch (G1→G2→M, *purple*) and a sequence of cell cycle events (*orange*). We summarized this with a mitotic spindle icon (*red-outlined cell*).

**Figure 6.**
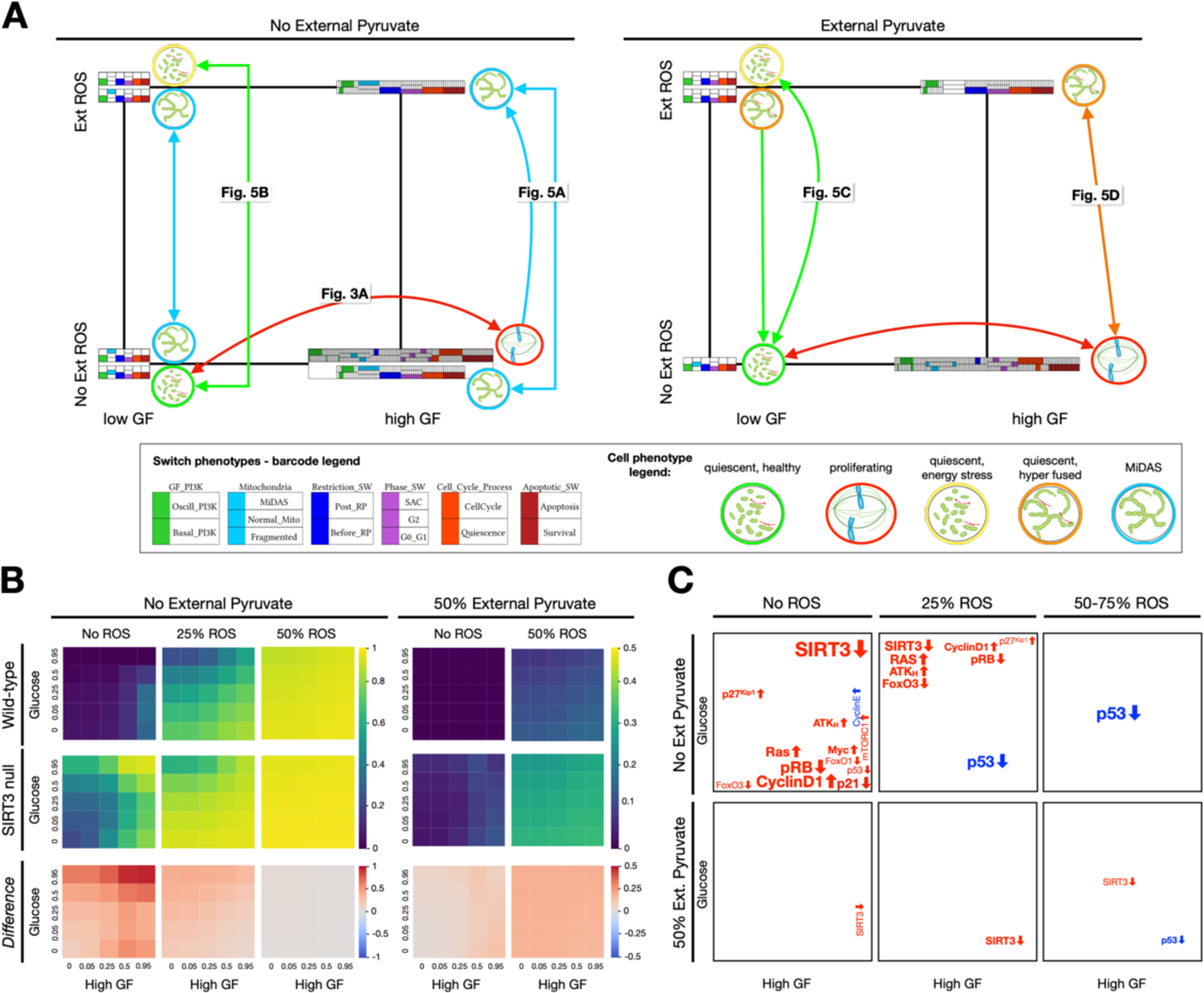
Model produces a heterogeneous mix of cell phenotypes modulated by mitogens, ROS, and pyruvate exposure, in which the prevalence of MiDAS increased by most oncogenic mutations. **A)** Summary of model cell states detected in every combination of no/low/high growth-factor (*x axis*), absence/presence of external ROS (*y axis*) and absence/presence of external pyruvate (*left/right*). Apoptotic and tetraploid quiescent cell states were omitted for clarity (see **SM** Fig. 15**-16** for more complete maps; **SM Files 7-8** contain all synchronous attractors). Barcodes representing each attractor were derived by comparing the expression of nodes in relevant modules to predetermined molecular signatures (e.g., apoptosis vs. survival), encoded in the *dmms* model file (barcode legend, *bottom left*). Oscillatory phenotypes have expanded barcodes that mark the transitions their regulatory switches undergo during the cycle (*high GF areas, no ROS*). Visual summaries of overall cell states indicate the shape and dynamics of the mitochondrial network, distinguish quiescent vs. cycling cells, as well as normal vs. dysfunctional mitochondria (cell phenotype legend, *bottom right*). *Figure labels:* time-courses with molecule-level view of state. *State transition arrows: light blue:* irreversible transition to MiDAS; *green:* reversible transitions from quiescence with healthy mitochondria & transition to it; *orange:* transitions between energy stressed non-MiDAS states; *red:* reversible cell cycle arrest upon glucose withdrawal. **B)** Fraction of time cells spend in a MiDAS state at varying levels of mitogen stimulation (*x* axis) and glucose (*y* axis) with no (0%), mild (5%) or strong (50%) ROS exposure, in the absence (left 3 columns) vs. presence of 50% external pyruvate (right 2 column). *Top/middle row:* wild-type/*SIRT3* null cells; *bottom row:* increase in the fraction of time spent in MiDAS in *SIRT3*-null cells. *Length of time-window for continuous runs:* 100 steps (10 wild-type cell cycle lengths); *total sampled live cell time:* 500,000 steps; synchronous update; *initial state for sampling runs:* cycling cell in high glucose and no external pyruvate, ROS, or *Trail*. **C)** Summary of changes to the frequency of MiDAS across 4 environmental dimensions for 13 oncogenic and 11 tumor-suppressive mutations [110] (data in **SM File 13;** hyper-activated: PI3K_H_, Ras, RAF, mTORC1, AKT_H_, Myc, HIF1, MEK, Cyclin E, Cyclin D1, ERK, p21_H_, p27^Kip1^; knocked out: p53, ATM, ATR, RB, p21, TSC2, FoxO1, FoxO3, Caspase 8, Caspase 9). *Top vs. bottom rows:* no external pyruvate vs. 50% of saturating external pyruvate for no ROS vs. mild (5%) vs. strong (50% to 95%) external ROS. *Gene label position:* area of growth factor/glucose environment map where the mutation’s effects are strongest; *red/blue:* mutation increases/decreases the frequency of MiDAS; *up/down arrows after gene:* knockdown/hyper-activation; *font size:* effect strength ∼10% / font increment. *Image credits:* apoptotic cell: https://en.wikiversity.org/wiki/WikiJournal_of_Medicine/Cell_disassembly_during_apoptosis; mitochondrial networks: PMID 25847815; mitotic spindle: https://en.wikipedia.org/wiki/Spindle_apparatus.

As the left panel of Fig. 6A indicates, in conditions with sufficient glucose and no external ROS or pyruvate our model predicts growth signal-dependent quiescence vs. cell cycle (*red arrow*). In addition, MiDAS cells are stable in all these environments, as expected from a non-reversible cell state (*blue-bordered cells with static hyperfused mitochondria*). ROS exposure in the presence of strong growth signals always results in MiDAS (*light-blue directed arrow*). Low growth signals, in contrast, protect quiescent cells from ROS-induced MIDAS; instead, these enter an energy-stressed state reversible upon relief from ROS (*green arrow*). While the MiDAS phenotype is stable across all environments without pyruvate, its presence drastically changes the fate of our model cells (Fig. 6A, *right panel*). Cell cycle entry and exit remain unaltered (*red arrow*), but MiDAS disappears in favor of a reversible hyperfused state. Moreover, loss of pyruvate only triggers MiDAS from this hyperfused state in the presence of ROS (transitions between orange to blue states between panels; *arrows not shown*). Finally, the absence of glucose further promotes the non-MiDAS hyperfused state, as detailed in *Suppl. Mat. 5* (**SM Figs. 15, 16**).

### 6. Oncogene hyperactivation and tumor suppressor loss (except p53) tend to increase MiDAS in the absence of external pyruvate

To probe the effect of mutations such as *SIRT3* loss on the full phenotype-repertoire of our model, we measured the average time spent in a MiDAS state during a 250-step (10 cell cycle length) window upon entering a new environment, which varied along a four-dimensional grid of external conditions. First, we measured the prevalence of MiDAS for every combination of *GF_High*, *Glucose*, and *ROS_Ext* at 0, 25%, 50%, and 75% saturation with no external pyruvate, then we repeated the full scan for 50% and 100% pyruvate. Figure 6B shows the relevant heat maps, comparing wild-type cells (*top row*) to those in which we knock out *SIRT3* at time 0 (*middle row*). The change caused by *SIRT3* knockdown indicates that in the absence of external ROS *SIRT3* knockdown boosts MiDAS most in high growth factor environments (*bottom row, leftmost column*). In contrast, mild ROS (25%, *middle*) shifts the effects of *SIRT3* knockdown to low-growth factor/high glucose cells. Strong ROS erases the effects of *SIRT3,* as it drives MiDAS regardless. In contrast, 50% pyruvate tempers the *SIRT3* effect (*rightmost columns*), while 100% completely blocks MiDAS (**SM File 13**).

Next we repeated this experiment on 13 oncogenic and 11 tumor-suppressive mutations from the top 500 of cancer-driver genes across human cancers for which our model has a corresponding node (**SM File 13**) [110]. We summarized our findings using an abstract version of the 4D environmental space, placing the name of proto-oncogenes and/or tumor suppressors that *altered* the model’s tendency to enter MiDAS into the position in environment-space where their effect was strongest (Fig. 6C). Thus, *SIRT3* knockdown appears in red (it up-regulates MiDAS), and its effect is most pronounced in high GF, high glucose, and no pyruvate (*bold SIRT3 labels*). In contrast, p53 knockdown reduces ROS-induced MiDAS, though its effects in pyruvate are strongest at high GF (*blue p53 labels*, **SM** Fig. 17). These results are in line with experimental data showing that p53 is required for mitochondrial hyperfusion in senescence [22]. Overall, our model predicts that oncogene activation or tumor suppressor loss that helps bypass the G1/S checkpoint (excess Cyclin D1, Ras, AKT_H_, Myc, mTORC1; loss of pRB, FoxO3/1 or p21) increases the incidence of MiDAS, especially in low glucose/ no ROS, or in mild ROS not capable of triggering it MiDAS in otherwise quiescent cells. External pyruvate erases most of these effects, except for SIRT3 and p53 loss (effects much weaker). The only cancer-associated mutations, other than p53, capable of reducing MiDAS were p21 overexpression in low glucose, and to a lesser extent Cyclin E hyper-activation – also known to block the cell cycle due to a failure in origin re-licensing [111]. The remaining mutations have no effect on MiDAS in any condition we model, though many alter cell cycle progression and/or apoptosis, as expected (loss of Casp8/9, TSC2, ATM/ART; activation of RAF, MEK, ERK, HIF1ɑ, PI3K_H_ or p21_H_).

## Discussion

Our model brings together insights from the in-depth MiDAS study in [17] and a wide variety of experimental data on mitochondrial dynamics during cell cycle, apoptosis, and DNA damage-induced senescence with a previously published Boolean network model of cell cycle control linked to apoptosis [45]. The resulting 134-node Boolean model (Fig. 2) is novel in that it **a)** reproduces the two-way connection between cell cycle progression and mitochondrial dynamics (Fig. 3A), **b)** accounts for the role of mitochondrial fragmentation in apoptosis (including mitotic catastrophe), **c)** reproduces the reversible hyperfusion observed in glucose-starved cells (Fig. 3B), **d)** matches experimental data on *SIRT3* knockout and mitochondrial damage-induced MiDAS, including the role of NAD^+^ and its rescue by external pyruvate (Fig. 4), and **e)** models ROS-induced MiDAS as a the first stage of a DNA-damage mediated senescence program (Fig 5).

Our model offers a series of testable predictions:

a. In our model, the molecular drivers of MiDAS, such as the loss of *SIRT3*, rely on the feedback between hyperfusion, low mitophagy and mitochondrial damage caused by high ROS/low NAD^+^. This proposed mechanism leads to several related predictions:

− Mitochondria in *SIRT3*-null cells that enter *MiDAS* are hyperfused, with low ΔΨ_M_ / high mROS (Fig. 4).
− p53 activity in MiDAS is not oscillatory, as neither AMPK or ATR (active in the presence of ROS-induced damage) are periodically inhibited by p53-induced Wip1 [109] (Figs. 4B, 5).
− *SIRT3*-null cells arrest in *MiDAS* from G2, with 4N DNA content (Fig. 4B).
− Quiescent cells with access to glucose are less susceptible to SIRT3 loss- and ROS-induced MiDAS due to FoxO-mediated mitophagy, which blocks MFN1/2 from inducing hyperfusion (**SM** Fig. 7B).
− Glucose-starved quiescent cells enter MiDAS in response to *SIRT3* knockdown (**SM** Fig. 7C). These predictions can be tested by monitoring mitochondrial morphology, ΔΨ_M_, and MFN1/2 localization in response to *SIRT3* loss, as well as H_2_O_2_ exposure in conditions involving various combinations of growth stimuli and glucose.
b. MiDAS cells that have not yet established deep senescence (i.e. 2-3 days post induction by ROS or ETC inhibitors) are rescued from MiDAS by subsequent pyruvate exposure (**SM Figs. 8-9**), or by forced SIRT3 activation (**SM** Fig. 14B). This parallels the MiDAS rescue by NAD^+^ precursor nicotinamide mononucleotide [17], or the increased expression of NAMPT in deep senescence required for proper SASP production [20]; both of which restore NAD^+^.
c. Cycling cells with hyperfused mitochondria near the G1/S boundary are highly susceptible to sub-lethal apoptotic signals that briefly lower ΔΨ_M_ but do not cause significant effector caspase activation (**SM** Fig. 10). A study documenting minority mitochondrial outer membrane permeabilization (MOMP) in senescent cells offers some indirect support to this prediction [89]. Minority MOMP only occurs in a small subset of mitochondria, it involves loss of ΔΨ_M_ and some cytochrome C leakage but no commitment to apoptosis, similar to the model’s dynamics. That said, a causal relationship between minority MOMP and senescence entry and/or MiDAS has not been established.
d. Modeling glucose withdrawal led to a cluster of predictions involving the timing of the withdrawal with respect to the cell cycle; detailed in ***SM Table 3*** (*blue font*). These include mitotic catastrophe and endoreduplication upon glucose re-exposure in a small subset of cells. These predictions could be tested by live imaging experiments that test the proposed cause of mitotic catastrophe (lack of timely mitochondrial fission), and by monitoring the ploidy of cells exposed to strong mitogens, along with cyclic glucose withdrawal / re-exposure.
e. Our cancer-related mutations screen predicts that the activation of oncogenes and/or loss of tumor suppressors that result in G1/S bypass tend to increase MiDAS, an effect blocked by saturating pyruvate exposure (Fig. 6C). These include Cyclin D1, Ras, AKT_H_, or Myc hyperactivation, and pRB, FoxO3 or p21 loss. A notable exception, as expected from the literature, is p53 loss, which blocks MiDAS [22].

To achieve a model complexity capable of reproducing the above range of observed cell behaviors, we found the Boolean modeling approach necessary as a constraint on the network’s dynamical rules. The type of data that could effectively constrain the large number of parameters needed for a concentration-based (continuous) model, such as detailed kinetic data on individual interactions and/or fine-grained time courses of molecular changes, are not currently available. That said, the Boolean framework is not without limitations. Key among these is the challenge of capturing dynamic homeostatic equilibria, such as the precise balance of mitochondrial fission and fusion that keeps cells healthy. While we account for *cell-wide* shifts in the balance leading to hyperfusion or excess fragmentation, our model does not capture subtler changes to mitochondrial dynamics such as the rapid, flexible fusion/fission cycle seen in dividing cells versus the more frozen morphology in aging cells [29]. Related to this, we purposefully left out PGC1α activation by mTORC1 [112], as this would force our model to activate MFN1/2 any time mTORC1 was active, and either cause hyperfusion well ahead of the G1/S boundary or require us to also account for mTORC1’s effect on increasing *Drp1-*driven fission. Overall, it appears that mTORC1 balances the induction of mitochondrial biogenesis downstream of PGC1α with increased fission [113], rendering the mitochondrial network more dynamic but not hyperfused. Here we chose to omit this balancing act, and thus are likely missing the role of mTORC1 in the induction and maintenance of senescence. Future work exploring a hybrid model that links the Boolean dynamics of enzyme expression / activity to a metabolic flux balance model could help mitigate these drawbacks, offer a precise accounting of oxidative phosphorylation as well as the Warburg effect [114], and probe the ROS-driven breakdown of mitochondrial energy metabolism during MiDAS.

Another limitation of our model’s applicability, albeit by design, is that it does not address the processes leading to deep senescence. This design allowed us to examine the conditions and feedback loops that drive mitochondrial dysfunction and stabilize MiDAS in a modular way, without the parallel influence of p53- and p21-dependent changes leading to damage-induced deep-senescence [15]; clearly distinct from MiDAS [17]. Briefly, the omitted process involves high p21-mediated p38 MAPK activation, leading to increased cytosolic ROS that locks in a p21 → p38 → ROS & p21 feedback loop [13]. This in turn damages both DNA and mitochondria. The loop is stable without ongoing p53 activity, maintains early senescence [115], and slowly activates deep-senescence promoting transcription via FoxA1, HBP1, and HMGA1, up-regulating p16 [24,116,117]. Together they assemble Senescence-Associated Heterochromatin Foci (SAHF), creating deep senescence. About 7 days following the initial damage HMGA1 induces NAMPT [20], while an unknown process downregulates p53 [26]. The resulting functional mitochondria and NF-κB reactivation change the SASP from primarily growth- and EMT-promoting (growth factors, TGFβ) to pro-inflammatory (IL-1/6/8) [118]. Without these pathways the phenotypic predictions of our model related to the fate of wild-type or mutant cells are limited to predicting MiDAS. The drawback is that the reversal of MiDAS in our model can reestablish proliferative capacity (**SM** Fig. 14), while cells *in vitro* remain arrested and senescent due to the non-mitochondrial p21/p38/ROS feedback [15]. Thus, incorporating this deep-senescence circuit downstream of ionizing radiation and/or UV DNA damage is an immediate future direction. The main challenges are the unknown mechanisms that lower p53 in deep senescence, and delineating the precise conditions that control the boundary between non-lethal vs. lethal damage, as the latter is known to be cell-type specific [119].

Our model raises the intriguing question: is MiDAS always reversed in deep senescence? Our model implicitly assumes that if / when NAD^+^ levels are restored, the cell’s mitochondria have an adequate reservoir of healthy mtDNA to cycle out of their dysfunctional state via enhanced mitophagy and mitochondrial biogenesis. Yet, sustained oxidative stress compromises mtDNA, and while mitochondria can tolerate a high mutation load, the system has a breaking point that permanently crashes the ΔΨ_M_ [120]. Thus, we expect a population of cells to respond to both the initial DNA damage and the subsequent chronically high ROS in a heterogeneous manner, generating some cells that reverse MiDAS to establish a canonical pro-inflammatory SASP alongside others that remain permanently stuck in MiDAS. Given the role of a pro-inflammatory SASP in recruiting the immune cells that clear senescent cells from a healthy tissue [9], we can further ask: *can deep-senescent MiDAS cells hide from the immune system?* This could provide a mechanism for the observed decline of senescent cell clearance with age [121]. Due to their enhanced ROS production, MiDAS cells may also be responsible for the population-dependent increase in senescent cell accumulation [122], shown to explain organism-level features of aging [123]. We expect mutations in a tumor setting to boost this heterogeneity, as several cancer-associated mutations guarantee permanent MiDAS (Fig. 6C). A stochastic version of our model with a weak noisy link from mitochondrial ROS to an mtDNA node that eventually shuts off the ETC and locks the system into a low-ΔΨ_M_ state could model this heterogeneity. Such a model would support the Stochastic Step Model of Replicative Senescence [124], offering specific molecular mechanisms responsible for tissue homeostasis versus aging. A hybrid Boolean/metabolic model proposed above could provide a more precise framework for capturing the metabolic feedback between MiDAS, deep senescence, and mtDNA homeostasis.

A long-term goal is to model cellular heterogeneity within a multicellular tissue. With our Boolean models at its heart, a multi-scale model of tissue homeostasis, aging, or cancer evolution could bring clarity to the structure of cellular heterogeneity, division of labor, and cooperation in micro-environments characteristic to each. Ideally done after deep senescence is accounted for, this project could bring together the conceptual clarity of population-only tissue aging models [123,124] with the predictive power displayed by earlier efforts at such multi-scale integration based on the models of mitochondrial dysfunction that inspired ours [29]. This multi-scale model could then examine the interplay between healing (involving damage, apoptosis, senescence, as well as cell cycle entry) and aging (involving senescent cell accumulation).

Furthermore, it could help probe the effects of cancer microenvironments on the behavior and heterogeneity of genetically identical cells (e.g., glucose and pyruvate availability and its effects on susceptibility to ROS [125]). Tying it all together, this multi scale model could predict the direct and/or combinatorial contribution of several cancer-associated mutations on cancer evolution, along with their ability to alter healing and aging in a tumor’s neighborhood.

## Acknowledgements

The authors would like to thank Samuel Pastva for his help with *AEON.py* in finding the asynchronous attractors of the model, and Ruthie Ressler for providing a Notepad++ language module for *.dmms* and *.vex* files.

## Author Contributions

H.S. — Conceptualization, Data curation; Formal analysis; Investigation; Writing - review & editing (designed the first version of a Boolean MiDAS switch, recognizing the relationship between hyperfusion, reduced mitophagy and senescence; provided revisions and manuscript edits).

D.D. — Data curation; Formal analysis; Methodology; Investigation; Writing - review & editing (helped refine the MiDAS module, build metabolic inputs accounting for the dynamics of NAD^+^; ran AEON, created Jupyter notebook, edited/restructured manuscript).

M.H. — Data curation; Formal analysis; Investigation (worked on the integration of the MiDAS module with cell cycle control and apoptosis).

K.F. — Data curation; Formal analysis; Investigation; Writing - review & editing (worked on the integration of the MiDAS module with cell cycle control and apoptosis, edited manuscript).

P.L.R. — Software; Methodology; Visualization; Writing - review & editing (developed the *dynmod* software package that simulated the model’s dynamical behavior, edited manuscript).

K.G. — Supervision; Resources (supported and supervised D.D., edited manuscript).

E.R.R. — Project administration; Conceptualization, Data curation; Formal analysis; Investigation; Methodology; Visualization; Software; Supervision; Resources; Writing - original draft; Writing - review & editing (helped conceive project, supervised H.S., M.H., K.F. and P.L.R., acquired funding, developed the *Boolean_code_for_MiDAS* software, analyzed/refined final model and drafted the paper).

## Abbreviations

ROS: reactive oxygen species

MiDAS: Mitochondrial Dysfunction-Associated Senescence

NAD: Nicotinamide Adenine Dinucleotide

NADH: nicotinamide adenine dinucleotide (NAD) + hydrogen (H)

ECT: Electron Transport Chain

TCA cycle: tricarboxylic acid cycle

ΔΨ_M_: mitochondrial membrane potential

SASP: Senescence-Associated Secretory Phenotype

EMT: epithelial-mesenchymal transition

MOMP: mitochondrial outer membrane permeabilization

SAC: Spindle Assembly Checkpoint

## Supplementary Materials

**Supplementary Table 1** -- Large (107-page), formatted and referenced table describing the biological evidence behind each node, link, and logic gate of our model, organized by regulatory module.

**Supplementary Figures and File list** – *MiDAS Supp_Figures_And_Files_list.pdf* file containing 16 supplementary figures with captions, followed by a list of supplementary files with brief descriptions (16 files).

