## Supplementary Material for "Unlocking Mitochondrial Dysfunction-Associated Senescence (MiDAS) with NAD^+^ – a Boolean Model of Mitochondrial Dynamics and Cell Cycle Control"

**Table of Contents**

1. ***Supplementary Methods*** **3**
   a) Advantages of using *dynmod* to model large biological regulatory networks 3
   b) Installing Haskell and *dynmod* 4
   c) Boolean model building in Dynamically Modular Specification (*.dmms*) format 5
   - SM Table 1 — SM Table 1 - MiDAS_Cell_Cycle_Arrests_Apoptosis.pdf
   - SM File 1 — MiDAS_Cell_Cycle_Arrests_Apoptosis.sbml
   - SM File 2 — MiDAS_Cell_Cycle_Arrests_Apoptosis.dmms
   - SM File 3 — MiDAS_Cell_Cycle_Arrests_Apoptosis_Fine.booleannet
   - SM File 4 — MiDAS_Cell_Cycle_Arrests_Apoptosis.gml
   - SM File 5 — Dynamically Modular Model Specification.plist
   - SM File 6 — DMMS_VEX_Language_Notepad++.xml

   d) Synchronous attractor detection with *dynmod* 7
   - SM File 7 — MiDAS_Cell_Cycle_Arrests_Apoptosis_attr_gr20_10_2.0e-2_10.csv
   - SM File 8 — MiDAS_Cell_Cycle_Arrests_Apoptosis_attr_gr20_10_2.0e-2_10.xlsx
   - SM File 9 — Virtual Experiment.plist
   - SM File 10 — MiDAS__Main_Figures.vex

   e) Attractor to cell phenotype mapping, visualization and evaluation of model attractors 8
   f) Running simulations with *dynmod* 9
   - SM File 9 — Virtual Experiment.plist
   - SM File 10 — MiDAS__Main_Figures.vex
   - SM File 11 — MiDAS__SM_Figures.vex
   - SM File 12 — MiDAS__Sizek_Results.vex

2. ***Results, section 2 - Cell Cycle-Dependence of Mitochondrial Morphology*** **12**
   a) Model reproduces experimentally observed mitochondrial dynamics during the cell cycle 12
   - SM Table 2 — summary of model behavior vs. experiments
   - SM Figure 1 — full version of Fig. 3

   b) Model reproduces experimentally observed mitochondrial dynamics under cell cycle perturbations 14
   - SM Table 3 — summary of model behavior vs. experiments
   - SM Figure 2 — phase-dependent effects of Plk1 knockout
   - SM Figure 3 — MFN1/2 knockdown or forced lowering of $\Delta\Psi_M$
   - SM Figure 4 — Drp1 knockdown and/or forced mitochondrial hyperfusion
   - SM Figure 5 — cell cycle arrest and apoptosis with mitochondrial hyperfusion

   c) Model reproduces experimentally observed mitochondrial dynamics under glucose starvation 20
   - SM Table 4 — summary of model behavior vs. experiments
   - SM Figure 6 — endoreduplication or apoptosis prediction following glucose withdrawal

3. ***Results, section 3 - MiDAS*** **22**
   - SM Figure 7 — full version of Fig. 4B (MiDAS in response of SIRT3 loss) and protection in quiescence
   - SM Figure 8 — low ΔΨM-induced MiDAS prevented / rescued by pyruvate
   - SM Figure 9 — external pyruvate protects cells from and can reverse MiDAS
   - SM Figure 10 — sub-lethal MOMP triggers MiDAS

4. ***Results, section 4 - ROS-induced MiDAS*** **27**
   - SM Figure 11 — modeling ROS
   - SM Figure 12 — full version of Fig. 5
   - SM Figure 13 — reversible G2 arrest in brief ROS

- SM Figure 14 — SIRT3 hyper activation protects from / reverses ROS-induced MiDAS

5. ***Results, section 5 - Context-dependence of MiDAS*** **31**
    - SM Figure 15 — model phenotype (attractor) map with all live, diploid phenotypes in all environments
    - SM Figure 16 — model phenotype (attractor) map with all attractors in all environments

6. ***Results, section 6 - MiDAS in cells with cancer-related mutations*** **33**
    - SM Figure 17 — p53 loss blocks MiDAS
    - SM File 13 — MiDAS in cancer-associated mutants.pdf

7. ***Results Reproduced with Biased Asynchronous Update*** **34**
    - SM File 14 — MiDAS__Asynch.vex
    - SM Figure 18 — synchronous limit cycles vs. asynchronous complex attractors
    - SM Figure 19 — asynchronous version of Fig. 3
    - SM Figure 20 — asynchronous version of Fig. 4B, 4D
    - SM Figure 21 — asynchronous version of Fig. 5A-B
    - SM Figure 22 — asynchronous version of Fig. 5C-D

8. ***Model Behavior in Response to Random Network Errors*** **40**
    - SM Figure 23 — cell cycle and MiDAS in ensembles of randomly mutated networks

### 1. Supplementary Methods

#### *a) Advantages of using dynmod to model large biological regulatory networks*

Widely used Boolean modeling software such as GinSim [57], Cell Collective [58], BooleanNet [59], BoolNet [61], CellNetAnalyzer [62] already allow for efficient, probabilistic modeling of regulatory networks representing cell ensembles. Capabilities of these platforms include graphical user interfaces to create, document and simulate models, libraries in common programming languages such as Python or R, synchronous, asynchronous and probabilistic Boolean update options, efficient attractor detection and visualization of the state transition graph. Several of these software packages also handle reconstruction of networks from time series, generation of random network ensembles, robustness analysis via perturbation, or network control [57-59,61,62].

In contrast, our focus in developing *dynmod* was to automate the detection, evaluation and analysis of **biological phenotypes** represented by attractors and/or time-series of our models. While we do replicate some of the functionality common to existing platforms in order to allow *dynmod* autonomy (e.g., attractor detection with synchronous update, synchronous vs. asynchronous time series, or stimulating ensembles), there are a few key reasons for using *dynmod* with a *dmms* model file format (rather than SBML):

a) In the *dmms* model file format, the modeler specify a hierarchy of user-defined signatures attached to regulatory switches; the example in *1a* above shows the Mitochondrial switch. With these profiles *dynmod* can automatically map each attractor to a combinatorial phenotype profile (e.g., quiescent, alive, MiDAS).

b) Using the combinatorial phenotype profile of each attractor, *dynmod* can visualize but also filter attractors of interest. One of its key functions is to organize attractors within a coordinate system of independent environmental input-combinations, and visualize them using barcodes of their combinatorial phenotypes (SM Fig. 16). As not all biological attractors are of equal interest at all times (e.g., robust apoptotic attractors when studying other processes), *dynmod* can filter the list to visualize a user-specified subset (Fig. 6A).

c) As *dynmod* can phenotype arbitrary network states (or sequences to detect oscillations), it can automatically evaluate cell behavior in an arbitrary simulation sequence. To do this, *dynmod* compares the state of the network in each Boolean time-step to phenotypes of interest, and measures the fraction of time particular phenotypes are expressed, as well as the number of times cells undergo specific transitions (e.g., apoptosis or a full cell cycle).

d) As most other software, *dynmod* can collect statistics on large cell ensembles (independent simulation runs) in non-saturating environments and/or non-saturating perturbations (e.g., 10% Trail, 50% AMPK inhibition). While most software does this for network nodes only, *dynmod* can also detect complex phenotypes such as normal vs. erroneous cell cycle progression.

e) *Dynmod* allows us to set up simulations by specifying the initial environment and cell phenotype-combination (rather than each node's state, or an attractor ID), which allows us to use the same "*in silico* experimental protocol" as we iteratively improve our model (see *.vex* files described below).

f) Much like the *dmms* model file format, several other model formats include node and link metadata. That said, we designed *dmms* metadata fields to be plain text with LaTeX formatting tags, including in-line citations, and wrote a *dynmod* module that transform these metadata fields into a formatted, publication-ready supplementary table containing all biological justification and references used to build the model (SM Table 1).

##### *b) Installing Haskell and dynmod*

*Note:* This font, marks commands to copy-paste into command line or configuration file (e.g., .zshrc file on macOS).

- **macOS**

  i. Check that Terminal is running z-shell; if not change your shell with the command:

  chsh –s /bin/zsh

  ii. Install Xcode developer tools with: xcode-select –install

  iii. Install HomeBrew (simplifies installing libraries required by *dynmod*):

  /bin/bash -c "$(curl -fsSL https://raw.githubusercontent.com/Homebrew/install/HEAD/install.sh)"

  If you already have HomeBrew, run:

  brew update
  brew upgrade

  iv. Add HomeBrew's path to your .zshrc file by including:

  export PATH=/opt/homebrew/bin:$PATH
  export PATH=/opt/homebrew/opt/python/libexec/bin:$PATH

  v. Install the pdf-making packages *dynmod* needs by running:

  brew install cairo pkg-config pango

  iv. Install Haskell by running: curl -sSL https://get.haskellstack.org/ | sh

  v. Specify the path to Haskell and *dynmod* in your .zshrc file and include an update function:

```
# For Haskell Stack:
PATH=~/.local/bin:${PATH}

update_dynmod() {
   rm -rf dynmod
   git clone https://github.com/Ravasz-Regan-Group/dynmod
   cd dynmod
   stack install
}
```

  vi. Install and/or update *dynmod* by navigating to the pace you plan to keep the dynmod source code, then run: update_dynmod ; check your success by running dynmod . If this results in "command not found", you may need to quit and reload terminal. If the issues persist, there are likely errors in your .zshrc file. In this case, run the commands from the update_dynmod() function one at a time.

- **Windows:**

  i. Download "Haskell stack" for windows using Windows Installer from this address: https://docs.haskellstack.org/en/stable/

  ii. To check that stack was installed, write stack -- help into the command line (if the command is found, the output will contain useful command hints):

  iii. Update the MSYS2 package manager pacman, then install required packages:

  pacman: stack exec -- pacman -Syu
  git: stack exec -- pacman -S git
  cairo: stack exec -- pacman -S mingw-w64-x86_64-cairo
  pkg-config: stack exec -- pacman -S mingw-w64-x86_64-pkg-config
  pango: stack exec -- pacman -S mingw-w64-x86_64-pango

v. Specify file paths to these installations[1].

vi. Download *dynmod* from GitHub: stack exec -- git clone https://github.com/Ravasz-Regan-Group/dynmod

vii. Install dynmod:

· Close and restart the command prompt.

· Change directory to the new folder you downloaded dynmod to (cd yourfile_path)

stack install

· Test using the dynmod command. If everything is correctly installed, dynmod will generate a warning for a missing model file followed by use instructions.

##### *c) Boolean model building in Dynamically Modular Specification (.dmms) format*

Relevant SM Tables & Files:

- **SM Table 1**, included as *SM Table 1 - MiDAS_Cell_Cycle_Arrests_Apoptosis.pdf*: Large (107-page), formatted and referenced table describing the biological evidence behind each node, link and logic gate of our model, organized by regulatory module.
- **SM File 1**, included as *MiDAS_Cell_Cycle_Arrests_Apoptosis.sbml:* Model file in SBML-qual format used by other Boolean modeling software such as GinSim [PMID: 22144167] or The Cell Collective [PMID: 22871178]; also uploaded to BioModels (MODEL2312140001).
- **SM File 2**, included as *MiDAS_Cell_Cycle_Arrests_Apoptosis.dmms*: Model file in *.dmms* format, containing the model's modular organization (including module and node order), phenotype signatures for relevant regulatory switches or modules, all nodes with logic expression, node description, input link annotation and relevant citations used to auto-generate **SM Table 1**, as well as 2D node coordinates and colors used to generate **SM File 4**.
- **SM File 3**, included as *MiDAS_Cell_Cycle_Arrests_Apoptosis_Fine.booleannet*: Model file in BooleanNet format used by other Boolean modeling software such as BooleanNet [PMID: 19014577] (list of Boolean logic gates with no annotation).
- **SM File 4**, included as *MiDAS_Cell_Cycle_Arrests_Apoptosis.gml:* Visual network layout in *.gml* format used to generate **Fig. 2**, readable by yED (*https://www.yworks.com/products/yed*) or Cytoscape [PMID: 14597658].
- **SM File 5**, included as *Dynamically Modular Model Specification.plist:* Language module file for the BBEdit text editor (Mac OS X), allowing it fold blocks, mark keywords, and recognize comments in *.dmms* files. Upon installing BBEdit, place this file in /Users/*yourusername*/Library/Application Support/BBEdit/Language Modules, then associate this language specification to the *dmms* extension in BBEdit's Language preferences.
- **SM File 6**, included as *DMMS_VEX_Language_Notepad++.xml:* Language module file for the NotePad++ text editor (Windows), allowing it fold blocks, mark keywords, and recognize comments in *.dmms* and *.vex* files. Upon installing NotePad++, import this file from the Language/User Defined Language tab.

---

[1] An easy way to find the path to stack, use: stack uninstall .This will not uninstall stack; merely list all the things you would delete if you were to uninstall it; this includes the directory path.

- Copy the directory containing stack's tools. It should read: C:\Users\USERNAME\AppData\Local\Programs\stack\
- List the contents of this folder to find out the CPU architecture of your machine (e.g., x86_64-windows):
  dir C:\Users\USERNAME\AppData\Local\Programs\stack\
- Add the folder name listed by the above command to the path to stack, then list its contents:
  dir C:\Users\USERNAME\AppData\Local\Programs\stack\x86_64-windows\
- Add the name of the package manager directory starting with msys to the path, append mingw64\bin\, then list its contents yet again (DATE is the date of your current msys install, listed by the above command):
  C:\Users\USERNAME\AppData\Local\Programs\stack\x86_64-windows\msys2-DATE\mingw64\bin\
- Open Settings and search for "environment". Select "edit environment variables for your account".
- Open this, select path, then edit and paste the path you built above into this section.

Our focus in building the *dmms* file structure and *dynmod* was to generate an easily human-editable, but also machine readable file format specific to discrete-state models (SBML-qual is a more general framework, but not friendly to direct editing without a Graphical User Interface). **SM File 2**, with the filename *MiDAS_Cell_Cycle_Arrests_Apoptosis.dmms (*do not rename as it must match internal model name) contains our model in *.dmms* format. This file keeps track of all model-relevant data and metadata, such as our model's hierarchically modular structure, phenotype profiles for regulatory switches, experimental data justifying each node, link and gate, and node visualization details. The top-level structure of this file is organized as follows:

```
Model{
ModelMetaData{ … }
ModelMapping{// organizes molecules of the embedded model into modules or regulatory switches
    Switch: Restriction_SW (Myc, CyclinD1, E2F1, CyclinE, p27Kip1, pRB, p21_B)
    ….
ModelMapping}
SwitchProfiles{ // specifies molecular activity/expression signatures of modules that determine the cell's phenotype / behavior
SwitchPhenotypes{
    SwitchName: Mitochondria
        Fragmented:0 *= (MFN1_2:0, Hyperfused_OM:0, Drp1:1, Unfused:1)
        Normal_Mito:1 *= (TCA_cycle:1, MFN1_2:0, Hyperfused_OM:0, MP_Low:0, AMPK:0, Drp1:0, Unfused:0, mROS:0, NADp_m:1)
        MiDAS:2 *= (TCA_cycle:0, MFN1_2:1, Hyperfused_OM:1, MP_Low:1, AMPK:1, Drp1:0, Unfused:0, mROS:1, NADp_m:0)
SwitchPhenotypes}
SwitchProfiles}
ModelGraph{ // consists of a Node{ …} block for each regulatory switch; allows modeling the network at switch level}
Model{ // consists of a Node{ …} block for each molecule in the Boolean (or discrete-state) model
Node{
    NodeMetaData{…}
    NodeGate{
      DiscreteLogic{
            Myc *= GF or E2F1
      DiscreteLogic}
    NodeGate}
    InLink{ … }
Model} // closes the molecular model inside the outer one involving regulatory modules or switches
Model} // closes the switch-level model
CitationDictionary{ … } // references in BibTeX format, exported from a citation manager such as Zotero
```

For editing *.dmms* files on Mac OX / Windows, we recommend *BBEdit* / *Notepad++* with DMMS language modules included as **SM File 4** (copy to *~/Library/Application Support/BBEdit/ Language Modules*) and **SM File 5** (import under the *Language/User Defined Language* tab).

To parse and validate the *dmms* file using *dynmod*, run:

```
dynmod MiDAS_Cell_Cycle_Arrests_Apoptosis.dmms
```

Key export functions include:

**-t** to export the molecular-scale model to **.BooleanNet** format (simple list of logic gates; **SM File 3**):

```
dynmod -t MiDAS_Cell_Cycle_Arrests_Apoptosis.dmms
```

**-g** to export to **.gml** format (for network visualization; **SM File 4**):

dynmod **-t** MiDAS_Cell_Cycle_Arrests_Apoptosis.dmms

**-u** to update coordinates read from user-organized *.gml* and appended to metadata in *dmms* files (note that this requires NOT altering the network structure or hierarchy while organizing the *.gml* file):

dynmod **-u** MiDAS_Cell_Cycle_Arrests_Apoptosis.glm MiDAS_Cell_Cycle_Arrests_Apoptosis.dmms

**-s** to generate a formatted LaTeX document & references. A LaTeX editor then generates **SM Table S1**:

dynmod **-s** MiDAS_Cell_Cycle_Arrests_Apoptosis.dmms

To generate a SBML-qual format of the model (**SM File 1**, *MiDAS_Cell_Cycle_Arrests_Apoptosis.sbml*), install *bioLQM* [PMID: 30510517] (download the jar file associated to the latest release), and use Java 8 Runtime to run it. *BioLQM* can performs format conversions, including from a *.Booleannet* format (generated with *dynmod*) to SBML:

java -jar bioLQM-version.jar modelname.Booleannet modelname.sbml

##### *d) Synchronous attractor detection with* dynmod

Relevant SM Files:

- **SM File 7**, included as *MiDAS_Cell_Cycle_Arrests_Apoptosis_attr_gr20_10_2.0e-2_10.csv*: Table containing all attractors detected by *dynmod* across a grid of sampling runs with an increasing number of random seeds and time-course lengths to test for saturation ($N_{\rm rnd} \in \{10, 20, 35, \ldots, 200\}$ random initial conditions in each unique environment, $N_{\rm series} \in \{10, 20, 35, \ldots, 100\}$ noisy steps / initial condition, and $p_{\rm noise} = 0.02$).
- **SM File 8**, included as *MiDAS_Cell_Cycle_Arrests_Apoptosis_attr_gr20_10_2.0e-2_10.xlsx*: Formatted Excel version of SM File 7, containing all synchronous attractors detected by *dynmod,* where ON/OFF states are marked yellow/blue, rows containing environmental input nodes are gray, and headers mark each attractor by their global phenotype.
- **SM File 9**, included as *Virtual Experiment.plist*: Language module file for the BBEdit text editor (Mac OS X), allowing it fold blocks, mark keywords, and recognize comments in *.vex* files. Upon installing BBEdit, place this file in /Users/*yourusername*/Library/Application Support/BBEdit/Language Modules, then associate this language specification to the *vex* extension in BBEdit's Language preferences.
- **SM File 10**, included as *MiDAS__Main_Figures.vex*: Experiment file specifying all simulation *dynmod* ran to generate main figures for the paper, with annotated comments on how to use the *.vex* file format. File can be used to reproduce all main time-course figures by running dynmod -e MiDAS__Main_Figures.vex MiDAS_Cell_Cycle_Arrests_Apoptosis.dmms.

*Routine sampling. Dynmod* has two sampling modes. The first, quicker mode involves a single attempt at finding attractors by running *N* noisy time-courses of length *T* with noise *p* from different random initial conditions for each unique combination of environmental inputs (e.g., $N$=200, $T$=20, $p$=0.02) [56,65,66]. In each noisy time-step, each node gives an erroneous output with probability *p*, allowing the dynamics to deviate from steepest descent. As *dynmod* does this, it also checks which synchronous attractor basin each state along the noisy trajectory belongs to. Once this is finished, *dymod* runs a synchronous time course from each attractor state by changing one environmental variable at a time. This increases its ability to detect attractors representing biological phenotypes that are robust in one environment (large attractor basin, easy to detect), and remain stable as the environment changes but their basin shrinks. All detected attractors are exported to a *.csv* file *dynmod* can read and reuse (instead of resampling). We use this mode as we iteratively improve our models, and we specify its parameters in the experiment (.vex) files we also use to run simulations. Details on setting up this sampling can be found at the top of **SM File 9**, *MiDAS__Main_Figures.vex.*

*Testing the convergence of sampling.* The second, slower sampling mode repeats the above procedure multiple times, with increasing N and T. This can test the convergence of the number of attractors detected with increasing N & T, while also collecting all unique attractors from these separate sampling runs. Here we used a 20 by 10 grid with $N \in \{10, 20, 30, . . , 200\}$, $T \in \{10, 20, 30, . . , 100\}$, and p = 0.02, and the command-line tag **-- grid**:

```
dynmod --grid '20 10 0.02 10' MiDAS_Cell_Cycle_Arrests_Apoptosis.dmms
```

(the last number in --grid '20 10 0.02 10' is the gap between increasing N and T values). The outputs of this command are a heat-map of the number of attractors in each independent sampling run along the grid (indicating that most attempts find 96 attractors; not shown), and the full attractor list collated into a single *.csv* (**SM File 7**). We use this mode when our models are largely finalized, to perform a final, extensive synchronous attractor search.

*Update dependence of attractors.* The fixed point attractors detected by synchronous update are solutions of the Boolean gate equations and thus stable regardless of update. In contrast, synchronous limit cycles do not always correspond to unique complex attractors, and there can be complex attractors with no synchronous limit cycle to match. To test whether *dynmod* detected all fixed points and probe whether its limit cycles are robust to update, we also ran an exhaustive asynchronous attractor search using AEON [63] (see *Suppl. Mat. 7* for comparison).

##### e) Attractor to cell phenotype mapping, visualization and evaluation of model attractors

*Attractor to cell phenotype mapping.* The idea that each attractor of a well-built biological network model represents a unique stable cell phenotype in a given environment is conceptually elegant, but difficult to achieve while building large models; especially in projects that focus on the context-dependent responses of large networks in 5 or more independent environments [56]. For models that involve irreversible apoptosis [45,56,66], this guarantees a minimum of 32 apoptotic attractors (doubled by each additional Boolean input). Thus, even a well-designed model with no non-biological attractor can have 50-100 or more attractors ([56] has ~ 480). Early stages of model-building often generate many more.

To help us with this problem, we designed *dynmod* in tandem with the *.dmms* model format to automatically evaluate the phenotype of each attractor in a principled, semi-automatic way. It does this by using the hierarchically modular structure of *dmms* files, paired with user-defined '*SwitchPhenotypes*' (*Suppl. Methods 1a*). These blocks allow the user to define and name specific signatures (sub-spaces) of regulatory modules of interest. For example, the Apoptosis label is associated with (Casp3:1, Casp9:1), one of two mutually exclusive signatures of the apoptotic switch (dead / alive). *Dynmod* uses this structured metadata to assign a global, combinatorial phenotype to each attractor, such as 'quiescent, alive, MiDAS'.

*Visualizing attractors by external input and phenotype.* Next, *dynmod* organizes all model attractors in a 1 to 5 dimensional environment-coordinate space (inputs above 5, if any, are selected and locked by user) and visualizes each attractor using a barcode of its modular phenotype (see **Fig. 6A**). With all attractors within an environment grouped and translated into cell behavior, iteratively evaluating whether a large model generates experimental documented states across multiple environments is faster and less error-prone than examining attractors one figure or table-column at a time. It is also a quick way to probe the model for biologically relevant multi-stability, such as the coexistence of healthy and MiDAS cells in the same environment (presumably due to their different history). In addition, *dynmod* can filter the list of visualized attractors by excluding/including a user-selected list of phenotypes (e.g., no apoptotic states), and can restrict the environmental inputs to those of primary interest (e.g., not showing Trail = 1 once we have checked that it always forces apoptosis). Methods to do this are detailed below.

##### *f) Running simulations*

Relevant SM File:

- **SM File 9**, included as *Virtual Experiment.plist*: Language module file for the BBEdit text editor (Mac OS X), allowing it fold blocks, mark keywords, and recognize comments in *.vex* files. Upon installing BBEdit, place this file in /Users/*yourusername*/Library/Application Support/BBEdit/Language Modules, then associate this language specification to the *vex* extension in BBEdit's Language preferences.
- **SM File 10**, included as *MiDAS__Main_Figures.vex*: Experiment file specifying all simulation *dynmod* ran to generate main figures for the paper, with annotated comments on how to use the *.vex* file format. File can be used to reproduce all main time-course figures by running dynmod -e MiDAS__Main_Figures.vex MiDAS_Cell_Cycle_Arrests_Apoptosis.dmms.
- **SM File 11**, included as *MiDAS__SM_Figures.vex*: Experiment file specifying all simulation *dynmod* ran to generate supplemental figures of the paper, with annotated comments on how to use the *.vex* file format. File can be used to reproduce all supplemental time-courses via dynmod -e MiDAS__SM_Figures.vex MiDAS_Cell_Cycle_Arrests_Apoptosis.dmms.
- **SM File 12**, included as *MiDAS__Sizek_Results.vex*: Experiment file specifying all single time-course simulation by which *dynmod* ran reproduce previpsuyl published cell behaviors our model inherited from its predecessor in [PMID: 30875364]. File can be used to reproduce all supplemental time-courses via dynmod -e MiDAS__Sizek_Results.vex MiDAS_Cell_Cycle_Arrests_Apoptosis.dmms.
- *Jupyter notebook in Python* to reproduce main time-course figures with BooleanNet [59]: LINK

*Setting up in silico experiments in .vex files*. To use dynmod for simulations, we designed an "in silico experiments" file format (*.vex* extension). Much like dynmod, this human-editable but machine readable file has a hierarchical organization, with blocks that specify attractor detection or reuse, as well as a series of experiment types. For editing *.vex* files on Mac OX / Windows, we recommend BBEdit / Notepad++ with DMMS language modules included as **SM File 9** and **SM File 6** (same as *.dmms*). Detailed instructions on how to set up each block are included in the *.vex* files that reproduce all our results (main figure results in **SM File 10**; supplementary figure results in **SM File 11**; additional simulations mentioned in Table 2 but not shown in **SM File 12**). Key *.vex* command blocks include:

• ***Sampling.*** This block either instructs *dynmod* to perform a new sampling, or to use of previously generated attractors from a specified *.csv* file (these attractors are verified to assure they match the model). This block also provided options to only sample a subspace of the model's full environment-space to speed iterative model constitution, and expand iteratively with a read-and-sample option.

• ***InputSpaceDiagram.*** This block generates the attractor visualization by phenotype barcodes (specified in the model *.dmms* file under *SwitchProfiles*) and environment-combinations (see *1e* above).

• ***Initial conditions for all experiment blocks.*** Dynmod's attractor profiling allows us to set up simulations by specifying the initial environment along with a desired starting phenotype-combination, rather than each individual node's state. This setup is robust to model tweaks during development, and automatically sets up runs starting from each unique state of a limit cycle attractor, simulating the arrival of a signal at different stages along a cycle (for details, see comments explaining the precise setup in **SM Files 10-12**).

• ***Time course experiment types.***

— *Pulse1* block generates a synchronous time course with three intervals: a) initial cell state(s), b) response to a change in a single environmental signal of user-specified duration (including non-saturating signals such as *GF_High* = 0.6); and c) response to the reversal of the same signal. To simulate non-saturating inputs, *dynmod* overrides their value with a stochastic ON/OFF sequence with a fixed probability, tuning the cell's average exposure. In case the initial state is a limit cycle, it runs multiple time courses to show the response to the environment change at all points along the cycle.

— *KDOE* block generates a synchronous time courses with two intervals: a) initial cell state(s), b) response the simultaneous partial or full knockdown/hyper-activation of a set of internal nodes. Partial knockdown/ hyper-activation of a node involves forcing it OFF/ON with a fixed probability in each time-step; otherwise allowing it to obey its Boolean rule [45,56,65,66] (justification / limitations of approach detailed in [65,66]). In case the initial state is a limit cycle, it runs multiple time courses to show the response to the environment change at all points along the cycle.

— *GeneralExperiment* block generates synchronous or asynchronous time courses from a subset of cell states in a given initial environment, exposed to an arbitrary sequence of manipulations defined in distinct time windows. Within each window the cell's external environment may be set to an arbitrary non-saturating value-combination (e.g., *GF_High* = 0.5 & *Glucose* = 0.3). In addition, a subset of internal nodes can be partially or fully knocked down / hyper-activated.

• ***Output from time course experiments.*** All types of time-course experiments run a single time-course by default, and show the time-dependent activity of each node (organized by modules) as a function of time (e.g., Fig. 3). Optional settings also include:

• averaging node activity in each tilmestep across an ensemble of cells (multiple simulations specified with a *SampeSize* parameter), resulting in an averaged time course (e.g., SM Fig. 18A)

• not exporting a node-level time course figure (*NodeTimeCourse*: False)

• exporting a time course at the switch phenotypes level, showing whether the model's state matching each phenotype at each time-point (can be averaged across an ensemble; e.g., SM Fig. 18B)

• averaging the activity of a user-specified subset of nodes within each distinct time-interval, resulting in a bar chart of activity for each interval (*AvgBarChartNodes;* e.g., SM Fig. 18C).

• ***Sampling the model's behavior with increasing input and/or knockout/hyper-activation*** *(coming to* dynmod *soon; SM Files 10-11, 15 offer ways to use time-course outputs to generate similar data).* In order to get a sense of how a large ensemble of cells behaves across a range of non-saturating environments and/or node knockdown/hyper-activation treatments, we designed a series of sampling runs that evaluate a cell ensemble across the desired range of conditions by calculating:

• the average time cells spend in any static phenotype of interest (e.g., MiDAS)

• how many times cells complete a biological cycle (e.g., cell cycle, normalized to the length of the wild-type cell cycle)

• how often cells complete a cycle with a biologically relevant error (e.g., resetting to G1 from G2; premature anaphase, skipped cytokinesis)

• and the average time cells take to undergo a particular, typically irreversible transition (e.g., apoptosis).

The intuitive way to set up the ensemble is to start with $N_{\mathrm{sample}}$ cells in the same initial environment and biological state (attractor), and simulate their dynamics with the desired environment / knockdown/ hyper-activation for $T_{\max}$ update steps. The problem with this is that if our cells undergo apoptosis during this time-course, then the data we get on other phenotypes following these events is no longer relevant to real cell behavior (e.g., a cell might die in MiDAS but after death its mitochondria are fragmented, so the length of our simulation post apoptosis could heavily influence our pre-death results). To avoid this, we actually set up our ensemble as follows:

- We specify a STOP condition at which we stop tracking the dynamics of individual cells; the first time a simulated cell's state matches one any phenotype on the STOP list (e.g., `Apoptotic_SW:Apoptosis`, or `Apoptotic_SW:Apoptosis, Mitochondria:MiDAS`). Note: these STOP phenotypes may *not* be limit cycles.
- We specify the maximum simulation time for cells that do not reach the STOP condition (e.g., $T_{max}$ = 250 update steps used here; corresponds to 10 wild-type cell cycle lengths).
- We specify a total sampled time across the ensemble, such as $T_{total}$ = 500,000 update steps, rather than the size of the ensemble. This guarantees that $N_{sample}$ is at least $T_{total}/T_{max}$ = 2,000 cells, or more if they tend to reach the STOP condition faster (the rationale is that in this case we need a better sampling of their briefer pre-stop dynamics).
- In addition to the scanned environment and/or environment / knockdown/ hyper-activation (simulated for a range of values), we can also specify a background environment (e.g., scan the model's behavior across all *ROS_ext* values, but do so in 10% external pyruvate), and/or a background mutation (e.g., do the above scan in cells heterozygous for p53: 50% knockdown).

Using this shared setup, we ran a series of sampling simulations of increasing combinatorial complexity, including (note, we used our older code to do this; see [56]; *dynmod* update for scans coming to GitHub soon):

- 1D scan along a singe input (e.g., cell cycle rate as a function of *GF_High*)
- 1D scan with increasing knockdown/over-expression of one (or more) molecule(s), as shown on Fig. 4D (when more than one molecule is specified for the scan, their levels of knockdown move in lockstep; this works well to simulate broader-target inhibitors that target more than 1 molecule in the model).
- 2D scan across two inputs
- 2D scan across an input and a knockdown/over-expression
- 3D scan across three inputs
- 3D scan across two inputs and a knockdown/over-expression
- comparative 3D scan across three inputs, computing the difference between wild-type cells and a specific knockdown/over-expression (e.g., Fig. 6B)

<u>Simulations with BooleanNet.</u> For readers who wish to use BooleanNet [59] to reproduce our main findings, please see the Jupyter notebook in Python at *<u>https://github.com/deriteidavid/midas_boolean_model</u>*. This reproduces the synchronous time-course simulations included in our main figures (Figs. 3, 4B, 5).

### 2. Results, section 2 - Cell Cycle-Dependence of Mitochondrial Morphology

#### *a) Model reproduces experimentally observed mitochondrial dynamics during the cell cycle*

Relevant SM Tables and Figures:

- **SM Table 2.** Model reproduces experimentally observed mitochondrial dynamics during cell cycle progression.
- **SM Figure 1.** Full version of Fig. 3: Model reproduces cell cycle-linked mitochondrial dynamics (hyperfusion at G1/S, fission in mitosis) and G1 arrest in response to glucose withdrawal.

**SM Table 2.** Mitochondrial dynamics during cell cycle progression

| | **Model behavior**<br>✤ new here / * inherited from [45] / ✲ prediction | **Figure** | **Experimentally observed cell behavior** |
|---|---|---|---|
| | • Stimulation with low mitogens (*GF* = ON but $GF_{High}$ = OFF) leads to quiescent cells (**Fig 3A**, *left*).<br><br>• Stimulation with $GF_{High}$ = ON leads to continuous cell cycle progression (**Fig 3A**,*middle*). | **Fig 3A**<br><br>**SM Fig 1** | – An increasing fraction of cells enter the cell cycle at increasing *EGF* or serum concentrations, reaching ~100% at 20 ng/mL *EGF* / 5% serum in MCF10A cells [78]; similar results were reported for rat embryonic fibroblasts [79].<br>– Mouse fetal fibroblasts display wide heterogeneity in the timing of the G1 / S transition regardless of the level or duration *of IGF-I, EGF, PDGF-AA*, or *PDGF-BB* treatment [80]. |
| | • Mitogen washout experiments ($GF_{High}$ = ON to $GF_{High}$ = OFF) show that a cell can pre-commit to another division cycle before finishing its current mitosis and execute a full cell cycle in the absence of mitogenic stimulation (**Fig 3A**, *right*). | **Fig 3A**<br><br>**SM Fig 1** | – Rapidly dividing mammalian cells (MCF10A, Swiss3T3) can pre-commit to a division cycle before finishing their current one; they often execute a full cell cycle after mitogen withdrawal, such that their last exposure to mitogens occurs sometime during the previous G2 phase [81]. |
| | ✤ Stimulation with $GF_{High}$ = ON leads to cyclic mitochondrial dynamics during cell cycle progression (**Fig 3A**, *middle*), involving:<br>✤ *E2F1*-induced hyperfusion of the mitochondrial network at the G1/S transition, leading to increased $\Delta\Psi_M$ and ATP generation, which helps activate *Cyclin E*<br>✲ Strong ETC activity in a healthy hyper- fused network increases SIRT3 activity to block runaway ROS production.<br>✤ Resetting of the mitochondrial network to its basal state in G2 (similar to G0) due to *E2F1*inhibition by *Cyclin A/Cdk2.*<br>✤ *Cyclin B/Cdk1*-mediated activation of the fission protein *Drp1*, resulting in an unfused (fragmented) state with relatively normal $\Delta\Psi_M$, which aids bipolar mitotic spindle formation and SAC passage. | **Fig 3A**<br><br>**SM Fig 1** | – *E2F1* is a direct transcriptional inducer of *MFN2,* which in turn increases mitochondrial fusion [28].<br>– Mitochondrial inner membrane fusion raises the efficiency of OXPHOS and increases the network's $\Delta\Psi_M$ [27].<br>– A high $\Delta\Psi_M$ mediated by hyper fusion and resulting in increased ATP production in late G1 are required for *Cyclin E* accumulation and S-phase entry [28].<br>– *Indirect support*: SIRT3 is necessary for healthy proliferation [17].<br>– During mitosis, *Drp1* is phosphorylated and activated by *Cdk1/Cyclin B*, resulting in mitochondrial fragmentation and proper bipolar spindle formation [28,31,82].<br>– TCA cycle metabolite levels oscillate during cell cycle progression to support high $\Delta\Psi_M$ at the G1/S boundary, but do not crash in mitosis [83]. |

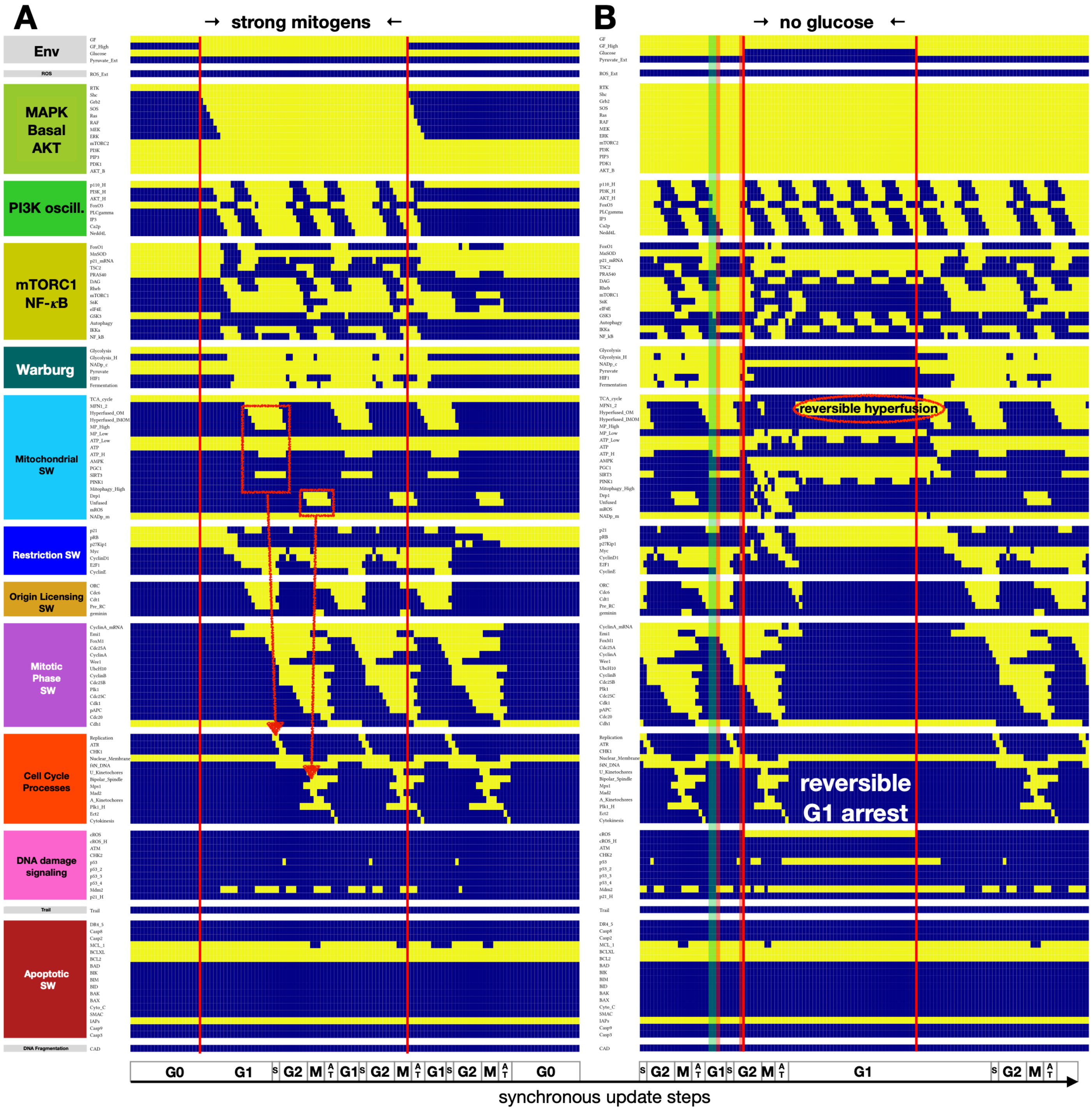


**SM Figure 1. Full version of Fig. 3: Model reproduces cell cycle-linked mitochondrial dynamics (hyperfusion at G1/S, fission in mitosis) and G1 arrest in response to glucose withdrawal. A)** Dynamics of regulatory molecule expression/activity during exposure of a quiescent cell to a saturating mitogenic signal for 60 update steps (full network). **B)** Dynamics of regulatory molecule expression/activity in a dividing cell responding to glucose withdrawal in early G1 for 50 update steps (full network). *X-axis:* time-steps; *y-axis:* nodes organized by regulatory modules; *yellow/dark blue:* ON/OFF; *vertical red lines:* start/end of signal; *red boxes with arrows:* morphological changes to mitochondria linked to cell cycle processes; *red oval*: reversibly hyperfused mitochondrial network; *shaded green box:* window of glucose withdrawal resulting in endoreduplicaiton (**SM Fig. 6A**); *shaded red bands:* windows of glucose withdrawal resulting in apoptosis (**SM Fig. 6B-C**).

##### *b) Model reproduces experimentally observed mitochondrial dynamics under cell cycle perturbations*

Relevant SM Tables and Figures:

- **SM Table 3.** Model reproduces experimentally observed mitochondrial dynamics under perturbations of cell cycle progression, accompanied by mitochondrial morphology change, and/or perturbations to mitochondrial fusion/fission proteins.
- **SM Figure 2.** Model reproduces the cell cycle phase-dependent effects of *Plk1* knockout and showcases difference between mitotic and apoptotic mitochondrial fragmentation.
- **SM Figure 3.** Model reproduces the cell cycle arrest induced by MFN1/2 knockdown or forced lowering of $\Delta\Psi_M$.
- **SM Figure 4.** Model reproduces loss of mitotic mitochondrial fragmentation and spindle assembly defects induced by Drp1 knockdown and/or forced mitochondrial hyperfusion.
- **SM Figure 5.** Model reproduces cell cycle arrest and apoptosis in cells with extreme mitochondrial hyperfusion.

**SM Table 3.** Perturbations of cell cycle progression accompanied by mitochondrial morphology change

| | **Model behavior**<br>✤ new here / * inherited from [45] / ✲ prediction | **Figure** | **Experimentally observed cell behavior** |
|---|---|---|---|
| | • Cycling cells experiencing *Plk1* knockdown at different points along the cell cycle, leading to:<br>— G2 arrest when *Plk1* is lost before the G2/M transition<br>— mitotic catastrophe when *Plk1* is lost in early metaphase<br>— aneuploidy when *Plk1* is lost in late metaphase<br>— no cytokinesis followed by endoreduplication when *Plk1* is lost after SAC passage. | **SM Fig. 2** | – *Plk1* activation by *Cyclin A/Cdk2* at the G2/M boundary is required for mitotic entry [84].<br>– *Plk1* is required for the formation and maintenance of microtubule-kinetochore attachments [85,86].<br>– Aneuploidy and genome duplication have been documented in *Plk1*-inhibited cells [87].<br>– Due to *Plk1*'s role in driving contractile ring assembly, *Plk1* knockdown in telophase blocks cytokinesis [88]. |
| | • Saturating *Trail* exposure kills quiescent as well as cycling cells.<br>• Non-saturating *Trail* leads to fractional killing<br>• Cells with prolonged metaphase are the most sensitive to *Trail*- mediated apoptosis. | virtual exp. in **SM File 10** | – Saturating *Trail* can kill ~100% of cells (immortalized human T lymphocytes [89], MCF10A [90], pancreatic cancer [91], glioblastoma & colon cancer cell lines [92]).<br>– *Trail* is synergistic with microtubule-targeting chemotherapy agents that trap cells in metaphase and delay SAC [92]. |
| | ✤ *Caspase 2*-mediated *BAK/BAX* activation during mitotic catastrophe activates *Drp1* to induce mitochondrial fragmentation, lowers $\Delta\Psi_M$, and increases mitochondrial ROS.<br>✤ In contrast to fragmentation in mitosis, apoptotic fragmented mitochondria aid *Cytochrome C* release and apoptosis. | **SM Fig. 2B** | – Mitotic catastrophe in IR-treated cells requires *Drp1*-induced mitochondrial fission [95]. |
| | ✤ Cycling cells experiencing *MFN1/2* knockdown show G1 arrest.<br>✤ Cycling cells experiencing forced low $\Delta\Psi_M$ (forced *MP_low* = ON) starting in late S-phase results in G1 arrest with no *Cyclin E* as well as increased *p21* and *p53*. | **SM Fig. 3** | – Cells infected with *Mfn2* shRNA show inhibited cell proliferation [96].<br>– Dividing cells treated with an oxidative phosphorylation uncoupler undergo G1 arrest, show a lack of *Cyclin E* accumulation and increased *p21* expression [28].<br>– Reducing $\Delta\Psi_M$ with FCCP results in G1–S arrest [28]. |

| | | | |
|---|---|---|---|
| | ✤ Cycling cells experiencing partial (60%) *Drp1* knockdown cannot sustain mitochondrial fragmentation during mitosis, resulting in problems with spindle assembly and prolonged SAC arrest.<br>✲ Cycling cells experiencing full *Drp1* knockdown arrest at the SAC long enough to undergo apoptosis.<br>✤ Cycling cells experiencing a brief period of *Drp1* knockdown in G2 do not arrest their cell cycle.<br>✤✲ Cycling cells experiencing prolonged *Drp1* knockdown arrest at the SAC and eventually undergo apoptosis.<br>✲ Cycling cells experiencing full *Drp1* knockdown undergo MiDAS.<br>✤ Cycling cells experiencing 50% forced outer membrane hyperfusion show prolonged SAC arrest due to spindle assembly delays and stochastically undergo apoptosis. | **SM Fig. 4 SM Fig. 5** | – Cells expressing dominant negative mutant *Drp1K38A* (DRP1m) fail to undergo mitochondrial fission during mitosis [31].<br>– *Indirect support*: high concentrations of mdivi-1 (mitochondrial division inhibitor) induce apoptosis [97].<br>– Cells overexpressing dominant negative *DRP1m* in a *brief* window before cells undergo mitosis experience no cell cycle inhibition [28].<br>– Perpetually hyperfused mitochondria generated by *prolonged DRP1m* expression significantly decreases proliferation [28].<br>– *Indirect support*: *Drp1* knockdown leads to mitochondrial dysfunction in muscle cells [97,98] and senescence in endothelial cells [99].<br>– Cells treated with a mitochondrial division inhibitor (mdivi-1) to induce hyperfusion have severely misaligned metaphase chromosomes [28].<br>– High concentrations of mdivi-1 not only block proliferation but also induce apoptosis [97]. |
| | ✤ *Trail*-induced apoptosis leads to *BAK/BAX*-mediated *Drp1* activation, mitochondrial fragmentation, and *Cytochrome C* release, along with the loss $\Delta\Psi_M$ and increased mROS.<br>✲ Partial Mitochondrial Outer Membrane Permeabilization (MOMP) via initially sub-lethal *Trail* can disrupt mitochondrial dynamics in G2/M, leading to mitotic catastrophe (discussed in *Results - 3*).<br><br>***Note:*** as most of our apoptosis results involve the intrinsic pathway, we chose not to include other extrinsic apoptotic signaling pathways such as FasL, TNF-α, or CD95-driven apoptosis. Given the similarities between FasL and Trail signaling, we hypothesize that our Trail results can be generalized to FasL signaling. In contrast, TNFR1 and CD95 also trigger pro-survival and pro-inflammatory signaling that may alter the fate of the cell and could alter the path to apoptosis and/or MiDAS (or trigger it by an alternate mechanism). The biological mechanisms needed to model this are not well explored in the experimental literature, and beyond our current scope. | **SM Fig. 10B** | – *Trail* induces *Drp1*-dependent mitochondrial fission by inducing Ser616 phosphorylation of *Drp1*, $\Delta\Psi_M$ loss, mROS, and cytochrome C release [94].<br>– *Indirect support*: non-lethal MOMP that only occurs in a small subset of mitochondria (minority MOMP) was documented in senescence, though it is unclear if it could be causal [94]. |

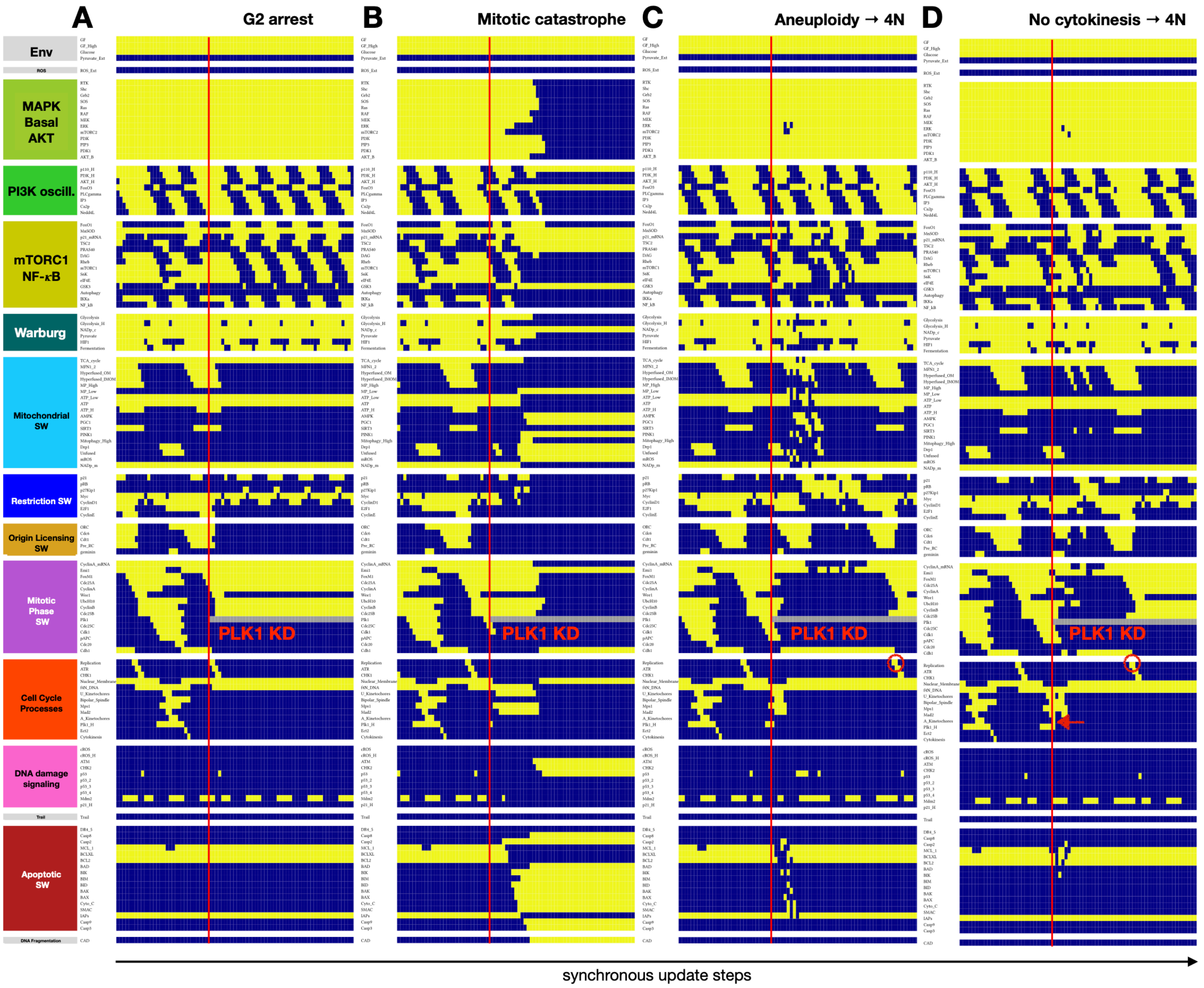


**SM Figure 2. Model reproduces the cell cycle phase-dependent effects of *Plk1* knockout and showcases difference between mitotic and apoptotic mitochondrial fragmentation. A-D)** Dynamics of regulatory molecule expression/ activity during full *Plk1* knockdown during cell cycle progression in (A) G2, leading to G2 arrest; *red box:* mitotic mitochondrial fission with normal $\Delta\Psi_M$; (B) prophase/early metaphase, leading to mitotic catastrophe and apoptosis; *red box:* apoptotic mitochondrial fission with low $\Delta\Psi_M$ and ROS production; (C) later in metaphase where *Cyclin A* loss results in $APC^{Cdh1}$ activation before apoptosis can set in, but also before the spindle is complete, leading to aneuploidy and endoreduplication; *red circle:* S-phase with 4N DNA content. (D) in anaphase past the SAC, leading to no cytokinesis and subsequent endoreduplication; *red circle:* S-phase with 4N DNA content; *red arrow:* node marking the completion of kinetochore attachment. *X-axis:* time-steps; *y-axis:* nodes organized in regulatory modules; *yellow/dark blue:* ON/OFF; *gray:* forced *Plk1* OFF state; *vertical red line:* start of *Plk1* knockdown.

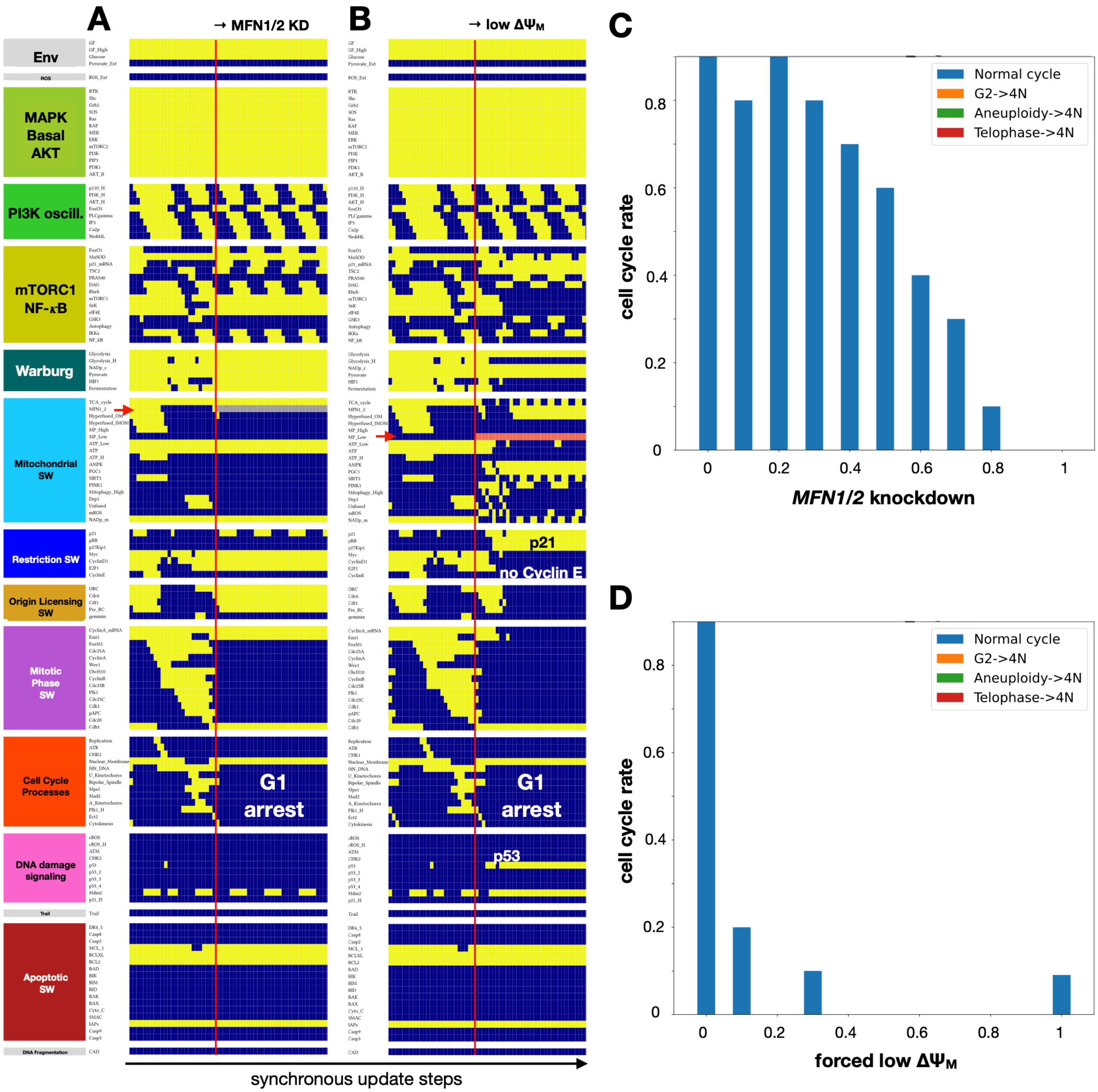


**SM Figure 3. Model reproduces the cell cycle arrest induced by MFN1/2 knockdown or forced lowering of $\Delta\Psi_M$. A)** Dynamics of regulatory molecule expression/activity during full *MFN1/2* knockdown in early G1, reproducing observations in [PMID: 25781899] that cells infected with *Mfn2* shRNA show inhibited cell proliferation; **B)** Dynamics of regulatory molecule expression/activity when $\Delta\Psi_M$ is forced to remain low starting before the previous cytokinesis, reproducing observations in [PMID: 19617534] that cells treated with the oxidative phosphorylation uncoupler FCCP arrest in G1, do not accumulate Cyclin E, and increase their *p21* expression. *X-axis:* time-steps; *y-axis:* nodes organized in regulatory modules; *yellow/dark blue:* ON/OFF; *gray/pink:* forced OFF/ON state; *vertical red line:* start of perturbation; *red arrows:* point of full *MFN1/2* knockdown, representing an inability to increase beyond its basal levels (A) and point of enforced low $\Delta\Psi_M$ (B); *white/black labels:* relevant outcomes. **C-D)** Response of cells dividing in 95% saturating growth signals to increasing *MFN1/2* knockdown (C) or increasingly low $\Delta\Psi_M$ (D). *Top/bottom:* rate of normal cell cycle completion (*blue*) vs. G2 → G1 reset (*orange,* not observed), aberrant mitosis (*green*, not observed), or failed cytokinesis followed by genome duplication (*red*, not observed), relative to the wild-type cell cycle length (25 time-steps), shown as stacked bar charts. *Initial state for sampling:* cycling cell in high glucose and no external pyruvate, ROS, or *Trail; sample size:* ≥ 2000 cells; *stop at:* apoptosis or MiDAS; *maximum length of single-cell tracks*: 250 steps (10 wild-type cell cycle lengths); *total sampled live cell time:* 500,000 steps; *update:* synchronous.

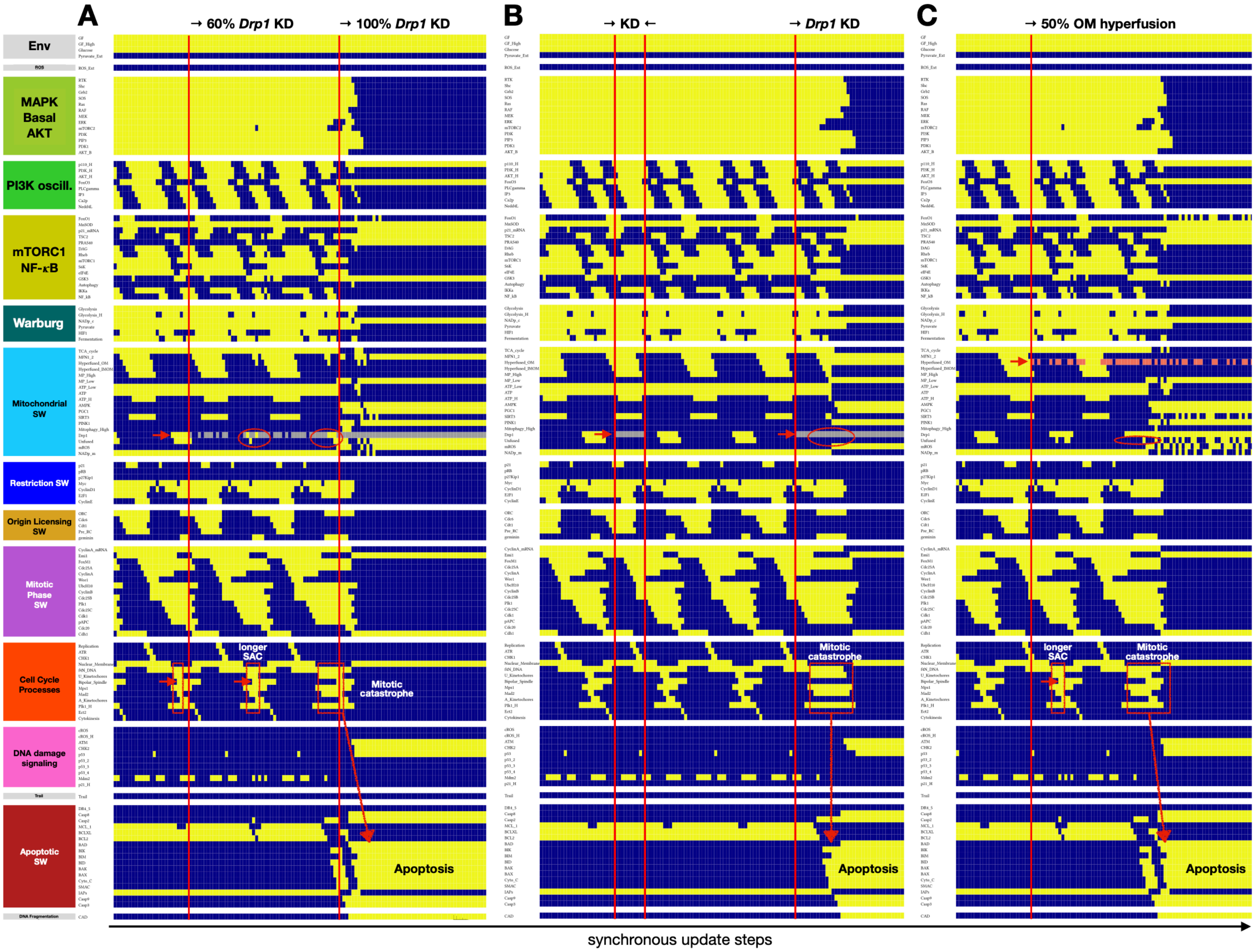


**SM Figure 4. Model reproduces loss of mitotic mitochondrial fragmentation and spindle assembly defects induced by Drp1 knockdown and/or forced mitochondrial hyperfusion. A-C)** Dynamics of regulatory molecule expression/activity during (A) partial vs. full *Drp1* knockdown (*middle:* for 50 steps/*last interval:* 50 update steps), (B) short vs. prolonged *Drp1* knockdown and (C) partial outer membrane hyperfusion in cycling cells, reproducing observations that: ***i)*** Cells expressing dominant negative mutant Drp1K38A (DRP1m) fail to undergo mitochondrial fission during mitosis (A, *red circles*) [PMID: 17301055]; ***ii)*** While there is no cell cycle inhibition in cells overexpressing DRP1m in a brief window before cells undergo mitosis (B, *left interval:* 10 update steps), perpetually hyperfused mitochondria (prolonged Drp1K38A expression) significantly decreases proliferation (B, *right interval*, predicting apoptosis) [PMID: 19617534]; ***iii)*** Cells treated with a mitochondrial division inhibitor (mdivi-1) to induce mitochondrial hyperfusion have severely misaligned metaphase chromosomes (C, *red box*) [PMID: 19617534]; ***iv)*** High concentrations of mdivi-1 not only block proliferation but also induce apoptosis (C) [PMID: 32147668]. *X-axis:* time-steps; *y-axis:* nodes organized in regulatory modules; *yellow/dark blue:* ON/OFF; *gray/pink:* forced OFF/ON states; *vertical red lines:* start/change in perturbation; *red arrows (top set):* start of *Drp1* knockdown (A,B) or outer membrane hyperfusion (C); *red boxes with arrow:* intervals between start of Metaphase (*U_Kinetochores* = ON) and Spindle Assembly Checkpoint passage (*Mad2* = OFF), with an arrow indicating the start of proper spindle alignment (*Bipolar_Spindle* = ON); *white/black labels:* relevant outcomes.

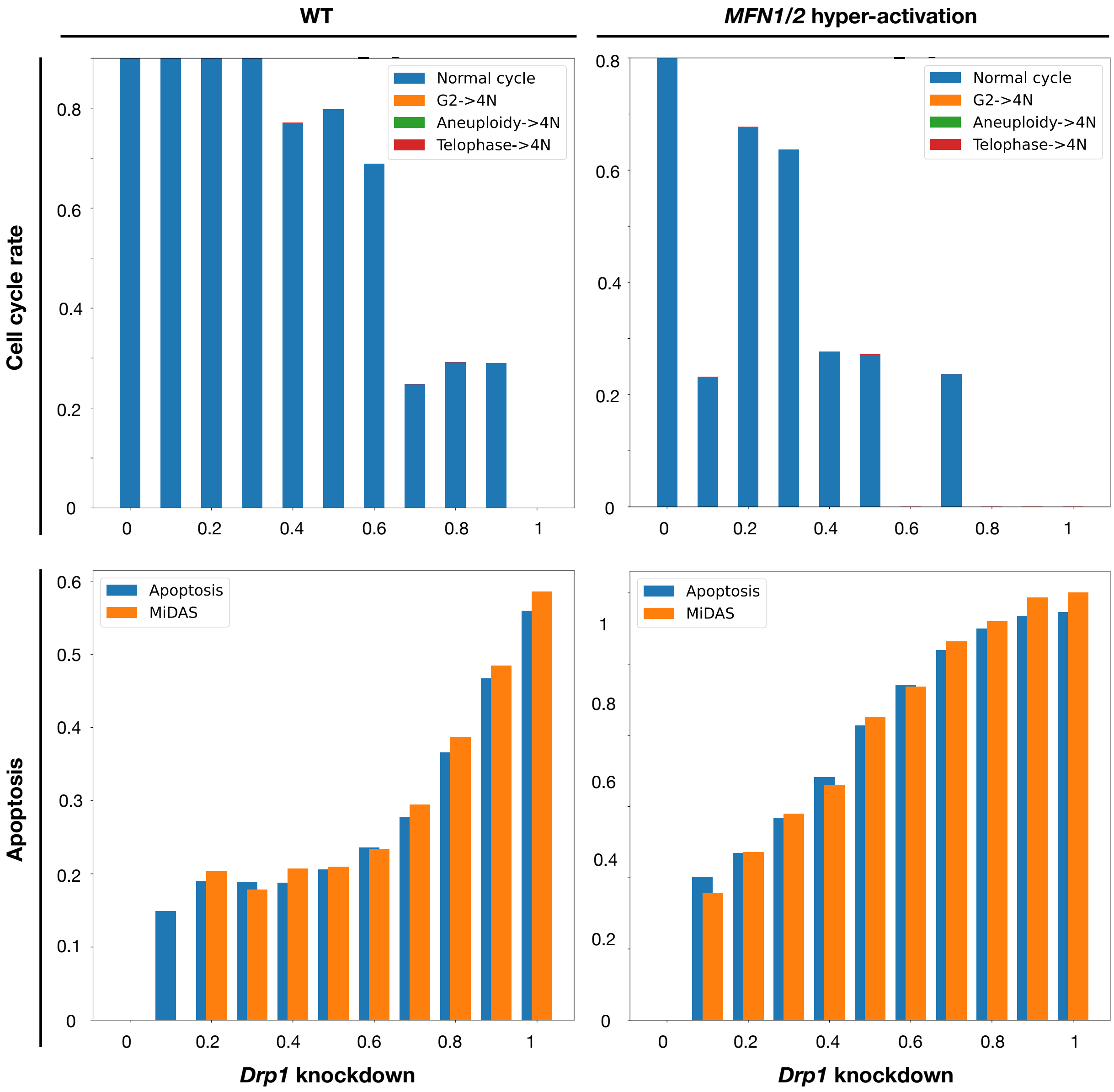


**SM Figure 5. Model reproduces cell cycle arrest and apoptosis in cells with extreme mitochondrial hyperfusion.** Response of cells dividing in 95% saturating growth stimuli to increasing levels of Drp1 knockdown alone (*left*) or in Mfn1/2 over-expressing cells (*right*). These two scenarios cover the range from normal dynamics to extreme hyper-fusion. Wild-type dynamics (*left, bars near 0*) and MFN1/2 hyper-activation that mitotic Drp1 can overpower (MFN1/2 *panels, bars near 0*) both show normal cell cycle progression. In contrast, the inability to form fragmented mitochondria without permanent hyperfusion (*wild-type, bars near 1 for 100%* Drp1 *knockdown*) and extreme hyperfusion (MFN1/2 *panels, bars near 1 for 100%* Drp1 *knockdown*) both block cell cycle progression and induce either apoptosis or MiDAS. Results reproduce the observed cell cycle arrest in cells with long-term *DRP1m* expression, where the mitochondrial network is hyperfused [PMID: 19617534]. They also *predict* a mix of apoptosis by mitotic catastrophe and MiDAS, supported by observations of apoptosis and mitochondrial dysfunction/senescence following Drp1 inhibition [PMIDs: 32147668, 32539155, 37517319] (these studies do not probe for specifically for MiDAS). *Top row:* rate of normal cell cycle completion (*blue*) vs. G2 → G1 reset (*orange,* not observed), aberrant mitosis (*green*, not observed), or failed cytokinesis followed by genome duplication (*red*, not observed), relative to the wild-type cell cycle length (25 time-steps) in wild-type (*left*) vs. *Mfn1/2* over-expressing cells (*right*), shown as stacked bar charts. *Bottom row:* rate of apoptosis (*blue*) relative to the wild-type cell cycle length (25 time-steps) in wild-type (*left*) vs. *Mfn1/2* over-expressing cells (*right*). *Initial state for sampling:* cycling cell in high glucose and no external pyruvate, ROS, or *Trail; sample size:* ≥ 2000 cells; *stop at:* apoptosis or MiDAS; *maximum length of single-cell tracks*: 250 update steps (10 wild-type cell cycle lengths); *total sampled time:* 500,000 steps; *update:* synchronous.

##### c) *Model reproduces experimentally observed mitochondrial dynamics under glucose starvation*

Relevant SM Tables and Figures:

- **SM Table 4.** Model reproduces experimentally observed mitochondrial dynamics under glucose starvation.
- **SM Figure 6.** Model predicts endoreduplication or apoptosis in rapidly cycling cells upon glucose withdrawal at different points along the G1/early-G2 window.

**SM Table 4.** Mitochondrial dynamics during glucose starvation

| | **Model behavior** ✤ new here / * inherited from [45] / ✲ prediction | **Figure** | **Experimentally observed cell behavior** |
|---|---|---|---|
| | ✤ Cycling cells glucose withdrawal undergo reversible G1 arrest, activate AMPK, which leads to PGC1α-mediated MFN1/2 up-regulation and indirect Drp1 inhibition mediated by *p53* (itself induced by AMPK and increased cellular ROS production).<br>✤ MFN1/2 up-regulation and Drp1 inactivation in response to glucose withdrawal leads to reversible mitochondrial hyperfusion and no increase in mitophagy (in spite of low $\Delta\Psi_M$ due to lack of glucose).<br>✲ Hyperfused networks in glucose-starved cells fuse their outer membranes only, as inner membrane fusion requires normal $\Delta\Psi_M$. | **Fig. 3B** | – Non-lethal glucose starvation induces reversible G1 arrest [100] with strong AMPK activation [101].<br>– Nutrient depletion, including glucose starvation, leads to significantly elongated and interconnected mitochondria, mediated by down-regulation of Drp1 and unopposed mitochondrial fusion [101].<br>– Mitochondrial hyperfusion upon nutrient deprivation protects mitochondria from mitophagy [101]. |
| | ✲ Cells that experience glucose withdrawal in a pre-committed G1 state; one that passed the restriction point *before* finishing its previous cytokinesis, undergo G2 arrest followed by endo-reduplication. Predicted mechanism (positive feedback on **SM Fig. 6B**; regulatory link details in **SM Table 1**):<br>— Glucose withdrawal leads to ROS accumulation and p53-mediated Myc repression.<br>— Loss of Myc decreases FoxM1, a critical G2 transcription factor responsible for Cdc25A/B, as well as Cyclin A induction.<br>— Positive feedback from Cyclin A/Cdk2 aided by Cdc25A/B can keep an existing FoxM1 pool active in G2 (**SM Fig. 6A**, *inset, gray arrows*), but glucose withdrawal interferes with this feedback and dooms the cell to an internal G1 reset.<br>— Upon re-exposure to glucose, the cell starts another round of DNA replication (**SM Fig. 6A**, *bottom red circle*). | **SM Figs. 6A, 6B** | – *Indirect support*:<br>~ Harsh tumor environments involving both glucose withdrawal and acidosis resulted in a ~19.4% pool of survivor cells with 4N DNA content (7 days of nutrient withdrawal with acidosis) [97].<br>~ 24 hours after nutrient restoration the fraction of G2/M cells briefly peak to a level that is consistent with pre-withdrawal levels (diploid cells reaching G2/M) *plus* the percentage of cells stuck with 4N DNA at day 7 of the withdrawal (potentially slower to re-enter the cell cycle) [97]. |
| | ✲ Hyperfusion in response to glucose withdrawal, if it persists into mitosis, delays mitotic mitochondrial fragmentation long enough to cause spindle assembly defects and apoptosis by mitotic catastrophe. | **SM Fig. 6C** | – *Indirect support*: glucose-starved cells show a significant increase in spindle formation defects such as multipolar spindles [97]. |

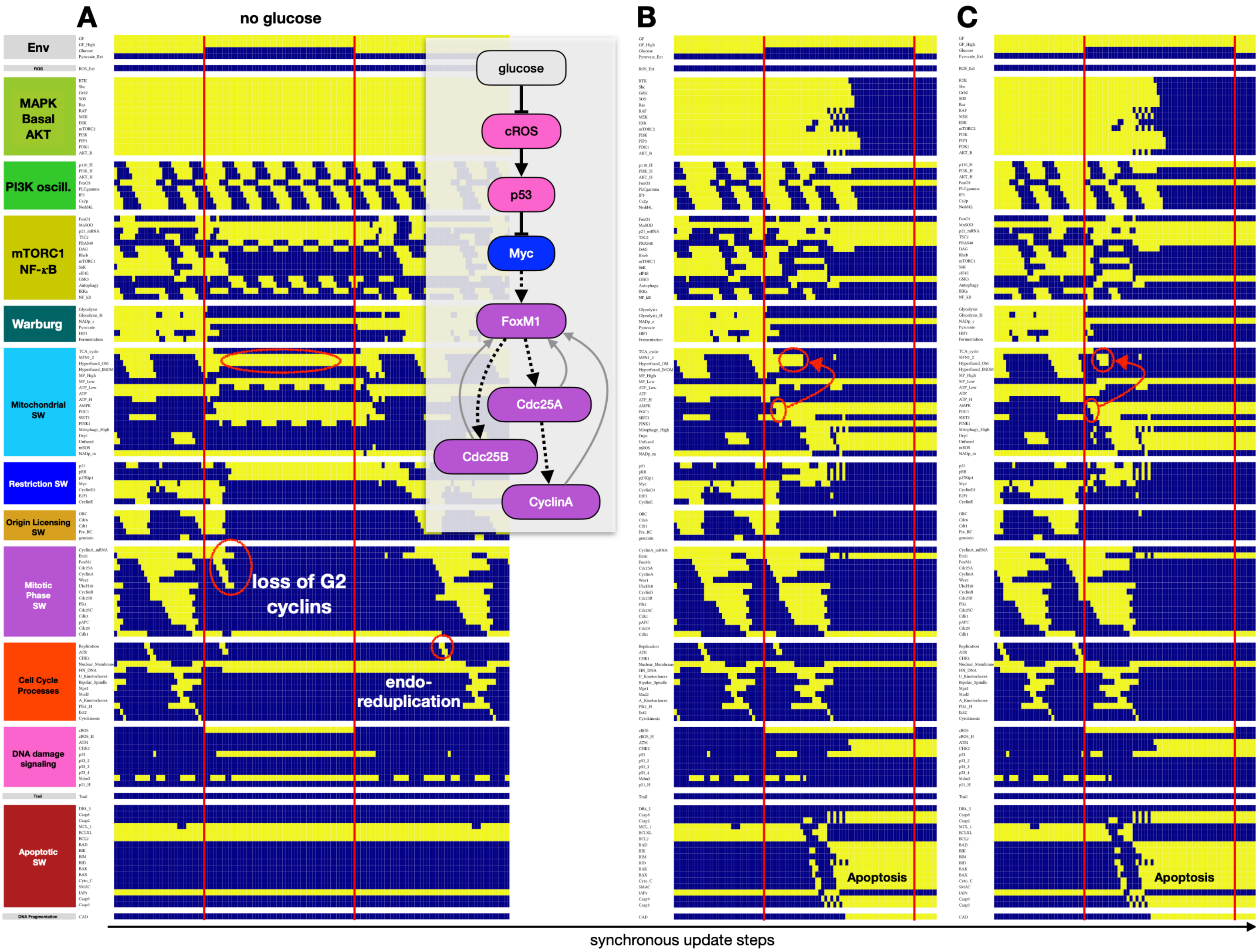


**SM Figure 6. Model predicts endoreduplication or apoptosis in rapidly cycling cells upon glucose withdrawal at different points along the G1/early-G2 window. A-C)** Dynamics of regulatory molecule expression/activity during glucose withdrawal for 50 update steps in (A) a pre-committed cell in early G1, which accumulates enough cyclin E to start DNA replication but cannot lock in a stable G2 state due to loss of Myc-induced FoxM1 accumulation (*inset*). (B-C) mid-G1 & early G2, where AMPK activation and hyperfusion prevent the timely mitotic fission of mitochondria (*red ovals with arrow*), causing bipolar spindle defects and mitotic catastrophe; *X-axis:* time-steps; *y-axis:* nodes organized in regulatory modules; *yellow/dark blue:* ON/OFF; *vertical red lines:* start/change in perturbation; *red ovals:* relevant molecular changes such as mitochondrial hyper-fusion (*top ovals*), loss of G2 cyclin A activity (A) and a second replication event with 4N DNA (A), AMPK → PGC1α activation (B, C); *white/black labels:* relevant outcomes.

#### 3. Results, section 3 - MiDAS

Relevant SM Tables and Figures:

- **SM Figure 7.** Model reproduces MiDAS in response to *SIRT3* loss and predicts that mitogen-arrested, but not glucose-arrested cells are protected from it.
- **SM Figure 8.** Model reproduces low $\Delta\Psi_M$-induced MiDAS prevented / rescued by pyruvate.
- **SM Figure 9.** Model predicts that external pyruvate can protect cells from MiDAS induced by *SIRT3* loss and reverse MiDAS to restore proliferation or healthy quiescence.
- **SM Figure 10.** Model predicts that sub-lethal MOMP that does not activate Caspase 9 but lowers $\Delta\Psi_M$ in cells with hyperfused mitochondria triggers MiDAS.

In addition to *SIRT3* knockdown, the electron transport chain inhibitor rotenone and mtDNA depletion were also shown to trigger MiDAS, a response that culturing cells in excess pyruvate could prevent [17]. To test whether our model can reproduce these experiments, we first mimicked lowered ETC activity by forcibly lowering the model cell's $\Delta\Psi_M$. Indeed, loss of $\Delta\Psi_M$ at the G1/S boundary in a cell committed to cell cycle triggered MiDAS, an effect rescued by pyruvate (**SM Fig. 8,** *middle vs. right window*). External pyruvate also prevented MiDAS following the loss of *SIRT3* (**SM Fig. 9A**). Moreover, the model predicted that pyruvate could rescue wild-type cells from MiDAS (early senescence only), freeing them to enter the cell cycle when exposed to mitogens (**SM Fig. 9B**) or restoring ATP generation and their potential for future proliferation in quiescence (**SM Fig. 9C**).

Further underscoring the key role of low $\Delta\Psi_M$ combined with mitochondrial hyperfusion, we have found that several perturbations described above for their role in disrupting the cell cycle can also trigger MiDAS – depending on their timing along the cycle. These include *Plk1* knockdown just before the cell clears the Spindle Assembly Checkpoint (SAC) (**SM Fig. 10A**), sub-lethal *Trail* exposure in cells pre-committed to the next cell cycle (**SM Fig. 10B**; late mitosis, starting hyperfusion for G1/S), and sub-lethal *Drp1* knockdown or mitochondrial hyperfusion in metaphase (**SM Fig. 10C-D**). The common predicted cause of MiDAS is a sub-lethal mitochondrial membrane permeabilization that lowers the $\Delta\Psi_M$ without activating executioner caspases (which would trigger apoptosis) – at a time when the mitochondria are hyperfused. As the cells they react to low $\Delta\Psi_M$ with *AMPK* activation, they lock in MiDAS.

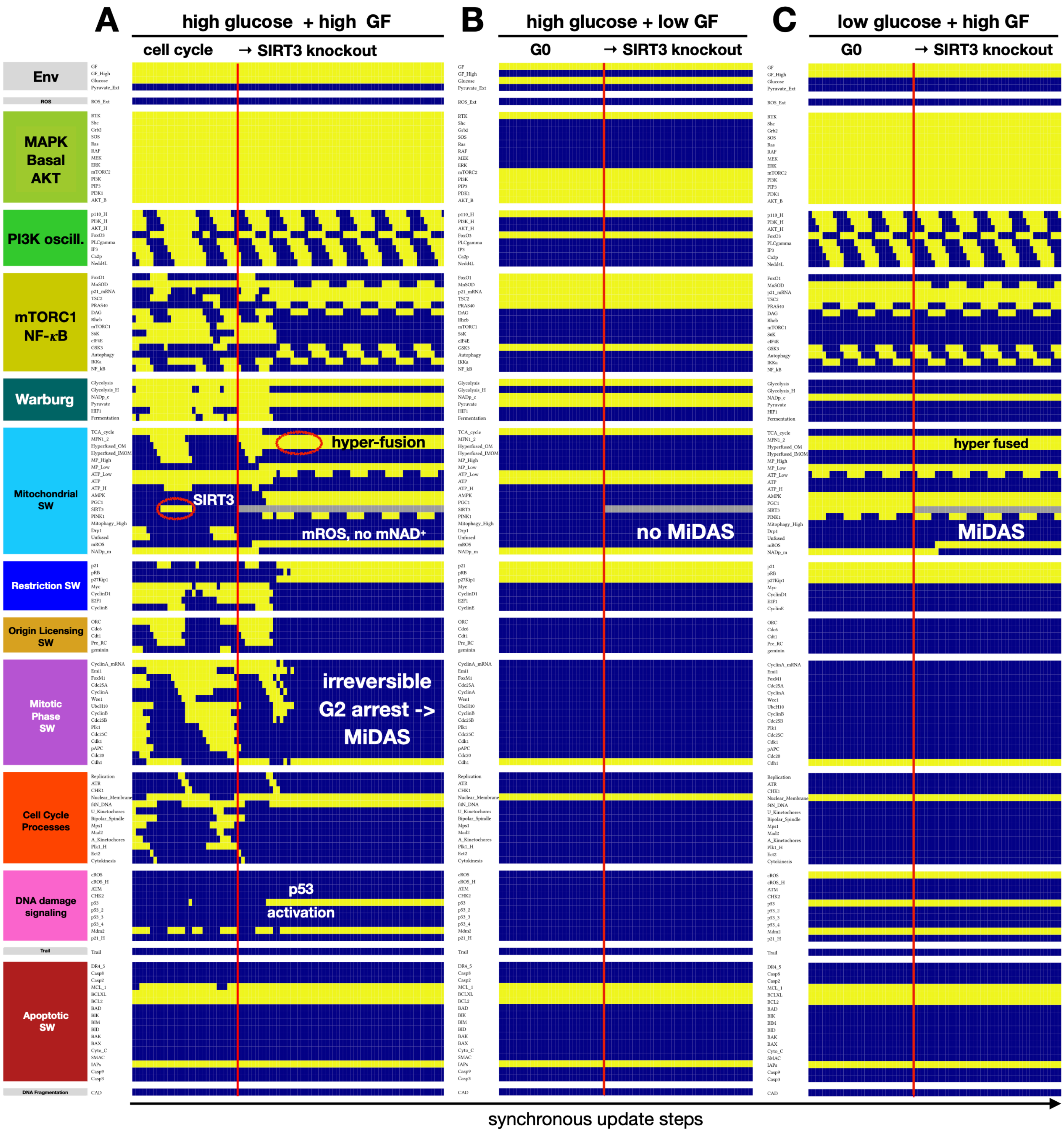


**SM Figure 7. Model reproduces MiDAS in response to *SIRT3* loss and predicts that mitogen-arrested, but not glucose-arrested cells are protected from it. A-C)** Dynamics of regulatory molecule expression/activity in: (A) a dividing cell responding to full *SIRT3* knockout in early G1 (full version of **Fig. 4B**); (B) a mitogen-arrested cell / (C) glucose-starved cell, responding to full *SIRT3* knockout, resulting in continued G0 / MiDAS, respectively. *X-axis:* time-steps; *y-axis:* nodes organized in regulatory modules; *yellow/dark blue:* ON/OFF; *pink/gray:* time-steps in which stochastic forced activation / knockdown has an effect; *vertical red lines:* start/change in perturbation; *white/black labels:* relevant outcomes.

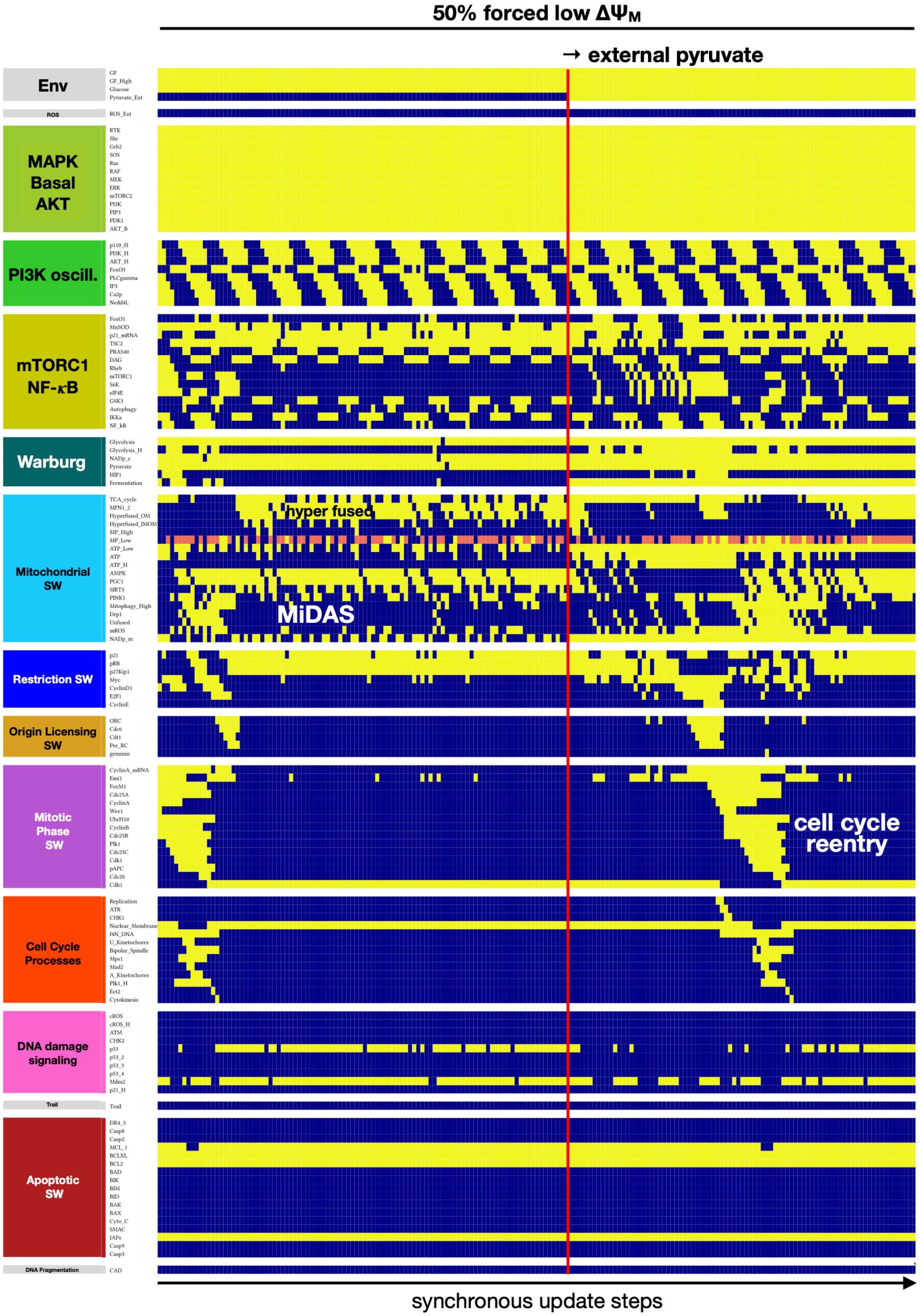


**SM Figure 8. Model reproduces low $\Delta\Psi_M$-induced MiDAS prevented / rescued by pyruvate.** Dynamics of regulatory molecule expression/activity in mitogen-stimulated cells exposed to an ETC inhibitor that lowers the $\Delta\Psi_M$ in 50% of the time steps, in the absence (*left*) vs. presence (*right*) of saturating external pyruvate. *X-axis:* time-steps; *y-axis:* nodes organized in regulatory modules; *yellow/dark blue:* ON/OFF; *pink/gray:* time-steps in which stochastic forced activation / knockdown has an effect; *vertical red lines:* start/change in perturbation; *white/black labels:* relevant outcomes.

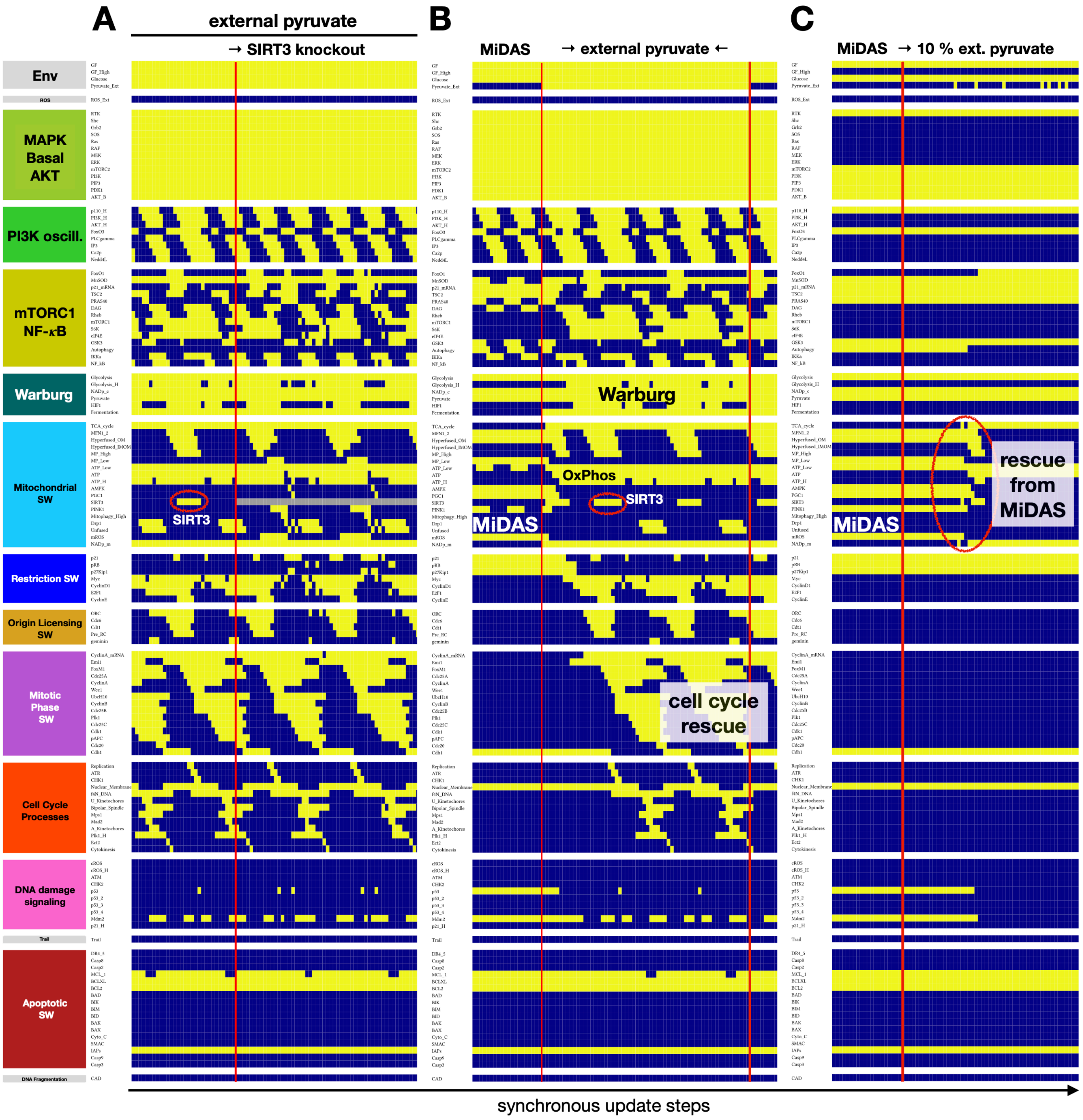


**SM Figure 9. Model predicts that external pyruvate can protect cells from MiDAS induced by *SIRT3* loss and reverse MiDAS to restore proliferation or healthy quiescence. A)** Dynamics of regulatory molecule expression/activity in cycling cells with access to saturating external pyruvate, in response to full *SIRT3* knockout. **B)** Dynamics of regulatory molecule expression/activity in mitogen-exposed MiDAS cells in response to a prolonged pulse (60 steps) of external pyruvate. **C)** Dynamics of regulatory molecule expression/activity in MiDAS cells under low growth factor exposure, in exposed to low-dose (10%) external pyruvate. *X-axis:* time-steps; *y-axis:* nodes organized in regulatory modules; *yellow/dark blue:* ON/OFF; *gray:* full SIRT3 knockdown; *vertical red lines:* start/change in perturbation; *white/black labels:* relevant outcomes.

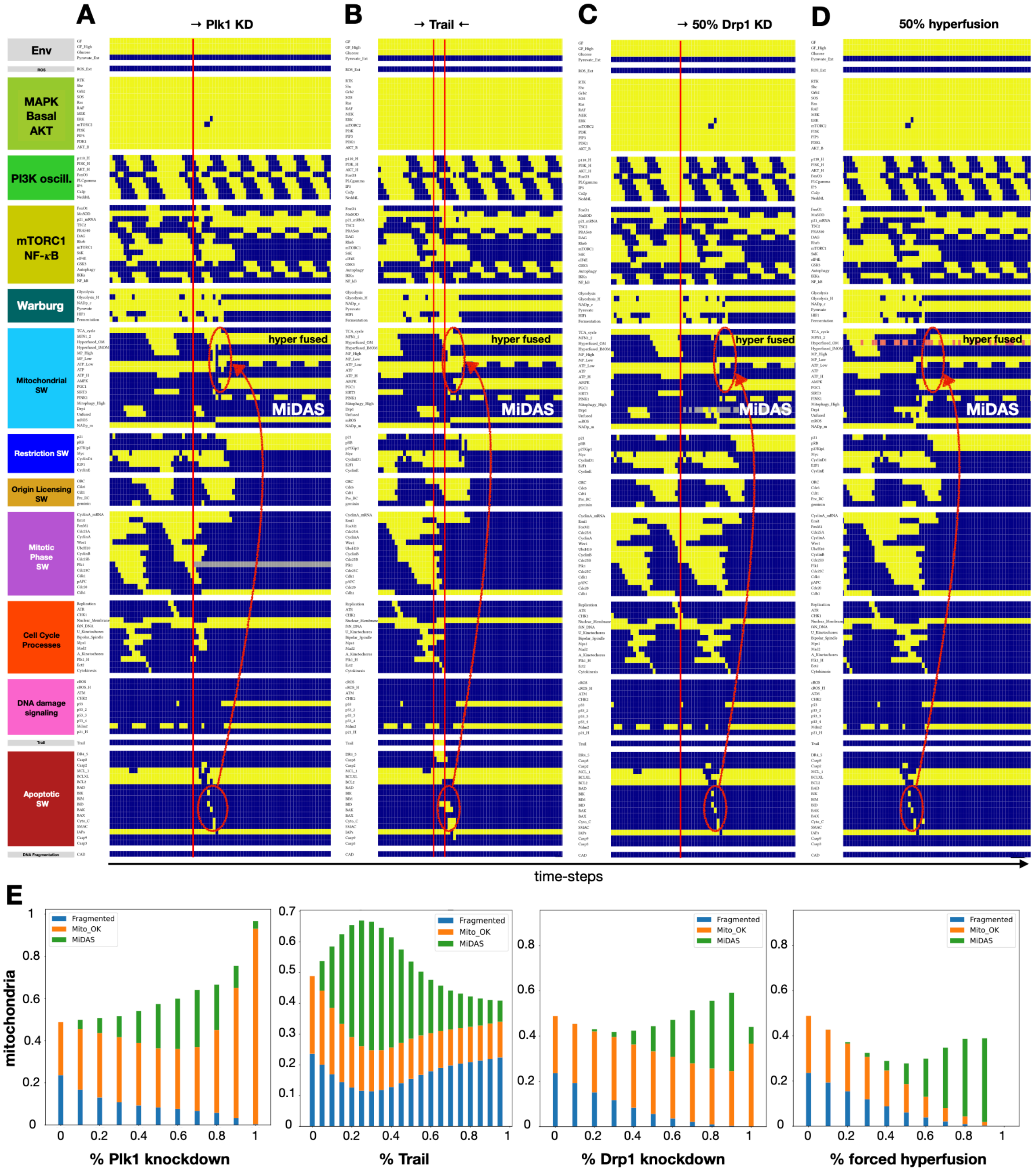


**SM Figure 10. Model predicts that sub-lethal MOMP that does not activate Caspase 9 but lowers $\Delta\Psi_M$ in cells with hyperfused mitochondria triggers MiDAS. A)** Dynamics of regulatory molecule expression/activity in cycling cells in response to full *Plk1* knockout in late metaphase, allowing them to narrowly avoid mitotic catastrophe but enter MiDAS. **B)** Dynamics of regulatory molecule expression/activity in cycling cells in response to a near-lethal *Trail* dose (4 steps) in anaphase, leading to MiDAS. **C)** Dynamics of regulatory molecule expression/activity in cycling cells in response to 50% *Drp1* knockdown, leading to prolonged SAC followed by MiDAS. **D)** Dynamics of regulatory molecule expression/activity in cycling cells in response to 50% outer mitochondrial membrane hyperfusion, leading to prolonged SAC followed by MiDAS. *X-axis:* time-steps; *y-axis:* nodes organized in regulatory modules; *yellow/dark blue:* ON/OFF; *gray/pink:* forced OFF/ON state; *red ovals & arrows:* start of MOMP (*bottom ovals*) leading to low $\Delta\Psi_M$ at a time of hyperfusion (*top ovals*); *vertical red lines:* start/change in perturbation; *white/black labels:* relevant outcomes. **E)** Response of cells dividing in 95% saturating growth stimuli to *Plk1* knockdown, *Trail, Drp1* knockdown and forced hyperfusion of the outer mitochondrial membrane (*left to right*), showing the fraction of time cells display normal mitochondria (*orange*), MiDAS (*green*), or fragmented mitochondria (*blue*). *Initial state for sampling:* cycling cell in high glucose, no external pyruvate, ROS, or *Trail; sample size:* $\geq$ 2000 cells; *stop at:* apoptosis; *maximum length of single-cell tracks*: 250 update steps (10 wild-type cycles); *total sampled time:* 500,000 steps; *update:* synchronous.

#### 4. Results, section 4 - ROS-induced MiDAS

Relevant SM Figures:

- **SM Figure 11.** Network neighborhood of the internal reactive oxygen species nodes *ROS* and *cROS_H*.
- **SM Figure 12.** Full version of Fig. 5: Model reproduces ROS-induced MiDAS in cycling cells and predicts protection from MiDAS in external pyruvate.
- **SM Figure 13.** Model reproduces reversible G2 arrest in response to brief / weak ROS exposure.
- **SM Figure 14.** Model predicts that *SIRT3* hyper-activation can protect from or reverse ROS-induced MiDAS, and reproduces MiDAS rescue by NAD+ boost.

In contrast to MiDAS in response to prolonged ROS, brief exposure led to *reversible* G2 arrest followed by mitosis (**SM Fig. 13**). Our model captures this difference via its four-node tracking of *p53* accumulation. In addition, our model predicted that quiescent cells were not as susceptible to ROS-induced MiDAS, owing their protection to continuous mitophagy (**Figs. 5C, SM 12C**). This was due to the lack of mitogen-induced $AKT_H$ oscillations, aiding *FoxO1/3*-mediated mitophagy in blocking *MFN1/2*-driven hyperfusion. That said, cells that exited the cell cycle into MiDAS maintained MiDAS even in the absence of mitogens; (**Figs. 5A-B,** *right*).

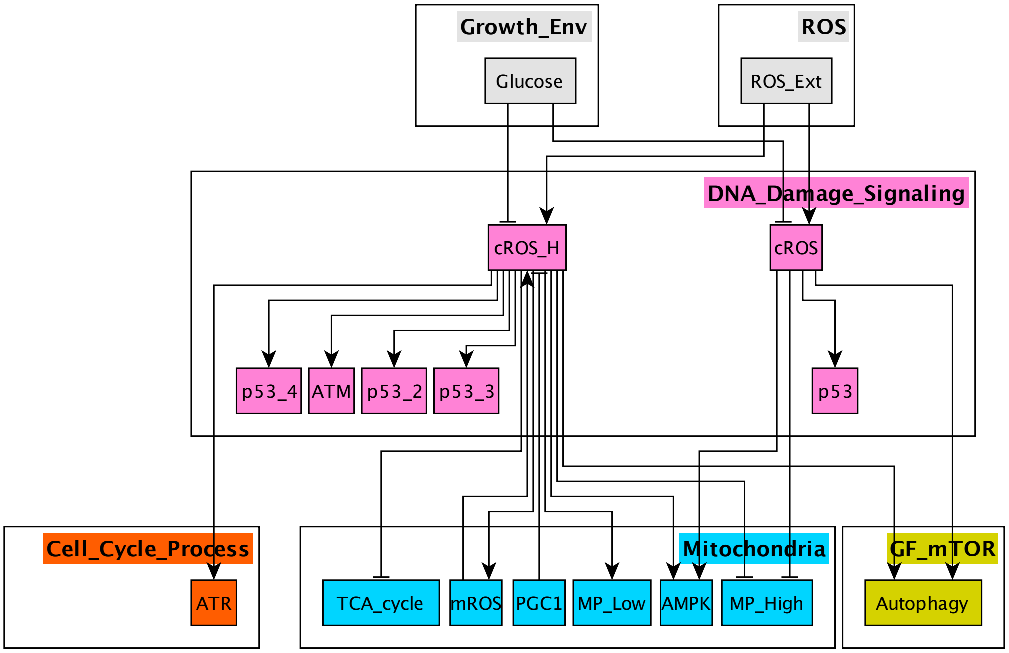


**SM Figure 11. Network neighborhood of the internal reactive oxygen species nodes *ROS* and *cROS_H*.** Part of our regulatory network model directly connected to the cROS and cROS_H nodes, including incoming and outgoing links. *Gray:* inputs representing environmental factors; *pink*: DNA damage response; *teal:* mitochondrial switch; *dark orange:* cell cycle processes; *mustard*: autophagy (new node in the mTORC1 growth module); → : activation; ⊣ : inhibition.

Given that external pyruvate could protect *SIRT3*-knockout cells from MiDAS by maintaining $NAD^+$ levels, we next asked whether pyruvate could also prevent or reverse ROS-induced MiDAS. Indeed, excess pyruvate prevented the drop in mitochondrial $NAD^+$ observed in our ROS-exposed model cells, allowing $NAD^+$ dependent *SIRT3* activity to prevent MiDAS (**Fig. 5D**). To test whether *SIRT3* hyper-activation itself could act in a similar way, we simultaneously exposed our model cell to external ROS and forced *SIRT3* activation (**SM Fig. 14A**). Our model predicted that cells with hyperactive *SIRT3* were protected from ROS-induced MiDAS, and that forced *SIRT3* activation could reverse MiDAS (**SM Fig. 14B**).

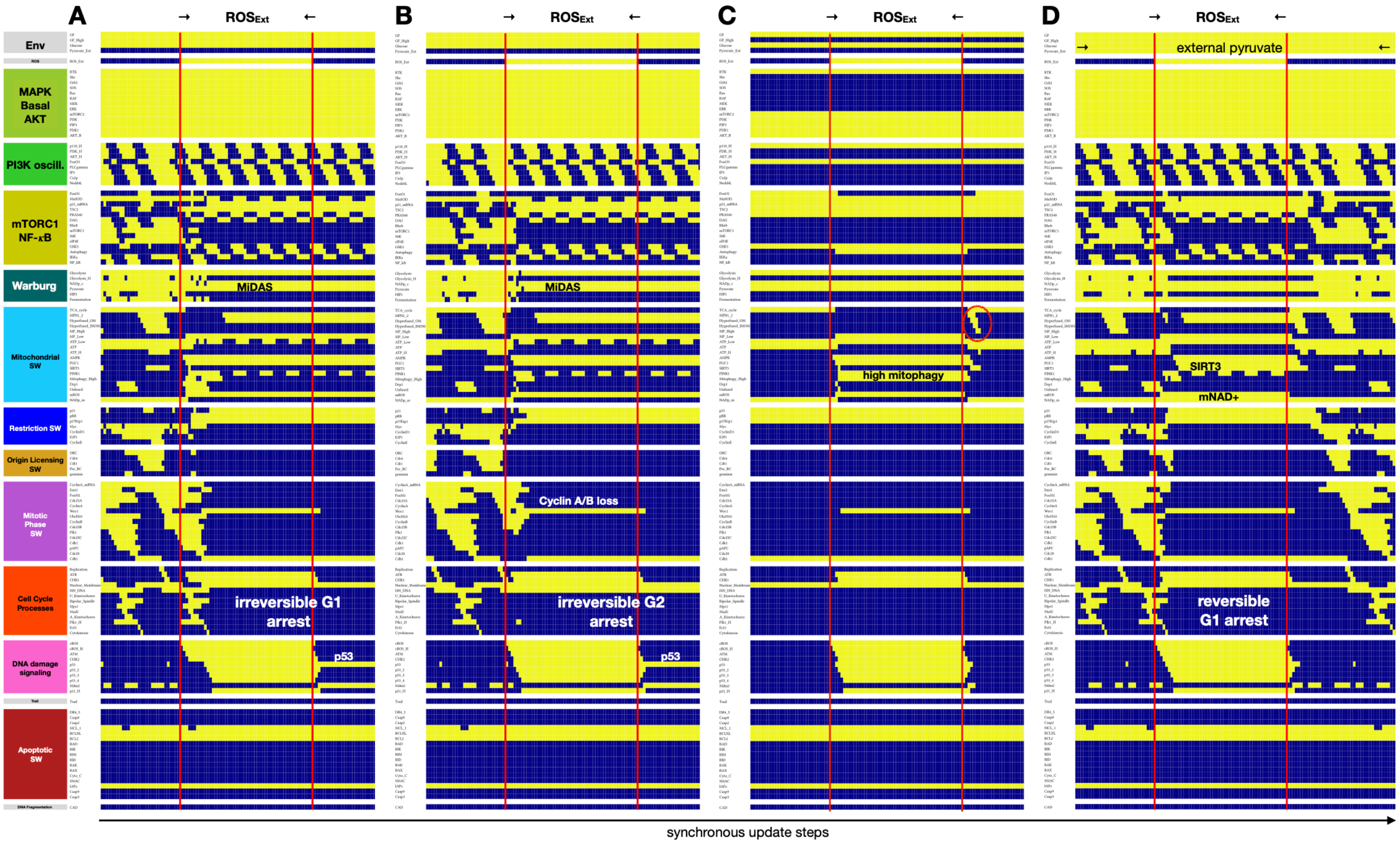


**SM Figure 12. Full version of Fig. 5: Model reproduces ROS-induced MiDAS in cycling cells and predicts protection from MiDAS in external pyruvate. A-B)** Dynamics of regulatory molecule expression/activity during exposure of a cycling cell to external ROS for 50 update steps in (A) prometaphase, leading to cytokinesis followed by MiDAS (2N DNA), vs. (B) early G2, leading to irreversible G2 arrest and MiDAS with 4N DNA. **C)** Dynamics of regulatory molecule expression/activity during exposure of a quiescent cell to external ROS for 50 update steps, leading to high mitophagy and short-term hyperfusion to restore ATP levels. **D)** Dynamics of regulatory molecule expression/activity during exposure of a cycling cell exposed to saturating levels of external pyruvate to external ROS near the SAC, for 50 update steps, leading to reversible G1 arrest (cell cycle entry after external ROS is removed). *X-axis:* time-steps; *y-axis:* nodes organized by regulatory modules; *yellow/dark blue:* ON/OFF; *vertical red lines:* start/end of ROS exposure; *red oval on (C):* brief, homeostatic hyperfusion involving both outer and inner mitochondrial membranes; *white/black labels:* relevant molecular changes or outcomes.

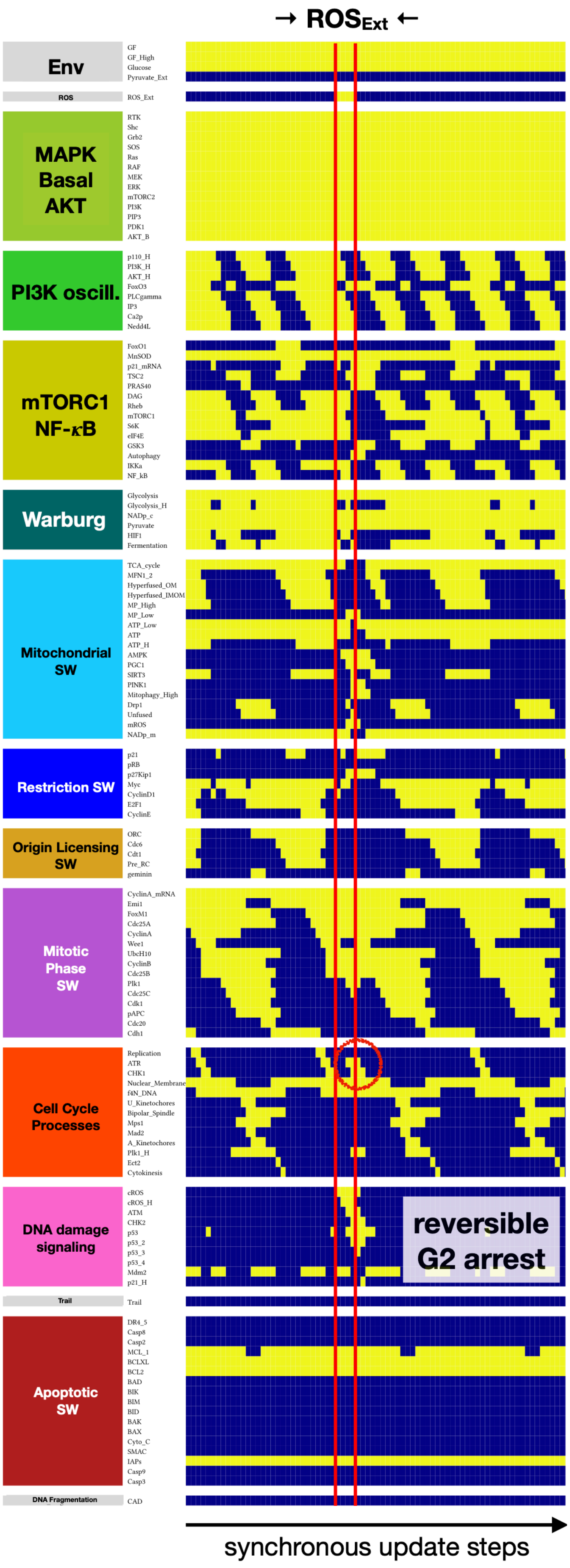


**SM Figure 13. Model reproduces reversible G2 arrest in response to brief / weak ROS exposure.** Dynamics of regulatory molecule expression/activity in cycling cells exposed to a brief (4 update step) pulse of external ROS. *y-axis:* nodes organized in regulatory modules; *yellow/dark blue:* ON/OFF; *vertical red lines:* start/end of high external ROS; *red oval:* prolonged G2, reversible arrest.

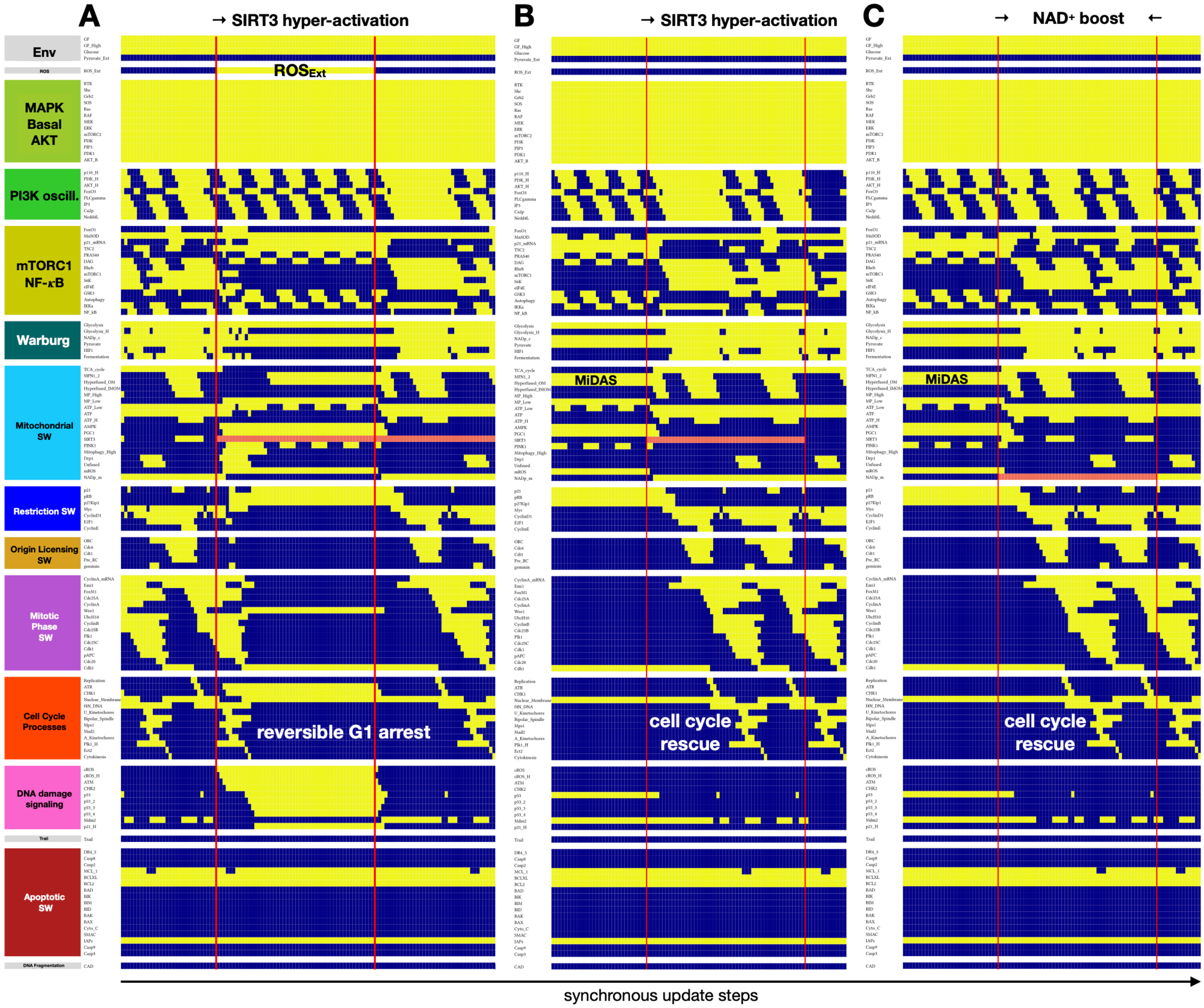


**SM Figure 14. Model predicts that *SIRT3* hyper-activation can protect from or reverse ROS-induced MiDAS, and reproduces MiDAS rescue by NAD+ boost. A)** Dynamics of regulatory molecule expression/activity in cycling cells exposed to external ROS (50 update steps) at the same time as *SIRT3* hyper-activation. **B-C)** Dynamics of regulatory molecule expression/activity in mitogen-exposed MiDAS cells in response to (B) *SIRT3* hyper-activation (50 update steps); (C) a boost in mitochondrial $NAD^+$ (50 update steps). *X-axis:* time-steps; *y-axis:* nodes organized in regulatory modules; *yellow/dark blue:* ON/OFF; *pink:* forced activation; *vertical red lines:* start/change in perturbation; *white/black labels:* relevant outcomes.

### 5. Results, section 5 - Context-dependence of MiDAS

<u>Relevant SM Figures:</u>

- **SM Figure 15.** Model produces a heterogeneous mix of cell phenotypes modulated by mitogen, glucose, ROS, and pyruvate exposure.
- **SM Figure 16.** Full repertoire of model attractors organized by the environmental input combinations in which they are stable.

Loss of glucose sends healthy quiescent cells into a state characterized by energy stress with active *AMPK*, low $\Delta\Psi_M$, high mitophagy, and no hyperfusion (**SM Fig. 15**, *bottom plane, <u>no pyruvate,</u> green arrow* to *yellow-bordered cell*). Cycling cells, on the other hand, generally arrest in a state with reversible mitochondrial hyperfusion, distinguished from MiDAS by *SIRT3* mediated control of ROS paired with $NAD^+$ maintenance (*red arrow* to *orange-bordered cell;* **Fig. 3A**), Indeed, this state is reversible by glucose exposure — either during cell cycle re-entry (*bidirectional red arrow*), or quiescence.

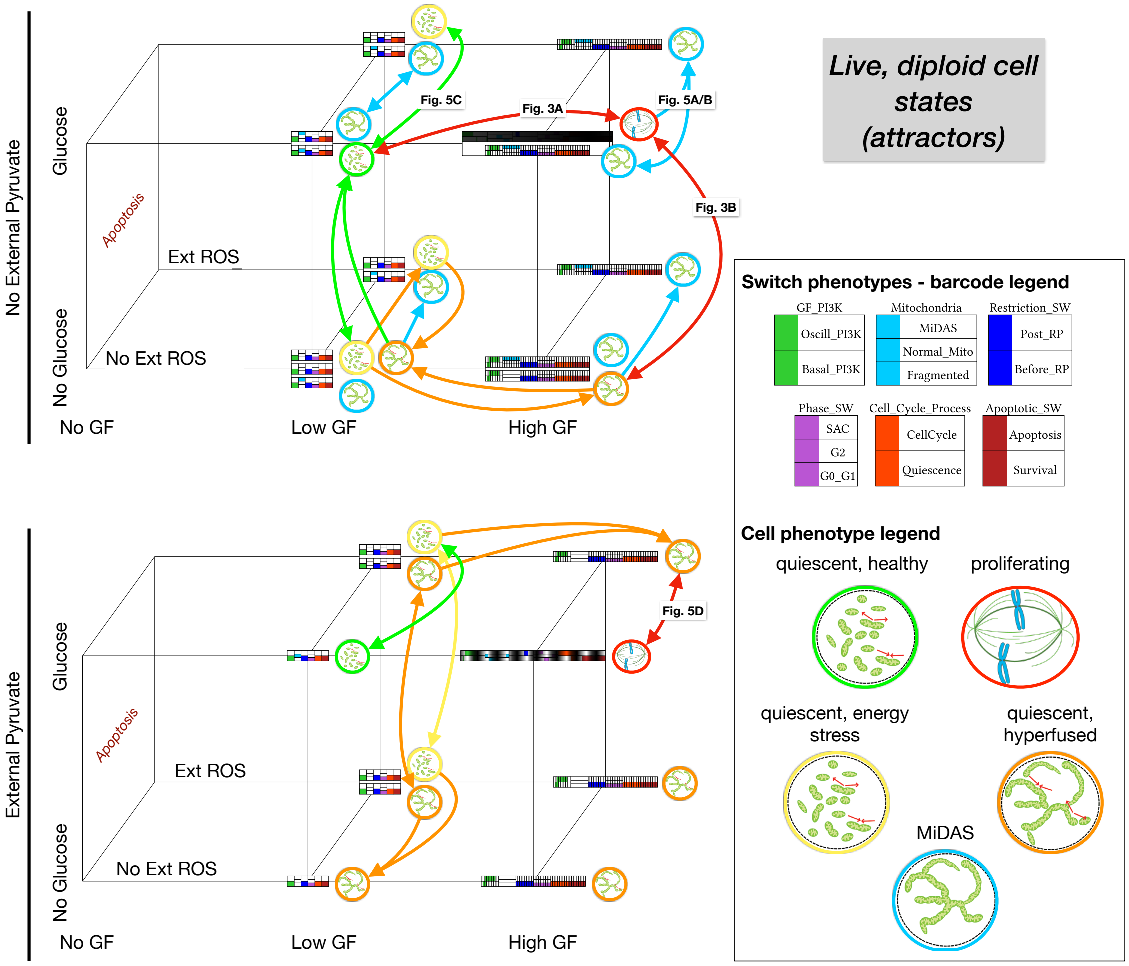


**SM Figure 15. Model produces a heterogeneous mix of cell phenotypes modulated by mitogen, glucose, ROS, and pyruvate exposure. A)** Summary of model cell states detected in every combination of no/low/high growth-factor (*x axis*), absence/presence of external ROS (*y axis*), low/high glucose (*z axis*), and absence/presence of external pyruvate (*left/right*). Apoptotic and tetraploid quiescent cell states were omitted for clarity (all attractors: **SM Fig. 16**. Barcodes representing each attractor were derived by comparing the expression of nodes in relevant modules to predetermined molecular signatures encoded in the model's *.dmms* file (barcode legend, *bottom left*). Oscillatory phenotypes have expanded barcodes that mark the transitions their regulatory switches undergo during the cycle (*left, high GF area*). Visual summaries of overall cell states indicate the shape and dynamics of the mitochondrial network, distinguish quiescent vs. cycling cells, as well as normal vs. dysfunctional mitochondria (cell phenotype legend, *bottom right*). *Figure labels:* time-courses with molecule-level view of state. *State transition arrows: light blue:* irreversible transition to MiDAS; *green:* reversible transitions from quiescence with healthy mitochondria & transition to it; *orange:* transitions between energy stressed non-MiDAS states; *red:* reversible cell cycle arrest upon glucose withdrawal.

Quiescent cells recovering from ROS exposure in the absence of glucose (**Fig. 15**, *bottom plane, no pyruvate*), get stuck with a hyperfused state that is not quite MiDAS (*orange-outlined cells, bottom plane*), given that subsequent glucose exposure can reestablish healthy mitochondria (*green up-arrow, front plane*). This protection is fragile however, as re-exposure to ROS in this state triggers MiDAS, regardless of growth signals strength (*light blue arrows, bottom plane*). Interestingly, lack of glucose but access to external pyruvate (*bottom plane, pyruvate*) forces this reversible hyperfusion in all but high ROS/low growth factor conditions, and loss of pyruvate in high growth factors triggers MiDAS (*arrow not shown*).

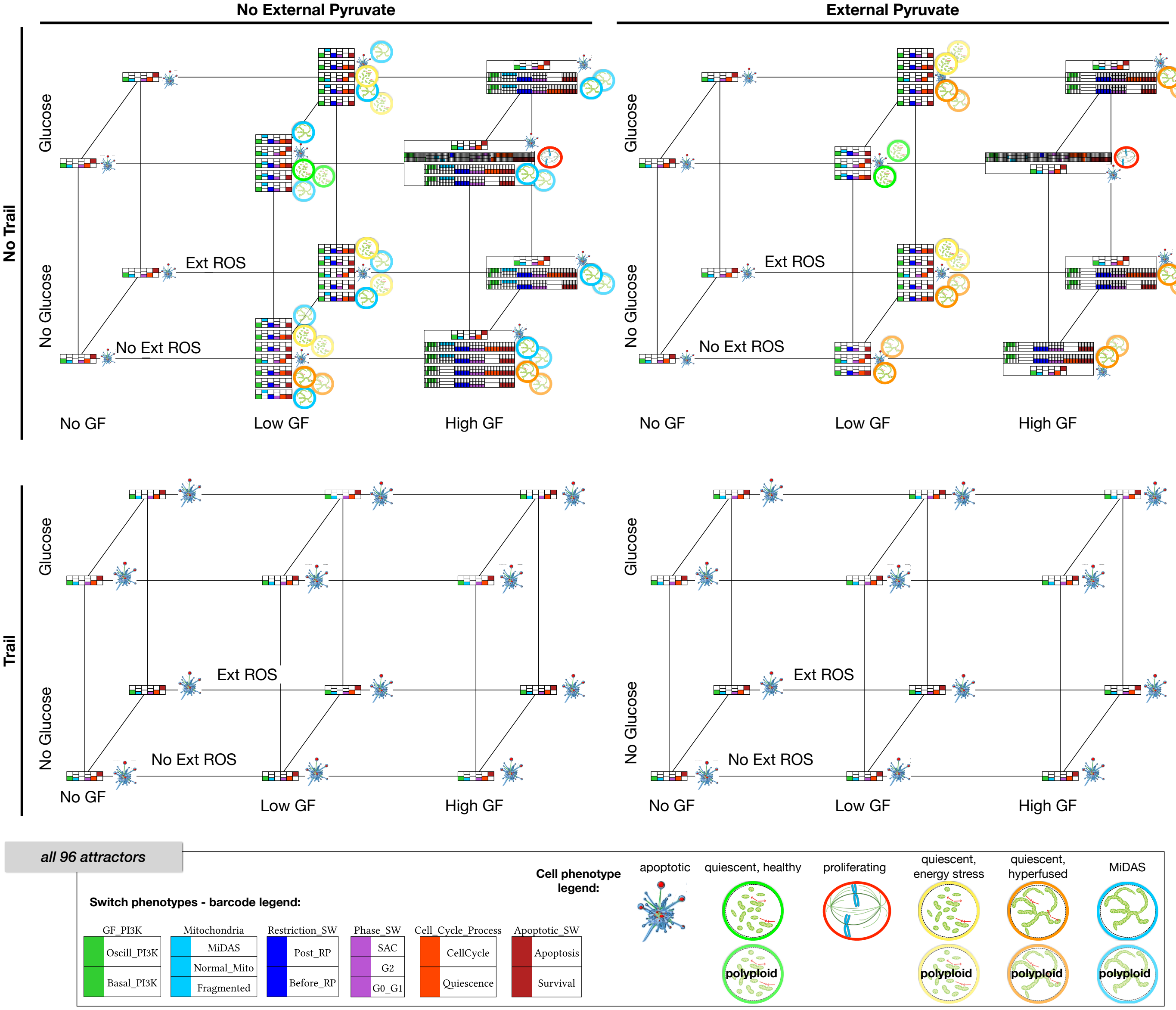


**Figure 16. Full repertoire of model attractors organized by the environmental input combinations in which they are stable. A)** Model cell states detected in every combination of no/low/high growth-factor (*x axis*), absence/presence of external ROS (*y axis*), low/high glucose (*z axis*), absence/presence of external pyruvate (*left/right*), and absence/presence of Trail (*top/bottom*). Barcodes representing each attractor were derived by comparing the expression of nodes in each relevant module to a pre-determined molecular signature known to represent a cell phenotype (e.g., apoptosis vs. survival) and encoded in the model's *.dmms* file (barcode legend, *bottom left*). Visual summaries of overall cell states indicate the shape and dynamics of the mitochondrial network, distinguish quiescent vs. cycling vs. apoptotic cells, normal vs. dysfunctional vs. fragmented mitochondria, and wild-type vs. tetraploid quiescence (cell phenotype legend, *bottom right*). Molecule-level ON/OFF state of the network in each attractor included in **SM File 7**.

### 6. Results, section 6 - MiDAS in cells with cancer-related mutations

Relevant SM Figures and Files:

- **SM Figure 17.** *p53* loss blocks MiDAS in response to mild/strong ROS in mitogen-stimulated cells.
- **SM File 14**, included as *MiDAS in cancer-associated mutants.pdf*: Simulation results for all cancer-related mutant cells for every combination of *GF_High*, *Glucose*, and *ROS_Ext* at 0, 5%, 25%, 50%, and 95% saturation with 0% and 50% external pyruvate (100% pyruvate blocked MiDAS, an affect unchanged by any of our perturbations; data not shown but can be generated with commands in **SM File 11**). *Hyper-activated nodes:* $PI3K_H$, Ras, RAF, mTORC1, $AKT_H$, Myc, HIF1, MEK, Cyclin E, Cyclin D1, ERK, $p21_H$, $p27^{Kip1}$; *knockouts:* SIRT3, p53, ATM, ATR, RB, p21, TSC2, FoxO1, FoxO3, Caspase 8, Caspase 9, Plk1 (results ordered by effect).

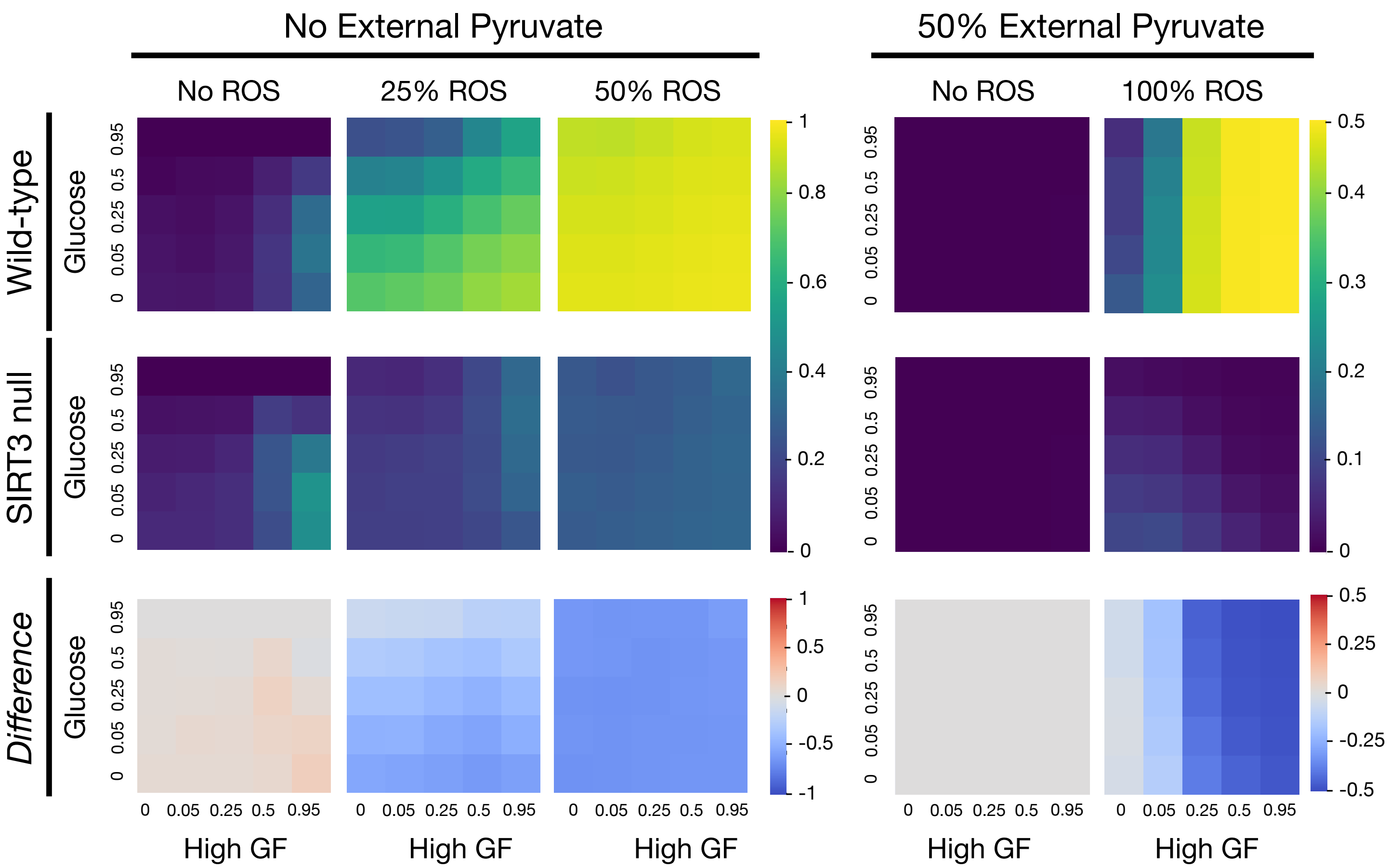


**SM Figure 17.** ***p53*** **loss blocks MiDAS in response to mild/strong ROS in mitogen-stimulated cells.** Fraction of time cells spend in a MiDAS state at varying levels of mitogen stimulation (*x* axis) and glucose (*y* axis) with no (0%), mild (5%) or strong (50% / 95%) ROS exposure, in the absence (left 3 columns) vs. presence of 50% external pyruvate (right 2 column). *Top/middle row:* wild-type/*p53* null cells; *bottom row:* increase in the fraction of time spent in MiDAS in *p53*-null cells. *Length of time-window for continuous runs:* 100 steps (10 wild-type cell cycle lengths); *total sampled time:* 500,000 steps; synchronous update; *initial state for sampling runs:* cycling cell in high glucose and no external pyruvate, ROS, or *Trail*.

### 7. Results Reproduced with Biased Asynchronous Update

Relevant SM Figures and Files:

- **SM Figure 18.** PI3K oscillations correspond to complex attractors, while the cell cycle is a metastable oscillation under asynchronous update (asynchronous version of Fig. 3A).
- **SM Figure 19.** Asynchronous version of Fig. 3, ensemble average (model reproduces cell cycle-linked mitochondrial dynamics and G1 arrest in response to glucose withdrawal).
- **SM Figure 20.** Asynchronous version of Fig. 4B,D (model reproduces MiDAS in response to SIRT3 knockdown).
- **SM Figure 21.** Asynchronous version of Fig. 5A-B (model reproduces ROS-induced MiDAS in cycling cells).
- **SM Figure 22.** Asynchronous version of Fig. 5C-D (model predicts protection from MiDAS in quiescence or in external pyruvate).

*Comparison of synchronous vs. asynchronous attractors.* *Dynmod* detected 96 synchronous attractors: 78 fixed points, 16 cycles involving PI3K oscillations at high growth factor, and two cell cycles (with/without external pyruvate). AEON (https://github.com/sybila/biodivine-aeon-py) [63], the fastest asynchronous attractor detection algorithms we are aware of, is guaranteed to find *all* fixed-point and complex attractors under general asynchronous update for any network it can complete its algorithm on in a reasonable time [67]. This was indeed the case for our model, confirming that the only fixed points and complex attractors of this network match the 78 fixed points and 16 PI3K oscillations found by *dynmod* (**SM Fig. 17A**). We have previously shown that our cell cycle regulatory modules generate a meta-stable circular "valley" in state space akin to a 'Mexican hat' landscape, which can temporarily trap the asynchronous dynamics to mimic the cell cycle (SM. Fig. 5 in [45]). It is not a real trap space however, as there are a small number of update-orders that send the model into an apoptotic fixed point, or in the current model, also into MiDAS. While this occurs too often with random-order asynchronous update to be realistic, the paths the model follows to apoptosis and its 'shortcuts through the ring' mimic known cell cycle errors [45]. By contrast, limit cycles involving oscillations driven by the PI3K module are robust to asynchronous update (see [45] for details).

*Robustness of modeling results to asynchronous update.* In order to test whether the model can generate qualitatively and thus biologically similar responses to signal under synchronous vs. asynchronous update, we re-ran our simulations with a biased version of random-order asynchronous update. Namely, we randomized the update order of nodes in each time-step, but moved a small, select set nodes to the start and/or end of the update order, depending on their current state (encoded in the .dmms file in the model's *Metadata* block as follows; red is new since [45]):

```
BiasOrderFirst: (Cytokinesis, 0), (Pre_RC, 1), (Replication, 0), (U_Kinetochores, 0),
(A_Kinetochores, 1), (Plk1_H, 1), (CyclinB, 1), (Cdc20, 0)
BiasOrderLast: (Replication, 1), (f4N_DNA, 0), (f4N_DNA, 1), (Ect2, 0), (A_Kinetochores, 0),
(U_Kinetochores, 1), (FoxM1, 1), (CyclinE, 1), (Cdc20, 1), (Plk1_H, 0), (MP_Low, 0)
```

This change prevents a series of unrealistic node orders from occurring, and increases the stability of the metastable cell cycle trap region. The result is a drop in the incidence of cell cycle errors, apoptosis and MiDAS, due to unrealistic breaks in signal propagation. For example, rerunning Fig. 3A with an ensemble of 10,000 independently simulated biased asynchronous update time-courses generates individual cell fate tracks that cycle and return to quiescence (**SM Fig. 17B**), end up as polyploid quiescent cells indicating a cell cycle error (**SM Fig. 17C**), undergo apoptosis (**SM Fig. 17D**), or undergo MiDAS (**SM Fig. 17E**). Yet, the average time course of the entire ensemble (**SM Fig. 18A**), as well as the time spent in each phenotype of relevant switches (**SM Fig. 18B**), indicate that most cells return to quiescence with 2N DNA. As expected from [45], the relative rate of events that deviate from a healthy cell cycle is lower with biased vs. fully random-order update (**SM Fig. 18C**). Similarly, loss of glucose (asynchronous version of Fig. 3B) results in a reversible G1 arrest with hyperfused mitochondria in the majority of cells (**SM Fig. 18D**), in accordance with Fig. 3B. Yet, the figure also indicates that a small fraction of cells undergoes apoptosis (observed with synchronous update, SM Fig. 6C), or enters MiDAS upon glucose re-exposure.

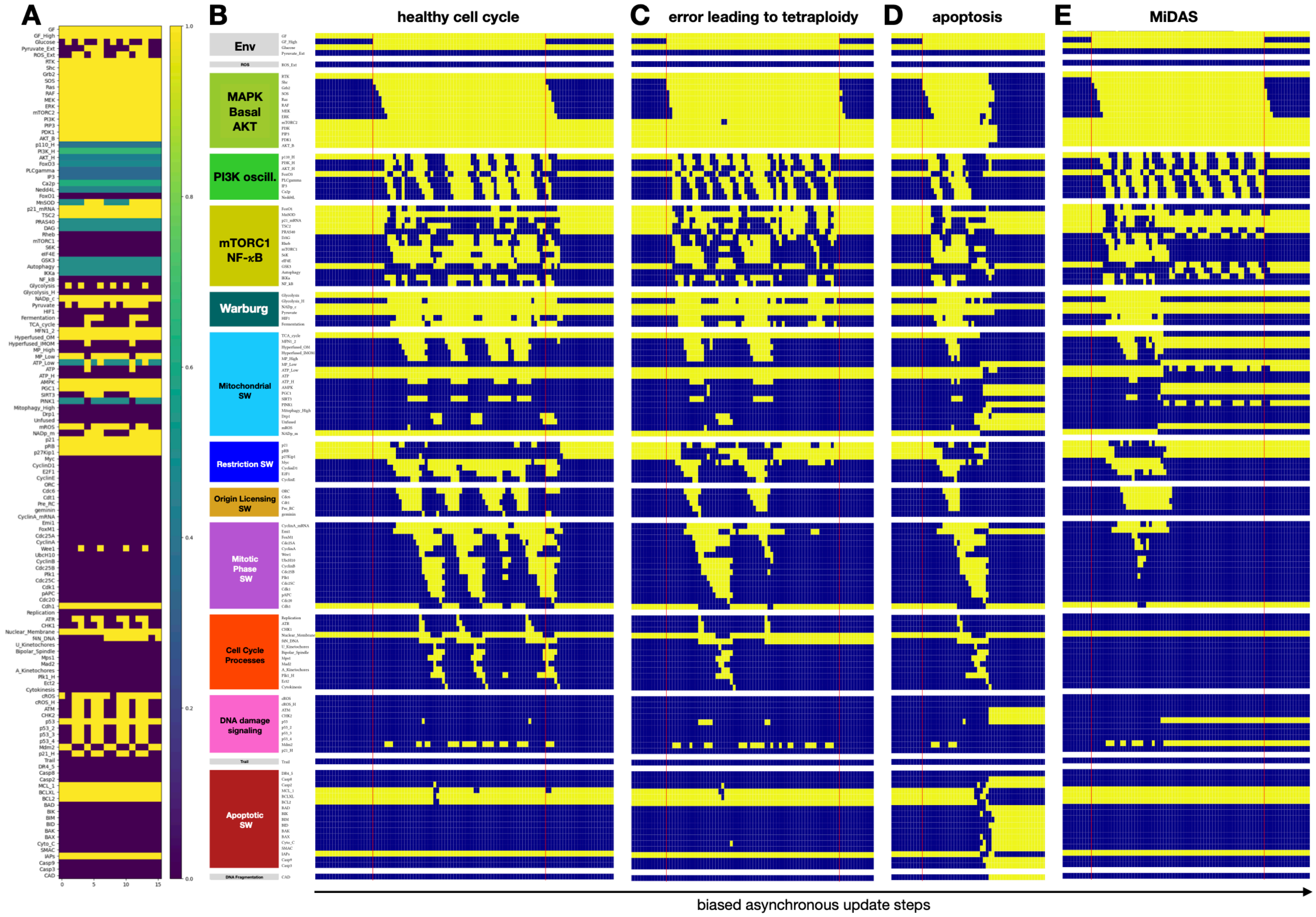


**SM Figure 18. PI3K oscillations correspond to complex attractors, while the cell cycle is a metastable oscillation under asynchronous update (asynchronous version of Fig. 3A). A)** Regulatory molecule expression/activity (*y axis*) averaged over all network states in the trap space of the 16 complex attractors (*x axis*) detected with AEON. *Yellow*: stable ON; purple: stable OFF; *teal*: toggling ON/OFF. **B-E)** Dynamics of regulatory molecule expression/activity in individual quiescent cells responding to strong growth signals (50 update steps) under biased asynchronous update, showing a heterogeneous mix of cell fates: (B) repeated normal cell cycle and return to quiescence; (C) error leading to polyploid cells that can undergo endo-reduplication; (D) spontaneous mitotic catastrophe and apoptosis; (E) spontaneous MiDAS. *X-axis:* biased random-order asynchronous update-steps; *y-axis:* nodes organized by regulatory modules; *yellow/dark blue:* ON/OFF; *vertical red lines:* start/end of strong growth signal.

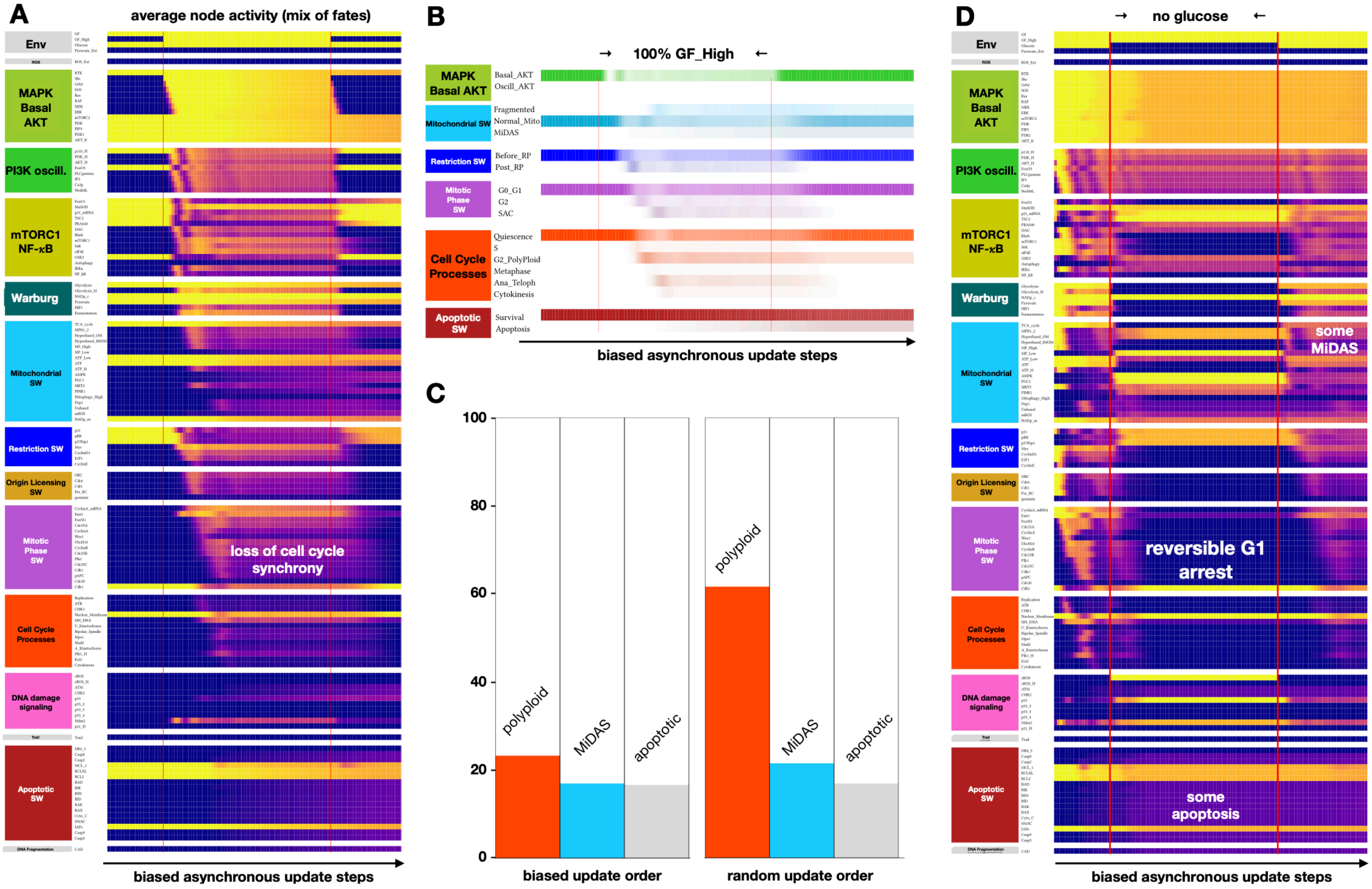


**SM Figure 19. Asynchronous version of Fig. 3, ensemble average (model reproduces cell cycle-linked mitochondrial dynamics and G1 arrest in response to glucose withdrawal). A)** Dynamics of regulatory molecule expression/activity in an ensemble of 10,000 cells responding to strong growth signals (50 update steps) with cell cycle entry, followed by de-synchronization of cell cycle progression and ~15% apoptosis. *X-axis:* biased random-order asynchronous update-steps; *y-axis:* nodes organized by regulatory modules; *yellow to dark blue:* ON to OFF; *vertical red line:* start/end of strong growth signals; **B)** Average number of cells with an internal state matching each regulatory switch-phenotype, as a function of biased random-order asynchronous update-steps in the simulation on (A). **C)** Average ON state of the 4N DNA node (*red*, marking tetraploid cells), the Hyperfused_OM node (*light blue*, marking MiDAS cells), and the CAD node (*gray*, marking apoptotic cells), averaged over 10,000 cells as well as a 50 time-step window following the removal of strong growth signals (last interval on A). **D)** Dynamics of regulatory molecule expression/activity in an ensemble of 10,000 dividing cells responding to glucose withdrawal (50 update steps) and its reversal, showing reversible cell cycle arrest, as well as a small uptick in apoptosis and MiDAS. *X-axis:* biased random-order asynchronous update-steps; *y-axis:* nodes organized by regulatory modules; *yellow to dark blue:* ON to OFF; *vertical red line:* start/end of glucose withdrawal; *labels:* relevant phenotypic changes.

Our main MiDAS results are generally recapitulated with biased asynchronous update:

— **SM Fig. 20:** asynchronous version of Fig. 4 (SIRT3 knockdown-induced MiDAS + some apoptosis)

— **SM Fig. 21**: asynchronous version of Fig. 5A-B (ROS-induced MiDAS following G1 or G2 arrest + some apoptosis)

— **SM Fig. 22**: asynchronous version of Fig. 5C-D (robust but not complete protection from ROS-induced MiDAS in quiescent cells; near-complete protection in cycling cells exposed to external pyruvate).

— **SM Fig. 23**: asynchronous version of Fig. 6B (context-dependent effect of SIRT3 knockdown).

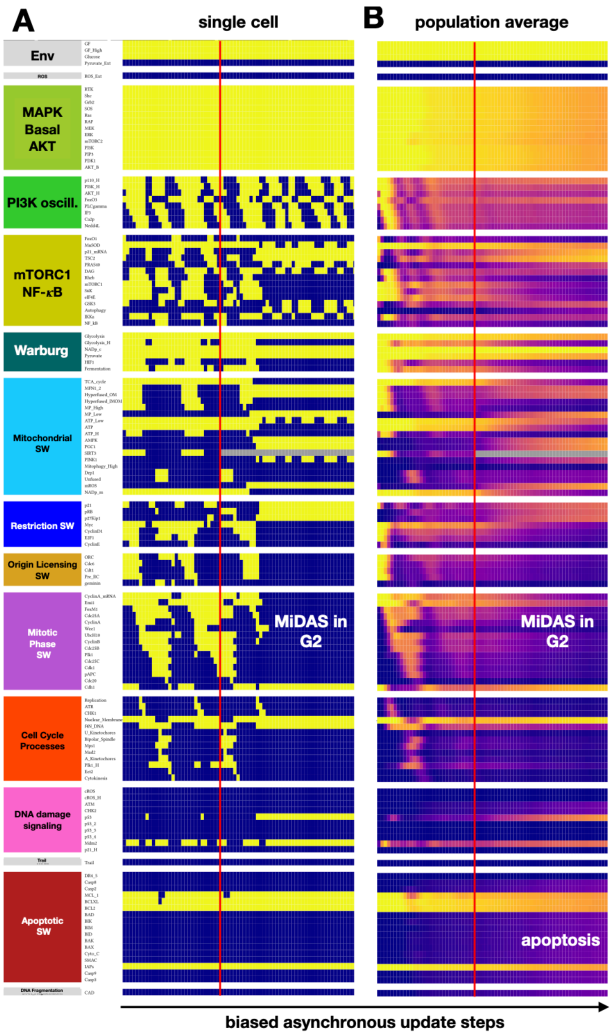


**SM Figure 20. Asynchronous version of Fig. 4B,D (model reproduces MiDAS in response to SIRT3 knockdown).** **A)** Dynamics of regulatory molecule expression/activity in a single dividing cell responding to full *SIRT3* knockout in early metaphase. **B)** Dynamics of regulatory molecule expression/activity in an ensemble of 10,000 asynchronously dividing cells responding to full *SIRT3* knockout. *X-axis:* biased random-order asynchronous update-steps; *y-axis:* nodes organized by regulatory modules; *yellow/dark blue:* ON/OFF; *vertical red line:* start of *SIRT3* knockout; *labels:* relevant molecular or phenotypic changes in response to *SIRT3* knockout, leading to irreversible G2 arrest and MiDAS with a mix of 2N and 4N DNA content, as well as ~35% apoptosis. **C)** Average number of cells with an internal state matching each regulatory switch-phenotype, as a function of biased random-order asynchronous update-steps in the simulation on (B). **D)** Response of cells dividing in 95% saturating growth stimuli to increasing SIRT3 knockdown, quantified as % time spent in each phenotype (*left:* quiescence vs. in cell cycle; *middle*: alive vs. apoptotic; *right*: fragmented vs. normal mitochondria vs. MiDAS), averaged over 1000 independent runs in each consecutive interval of SIRT3 knockdown (*x* axis), shown as stacked bar charts. *update:* biased asynchronous; *initial state for sampling runs:* cycling cell in high glucose and no external pyruvate, ROS, or *Trail*.

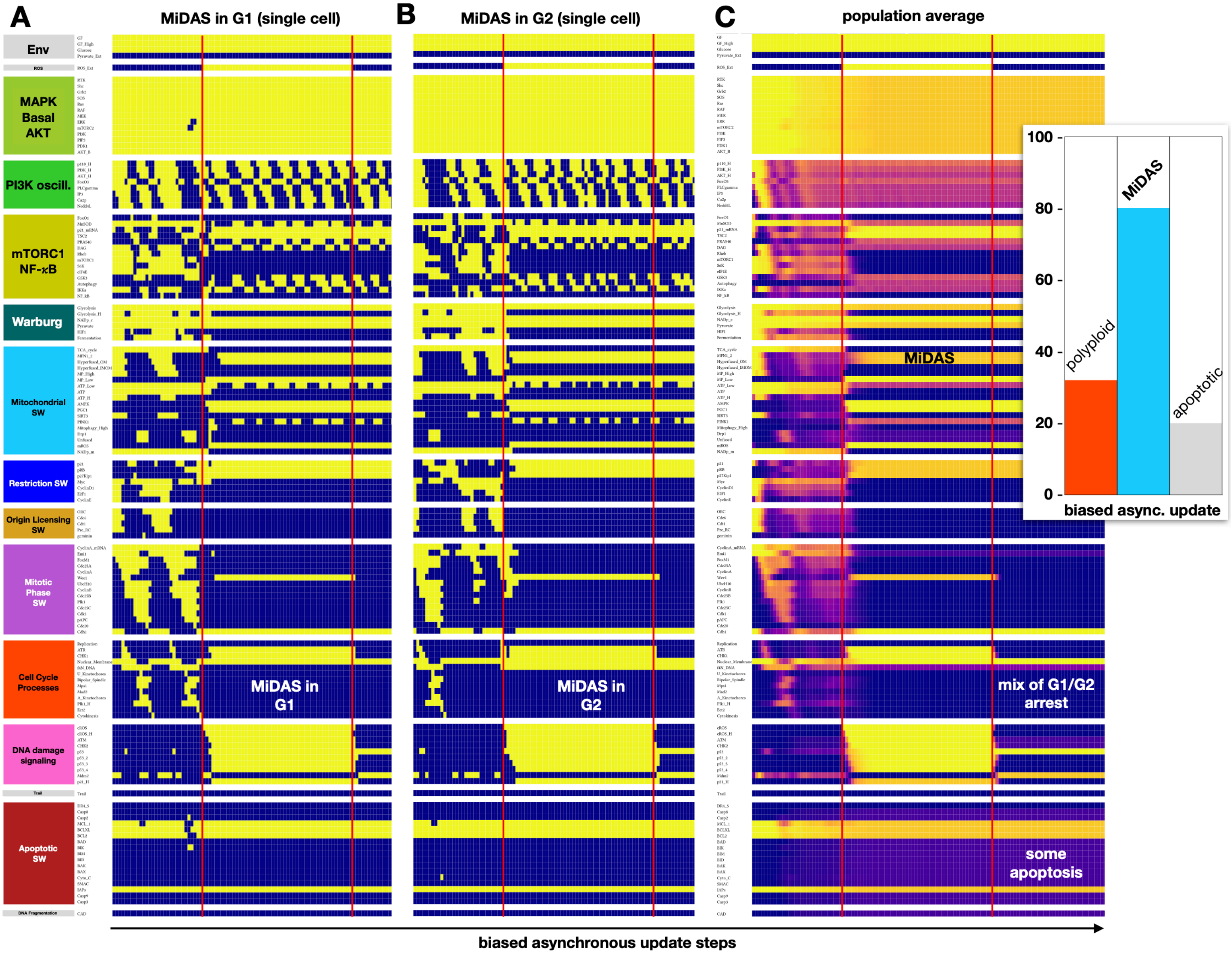


**SM Figure 21. Asynchronous version of Fig. 5A-B (model reproduces ROS-induced MiDAS in cycling cells). A-B)** Dynamics of regulatory molecule expression/activity during exposure of a cycling cell to external ROS in (A) early G1, leading to irreversible G1 arrest and MiDAS with 2N DNA, vs. (B) early G2, leading to irreversible G2 arrest and MiDAS with 4N DNA. **C)** Dynamics of regulatory molecule expression/activity during exposure of an ensemble of 10,000 asynchronously cycling cells to external ROS, leading to irreversible G1/G2 arrest and MiDAS with a mix of 2N (~65%) and 4N (~35%) DNA content, as well as ~20% apoptosis. *Inset:* average ON state of the 4N DNA node (*red*, marking tetraploid cells), the Hyperfused_OM node (*light blue,* marking MiDAS cells), and the CAD node (*gray*, marking apoptotic cells), averaged over 10,000 cells as well as the 50 time-step window following the removal of external ROS (last interval on C).

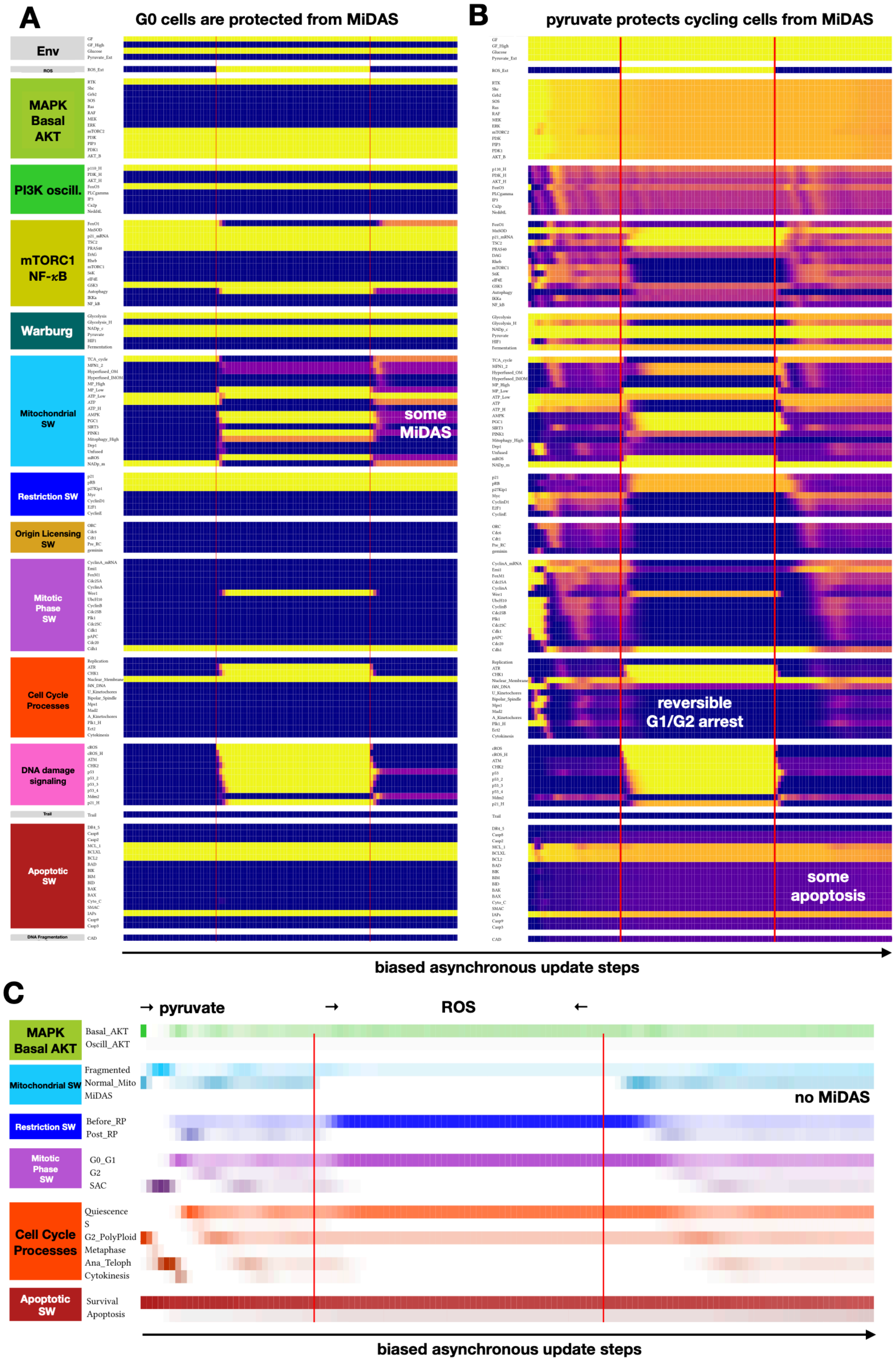


**SM Figure 22. Asynchronous version of Fig. 5C-D (model predicts protection from MiDAS in quiescence or in external pyruvate). A)** Dynamics of regulatory molecule expression/activity in an ensemble of 10,000 quiescent cells during exposure to external ROS, leading to high mitophagy and short-term hyperfusion to restore ATP levels. **B)** Dynamics of regulatory molecule expression/activity during exposure of an ensemble of 10,000 simulated cycling cells in saturating external pyruvate to external ROS, leading to reversible G1/G2 arrest. *X-axis:* biased random-order asynchronous update-steps; *y-axis:* nodes organized by regulatory modules; *yellow/dark blue:* ON/OFF; *vertical red lines:* start/end of ROS exposure; *white/black labels:* relevant molecular changes or outcomes. **C)** Average number of cells with an internal state matching each regulatory switch-phenotype, as a function of biased random-order asynchronous update-steps in the simulation on (B).

#### 8. Model Behavior in Response to Random Network Errors

Relevant SM Figure:

- **SM Figure 23.** Cell cycle and MiDAS in response to SIRT3 knockdown are robust to random mutations and/ or errors in model construction.

We tested the model's robustness to structural errors by generating three distinct ensembles of random mutants:

— **node knockout / hyper-activation:** involved locking a number $n_{\text{Node}}$ of random nodes ON or OFF;

— **link knockout:** removing a number $n_{\text{Link}}$ of randomly chosen links;

— **gate error:** introducing a number $n_{\text{Gate}}$ of random errors in gate output by flipping the expected output of a node for a single combination of inputs.

Averaging the dynamical behavior of the resulting mutant ensembles in a simulation that started with dividing cells, then entered MiDAS due to the loss of SIRT3, allowed us to probe the network's structural robustness during cell cycle (before SIRT3 knockdown) as well as its ability to enter / maintain MiDAS (after SIRT3 knockdown). In other words, we tested how Fig. 4B would change in our mutant ensembles. **SM Fig. 24A** indicates that our model can tolerate 1-2 full node knockout / hyper-activation mutations/cell before both the cell cycle and MiDAS from cell cycle becomes hard to detect in a population. It tolerates link removals best, starting to loose both cycling cells and the MiDAS phenotype by 15 link removals/ cell (**SM. Fig. 24B**). Altering the output of gates is similar but more potent; by the time 15 random nodes are responding incorrectly to a single combination of their inputs, the ensemble looses most of its cycling cells, some apoptosis appears and most cells show energy stress with active AMPK (**SM Fig. 24C**).

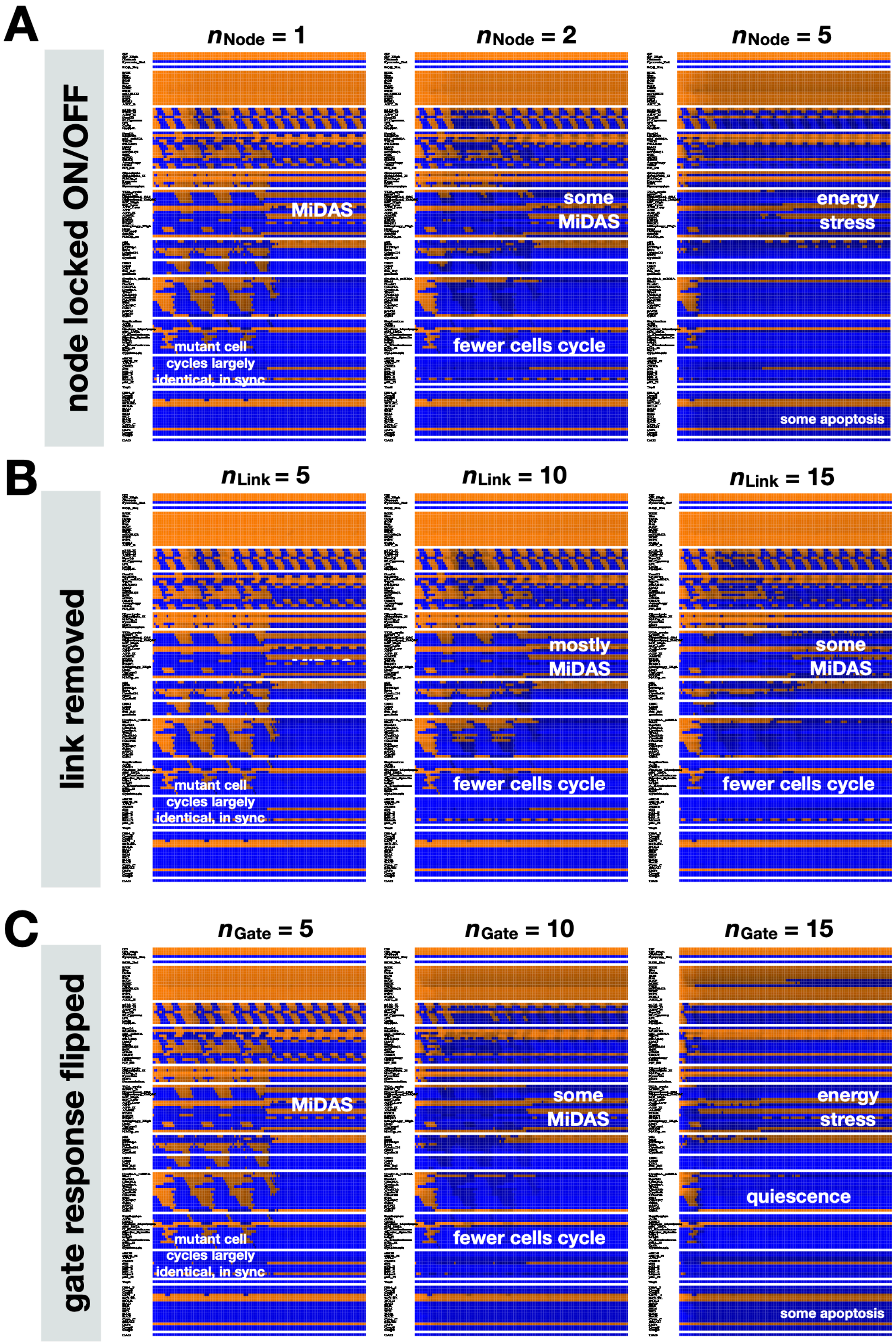


**SM Figure 23. Cell cycle and MiDAS in response to SIRT3 knockdown are robust to random mutations and/or errors in model construction.** Synchronous dynamics of regulatory molecule expression/activity during SIRT3 knockdown in an ensemble of growth-stimulated, cycling cells, averaged over 1000 distinct mutant networks with **A)** $n_{\mathrm{Node}} \in \{1,2,5\}$ random nodes per network locked ON or OFF; **B)** $n_{\mathrm{Link}} \in \{2,5,15\}$ random links per network removed; **C)** $n_{\mathrm{Gate}} \in \{2,5,15\}$ random gate outputs per network flipped. *X-axis*: synchronous update steps; *y-axis*: nodes organized in regulatory modules; *orange/black/blue color-scale*: average expression of each molecule across 1000 time-courses from independently generated mutant models (orange = all ON; black = 50% ON/OFF; blue = all OFF); synchronous update; black/white labels: relevant molecular patterns.
