## Supplementary material for "Unlocking Mitochondrial Dysfunction-Associated Senescence (MiDAS) with NAD^+^ – a Boolean Model of Mitochondrial Dynamics and Cell Cycle Control": SM Files: SM File 13.pdf

### MiDAS: WT (top) vs. SIRT3 KD - WT (bottom)

**TS loss promotes MiDAS**

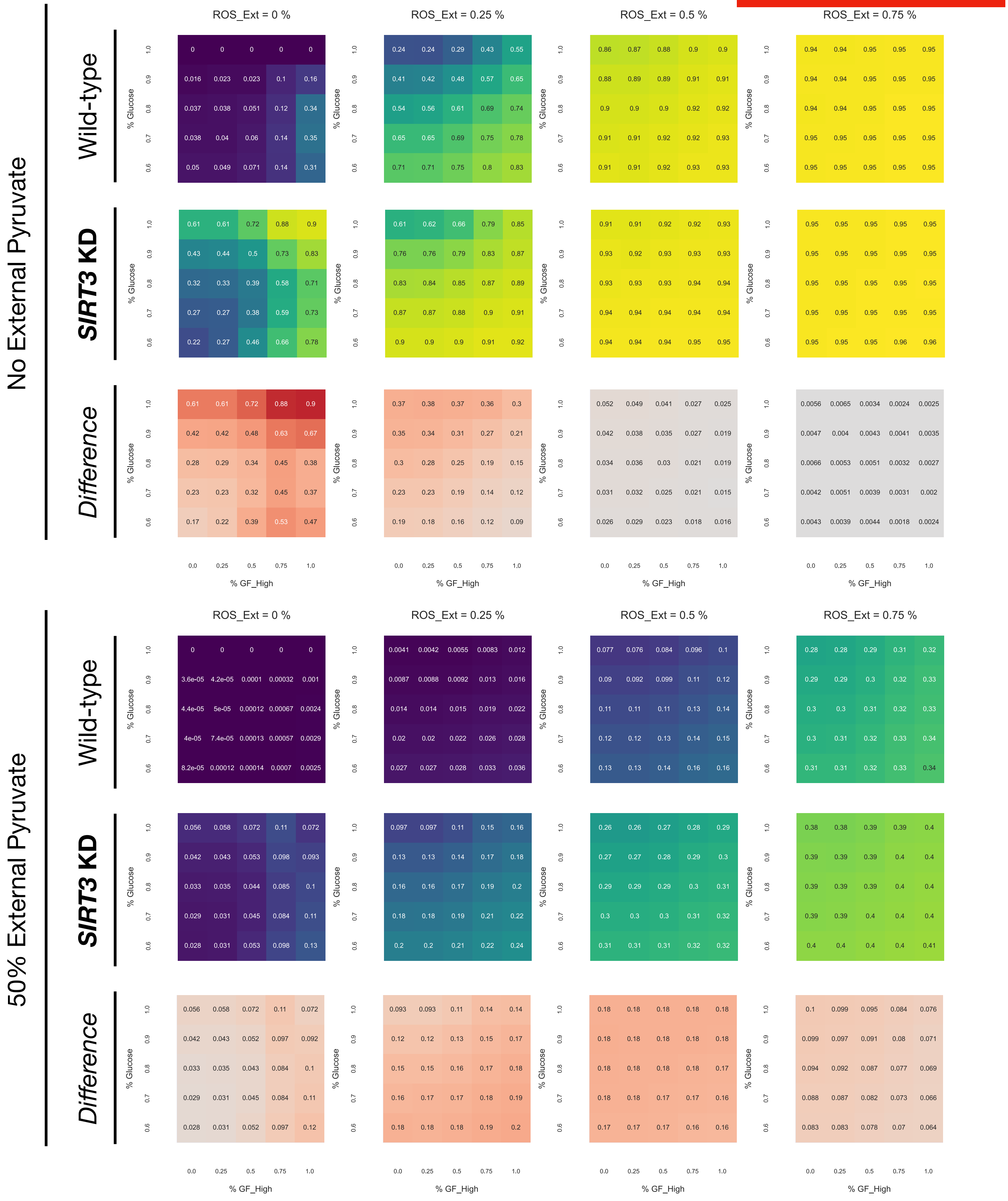

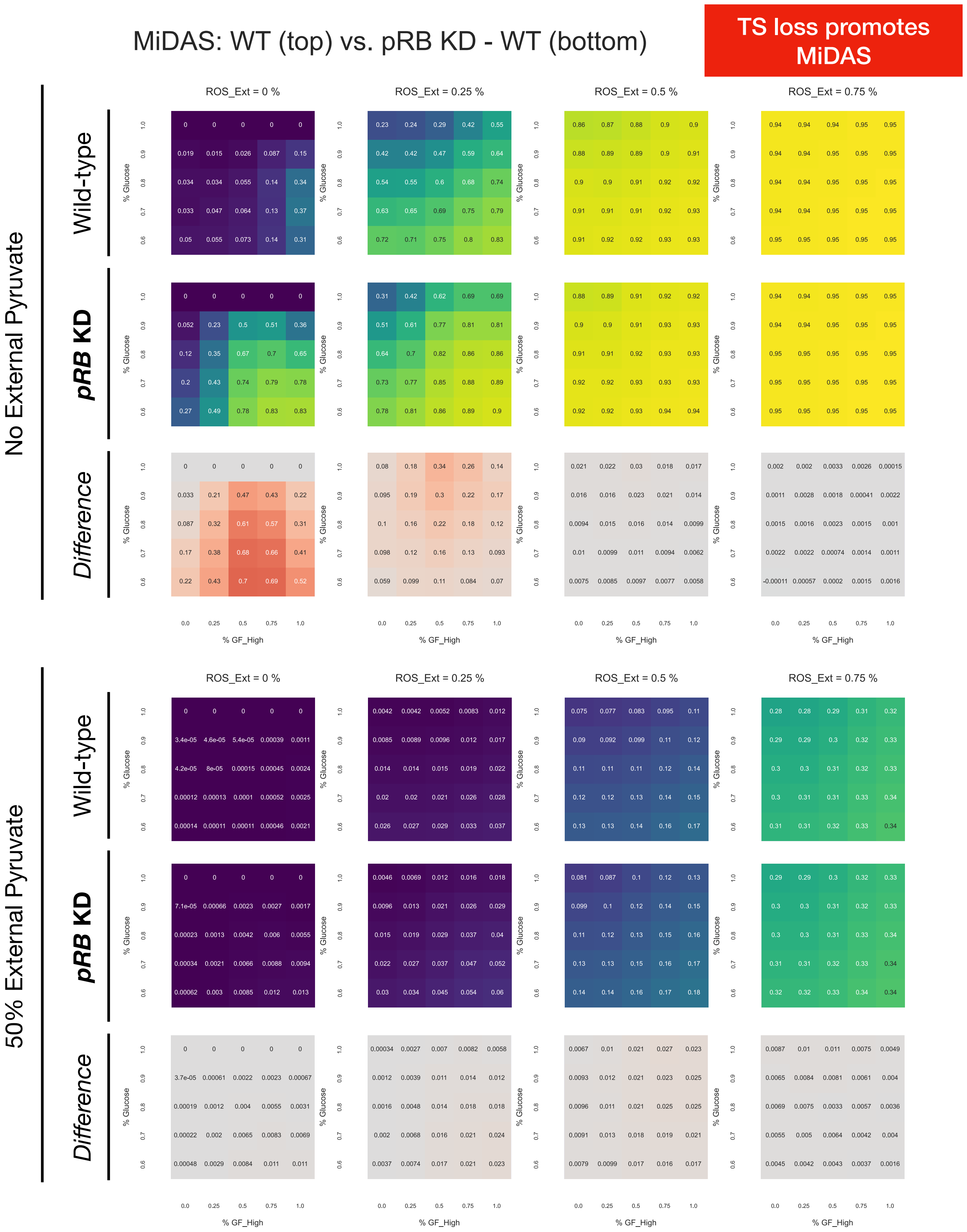
MiDAS: WT (top) vs. pRB KD - WT (bottom)
TS loss promotes MiDAS
No External Pyruvate
50% External Pyruvate
Wild-type
pRB KD
Difference
ROS_Ext = 0 %
ROS_Ext = 0.25 %
ROS_Ext = 0.5 %
ROS_Ext = 0.75 %
% Glucose
% GF_High

MiDAS: WT (top) vs. CyclinD1 OE - WT (bottom)

oncogene promotes MiDAS

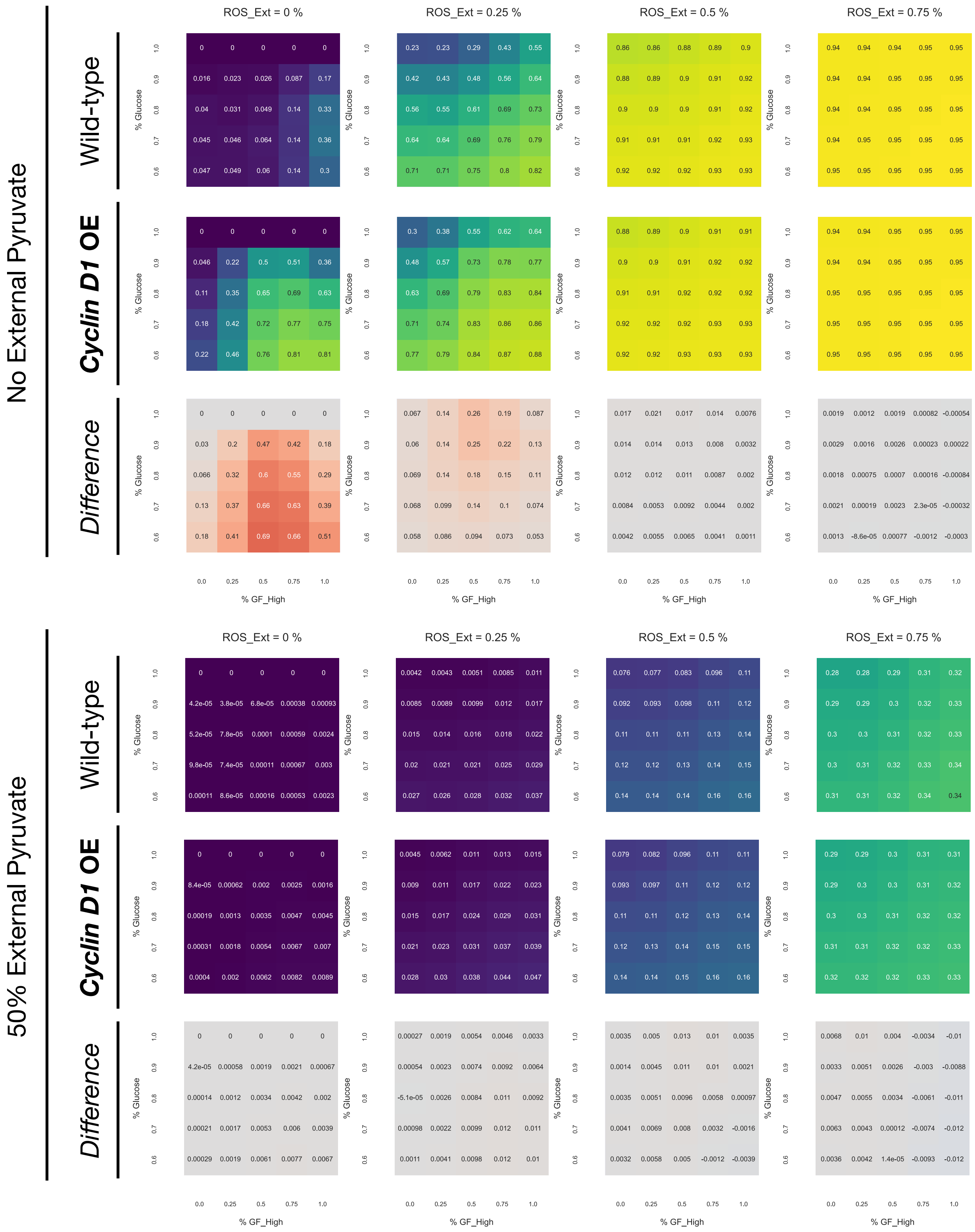

### MiDAS: WT (top) vs. p21 KD - WT (bottom)

**TS loss promotes MiDAS**

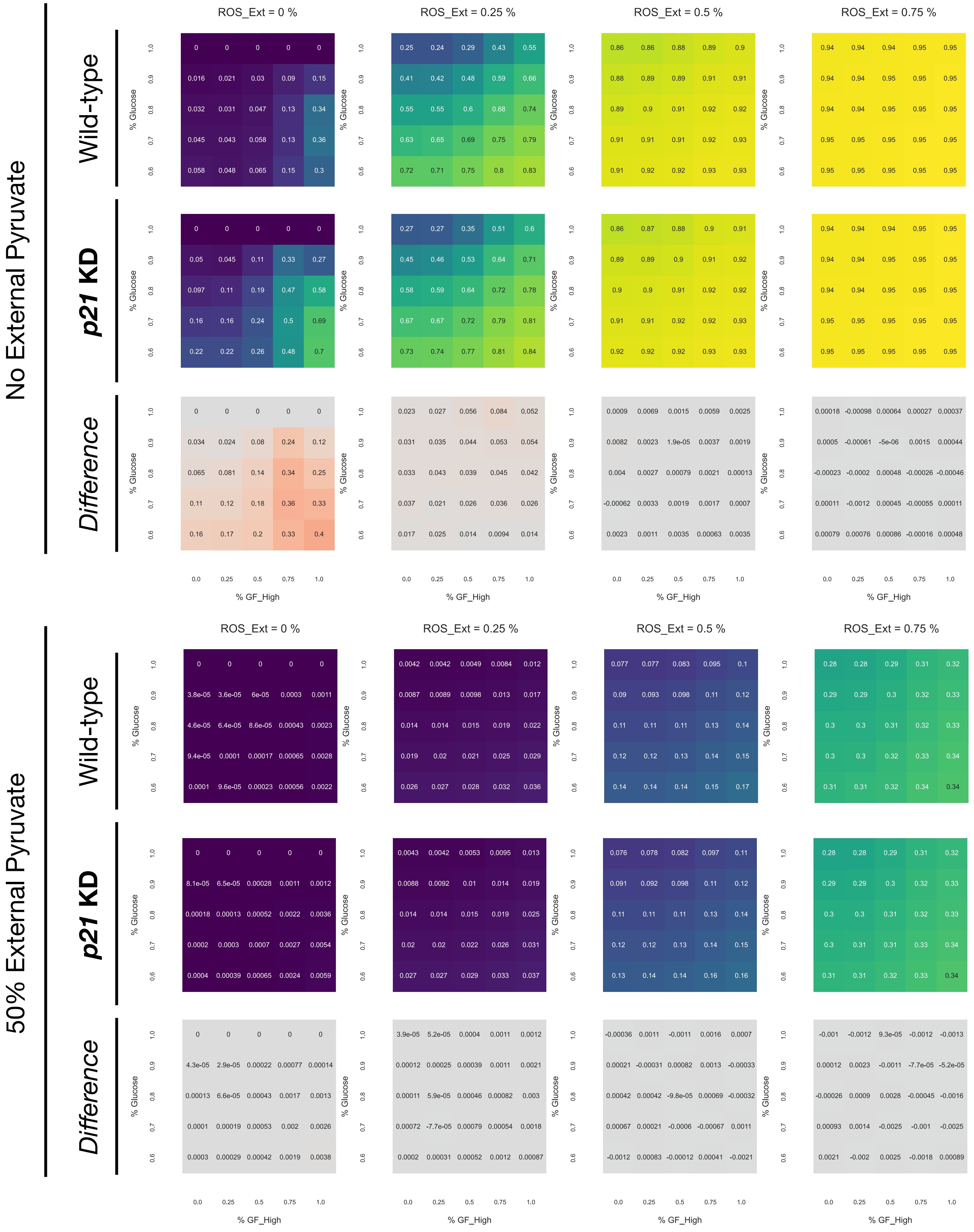

### MiDAS: WT (top) vs. Ras OE - WT (bottom)

**oncogene promotes MiDAS**

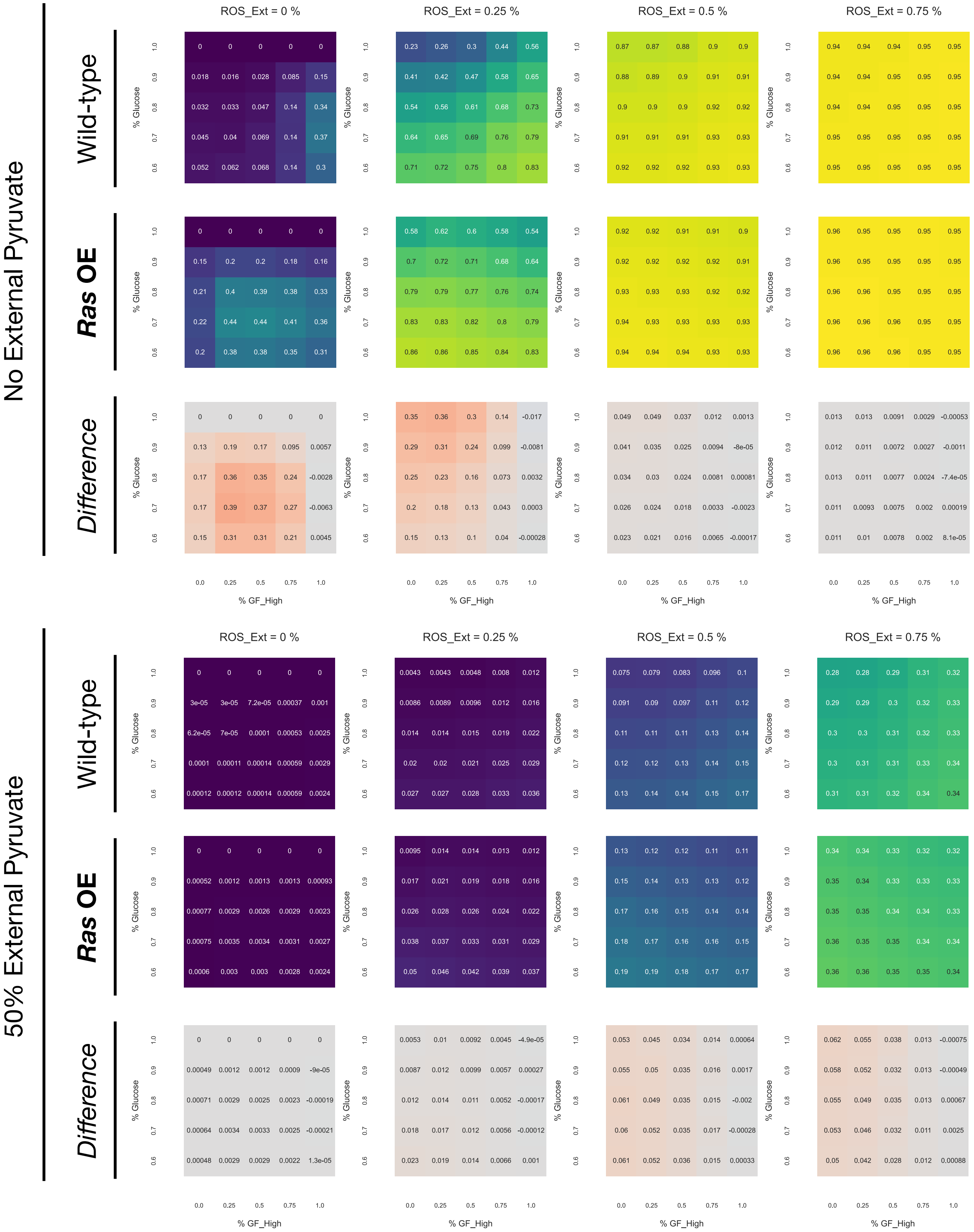

### MiDAS: WT (top) vs. AKT_H OE - WT (bottom)

**oncogene promotes MiDAS**

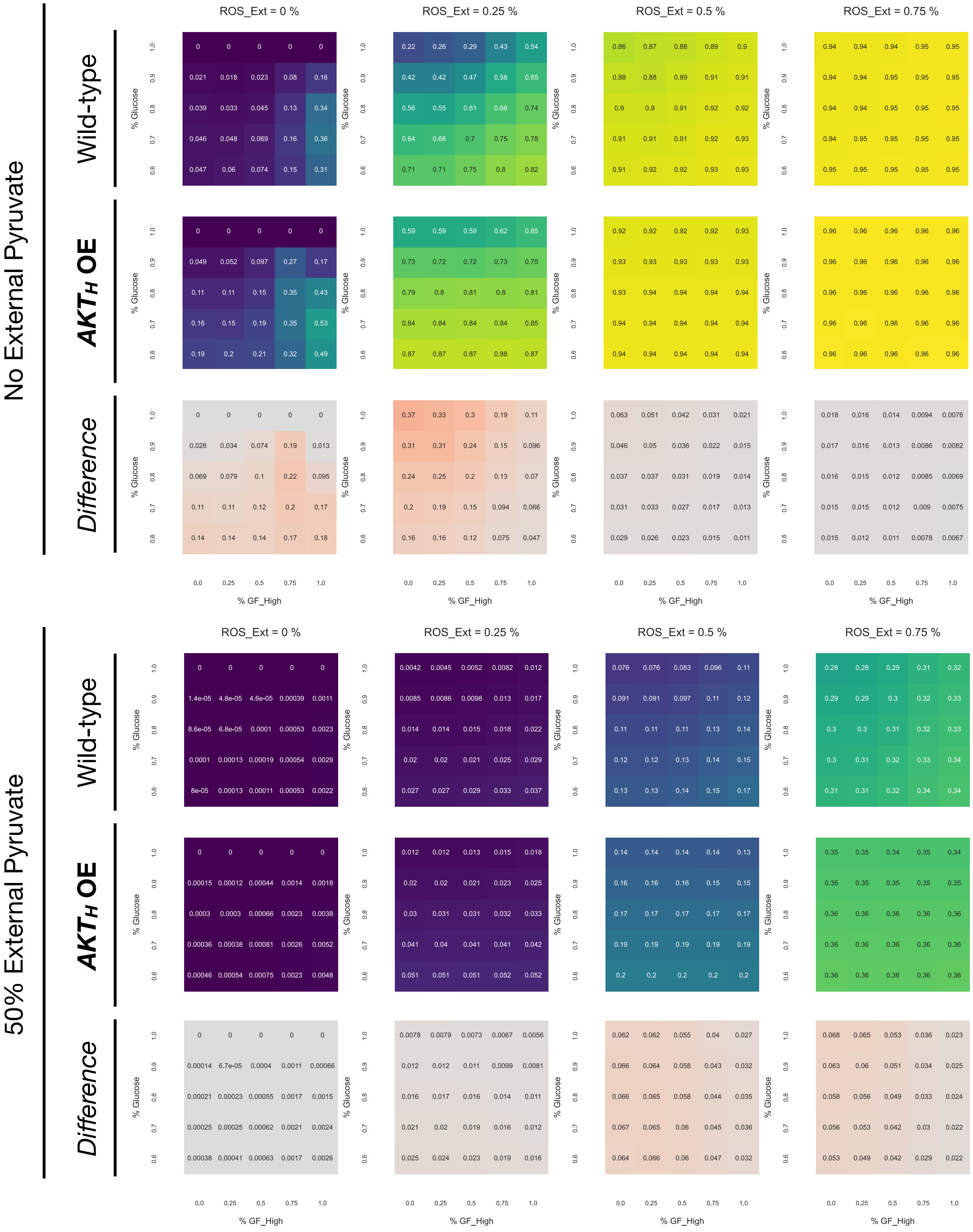

### MiDAS: WT (top) vs. FoxO3 KD - WT (bottom)

**TS loss promotes MiDAS**

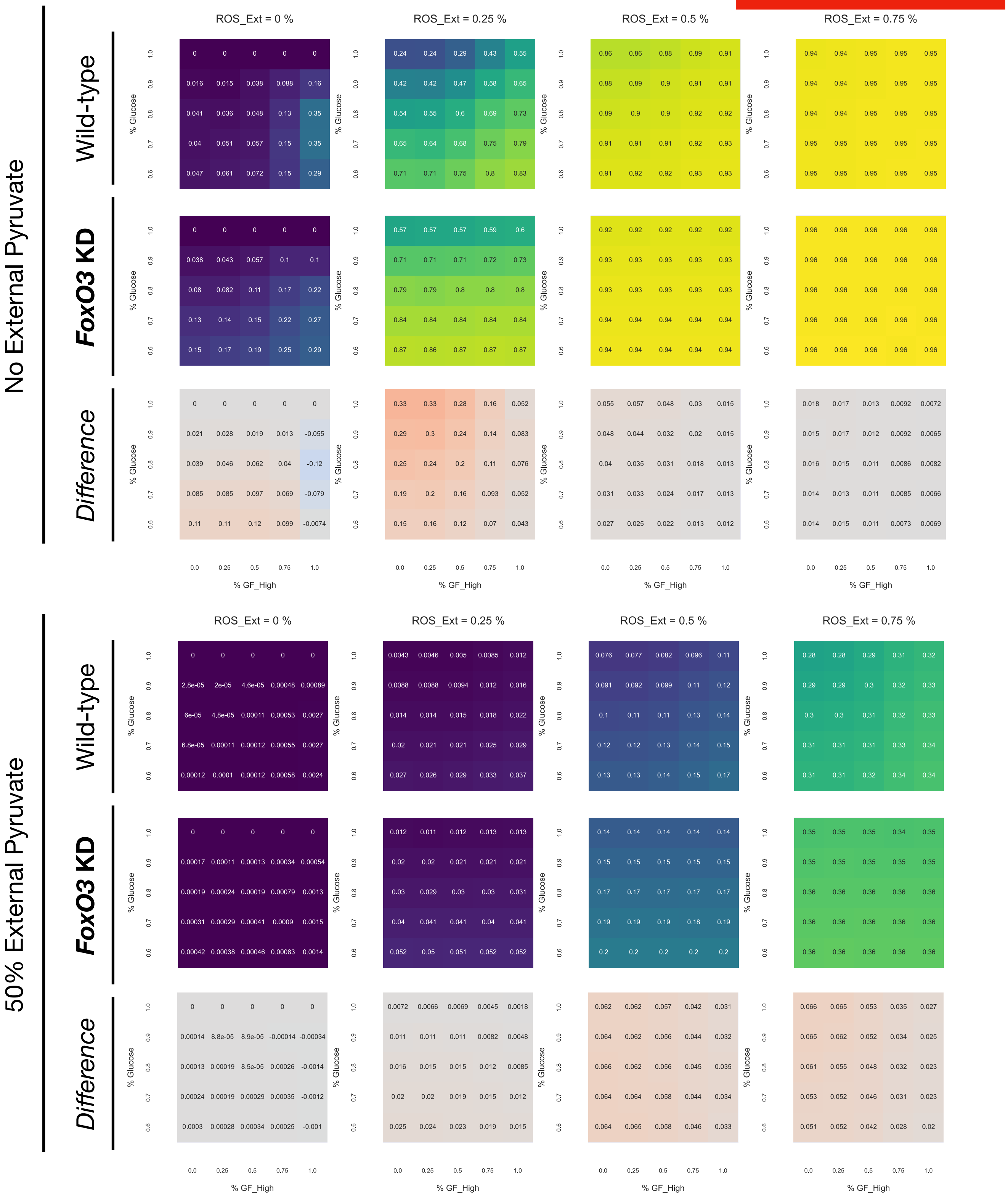

### MiDAS: WT (top) vs. p27Kip1 OE - WT (bottom)

**TS loss promotes MiDAS**

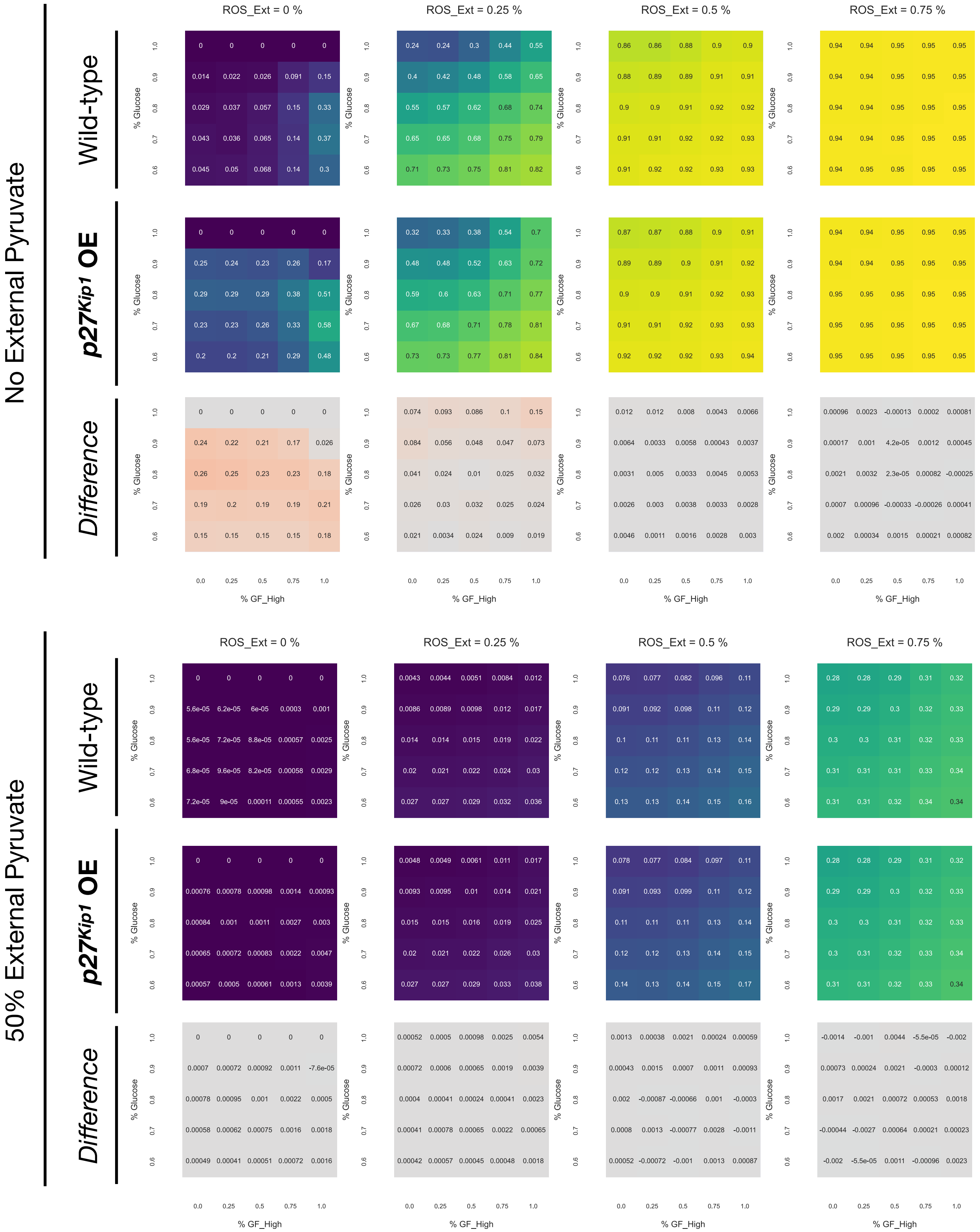

### MiDAS: WT (top) vs. Myc OE - WT (bottom)

**oncogene promotes MiDAS**

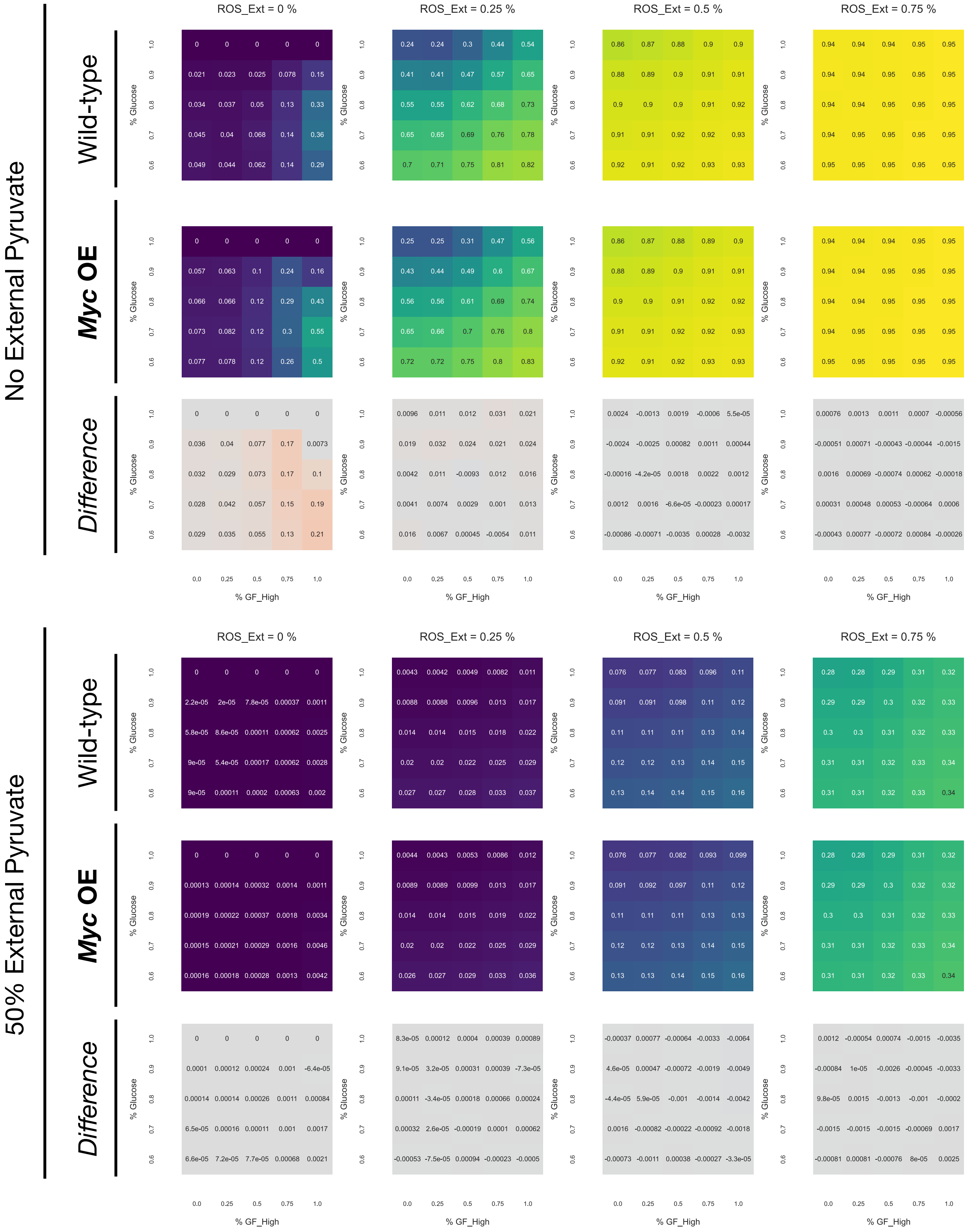

MiDAS: WT (top) vs. mTORC1 OE - WT (bottom)

oncogene promotes MiDAS

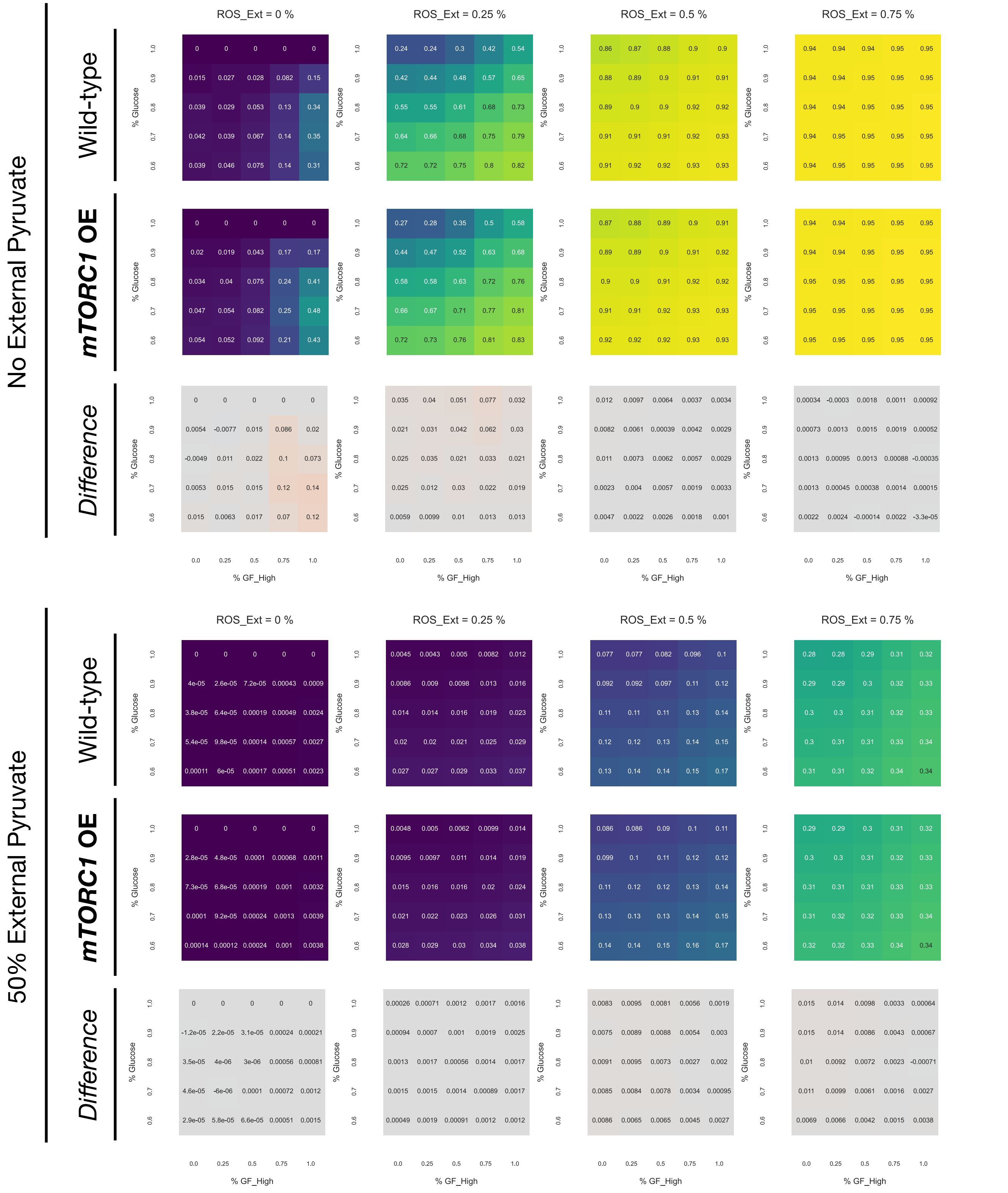

MiDAS: WT (top) vs. FoxO1 KD - WT (bottom)

**TS loss promotes MiDAS**

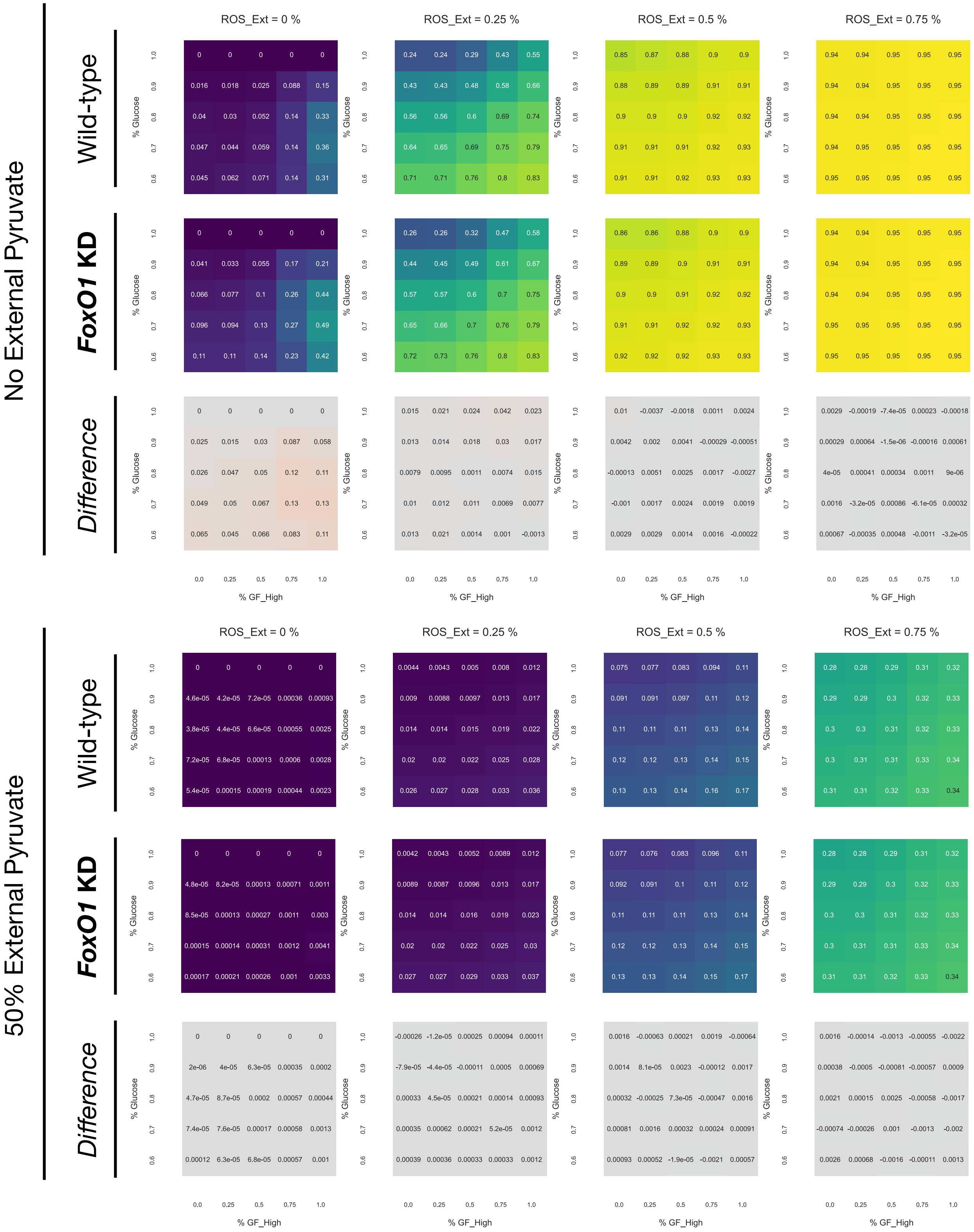

### MiDAS: WT (top) vs. p53 KD - WT (bottom)

**TS loss blocks MiDAS**

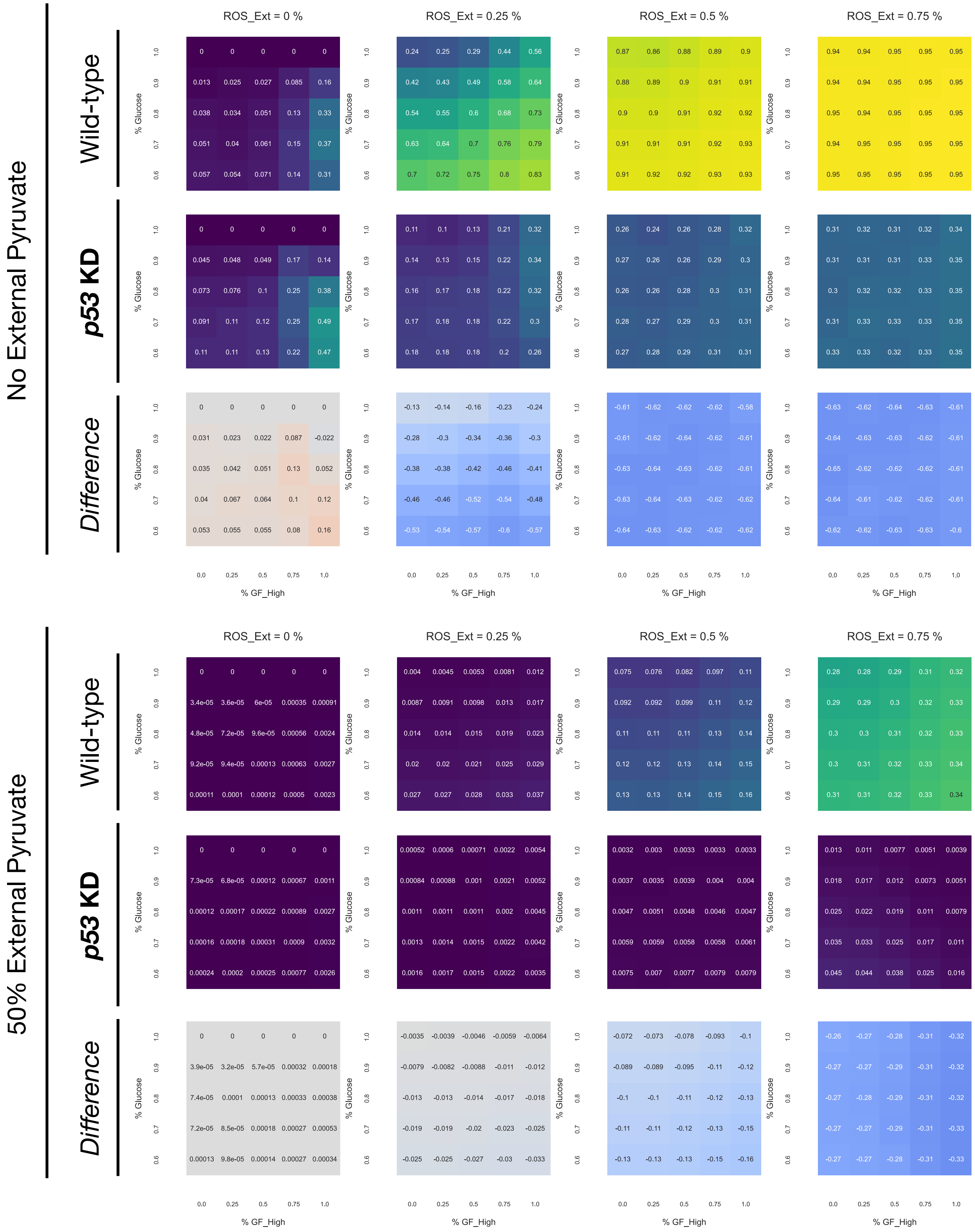

MiDAS: WT (top) vs. p21_H OE - WT (bottom)

**oncogene (?) blocks MiDAS**

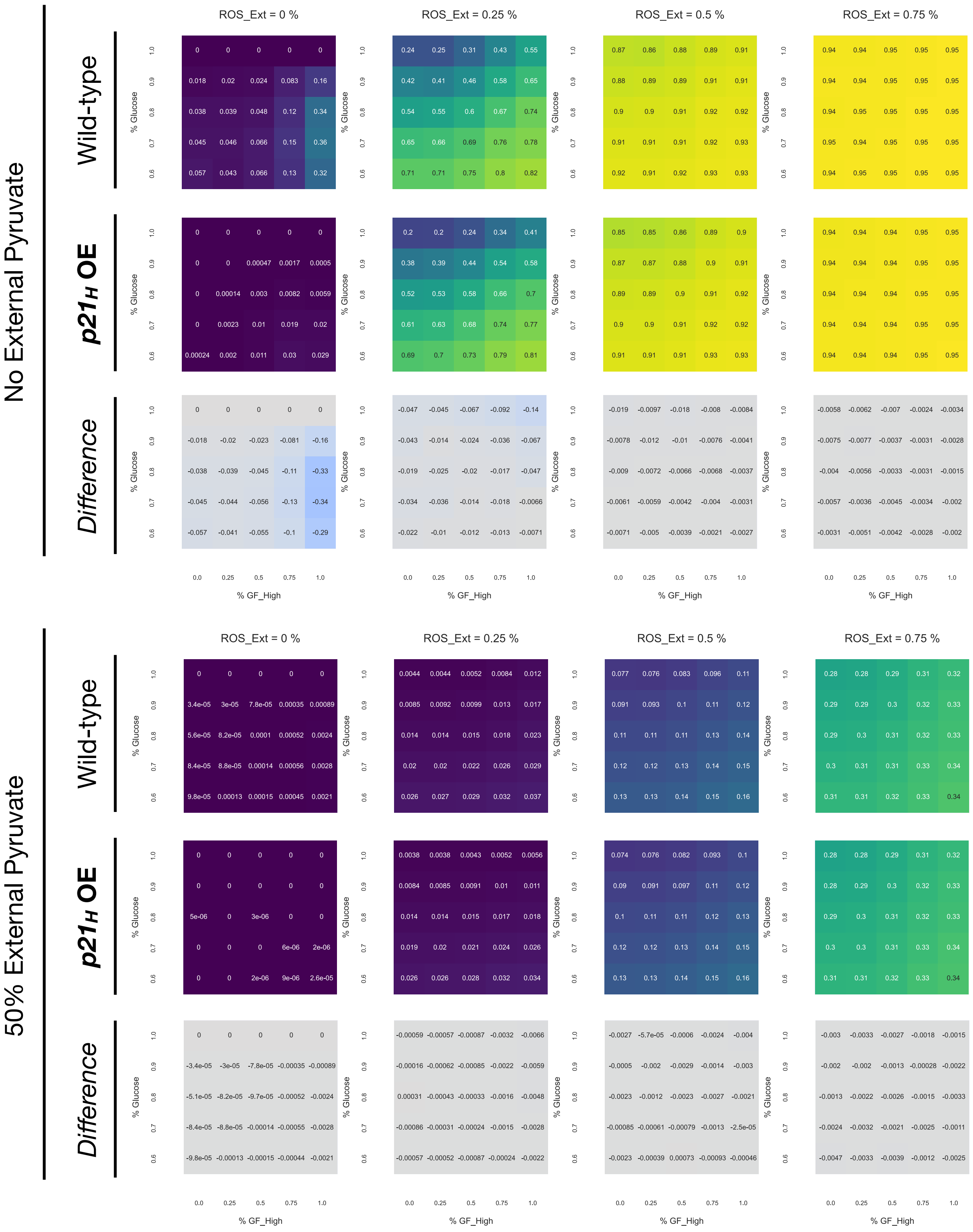

MiDAS: WT (top) vs. CyclinE OE - WT (bottom)

**oncogene blocks MiDAS**

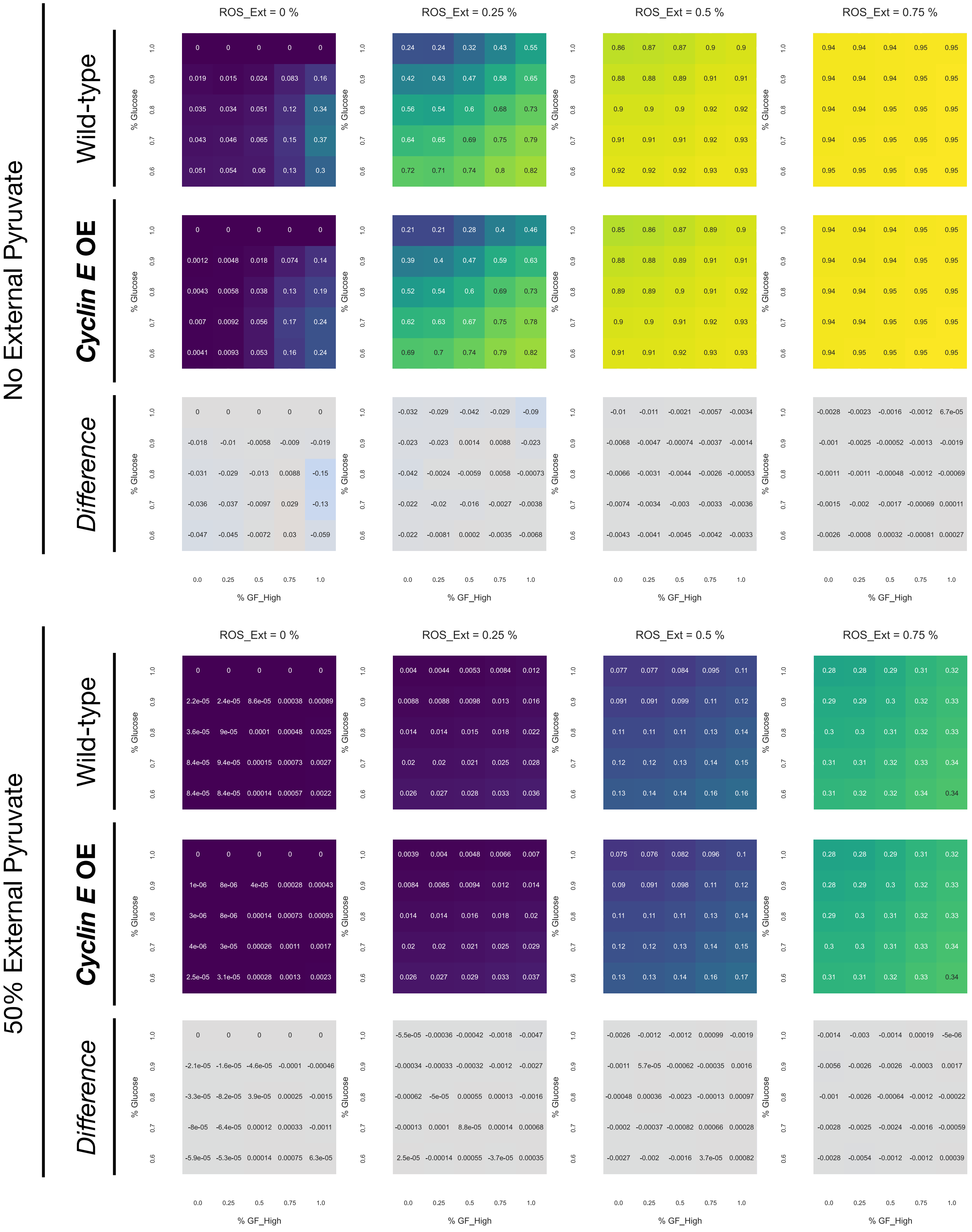

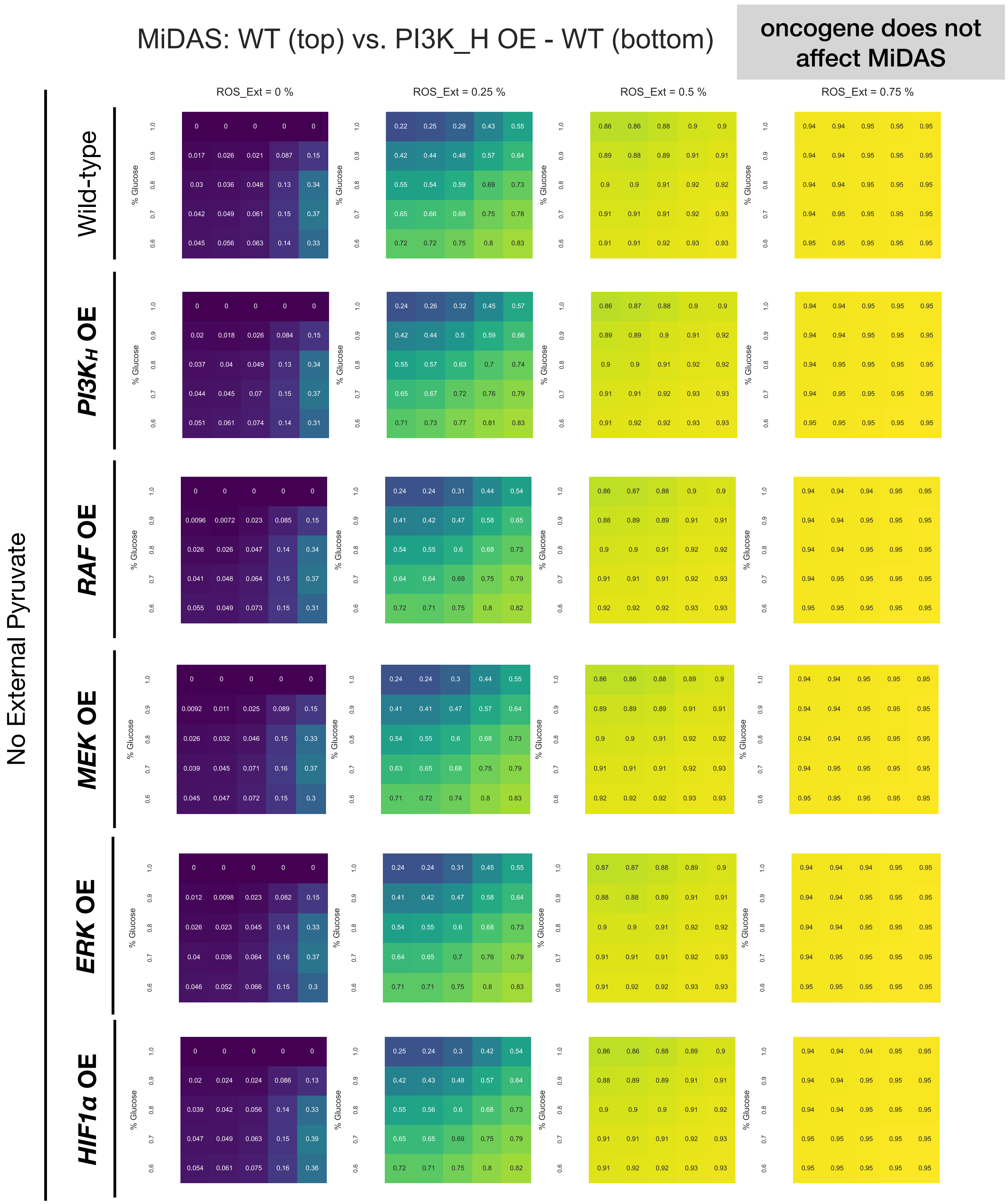
MiDAS: WT (top) vs. PI3K_H OE - WT (bottom)
oncogene does not affect MiDAS
ROS_Ext = 0 %
ROS_Ext = 0.25 %
ROS_Ext = 0.5 %
ROS_Ext = 0.75 %
No External Pyruvate
Wild-type
PI3KH OE
RAF OE
MEK OE
ERK OE
HIF1α OE
% Glucose
1.0
0.9
0.8
0.7
0.6

MiDAS: WT (top) vs. TSC2 KD - WT (bottom)

**TS loss does not affect MiDAS**

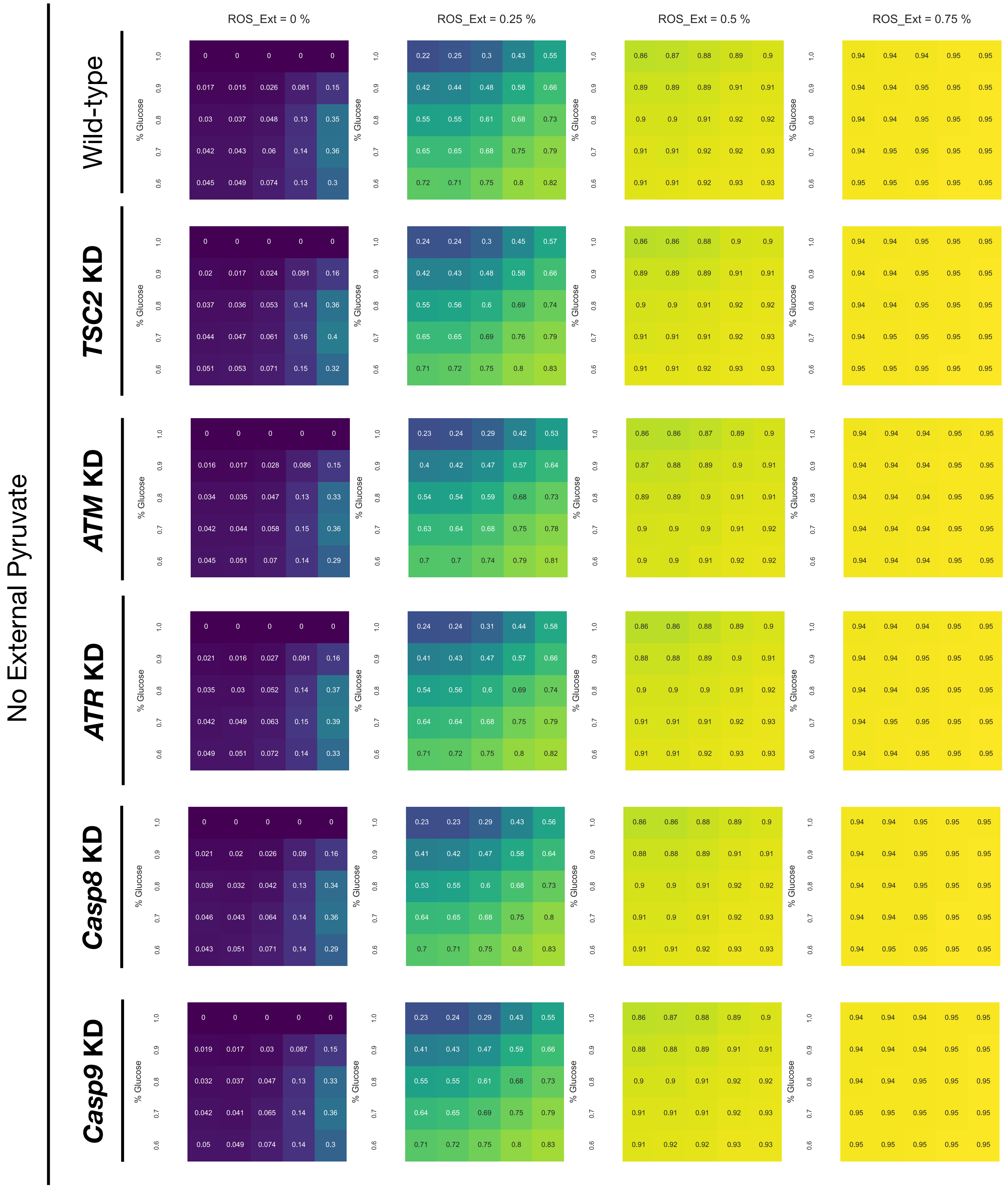
