## Supplementary material for "Unlocking Mitochondrial Dysfunction-Associated Senescence (MiDAS) with NAD^+^ – a Boolean Model of Mitochondrial Dynamics and Cell Cycle Control": SM Table 1

### S1 Description & experimental support for the modules of MiDAS_Cell_Cycle_Arrests_Apoptosis_Fine.

**Table S1a: Growth_Env module**

| Target Node | Node Gate<br>Node Type<br>Link Type | Node Description<br>Input Node | Link Description |
|---|---|---|---|
| GF | **GF** = **GF** or **GF_High** | | |
| | Env | The *GF* node represents an extracellular environment with low levels of growth factors capable of sustaining survival signaling. Thus, the *GF* input node is self-sustaining in the absence of in silico perturbation. | |
| | ← Env | GF | The *GF* input node is self-sustaining in the absence of in silico perturbation. |
| | ← Env | GF_High | The *GF* node represents an extracellular environment with low levels of growth factors capable of sustaining survival signaling. Thus, this node is ON in high growth factor as well. |
| GF_High | **GF_High** = **GF_High** | | |
| | Env | The $GF_{High}$ node in our model represents an extracellular environment with saturating levels of growth factors; this input node is self-sustaining in the absence of *in silico* perturbation. | |
| | ← Env | GF_High | The $GF_{High}$ input node is self-sustaining in the absence of *in silico* perturbation. |
| Glucose | **Glucose** = **Glucose** | | |
| | Env | The *Glucose* = OFF input node represents sufficent nutrient for quiescent cell survival aided by autophagy. Cell cycle entry, in contrast, requires *Glucose* = ON to reduce the AMP/ATP ratio and deactivate *AMPK* [1]. | |
| | ← Env | Glucose | The *Glucose* input node is self-sustaining in the absence of *in silico* perturbation. |
| Pyruvate _Ext | **Pyruvate_Ext** = **Pyruvate_Ext** | | |
| | Env | The *Pyruvate_Ext* node represents saturating access to extracellular pyruvate. | |
| | ← Env | Pyruvate _Ext | The *Pyruvate_Ext* input node is self-sustaining in the absence of *in silico* perturbation. |

**Table S1b: ROS module**

| Target Node | Node Gate<br>Node Type | Node Description |
|---|---|---|

**Table S1b: ROS module**

| | Link Type | Input Node | Link Description |
|---|---|---|---|
| ROS_Ext | **ROS_Ext = ROS_Ext** | | |
| | Env | The *ROS_Ext* node in our model represents an extracellular environment with high levels of ROS; this input node is self-sustaining in the absence of *in silico* perturbation. | |
| | ← Env | ROS_Ext | The *ROS_Ext* input node is self-sustaining in the absence of *in silico* perturbation. |

**Table S1c: GF_Basal_MAPK module**

| Target Node | Node Gate<br>Node Type<br>Link Type | Node Description<br>Input Node | Link Description |
|---|---|---|---|
| RTK | **RTK** = (**GF_High** or **GF**) and not **CAD** | | |
| | Rec | The ON state of the *RTK* node in our model represents basal growth receptor activation (required to keep a normal cell alive). Thus it requires the absence of *CAD* and at least low growth levels of growth factors in the extracellular environment [2]. | |
| | ← Ligand | GF | The ON state of the *RTK* node in our model represents basal growth receptor activation by low / medium growth factor availability, encoded by the *GF* node (required to keep a normal cell alive). |
| | ← Ligand | GF_High | Similarly, high growth factor availability also keeps *RTK* on. |
| | ⊣ Per | CAD | Caspase-activated DNase (*CAD*) inhibition of receptor tyrosine kinases ensures that apoptotic cells no longer maintain even basal levels of growth signaling. |
| Shc | **Shc** = **RTK** and **GF_High** | | |
| | Adap | *Shc* is ON when *RTK*s are activated by high levels of extracellular growth factors (capable of driving proliferation) [3]. | |
| | ← Compl | GF_High | The ON state of *Shc* in our model encode the change from basal *Shc* recruitment to weakly stimulated *RTKs* to the level of recruitment seen in high growth factor environments, capable of mediating *Ras* activation [3]. |
| | ← Compl | RTK | *Shc* proteins are adaptors that binds to phosphor-tyrosine motifs, facilitating their recruitment to activated receptors such as receptor tyrosine kinases *RTKs* [3]. |
| Grb2 | **Grb2** = **RTK** and **Shc** | | |
| | Adap | *Grb2* is recruited to *RTKs* upon ligand binding and subsequent recruitment of *Shc* adaptors [4]. | |

**Table S1c: GF_Basal_MAPK module**

| | | | |
|---|---|---|---|
| | $\underset{\text{Compl}}{\leftarrow}$ | RTK | The SH2 domain of *Grb2* binds to a phosphotyrosine residue in the activated *RTK*, where it functions as an adaptor protein [4]. |
| | $\underset{\text{Compl}}{\leftarrow}$ | Shc | *Shc* proteins phosphorylated by tyrosine kinases represent binding sites for *Grb2*, aiding its recruitment to active RTKs [3]. |
| SOS | **SOS** = **Grb2** | | |
| | GEF | *RTK*-bound *Grb2* recruits *SOS*, a guanine nucleotide–exchange protein (GEF) that converts inactive *Ras* to its active GTP-bound form [4]. | |
| | $\underset{\text{Compl}}{\leftarrow}$ | Grb2 | *Grb2* recruits *SOS* to activated *RTK*s [4]. |
| Ras | **Ras** = **Grb2** and **SOS** | | |
| | GTPa | *Ras* activation requires the GEF activity of *SOS* and the *RTK*-linked (active) adaptor protein *Grb2* [4]. | |
| | $\underset{\text{Compl}}{\leftarrow}$ | Grb2 | *RTK*-bound *Grb2* is required to recruits *SOS*, the GEF responsible for converting inactive *Ras* to its GTP-bound active form [4]. |
| | $\underset{\text{GEF}}{\leftarrow}$ | SOS | *SOS* is a GEF that is recruited to activate *Ras* near ligand-bound, active *RTK*s [4]. |
| RAF | **RAF** = **Ras** and not **Casp3** | | |
| | K | *Raf* is active in response to *Ras* activity in the absence of *Caspase 3*. As active *Raf-1* is continuously dephosphorylated and bound by *14-3-3*, which translocates it to the cytoplasm from the plasma membrane (not modeled explicitly), ongoing *Ras* activity is necessary to keep it ON [5]. | |
| | $\underset{\text{P}}{\leftarrow}$ | Ras | Active *Ras* phosphorylates *Raf*, enhancing its kinase activity [5]. |
| | $\underset{\text{Lysis}}{\vdash}$ | Casp3 | *Raf-1* is cleaved and inhibited by *Caspase 3* [6]. |
| MEK | **MEK** = **RAF** | | |
| | K | *Raf* phosphorylates and activates the *MEK* kinase [5]. | |
| | $\underset{\text{P}}{\leftarrow}$ | RAF | *Raf* phosphorylates and activates the *MEK* kinase [5]. |
| ERK | **ERK** = **MEK** and not **BIK** | | |
| | K | The *ERK* kinase is active when phoisphotylated by *MEK* [5] and allowed to translocate to the nucleus in the absence of *BIK* [7]. | |
| | $\underset{\text{P}}{\leftarrow}$ | MEK | *MEK* phosphorylates and activates the *ERK* kinase [5]. |
| | $\underset{\text{IBind}}{\vdash}$ | BIK | *BIK* binds to active, phosphorylated *ERK1/2* and suppresses its nuclear translocation [7]. |
| mTORC2 | **mTORC2** = (**PIP3** or not **S6K**) and not **Casp2** | | |

#### Table S1c: GF_Basal_MAPK module

| | | | |
|---|---|---|---|
| | PC | | Our model assumes that *mTORC2* is active in quiescent cells with basal levels of *PI3K* activity leading to basal *PIP3* generation [8]. Alternatively, the absence of high growth factor-stimulated *mTORC1* and *S6K1* can also increase *mTORC2* activity [9, 10]. In contrast, *Caspase-2* degrades a key *mTORC2* component and deactivates the complex [11]. |
| | ← PBind | PIP3 | PtdIns(3,4,5)P3 (*PIP3*), interacts with the *mTORC2* component *Sin1* to release its inhibition on the *mTOR* kinase domain. Thus, *PIP3* is necessary for *mTORC2* activation [8]. |
| | ⊢ P | S6K | *Rictor*, a component of the *mTORC2 complex*, undergoes *S6K1*-mediated phosphorylation at T1135, dampening *mTORC2*-dependent phosphorylation of *Akt* [9, 10]. |
| | ⊢ Lysis | Casp2 | *Caspase-2* degrades a key *mTORC2* component (*Rictor*) and inhibits downstream *Akt* activation [11]. |
| PI3K | **PI3K** = **RTK** or **Ras** | | |
| | K | | In our model, basal *PI3K* activity can be maintained by active *RTK*s or active *Ras*, while high *PI3K* activity requires both (see below). |
| | ← BLoc | RTK | Active *RTK*s recruit *PI3K* to the signaling complex they nucleate, where *PI3K* catalyzes the production of PtdIns(3,4,5)P3 (PIP3) [2]. |
| | ← Compl | Ras | *Ras* binds the catalytic subunit of *PI3K* and *Ras* knockdown / over expression decreases /increases the *PI3K*-dependent generation of PIP3 [12, 13]. |
| PIP3 | **PIP3** = **PI3K** or **PI3K_H** | | |
| | Met | | In our model, PIP3 is ON as a result of basal or high *PI3K* activity. |
| | ← Cat | PI3K | Active *PI3K* recruited to the membrane catalyzes the production of membrane-bound PtdIns(3,4,5)P3 (PIP3) from PtdIns(4,5)P2 (PIP2) [2]. |
| | ← Cat | PI3K_H | Active *PI3K* recruited to the membrane catalyzes the production of membrane-bound PtdIns(3,4,5)P3 (PIP3) from PtdIns(4,5)P2 (PIP2) [2]. |
| PDK1 | **PDK1** = **PI3K** and **PIP3** | | |
| | K | | *PDK1* enzyme activation requires active (at least basal) *PI3K* and *PIP3* [14]. |
| | ← BLoc | PI3K | The *PDK1* kinase is recruited to the plasma membrane by *PIP3* at the sites of active *PI3K* activity [14]. |
| | ← BLoc | PIP3 | The *PDK1* kinase is recruited to the plasma membrane by *PIP3* at the sites of active *PI3K* activity [14]. |
| AKT_B | **AKT_B** = (**PDK1** or **mTORC2**) and **PIP3** and not **Casp3** | | |
| | K | | Basal *AKT1* activity in our model requires the absence of *Caspase 3*, the availability of at least basal levels of *PIP3*, and phosphorylation by *PDK1* or *mTORC2*. In contrast, full mitogen-stimulated *AKT1* activation requires phosphorylation by both (see *AKT_H*) [14]. |

**Table S1c: GF_Basal_MAPK module**

| Target Node | Link Type | Input Node | Link Description |
|---|---|---|---|
| | ← P | PDK1 | Membrane-recruited *PDK1* phosphorylates *AKT1* at T308, a critical step in its activation [14]. |
| | ← P | mTORC2 | Maximal activation of *AKT1* requires phosphorylation of S473 by *mTORC2* [14]. |
| | ← BLoc | PIP3 | *PIP3* recruits *AKT1* to the plasma membrane and *PIP3* binding changes the conformation of *AKT1* such that it becomes accessible for T308 phosphorylation by *PDK1* [14]. |
| | ⊢ Lysis | Casp3 | *AKT1* is cleaved and inhibited by *Caspase 3* [6]. |

**Table S1d: GF_PI3K module**

| Target Node | Node Gate<br>Node Type<br>Link Type | Node Description<br>Input Node | Link Description |
|---|---|---|---|
| p110_H | **p110_H** = (**FoxO3** and not **Nedd4L**) or (**p110_H** and (**FoxO3** or not **Nedd4L**)) | | |
| | Prot | In order to capture the cyclic dynamics of *p110* protein expression, we make the assumption that high *p110* protein levels can be induced by *FoxO3* in the absence of the growth factor-activated *Nedd4L* ubiquitin ligase. Once present, high *p110* can be maintained by *FoxO3* transcription, or the absence of activated *Nedd4L*. | |
| | ← TR | FoxO3 | *FoxO3* is a direct inducer *p110*$\alpha$ (*PIK3CA*), the catalytic subunit of *PI3K* [15]. |
| | ⊢ Ubiq | Nedd4L | *p110*$\alpha$ (*PIK3CA*) is polyubiquitinated by the E3 ligase *Nedd4L*, leading to its proteasomal degradation. Both free *p110*$\alpha$ and the regulatory subunit-bound protein is subject to ubiquitination by *Nedd4L* [16]. |
| | ← Per | p110_H | Our model assumes that maintaining high *p110* levels is easier than driving the re-accumulation of the protein following its rapid destruction. |
| PI3K_H | **PI3K_H** = **p110_H** and **PI3K** and **RTK** and **Ras** | | |
| | K | Full, peak-level activation of *PI3K* requires high levels of *p110* protein, basal *PI3K* activation, active *RTK*s, and active *Ras*. As the ON-state of *Ras* in our model represents strong *Ras* activation in the presence of proliferation-inducing (high) growth factors, *PI3K_H* activation can only occur in these conditions. | |
| | ← Per | p110_H | High levels of *PI3K* activity in response to strong growth factor stimulation only occur in cells that express high levels of *p110* protein [17]. |
| | ← Per | PI3K | In our model, high *PI3K* activation is contingent on the ON-state of the basal *PI3K* node. |

**Table S1d: GF_PI3K module**

| | | | |
|---|---|---|---|
| | ← BLoc | RTK | High levels of *PI3K* activation only occur at growth factor-bound *RTK*s, which recruit and activate *PI3K* at the plasma membrane [14]. |
| | ← Compl | Ras | *Ras* binds the catalytic subunit of *PI3K* and *Ras* knockdown/over-expression decreases/increases the *PI3K*-dependent generation of PIP3 [12, 13]. |
| AKT_H | **AKT_H** = **AKT_B** and **p110_H** and **PI3K_H** and **PIP3** and **PDK1** and **mTORC2** and **Ras** | | |
| | K | | In contact to basal *AKT1*, high *AKT1* activity in our model requires basal *AKT1* (*AKT_B*), the ongoing presence of high *p110* protein levels along with active *PI3K_H* and *PIP3*. In addition this maximal *AKT1* activation requires phosphorylation by both *PDK1* and *mTORC2*, as well as active *Ras* [14]. |
| | ← Per | AKT_B | In our model, high *AKT1* activation is contingent on the ON-state of basal *AKT1* (*AKT_B*). |
| | ← P | p110_H | Ongoing high *p110* availability and *PI3K_H* activity are required to induce maximal activation of *AKT_H* [14]. |
| | ← P | PI3K_H | Ongoing high *p110* availability and *PI3K_H* activity are required to induce maximal activation of *AKT_H* [14]. |
| | ← BLoc | PIP3 | *PIP3* recruits *AKT1* to the plasma membrane and *PIP3* binding changes the conformation of *AKT1* such that it becomes accessible for T308 phosphorylation by *PDK1* [14]. |
| | ← P | PDK1 | Membrane-recruited *PDK1* phosphorylates *AKT1* at T308, a critical step in its activation [14]. |
| | ← P | mTORC2 | Maximal activation of *AKT1* requires phosphorylation of S473 by *mTORC2* [14]. |
| | ← Compl | Ras | *Ras* binding to the catalytic subunit of *PI3K* is required for its full potency in *PIP3* generation [12, 13]. Active *Ras* is thus required for inducing peak *AKT_H* activity. |
| FoxO3 | **FoxO3** = (not(**AKT_H** and (**ERK** or **AKT_B** or **Plk1** or **Plk1_H** or **p53**)) and not(**Plk1** and **Plk1_H** and **ERK**)) or ((**AMPK** or **SIRT3**) and not(**AKT_H** and (**Plk1** or **Plk1_H** or **p53**))) | | |
| | TF | | In order to account for all the influences on *FoxO3* activity, we used the following logic. In the absence of basal or high *AKT1* as well as *ERK*, *FoxO3* remains active. In addition, *FoxO3* can overcome peak (*AKT_H*) activation only if no other inhibitor is present and *AKT_B* is OFF (indicating that *AKT1* levels are falling)[14], or if *AMPK* [18, 19] or *SIRT3* [20] activate *FoxO3* in the absence of *Plk1* and *P53* [21, 22]. Finally, the joint activity of *ERK* and *Plk1* can also block *FoxO3* [21, 23]. |
| | ⊣ PLoc | AKT_H | *AKT1* mediates the translocation of the *FoxO3* our of the nucleus through direct phosphorylation of three conserved residues. These events create a recognition site for *14-3-3* family proteins, which export and sequester *FoxO3* in the cytosol [14]. |

**Table S1d: GF_PI3K module**

| | | | |
|---|---|---|---|
| | $\dashv$ PLoc | AKT_B | *AKT1* mediates the translocation of the *FoxO3* our of the nucleus through direct phosphorylation of three conserved residues. These events create a recognition site for *14-3-3* family proteins, which export and sequester *FoxO3* in the cytosol [14]. |
| | $\dashv$ P | ERK | ERK downregulates *FoxO3* transcriptional activity by phosphorylating it at three Serines, inducing its *MDM2*-mediated ubiquitination and degradation [23]. |
| | $\dashv$ PLoc | Plk1 | *Plk1* binds *FoxO3*, induces its translocation to the cytosol, phosphorylates it and suppresses its activity through most of the the cell cycle, but most significantly during G2 and M [21]. |
| | $\dashv$ PLoc | Plk1_H | *Plk1* binds *FoxO3*, induces its translocation to the cytosol, phosphorylates it and suppresses its activity through most of the the cell cycle, but most significantly during G2 and M [21]. |
| | $\dashv$ Ind | p53 | *p53*-induced increase in *MDM2*, an ubiquitin E3 ligase, promotes *FoxO1* and *FoxO3* ubiquitination and degradation in a manner dependent on *FoxO* phosphorylation at *AKT* sites [22, 24]. |
| | $\leftarrow$ P | AMPK | *AMPK* phosphorylation of *FoxO3* at its C-terminal portion (its the transactivation domain) increases its transcriptional activity without altering its cytoplasmic-nuclear shuttling [18, 19]. |
| | $\leftarrow$ Deacet | SIRT3 | *SIRT3* deacetylates *FoxO3* at K271 and K290, which promotes *FoxO3* nuclear localization and increases transcription [20]. |
| PLCgamma | **PLCgamma** = **RTK** and **Grb2** and **p110_H** and **PI3K_H** and **PIP3** | | |
| | Enz | | Peak activation of $PLC\gamma$ requires active an *RTK* receptor node bound by active *Grb2*, as well as high *PI3K* activity (including high *p110* availability and the presence of PIP3). |
| | $\leftarrow$ P | RTK | The SH2 domains of $PLC\gamma$ binds to active *RTK*s at tyrosine autophosphorylation sites, leading to tyrosine phosphorylation of $PLC\gamma$ and stimulation its enzymatic activity [25, 26]. |
| | $\leftarrow$ BLoc | Grb2 | *RTK* tyrosine autophosphorylation induces $PLC\gamma$ binding to the *Grb2* adaptor protein and likely aids the translocation of $PLC\gamma$ to the plasma membrane [27]. |
| | $\leftarrow$ BLoc | p110_H | Membrane targeting of $PLC\gamma$ to growth receptor stimulation requires *PI3K* activity and *PIP3* generation near growth receptors [28]. Thus, peak $PLC\gamma$ activity in our model requires high *p110* protein expression [29]. |
| | $\leftarrow$ BLoc | PI3K_H | In addition to high *p110* protein levels, high *PI3K* activation is also required to fully activate $PLC\gamma$ [28, 29]. |
| | $\leftarrow$ BLoc | PIP3 | Membrane targeting of $PLC\gamma$ to growth receptor stimulation is mediated by *PIP3* binding of $PLC\gamma$ [28, 29]. |

**Table S1d: GF_PI3K module**

| | | | |
|---|---|---|---|
| IP3 | **IP3** = **PLCgamma** | | |
| | Met | Membrane-bound, active $PLC\gamma$ is responsible for converting phosphatidylinositol(4,5)P2 (*PIP2*) to the second messenger inositol(1,4,5)P3 (*IP3*) responsible for triggering a sudden $Ca^{2+}$ influx from the endoplasmic reticulum, along with *DAG* (diacylglycerol, another second messenger) [30]. | |
| | ← Cat | PLCgamma | Membrane-bound, active $PLC\gamma$ is responsible for converting phosphatidylinositol(4,5)P2 (*PIP2*) to the second messenger inositol(1,4,5)P3 (*IP3*) responsible for triggering a sudden $Ca^{2+}$ influx from the endoplasmic reticulum, along with *DAG* (diacylglycerol, another second messenger) [30]. |
| Ca2p | **Ca2p** = **IP3** | | |
| | Met | *IP3* travels from the cell membrane to the endoplasmic reticulum where it opens *IP3*-sensitive $Ca^{2+}$ channels, releasing a sudden $Ca^{2+}$ efflux from the ER into the cytosol [31]. | |
| | ← Loc | IP3 | *IP3* travels from the cell membrane to the endoplasmic reticulum where it opens *IP3*-sensitive $Ca^{2+}$ channels, releasing a sudden $Ca^{2+}$ efflux from the ER into the cytosol [31]. |
| Nedd4L | **Nedd4L** = **Ca2p** and **IP3** | | |
| | UbL | Activation of *Nedd4L* requires both $Ca^{2+}$ and *IP3* binding [32]. | |
| | ← Compl | IP3 | In order to transition to its active form, the E3 ubiquitin ligase *Nedd4L* binds $Ca^{2+}$ and inositol 1,4,5-trisphosphate (*IP3*) [32]. |
| | ← Compl | Ca2p | In order to transition to its active form, the E3 ubiquitin ligase *Nedd4L* binds $Ca^{2+}$ and inositol 1,4,5-trisphosphate (*IP3*) [32]. |

**Table S1e: GF_mTOR module**

| Target Node | Node Gate<br>Node Type<br>Link Type | Node Description<br>Input Node | Link Description |
|---|---|---|---|
| FoxO1 | **FoxO1** = not(**AKT_H** or **Plk1** or **p53**) or (**AMPK** and **SIRT3** and not(**AKT_B** and **p53**)) | | |
| | TF | *FoxO1* is transcriptionally active in the absence of peak *AKT1* activation [14], *Plk1* activity [33], or *p53* activation [22]. Alternativelty, *AMPK* [34] and *SIRT3* [35] can activate *FoxO1* if there is no active *p53* (aided by at least basal *AKT*). | |
| | ⊢ PLoc | AKT_H | *AKT1* mediates the translocation of *FoxO1* our of the nucleus through direct phosphorylation of three conserved residues. These events create a recognition site for *14-3-3* family proteins, which export and sequester *FoxO1* in the cytosol [14]. |

**Table S1e: GF_mTOR module**

| | | | |
|---|---|---|---|
| | ⊣ PLoc | AKT_B | *AKT1* mediates the translocation of *FoxO1* our of the nucleus through direct phosphorylation of three conserved residues. These events create a recognition site for *14-3-3* family proteins, which export and sequester *FoxO1* in the cytosol [14]. |
| | ⊣ PLoc | Plk1 | *Plk1* interacts with and phosphorylates *FoxO1*, mainly at the G2/M phase of the cell cycle. *Plk1*-mediated phosphorylation leads to the impairment of *FoxO1*'s transcriptional activity in an *Akt*-independent manner. *Plk1*-induced *FoxO1* phosphorylation causes its nuclear exclusion [33]. |
| | ⊣ Ind | p53 | *p53*-induced *MDM2*, an ubiquitin E3 ligase, promotes *FoxO1* and *FoxO3* ubiquitination and degradation in a manner dependent on *FoxO* phosphorylation at *AKT* sites [22]. |
| | ← PLoc | AMPK | *AMPK* induces *FoxO1* activity by phosphorylation at Ser383, Thr649, and Ser22; where Ser22 phosphorylation prevents binding to the *14-3-3* proteins (these inhibit DNA binding and mask *FoxO1*'s nuclear localization signal) [34]. |
| | ← Deacet | SIRT3 | *SIRT3* binds to and deacetylates *FoxO1* and increases the transtiption of its antioxidant targets (catalase, MnSOD) [35]. |
| MnSOD | **MnSOD** = **FoxO1** or **FoxO3** or **FoxM1** or (**ERK** and not **p27Kip1**) or **SIRT3** | | |
| | Enz | | Mitochondrial superoxide dismutase 2 (*SOD2*), also known as manganese-dependent superoxide dismutase or *MnSOD*, is induced by *FoxO* transcription factors, as well as *FoxM1* during cell cycle progression [36, 37, 38]. In addition, increased *ERK* in the absence of $p27^{Kip1}$ can also induce textitMnSOD [39]. |
| | ← TR | FoxO1 | *FoxO1* is a direct transcriptional inducer of *MnSOD*, potentially dependent on the presence of the mechanosensitive *YAP* transcription factor (not explicitiy modeled, but assumed to be active in non-contact-inhibited cells we model) [37]. |
| | ← TR | FoxO3 | *FoxO3* is a direct transcriptional inducer of *MnSOD* [36]. Note that *FoxO3* does not appear to interact with, or require *YAP* to induce *MnSOD* [37]. |
| | ← TR | FoxM1 | *FoxM1* can substitute for *FoxOs* to keep *MnSOD* transctiption going during the cell cycle, when *FoxO* activity is blocked by *AKT* [38]. |
| | ← Ind | ERK | *ERK*-mediated induciton of *ATF1* upregulates *STAT3*, which in turn induces *MnSOD* expression; an effect observed in textitp27$^{Kip1}$-/- cells [39]. |
| | ⊣ TR | p27Kip1 | Loss of $p27^{Kip1}$ upregulates the expression of *STAT3* in an *ERK*-dependent manner (via *ATF1*). *STAT3*, in turn, is a direct transcriptional inducer of *MnSOD* [39]. As a result, textitp27$^{Kip1}$-/- cells have elevated *MnSOD* levels [39]. |
| | ← Deacet | SIRT3 | *SIRT3* deacetylates and activates *MnSOD* [40]. |
| p21 _mRNA | **p21_mRNA** = (**FoxO1** and **FoxO3**) or ((**FoxO1** or **FoxO3**) and not **Myc**) or **p53** | | |

**Table S1e: GF_mTOR module**

| | | | |
|---|---|---|---|
| | mRNA | Our model requires both *FoxOs* or *p53* to induce *p21*$^{Cip1}$ if *Myc* is active, but only one of the two *FoxOs* or *p53* if *Myc* is OFF. This is based on data showing that both *FoxO3* and *FoxO1* bind and induce the *p21*$^{Cip1}$ promoter, and that loss of *Myc* repression alone is not sufficient to induce *p21*$^{Cip1}$ [41]. | |
| | $\leftarrow$ TR | FoxO1 | *p21*$^{Cip1}$ is a direct transcriptional target of *FoxO1* [41]. |
| | $\leftarrow$ TR | FoxO3 | *p21*$^{Cip1}$ is a direct transcriptional target of *FoxO3* [41]. |
| | $\vdash$ TR | Myc | *Myc* is a direct transcriptional repressor of the *p21*$^{Cip1}$ promoter (it is recruited by the DNA-binding *Miz-1*) [42, 43]. |
| | $\leftarrow$ TR | p53 | *p21*$^{Cip1}$ is a direct transcriptional target of *p53* [44]. |
| TSC2 | **TSC2** = **AMPK** or **MP_Low** or not **AKT_H** or not(**AKT_B** or **ERK**) | | |
| | Prot | Blocking *TSC2* requires ongoing mitogen stimulation through *AKT* and/or *ERK*, as well as the absence of active *AMPK* [45] and low $\Delta\Psi_M$ [46]. In our model, *TSC2* inhibition in the absence of an energy deficit requires high (peak) *AKT* activity, supported by either *ERK* or basal *AKT* (assuring that complete loss of *AKT* activity is not impending) [47]. | |
| | $\leftarrow$ P | AMPK | *AMPK* phosphorylates *TSC2* to enhance its activity to block *mTORC1* in response to energy deprivation [45]. |
| | $\leftarrow$ ComplProc | MP_Low | $\Delta\Psi_M$ lowered with OXPHOS uncouplers results in *TSC2* activation and *mTORC1* inhibition even in the absence of *AMPK* [46]. |
| | $\vdash$ P | AKT_H | *TSC2* is phosphorylated by *AKT1*, inhibiting it by dissociating *TSC2* from lysosomal membranes [48], where it stimulates GTP hydrolysis of the small GTPase *Rheb*, this inactivating it [47]. |
| | $\vdash$ P | AKT_B | *TSC2* is phosphorylated by *AKT1*, inhibiting it by dissociating *TSC2* from lysosomal membranes [48], where it stimulates GTP hydrolysis of the small GTPase *Rheb*, this inactivating it [47]. |
| | $\vdash$ P | ERK | *ERK* phosphorylates *TSC2* directly, causing dissociation of the complex and inhibition of its activity [49]. In addition, the *ERK* target *p90RSK* can also inactivate *TSC2* [50]. |
| PRAS40 | **PRAS40** = not **AKT_H** and not(**AKT_B** and **mTORC1**) | | |
| | Prot | *PRAS40* is inhibited by peak *AKT1* activity aided by either basal *AKT1* (meaning *AKT_H* is on its way down), or ongoing *mTORC1* activation. Both *AKT1* and *mTORC1* phosphorylate *PRAS40*, leading to its dissociation from *mTORC1* [51]. | |
| | $\vdash$ P | AKT_H | *PRAS40* is an inhibitory component of the *mTORC1* complex. It is phosphorylated by *AKT*, triggering its dissociation from *mTORC1* and loss of *mTORC1* inhibition [52]. |
| | $\vdash$ P | AKT_B | *PRAS40* is phosphorylated by *AKT*, triggering its dissociation from *mTORC1* [52]. |

**Table S1e: GF_mTOR module**

| | | | |
|---|---|---|---|
| | ⊢ P | mTORC1 | *PRAS40* is a substrate of the *mTORC1* kinase; its phosphorylation aids its dissociation from *mTORC1* and its sequestration by 14-3-3 proteins [53]. |
| DAG | **DAG** = **PLCgamma** | | |
| | Met | Membrane-bound, active *PLC*$\gamma$ is responsible for converting phosphatidylinositol(4,5)P2 (*PIP2*) to the second messenger diacylglycerol (*DAG*), along with *IP3* [30]. | |
| | ← Cat | PLCgamma | Membrane-bound, active *PLC*$\gamma$ is responsible for converting phosphatidylinositol(4,5)P2 (*PIP2*) to the second messenger diacylglycerol (*DAG*), along with *IP3* [30]. |
| Rheb | **Rheb** = **DAG** and not **TSC2** | | |
| | GTPa | *PKC* (and *DAG*)-dependent activation of *mTORC1* recruits *mTORC1* to the site of *Rheb* activity (to prenuclear lysosomes), while *AKT* and *ERK*-mediated *TSC2* inhibition guarantees that *Rheb* remains potent [54, 55, 56]. | |
| | ← Loc | DAG | The second messenger *DAG* activates both classical and novel *PKC*s. One of its targets, *PKC*$\eta$, is responsible for the translocation and accumulation of *mTORC1* to perinuclear lysosomes, where the majority of *Rheb* is anchored. Thus, *DAG* brings *Rheb* in proximity with its target, *mTORC1* [55]. |
| | ⊢ GAP | TSC2 | *TSC2*, a key component of the heterotrimeric *TSC* complex, is a GTPase activating protein (GAP) that induces ATP hydrolysis and deactivation of the small GTPase *Rheb* [54]. |
| mTORC1 | **mTORC1** = ((not **AMPK** and not **PRAS40** and **Rheb**) or (**E2F1** and (not **AMPK** or **ERK**)) or (**CyclinB** and **Cdk1** and **GSK3**)) and not **Casp3** | | |
| | PC | *mTORC1* is activated by mitogenic signals via *Rheb* in the absence of both active *PRAS40* and *AMPK*. Independently, *E2F1* can promote *mTORC1* activity as long as *AMPK* is absent, or *ERK* is active (*E2F1* can override *mTORC1* inhibition by *TSC2* [57]; we assume it requires growth signaling to do so.). Finally, the mitotic *Cyclin B* and its *Cdk1* kinase can also activate *mTORC1*, aided by *GSK3*. | |
| | ⊢ P | AMPK | *AMPK* inhibits *mTORC1* by direct phosphorylation of the *mTORC1* member *Raptor*, as well as by activation of the *TSC1/2* pathway (also included in the model via its effects on *Rheb*) [58]. |
| | ⊢ IBind | PRAS40 | *PRAS40* is an inhibitory component of the *mTORC1* complex, removed by phosphorylation by *AKT* or *mTORC1* itself [47]. |
| | ← Compl | Rheb | The *Rheb* small GTPase binds *mTORC1* directly and activates the complex [59]. |
| | ← Loc | E2F1 | *E2F1* induces *mTORC1* activity by inducing *mTORC1* translocation to late endosomes. This effect does not require *AKT* and is not blocked by high levels of *TSC2* [57]. |

**Table S1e: GF_mTOR module**

| | | | |
|---|---|---|---|
| | ← P | ERK | *ERK1* and *ERK1* bind to the *mTORC1* component *Raptor* and phosphorylate it at Ser8, Ser696, and Ser863, modifications that promote *mTORC1* activity [60]. Here we assume that *ERK* can help *E2F1* maintain active *mTORC1* in the presence of *AMPK* (*E2F1* was shown to override *mTORC1* inhibition by *TSC2* [57]). |
| | ← Ind | CyclinB | During mitosis, *mTORC1* is activated by the G2/M-specific phosphorylation of *Raptor*, a component of *mTORC1*, by *CyclinB* / *Cdk1* complexes, aided by *GSK3* [61]. |
| | ← Ind | Cdk1 | During mitosis, *mTORC1* is activated by the G2/M-specific phosphorylation of *Raptor*, a component of *mTORC1*, by *CyclinB* / *Cdk1* complexes, aided by *GSK3* [61]. Here we assume that G2/M-specific phosphorylation of *Raptor* by *Cdk1* overrides direct inhibition by *AMPK*, keeping *mTORC1* active during mitosis. |
| | ← Ind | GSK3 | During mitosis, *mTORC1* is activated by the G2/M-specific phosphorylation of *Raptor*, a component of *mTORC1*, by *CyclinB* / *Cdk1* complexes, aided by *GSK3* [61]. |
| | ⊢ Lysis | Casp3 | *Raptor*, a key component of the *mTORC1* complex, is cleaved and inhibited by *Caspase 3* [62]. |
| S6K | **S6K** = **mTORC1** and not(**CyclinB** and **Cdk1**) and not **Casp3** | | |
| | K | *S6K* is activated by *mTORC1* in the absence of *Caspase 3*. | |
| | ← P | mTORC1 | *mTORC1* phosphorylates and activates 40S ribosomal S6 kinases (*S6K*s) [63]. |
| | ← Ind | CyclinB | During mitosis, p70 *S6K* is a physiological substrate of Cdk1/Cyclin, keeping its mitotic activity low. This interaction may be responsible for supressed protein synthesis during mitosis [64]. |
| | ← Ind | Cdk1 | During mitosis, p70 *S6K* is a physiological substrate of Cdk1/Cyclin, keeping its mitotic activity low. This interaction may be responsible for supressed protein synthesis during mitosis [64]. |
| | ⊢ Lysis | Casp3 | *S6K* is cleaved and inhibited by *Caspase 3* [65]. |
| eIF4E | **eIF4E** = **mTORC1** and not **Casp3** | | |
| | Prot | *eIF4E* is activated by *mTORC1*-mediated *4EBP* repression in the absence of *Caspase 3*. | |
| | ← Ind | mTORC1 | *mTORC1* phosphorylates *4EBP*, triggering its dissociation from *eIF4F*, and thus promoting translation initiation [47]. |
| | ⊢ Lysis | Casp3 | *eIF4E* is cleaved and inhibited by *Caspase 3* [66]. |
| GSK3 | **GSK3** = (not **AKT_H** or **U_Kinetochores**) and (**SIRT3** or not **AKT_B** or not(**S6K** and **ERK**)) | | |

**Table S1e: GF_mTOR module**

| | | | |
|---|---|---|---|
| | K | | *GSK3β* activity can be completely blocked by peak *AKT* activity (*AKT_H*), or by the joint action of *S6K* and *ERK*. Peak *AKT*-mediated *GSK3β* inhibition is overriden during metaphase (marked by the presence of unattached kinetochores, or *U_Kinetochores* = ON), when active *AKT* localizes primarily to the centrosomes and spindle poles, leaving cytosolic *GSK3β* active. In addition, *SIRT3* can deacetylate and activate *GSK3β*, overriding the joint inhibitory effects of *S6K* and *ERK*. |
| | ⊢ P | AKT_H | *AKT* blocks *GSK3β* kinase activity via an inhibitory phosphorylation on the amino terminus, which blocks the substrate accessibility of *GSK3β* [14]. |
| | ⊢ P | AKT_B | *AKT* blocks *GSK3β* kinase activity via an inhibitory phosphorylation on the amino terminus, which blocks the substrate accessibility of *GSK3β* [14]. We assume that loss of basal *AKT* activity overrides the joint effects of *S6K* and *ERK* on *GSK3* inhibition. |
| | ← Loc | U _Kinetochores | During metaphase, active *AKT* and phosphorylated (inactive) *GSK3β* localize primarily to the centrosomes and spindle poles, leaving cytosoling *GSK3β* active [67]. |
| | ← Deacet | SIRT3 | *SIRT3* activates *GSK3β* by deacetylating it at K15 [68]. |
| | ⊢ P | S6K | *GSK3β* is a direct phosphorylation target of *S6K1*, resulting in its inhibition [9]. |
| | ⊢ P | ERK | *ERK* binds and phosphorylates *GSK3β* at Thr-43, which primes it for subsequent phosphorylation by the *ERK* target *p90RSK* at Ser-9, which inactivates *GSK3β* [69]. |
| Autophagy | **Autophagy** = (**FoxO1** or **FoxO3**) and **AMPK** and not **mTORC1** and not **BIM** and (**mROS** or **cROS** or **cROS_H**) | | |
| | Proc | | To account for increased autophagy triggered by low ATP/AMP, we assume that *AMPK* activation aided by *FoxO1/3* in the absence of *mTORC1* or *BIK*, can increase autophagy from its basal levels to a state that can regenerate sufficient ATP for survival. In addition to energy stress, this increase also requires an increase in internal ROS. Note that the current model does not include the processes that eventualy kills the cell in response to prolonged glucose withdrawal, only the metastable homeostasis before the cell permanenly runs out of energy. |
| | ← ComplProc | FoxO3 | *FoxO3* stimulates lysosomal proteolysis by inducing the expression of several autophagy-related genes, such as *LC3b*, *Gabarapl1*, and *PI3KIII* (required for autophagy), *Ulk2*, *Atg12l*, and *Beclin1*. |
| | ← ComplProc | FoxO1 | *FoxO1* increased *Rab7* expression, which mediates late autophagosome-lysosome fusion [70]. In addition to its transcriptional role, *FoxO1* is also required for autophagy in human cancer cell lines in response to oxidative stress or serum starvation, where it is acetylated and bound to *Atg7* (essential autophagy effector enzyme) [71]. |

**Table S1e: GF_mTOR module**

| | | | |
|---|---|---|---|
| | ← ComplProc | AMPK | During glucose starvation, *AMPK* directly activates *Ulk1* through phosphorylation of Ser-317 and Ser-777, which in turn promotes autophagy [72]. |
| | ⊣ ComplProc | mTORC1 | High *mTORC1* activity prevents *Ulk1* activation by phosphorylating *Ulk1* Ser-757, which disrupts its interaction with *AMPK* [72]. |
| | ⊣ IBind | BIM | *BIM* inhibits autophagy by interacting with and changing the localization of *Beclin 1* [73]. |
| | ← ComplProc | mROS | ROS production, the main intrinsic source of which are mitochidria, aids autophagy (the exact mechanism of this is still debated) [74]. |
| | ← ComplProc | cROS | ROS production is induced immediately upon nutrient deprivation, where it aids autophagy (the exact mechanism of this is still debated) [74]. |
| | ← ComplProc | cROS_H | ROS production is induced immediately upon nutrient deprivation, where it aids autophagy (the exact mechanism of this is still debated) [74]. |
| IKKa | **IKKa = AKT_H** | | |
| | K | *IKKα* is a subunit of the *IKK* protein complex composed of two catalytic subunits, *IKKα* and *IKKβ*, and the regulatory protein *NEMO*. Activation of the transcription factor *NF-κB* is mediated by the IKK complex, which phosphorylates and degrades the inhibitory *IκB* proteins [75]. | |
| | ← P | AKT_H | *IKKα* is phosphorylated by *AKT* at T23, and as subsequent *NF-κB* activation is induced when high *AKT* activity is observed, we model it via the *AKT_H* input [76]. |
| NF_kB | **NF_kB = IKKa** and not **FoxO3** and not **p53** | | |
| | TF | *NF-κB* is a transcription factor primarily known as a master regulator of inflammatory signaling and the immune system. It is activated via the destruction of its inhibitory binding partner *IκB*, which is phosphorylated by the *IKK* complex subsequently destroted [75]. *FoxO3* and *p53* both limit *NF-κB*-mediated transcription by trapping one of its subunits in the cytosol and competing for a limited pool of cofactors, respectively [77, 78]. | |
| | ← Ind | IKKa | *IKKα*, part of the *IKK* complex, phosphorylates and degrades the inhibitory *IκB* proteins [75]; an action that can be independent of the *IKKβ* subunit [79]. |
| | ⊣ IBind | FoxO3 | *FoxO3* binds and traps *NF-κB*' *RelA* subunit in the cytosol [77]. |
| | ⊣ Ind | p53 | *p53*-mediated inhibition of *NF-κB* is likely the result of their competition for a limited pool of *p300/CBP complexes*, as they both require these transcriptional coactivators [78]. |

**Table S1f: Warburg module**

| Target Node | Node Gate<br>Node Type<br>Link Type | Node Description<br>Input Node | Link Description |
|---|---|---|---|
| Glycolysis | **Glycolysis** = (**Glucose** and **AKT_B** and **NADp_c**) or **Glycolysis_H** | | |
| | Proc | The *Glycolysis* node represents the metabolic process by which cells bring in glucose from the extracellular environment and convert it into pyruvate [80]. As a step of this pathway reduces NAD$^+$ [80], our model assumes that the *Glycolysis* node turns on when there is extracelluar glucose and the *NADp_c* node representing cytosolic NAD$^+$ is ON. | |
| | ← ComplProc | Glucose | *Glycolysis* requires glucose uptake from the envrironment, which is converted to pyruvate in the cytosol [80]. |
| | ← Ind | AKT_B | *Akt* increases glucose uptake by promoting *Glut1* localization to the cell surface [81], as well as glucose phosphorylation by increasing cellular hexokinase (*HK2*) levels (this effect requires mTORC1 activity) [82]. |
| | ← Cat | NADp_c | NAD$^+$ is converted to NADH at the glyceraldehyde 3-phosphate dehydrogenase (GAPDH) step of glycolysis [80]. Inhibition of *NAMPT*, an enzyme essential for NAD$^+$ biosynthesis leads to attenuation of glycolysis [83]. |
| | ← Per | Glycolysis_H | To keep our Boolean model conistent with the use of two nodes for three qualitatively distinct levels of glycolysis, *Glycolysis_H* = ON keeps this basal glycolisis node ON as well. |
| Glycolysis_H | **Glycolysis_H** = (**HIF1** or **Myc**) and **Glycolysis** and **Glucose** and **NADp_c** and **ERK** and not **p53** | | |
| | Proc | The *Glycolysis_H* node represents incrased glycolysis characteristic of proliferative metabolism, driven by increased glucose uptake and enhanced glycolytic enzyme expression downstream of *HIF1* (primarily under anaerobic conditions) or *Myc* (in proliferating cells under aerobic conditions); further aider by *ERK1/2*. In contrast, *p53* activation limits glycolysis. | |
| | ← TR | HIF1 | Hypoxia-induced factor-1 or *HIF-1*$\alpha$ is a transcriptional inducer of nealry every step of glycolysis, including *GLUT1* to facilitate increased glucose import [84]. |
| | ← TR | Myc | *Myc* is a transcriptional inducer of nealry every step of glycolysis, including *GLUT1/2/4* to facilitate increased glucose import [84]. |
| | ← Per | Glycolysis | We assume that basal *Glycolysis* must be active before the *Glycolysis_H* node can turn on. |
| | ← ComplProc | Glucose | *Glycolysis* requires glucose uptake from the envrironment, which is converted to pyruvate in the cytosol [80]. |
| | ← Cat | NADp_c | NAD$^+$ is converted to NADH at the glyceraldehyde 3-phosphate dehydrogenase (GAPDH) step of glycolysis [80]. Inhibition of *NAMPT*, an enzyme essential for NAD$^+$ biosynthesis leads to attenuation of glycolysis [83]. |

**Table S1f: Warburg module**

| | | | |
|---|---|---|---|
| | $\leftarrow$ Ind | ERK | *ERK1/2* phosphorylate *PKM2*, which aids the conversion of PKM2 from a tetramer to a monomer. In the nucleus this monomer leds acts as a protein kinase that phosphorylates histone H3 to boost the transcription of *c-Myc*, which then promotes glycolytic enzyme expression [85]. |
| | $\dashv$ Ind | p53 | *p53* slows down glycolysis by multiple mechanisms, including the attenuation of *HIF-1*$\alpha$ transactivation (moderate *p53* activation), *HIF-1*$\alpha$ degradation (high *p53*) [86], and induction of the *TIGAR* enzyme, which lowers fructose 2,6-bisphosphate step of glycolysis [84]. |
| NADp_c | **NADp_c** = not **Glycolysis_H** or **Fermentation** or **NADp_m** | | |
| | Met | | In mammalian cells intracellular $NAD^+$ levels typically fall between 0.2 and 0.5 mM [87]. $NAD^+$ is consumed by glycolysis but regenerated during pyruvate to lactate conversion (*Fermentation* node in our model). In addition, we assume that mitochodtial $NAD^+$ can replenish cytosolic $NAD^+$ via the malate-aspartate shuttle [88]. |
| | $\dashv$ Cat | Glycolysis_H | $NAD^+$ is converted to NADH during the glyceraldehyde 3-phosphate dehydrogenase (GAPDH) step of glycolysis [80], lowering cytosolic $NAD^+$. |
| | $\leftarrow$ Cat | Fermentation | When cells convert pyruvate into lactate rather than process it in the mitochodtia to drive the TCA cycle, the process uses NADH to regenerate $NAD^+$ [89, 90]. |
| | $\leftarrow$ Loc | NADp_m | The malate–aspartate shuttle aids the translocaton of electrons produced during glycolysis across the mitochondrial inner membrane. Its net effect is to oxidize NADH in the cytosol to NAD+ (thus raising cytosolic NAD+ levels) at the expense of reducing mitochondrial NAD+ to NADH [88]. |
| Pyruvate | **Pyruvate** = **Glycolysis** or **Glycolysis_H** or **Pyruvate_Ext** | | |
| | Met | | Intracellular pyruvate is the product of glycolysis [89]. In addition, extracellular pyruvate can enter the cytosol via monocarboxylate transporters [91]. |
| | $\leftarrow$ ComplProc | Glycolysis | Intracellular pyruvate is the product of glycolysis [89]. |
| | $\leftarrow$ ComplProc | Glycolysis_H | Intracellular pyruvate is the product of glycolysis [89]. |
| | $\leftarrow$ Loc | Pyruvate_Ext | Extracellular pyruvate enters the cytosol via monocarboxylate transporters [91]. |
| HIF1 | **HIF1** = **mTORC1** and **eIF4E** and **S6K** and (**ERK** or not(**GSK3** and **FoxO3**)) and not **p53** | | |

**Table S1f: Warburg module**

| | | | |
|---|---|---|---|
| | TF | | Hypoxia-inducible factor 1, *HIF-1α*, is induced under normoxic conditions by growth-promoting signals to drive the Warburg effect in proliferating cells. First, *AKT*-mediated *mTORC1* and its downtream targets (*eIF4E*, *S6K*) increase both transcription and translation of *HIF-1α* [92]. Second, transcriptional activity of *HIF-1α* is boosted my MAPK signaling through *ERK*-mediated phosphorylation [93], while freed from *GSK3β*-mediated degradation and *FoxO3* binding by deactivation of these repressors downstream of *PI3K/AKT* [94, 95]. In contrast, *p53* represses *HIF-1α*-mediated transactivation [86]. |
| | ← Ind | mTORC1 | *mTORC1* activation downtream of *AKT* increases *HIF-1α* protein expression [96] protein accumulation through enhanced *STAT3* dependent transcription [92]. |
| | ← TL | eIF4E | *mTORC1*-induced *eIF4E* activation increases *HIF-1α* protein translation [92]. |
| | ← TL | S6K | *mTORC1*-induced *S6K* activation increases *HIF-1α* protein translation [92]. |
| | ← P | ERK | *ERK*-mediated phosphorylation of *HIF-1α* regulates its physical interaction with *NPM1*, a histone chaperone and chromatin remodeler. This interaction is increases *HIF-1α* targeting to hypoxia target genes and their transcriptional activation [93]. |
| | ⊣ P | GSK3 | *GSK3β* phosphorylates and destabolizes *HIF-1α*, independently of thye hypoxia sensor *VHL* [94]. |
| | ⊣ IBind | FoxO3 | *FoxO3* interferes with and reduces *p300*-dependent *HIF-1α* transcription [95]. |
| | ⊣ Ind | p53 | Moderate *p53* activation attenuates *HIF-1α* transactivation by competing for the shared *p300* co-activator [86]. |
| Fermentation | **Fermentation** = **ERK** and ((**Glycolysis_H** and **Pyruvate**) or **Pyruvate_Ext**) | | |
| | Proc | | In our model *Fermentation* = ON represents high rates of pyruvate to lactate conversion. This requires *ERK*-mediated phosphorylatiom of *PKM2*, which decreases its enzymatic activity (required for estabolishing the metabolic fluxes of the Warbugr effect) and boosts the expression of glycolitic genes [97]. In additiom, *Fermentation* turns ON with increased glycolysis and intracellular pyruvate, or in cells with access to extracellular pyruvate. |
| | ← Ind | ERK | *ERK*-mediated phosphorylatiom of *PKM2* converts this enzyme from a tetramer to a monomer, decreasing its catalytic activity (observed to promote increased pyruvate to lactate conversion) and allowing it to localize to the nucleus where it aids the transcription of *cyclin D1*, *c-Myc*, and glycolytic enzymes [97]. |
| | ← ComplProc | Glycolysis_H | Increased glycolysis coupled with low-activity *PKM2* drives the conversion of pyruvate to lactate [97]. |
| | ← ComplProc | Pyruvate | The *Fermentation* node represents the conversion of pyruvate to lactate. |

**Table S1f: Warburg module**

| Target Node | Link Type | Input Node | Link Description |
|---|---|---|---|
| | ← ComplProc | Pyruvate _Ext | Extracellular pyruvate enters the cytosol via monocarboxylate transporters [91], where its excess can drive fermentation. |

**Table S1g: Mitochondria module**

| Target Node | Node Gate / Node Type / Link Type | Node Description / Input Node | Link Description |
|---|---|---|---|
| TCA_cycle | **TCA_cycle** = **Pyruvate** and (**NADp_m** or not **MP_Low**) and not(**Unfused** and **mROS**) and not **cROS_H** | | |
| | Proc | The tricarboxylic acid (TCA) cycle, also known as the citric acid or Krebs (or Szent-Györgyi-Krebs) cycle, consists of a series of reactions driven by pyruvate-derived acetyl-CoA. In eukaryotic cells, the TCA cycle occurs in the matrix of mitochondria. This cycle feeds the Electron Transport Chain (ETC) by transfering electrons to $NAD^+$ carriers (thus using $NAD^+$ to generate NADPH). The ETC, in turn, raises the $\Delta\Psi_M$. In our model we ommited an explicit ETC node, and linked the TCA cycle to $\Delta\Psi_M$ ($TCA\_Cycle \longrightarrow MP\_High$ and $TCA\_Cycle \dashv MP\_Low$). | |
| | ← ComplProc | Pyruvate | The TCA cycle is driven by pyruvate-derived acetyl-CoA. |
| | ← ComplProc | NADp_m | Two steps of the TCA cycle require $NAD^+$ (mitochondrial $NAD^+$ is represented in the model by *NADp_m*). |
| | ⊢ ComplProc | MP_Low | The TCA cycle removes electrons from acetyl-CoA and transfers them to carriers that bring them to the electron transport chain (ETC), which in turn generates the mitochondrial membrane potential. As the oxidized electron carriers required to continue to turn the cycle are regenerated by the ETC (e.g., $NAD^+$), we assume that the TCA cycle works normally if the $\Delta\Psi_M$ is not low, or the mitochondria still have $NAD^+$. |
| | ⊢ ComplProc | Unfused | Inhibition of fusion, such as *Mfn2* knockdown, leads to fragmented mitochondria with reduced pyruvate, glucose and fatty acid oxidation, and reduces mitochondrial membrane potential [98]. Thus, we assume that unfused mitochondria negatively impact the TCA cycle. That said, experiments in cycling cells indicate that while the speed of the TCA cycle orcillates along with the cell cycle, the main change is an increased glucose-derived flux into TCA cycle in G1. Notably, the *short-term* mitochondrial fragmentation during metaphase does not significantly alter the TCA cycle [99]. To model this lack of disruption, we assume that only unfused mitochondria with high mitochodtial ROS (a consequence of fission triggered by stressors such as hyperglycemic challenge [100]) have significanly disrupted TCA cycles. |

**Table S1g: Mitochondria module**

| | | | |
|---|---|---|---|
| | $\vdash$ ComplProc | mROS | Mitochondrial ROS can compromose the TCA cycle by blocking two key enzymes [101, 102]. First, the enzyme aconitase, which catalyzes the conversion of citrate to isocitrate, is vulnerable to reactive oxygen and reactive nitrogen species [103]. Second, $\alpha$ ketoglutarate dehydrogenase catalyzes conversion of $\alpha$-ketoglutarate and coenzyme A to succinyl coA, converting $NAD^+$ to NADH in the process. This enzyme also has reduced activity in the presence of ROS [104]. |
| | $\vdash$ ComplProc | cROS_H | $H_2O_2$ exposure revealed that $\alpha$ ketoglutarate dehydrogenase (*KGDH*), succinate dehydrogenase (*SDH*), and aconitase were all inactivated by ROS [102]. |
| MFN1_2 | **MFN1_2** = (**PGC1**or**E2F1**)andnot(**MP_Low**and**PINK1**and**Mitophagy_High**)and not **Drp1** | | |
| | GTPa | | The mitofusins *MFN1* and *MFN2* facilitate fusion of the outer mitochondrial membrane [105]. Their mRNA levels are increased by *PGC-1*$\alpha$ and/or *E2F1*. In contrast, low $\Delta\Psi_M$ on mitochondria subject to mitophagy triggers the activity of *PINK1*, a kinase that recruits the ubiquitin E3 ligase *Parkin* to target both *MFN*s [106, 107]. In our model, we assume that in the presence of low $\Delta\Psi_M$, high mitophagy and PINK1, *E2F1* or *PGC1* cannot accumulate sufficient *MFN1* and *MFN2* to upregulate them above their basal levels. Finally, *Drp1* recruits the fission protein *Fis1*, which binds to *Mfn1/2* and directly inhibits its GTPase activity [108]. |
| | $\leftarrow$ TR | PGC1 | *PGC-1*$\alpha$ is a direct transcriptional inducer of both *MFN1* [109] and *MFN2* [110, 111] by co-activating the estrogen-related receptor $\alpha$. |
| | $\leftarrow$ TR | E2F1 | *E2F1* is a direct transcriptional inducer of *MFN2* [112]. |
| | $\vdash$ ComplProc | MP_Low | Low $\Delta\Psi_M$ on mitochondria subject to mitophagy triggers the activity of the *PINK1* kinase, which recruits the ubiquitin E3 ligase *Parkin* to target both *MFN*s [106, 107]. |
| | $\vdash$ ComplProc | Mitophagy _High | Mitophagy triggers the activity of *PINK1* kinase, which recruits the ubiquitin E3 ligase *Parkin* to target both *MFN*s [107]. |
| | $\vdash$ Deg | PINK1 | Active *PINK1* kinase recruits the ubiquitin E3 ligase *Parkin* to target both *MFN*s [107]. |
| | $\vdash$ Ind | Drp1 | *Drp1* recruits the fission protein *Fis1*, which binds to *Mfn1*, *Mfn2*, and *OPA1* and inhibits their GTPase activity, blocking the fusion machinery [108]. |
| Hyperfused _OM | **Hyperfused_OM** = **MFN1_2** and not(**Unfused** or **Drp1** or (**PINK1** and **Mitophagy_High**)) | | |

**Table S1g: Mitochondria module**

| | | | |
|---|---|---|---|
| | MSt | In mammalian cells, mitochondrial fusion is a two-step process in which outer mitochondrial membrane fusion is not always followed by inner membrane fusion [113] (the latter is required for boosting the $\Delta\Psi$ of the fused structure [114]). Outer membrane fusion is driven by mitofusins *MFN1/2* [113] and blocked by the fission factor *Drp1*, as well as *PINK1*-mediated strong mitophagy [107]. In our model, a hyperfused state is also mutually exclusive with the fragmented state, represented by the *Unfused* node. | |
| | ← Compl | MFN1_2 | *MFN*s facilitate mitochondrial fusion [115]. Specifically *MFN1/2* facilitate outer mitochondrial membrane fusion, which can be uncoupled from inner membrane fusion in mammalian cells [113]. |
| | ⊢ Per | Unfused | As we model three distinct overall mitochondrial morphology states with two Boolean nodes we set *Unfused* = ON and *Hyperfused_OM* = ON to be mutually exclusive. Thus, a cell with *Unfused* (fragmented) must pass through a normal (*Unfused* = OFF and *Hyperfused_OM* = OFF) state before undergoing hyperfusion. |
| | ⊢ IBind | Drp1 | *Drp1* is a main driver of mitochondrial fission [116]. When recruited to mitochondria, *Drp1* dimers remodel the mitochondrial membrane, while *Drp1* multimers promote its GTPase activity and induce fission [117]. |
| | ⊢ ComplProc | PINK1 | Low $\Delta\Psi$ on mitochondria undergoing mitophagy triggers the activity of the *PINK1* kinase, which recruits the ubiquitin E3 ligase *Parkin* to target both *MFN*s [107]. |
| | ⊢ Ind | Mitophagy_High | As *MFN1/2* ubiquitinated in a *PINK1/Parkin*-dependent manner upon induction of mitophagy [106], here we assume that high levels of mitophagy locally block low-$\Delta\Psi_M$ mitochondria from undergoing *MFN*-induced outer membrane fusion. |
| Hyperfused_IMOM | **Hyperfused_IMOM** = **Hyperfused_OM** and **MFN1_2** and not(**MP_Low** or **Unfused** or **Drp1**) | | |
| | MSt | Fusion of the inner mitochondrial membrane requires outer membrane fusion maintained by *MFN1/2*, but blocked by active *Drp1* [116, 117] or low $\Delta\Psi_M$ [118]. In addition, we assume that cell-wide fragmentation of mitochodrtia needs to be resolved to a normal state before the transition to a connected hyperfused mitochondrial network. | |
| | ← ComplProc | Hyperfused_OM | Fusion of the outer mitochondrial membrane, maintained by *MFN1/2*, is required for *Opa1* cleavage and activation, and is thus a prerequisite for inner membrane fusion [118]. |
| | ← Compl | MFN1_2 | Fusion of the outer mitochondrial membrane, maintained by *MFN1/2*, is required for *Opa1* cleavage and activation, and is thus a prerequisite for inner membrane fusion [118]. |

**Table S1g: Mitochondria module**

| | | | |
|---|---|---|---|
| | $\dashv$ IBind | MP_Low | *Opa1* cleavage and activation requires OXPHOS-stimulated activation of *Yme1L*, which cleaves *Opa1* more efficiently under high OXPHOS conditions [118]. We model this by requiring normal or high $\Delta\Psi_M$ as a condition of inner membrane hyperfusion. In contrast, outter membrane fusion is insensitive to $\Delta\Psi_M$ and OXPHOS [118]. |
| | $\dashv$ IBind | Drp1 | *Drp1* is a main driver of mitochondrial fission [116, 117] and thus counteracts both inner and outer membrane fusion. |
| | $\dashv$ Per | Unfused | As we model three distinct overall mitochondrial morphology states with two Boolean nodes, we set *Unfused* = ON and *Hyperfused_OM* = ON to be mutually exclusive. Thus, a cell with *Unfused* (fragmented) must pass through a normal (*Unfused* = OFF and *Hyperfused_OM* = OFF) state before undergoing hyperfusion. |
| MP_High | **MP_High** = **TCA_cycle** and **Hyperfused_IMOM** and not(**MP_Low** or **Unfused** or **mROS** or **cROS** or **cROS_H** or **ROS_Ext**) | | |
| | MP | The high $\Delta\Psi_M$ able to drive proliferation requires a funcioning TCA cycle / ETC, a hypefused network in which the inner membranes are also fused [119, 120], as well as the absence of excess mitochondrial, cytosolic, or external ROS [121]. | |
| | $\leftarrow$ ComplProc | TCA_cycle | The TCA cycle feeds the Electron Transport Chain (ETC) by transfering electrons to $NAD^+$ carriers, and thus increases $\Delta\Psi_M$. |
| | $\leftarrow$ ComplProc | Hyperfused_IMOM | Mitochondrial inner membrane fusion raises the efficiency of OXPHOS and increases the network's $\Delta\Psi_M$ [119]. We assume that the high $\Delta\Psi_M$ required for *Cyclin E* accumulation and S-phase entry requires such hyperfusion [120]. |
| | $\dashv$ Per | MP_Low | As we model three distinct overall $\Delta\Psi_M$ levels with two Boolean nodes, we set *MP_High* = ON and *MP_Low* = ON to be mutually exclusive. Thus, a cell with low $\Delta\Psi_M$ (*MP_Low*) must pass through a state with normal $\Delta\Psi_M$ (*MP_Low* = OFF and *MP_High* = OFF) before achieving high $\Delta\Psi_M$. |
| | $\dashv$ ComplProc | Unfused | As mitochondrial inner membrane fusion raises the efficiency of OXPHOS and increases the network's $\Delta\Psi_M$ [119], we assume that a cell with fragmented overall mitochondrial morphology cannot have high $\Delta\Psi_M$. |
| | $\dashv$ ComplProc | mROS | Exposing mitochondria to excessive oxidative stress can push them past a threshold that triggers the opening of the mitochondrial permeability transition (MPT) pore and/or the inner membrane anion channel (IMAC). This results in the simultaneous collapse of the $\Delta\Psi_M$, and a transient increased mitochondrial ROS generation by the ETC. This mitochondria-derived ROS can be released into cytosol, where it can trigger "Reactive Oxygen Species (ROS)-induced ROS release" (RIRR) in neighboring mitochondria [121]. |

**Table S1g: Mitochondria module**

| | | | |
|---|---|---|---|
| | $\vdash$ ComplProc | cROS | Excessive cytosolic ROS can push mitochondria past a threshold that triggers the opening of the MPT and/or the IMAC pore(s), which collapses the $\Delta\Psi_M$ [121]. |
| | $\vdash$ ComplProc | cROS_H | Excessive cytosolic ROS can push mitochondria past a threshold that triggers the opening of the MPT and/or the IMAC pore(s), which collapses the $\Delta\Psi_M$ [121]. |
| | $\vdash$ ComplProc | ROS_Ext | Excessive cytosolic ROS can push mitochondria past a threshold that triggers the opening of the MPT and/or the IMAC pore(s), which collapses the $\Delta\Psi_M$ [121]. |
| MP_Low | **MP_Low** = **BAX** or **BAK** or **cROS_H** or (not **MP_High** and ((not **TCA_cycle** and **NADp_m**) or (**Unfused** and **mROS**) or (**Hyperfused_OM** and not **Mitophagy_High** and **mROS** and not(**MnSOD** and **SIRT3**)) or (**MP_Low** and **mROS** and not(**Mitophagy_High** or **PGC1**)))) | | |
| | MP | | Low $\Delta\Psi_M$ can arise from a variety of processes: i) permeabilization of the outter membrane by *BAX/BAK* activation [122, 123] or by excessive cytosolic ROS [121], ii) lack of mitochondrial $NAD^+$ and no ongoing generation by the TCA cycle, iii) unfused ROS-producing mitochondria [124], iv) a hyperfused network barred from mitophagy [125] and damaged by ROS production, not mitigated by *MnSOD* [126] and *SIRT3* [127, 40], or v) mitochondria with low $\Delta\Psi_M$ and high ROS production, not rescued by *PGC-1*$\alpha$-mediated biogenesis along with high mitophagic flux [128]. |
| | $\leftarrow$ ComplProc | BAK | *BAK* translocation to the mitochondrial membrane can induce the loss of $\Delta\Psi_M$ [122]. |
| | $\leftarrow$ ComplProc | BAX | *BAX* translocation to the mitochondrial membrane can induce the loss of $\Delta\Psi_M$[123]. |
| | $\leftarrow$ ComplProc | cROS_H | Excessive cytosolic ROS can push mitochondria past a threshold that triggers the opening of the MPT and/or the IMAC pore(s), which collapses the $\Delta\Psi_M$ [121]. Over time, excessive ROS also damages mtDNA. |
| | $\vdash$ Per | MP_High | As we model three distinct overall $\Delta\Psi_M$ levels with two Boolean nodes, we set *MP_Low* = ON and *MP_High* = ON to be mutually exclusive. Thus, a cell with high $\Delta\Psi_M$ (*MP_High*) must pass through a state with normal $\Delta\Psi_M$ (*MP_Low* = OFF and *MP_High* = OFF) before crashing to a low $\Delta\Psi_M$. |
| | $\vdash$ ComplProc | TCA_cycle | The TCA cycle feeds the Electron Transport Chain (ETC) by transfering electrons to $NAD^+$. The ETC, in turn, raises $\Delta\Psi_M$. |
| | $\leftarrow$ ComplProc | NADp_m | As the ETC requires NADH to increase $\Delta\Psi_M$, high mitochondrial $NAD^+$ in the *absence* of ongoing TCA cycle flux is expected to crash $\Delta\Psi_M$. |
| | $\leftarrow$ ComplProc | Unfused | Cells with severely fragmented mitochondrial generally have low $\Delta\Psi_M$, though there is some heterogeneity among individual mitochondria [124]. |

**Table S1g: Mitochondria module**

| | | | |
|---|---|---|---|
| | ← ComplProc | mROS | Exposing mitochondria to excessive oxidative stress can push them past a threshold that triggers the opening of the mitochondrial permeability transition (MPT) pore and/or the inner membrane anion channel (IMAC). This results in the simultaneous collapse of the $\Delta\Psi_M$, and a transient increased mitochondrial ROS generation by the ETC. |
| | ← ComplProc | Hyperfused _OM | As fusion of the inner membrane is needed to raise the efficiency of OXPHOS in hyperfused mitochodtia [119], we assume that outer membrane fusion in mitochondria that generate excess ROS and cannot detoxify it with the aid of both *MnSOD* and *SIRT3* become incapable of sustaining normal $\Delta\Psi_M$. |
| | ⊢ ComplProc | Mitophagy _High | High levels of mitophagy can remove damaged mitochondria and aid the reestablishment of $\Delta\Psi_M$ within its nominal range [125]. |
| | ⊢ ComplProc | MnSOD | *MnSOD* detoxifies the superoxide anions ($O_2^{\cdot -}$) generated by the ETC by converting them to $H_2O_2$, which is further reduced to water by *Gpx-1*. Mitochondria from mice heterozygous for *MnSOD* show increased proton leak, inhibition of respiration, and early and rapid accumulation of mitochondrial oxidative damage [126], while complete loss of *MnSOD* is embryonic lethal [129]. |
| | ⊢ ComplProc | SIRT3 | *SIRT3* activation increases oxidative phosphorylation [127]. It deacetylates several critical enzymes involved the TCA cycle (*IDH2*) and OXPHOS (*SDHB*, *complex I*, *II* and *V*), boosting their activity [40]. |
| | ← Per | MP_Low | We assume that low $\Delta\Psi_M$ in mitochondria with excess mROS does not recover in the absence of high levels of mitophagy, or *PGC-1*$\alpha$-mediated biogenesis. |
| | ⊢ ComplProc | PGC1 | *PGC-1*$\alpha$ works together with *NRF1/2* and mitochondrial transcription factor A (*TFAM*) to increase mitochondrial biogenesis and boost mitochondrial energy metabolism [128]. |
| ATP_Low | **ATP_Low** = not **MP_Low** or (**Glycolysis** and **Fermentation**) or (**Autophagy** and **PGC1**) | | |
| | Met | | The *ATP_Low* node in our model represents an ATP/ADP ratio required for avoiding necrotic cell death [130] (not modeled), but not sufficient to keep AMPK inactive. This requires ATP production by OXPHOS (i.e., at least normal $\Delta\Psi_M$), glycolysis coupled to fermentation, or autophagy coupled with *PGC-1*$\alpha$ activity. |
| | ⊢ ComplProc | MP_Low | We assume that mitochondria with cell-wide low $\Delta\Psi_M$ do not contribute to ATP production. |
| | ← ComplProc | Glycolysis | In the absence of normal $\Delta\Psi_M$, glycolysis coupled with fermentation can support ATP production. |
| | ← ComplProc | Fermentation | In the absence of normal $\Delta\Psi_M$, glycolysis coupled with fermentation can support ATP production. |

**Table S1g: Mitochondria module**

| | | | |
|---|---|---|---|
| | ← ComplProc | Autophagy | The process of autophagy breaks down existing cytoplasmic components to support ATP production under starvation [131, 132]. |
| | ← ComplProc | PGC1 | *PGC-1α* induces mitochondrial biogenesis [133], as well as mitochondrial fatty acid oxidation to boost ATP levels under fasting conditions [134]. |
| ATP | **ATP** = **ATP_Low** and ((not **MP_Low**) or (**Glycolysis_H** and **Fermentation**)) | | |
| | Met | The *ATP* node represents normal ATP/ADP ratios that keep AMPK inactive. While at least low ATP (*ATP_Low* node) is a prerequisite, maintaining nominal ATP levels also requires normal $\Delta\Psi_M$, or high levels of glycolysis coupled to fermentation. | |
| | ← Per | ATP_Low | As the *ATP* node represents normal ATP/ADP ratios, we designed the model suych that it cannot be ON while the node representing survival-level ATP (*ATP_Low*) is OFF. |
| | ⊢ ComplProc | MP_Low | We assume that mitochondria with cell-wide low $\Delta\Psi_M$ do not contribute to ATP production. |
| | ← ComplProc | Glycolysis_H | High levels of glycolysis coupled with fermentation can support normal ATP production. |
| | ← ComplProc | Fermentation | High levels of glycolysis coupled with fermentation can support normal ATP production. |
| ATP_H | **ATP_H** = **ATP** and ((not **MP_Low** and **MP_High** and **Hyperfused_IMOM**) or (**Glycolysis_H** and **Fermentation** and **Pyruvate_Ext**)) | | |
| | Met | The *ATP_High* node represents the increased ATP/ADP ratios required to enter S-phase [120]. While at least normal ATP (*ATP* node) is a prerequisite, maintaining high ATP levels also requires high $\Delta\Psi_M$ maintained by a hyperfused network of mitochondria [120], or high levels of glycolysis coupled to fermentation supported by excess external pyruvate. | |
| | ← Per | ATP | As the *ATP_High* node represents high ATP/ADP ratios, we designed the model suych that it cannot be ON while the node representing normal ATP levels (*ATP*) is OFF. |
| | ⊢ ComplProc | MP_Low | We assume that mitochondria with cell-wide low $\Delta\Psi_M$ do not contribute to ATP production. |
| | ← ComplProc | MP_High | Cell-wide high $\Delta\Psi_M$ can produce the high ATP levels required for S-phase entry [120]. |
| | ← ComplProc | Hyperfused_IMOM | A hyperfused mitochondrial state is required for the elevated ATP production leading to S-phase entry [120]. As inner membrane fusion does not occur at low $\Delta\Psi_M$ and helps raise $\Delta\Psi_M$ [119], we assume this hyperfusion invovles both membranes. |
| | ← ComplProc | Glycolysis_H | Culturing cells with compromised mitochondria in excess pyruvate rescues cell cycle progression [135]. We assume that conversion of excess pyruvate to lactate can maintain the cytosolic $NAD^+$/NADH ratios required to support high levels of glycolysis, which in turn can raise ATP levels to promote S-phase entry even in teh absence of high $\Delta\Psi_M$. |

**Table S1g: Mitochondria module**

| | | | |
|---|---|---|---|
| | ← ComplProc | Fermentation | The conversion of pyruvate to lactate regenerates cytosolic $NAD^+$, required to maintain high rates of glycolysis. When mitochondrial $NAD^+$ production (NADH use) is compromised by the lack of an active ETC, the cell's NADH shuttles saturate and fermentation is used to regenerate $NAD^+$ [136]. |
| | ← ComplProc | Pyruvate _Ext | Culturing cells with compromised mitochondria in excess pyruvate rescues cell cycle progression [135], presumably by allowing rapid $NAD^+$ generation to support high levels of glycolysis. |
| AMPK | **AMPK** = not(**ATP_H** or **MP_High**) and (not **ATP** or **mROS** or **cROS** or **cROS_H**) | | |
| | K | | The cellular energy sensor AMP-activated protein kinase (*AMPK*) is activated by AMP binding in cells with low ATP/AMP ratios, leading to its allosteric activation and Thr172 phosphorylation by *LKB1* kinase [137, 1]. Thus, we assume *AMPK* blocked by high ATP levels or high $\Delta\Psi_M$, while the lack of normal ATP levels activates it. In addition, mitochondrial and cellular ROS was also shown to activate *AMPK*, promoting its antioxidant functions[138]. |
| | ⊢ Ind | ATP_H | High ATP to AMP rations inhibit AMP binding to *AMPK*, blocking its allosteric activation and Thr172 phosphorylation [139]. |
| | ⊢ Ind | MP_High | High $\Delta\Psi_M$ can block *AMPK* localized to the mitochondrial outer membrane via a localized ATP boost [140]. |
| | ⊢ Ind | ATP | We assume that nominal ATP to AMP rations keep AMP binding to *AMPK* low, blocking its allosteric activation and Thr172 phosphorylation [139]. |
| | ← Loc | mROS | Mitochondrial ROS was shown to directly activate *AMPK*, triggering a *PGC-1*$\alpha$-dependent antioxidant response to limit mitochondrial ROS [138]. |
| | ← Ind | cROS | Cells cultured in Trolox, an antioxidant that reduces cellular ROS levels displayed reduced AMPK activation (Thr172 phosphorylation) [138]. |
| | ← Ind | cROS_H | Cells cultured in Trolox, an antioxidant that reduces cellular ROS levels displayed reduced AMPK activation (Thr172 phosphorylation) [138]. |
| PGC1 | **PGC1** = **AMPK** | | |
| | TF | | Peroxisome proliferator-activated receptor-gamma coactivator *PGC-1*$\alpha$ is a transcription coactivator with key roles in the regulation of energy metabolism and antioxidant response [141]. In our model, activation of *PGC-1*$\alpha$ requires *AMPK*, or a joint boost from *mTORC1* [142] and *SIRT3* deacetylation [40]. (Note that we choose to omit *mTORC1*-mediated *PGC-1*$\alpha$ activation, as this would force our model to activate *MFN1/2* any time *mTORC1* was active, a response that does not match observed cell behavior. Overall, it appears that *mTORC1* balances the induction of mitochondrial biogenesis downstream of *PGC-1*$\alpha$ with increased fission, rendering the network more dynamic but not hyperfused.) |
| | ← Ind | AMPK | Increased AMPK levels increase *PGC-1*$\alpha$ mRNA and protein levels [143, 144]. |

**Table S1g: Mitochondria module**

| | | | |
|---|---|---|---|
| SIRT3 | **SIRT3** = **NADp_m** and (**PGC1** or (**Hyperfused_IMOM** and not **MP_Low**)) | | |
| | Enz | The mitochondrial NAD-dependent deacetylase *SIRT3* helps maintain ATP levels by regulating mitochondrial electron transport [127], boosting antioxidant responses via deacetylaiton of *MnSOD*, *NRF2* and *PGC-1α* (which also promotes mitochondrial biogenesis) [40]. In our model, *SIRT3* activation requires mitochondrial $NAD^+$ and either *PGC-1α*-mediated transcription [145] or mitochondrial hyperfusion with at least nominal $\Delta\Psi_M$. (Note that we do not model the localized homeostatic role of *SIRT3*, where it its activity is locally boosted by the lowering of $\Delta\Psi_M$ in order cause rapid deacetylation of matrix proteins to recover membrane potential [146]. This effect, however, implicitly requires no drastic loss of mitochodtial $NAD^+$, and thus cannot be a lasting condition.) | |
| | ← Cat | NADp_m | *SIRT3* activity requires mitochondrial $NAD^+$ [127]. |
| | ← TR | PGC1 | *PGC-1α* is a direct transcriptional inducer of *SIRT3* [145]. |
| | ← ComplProc | Hyperfused_IMOM | In the absence of a *PGC-1α*-mediated increase in *SIRT3* levels, we assume that a hyperfused mitochondrial network with strong ETC activity and at least normal $\Delta\Psi_M$ are required to keep mitochondrial $NAD^+$ levels high for sustained *SIRT3* activity. |
| | ⊣ ComplProc | MP_Low | In the absence of a *PGC-1α*-mediated increase in *SIRT3* levels, we assume that a hyperfused mitochondrial network with strong ETC activity and at least normal $\Delta\Psi_M$ are required to keep mitochondrial $NAD^+$ levels high for sustained *SIRT3* activity. |
| PINK1 | **PINK1** = **MP_Low** and (**NF_kB** or **FoxO1** or **FoxO3**) | | |
| | K | PTEN-induced putative kinase 1 *PINK1* phosphorylates ubiquitin to activate the ubiquitin ligase *Parkin*, which builds ubiquitin chains on mitochondrial outer membrane proteins to initiate mitophagy [147]. *PINK1* is activated by mitochondrial membrane depolarization [148], while its transcriptional activation (for increased levels of mitophagy modeled here) is induced by *NF-κB* or *FoxO1/3* [46, 149]. | |
| | ← Ind | MP_Low | *PINK1* is activated by mitochondrial membrane depolarization and stimulates Parkin E3 ligase activity by phosphorylating it at Ser65 [148]. |
| | ← TR | NF_kB | *NF-κB* is a direct transcriptional inducer of *PINK1*[150]. |
| | ← TR | FoxO1 | *FoxO1* is a transcriptional inducer of *PINK1* [46]. |
| | ← TR | FoxO3 | *FoxO3* is a transcriptional inducer of *PINK1* [149]. |
| Mitophagy_High | **Mitophagy_High** = (**PINK1** or **Drp1**) and not **Hyperfused_OM** and (**AMPK** or **MP_Low**) and (**FoxO3** or **FoxO1** or not **mTORC1**) and not **Casp8** | | |

#### Table S1g: Mitochondria module

| | | | |
|---|---|---|---|
| | Proc | | In out model, high (above-basal) mitophagy requires either *PINK1/Parkin* [147] or *Drp1* activation [151], lack of cell-wide hyperfusion of the outer mitochondrial membrane, either *AMPK* [152] or low $\Delta\Psi_M$ [153], and transcriptional upregulation of autophagy/mitophagy components by *FoxO1/3* [154] or *mTORC1* inhibition [46]. Finally, *Caspase 8* blocks high mitophagy by inhibiting *Parkin* [155]. |
| | ← Ind | PINK1 | *PINK1* phosphorylates ubiquitin to activate *Parkin*, which tags mitochondrial outer membrane proteins with ubiquitin chains to initiate mitophagy [147]. |
| | ← Ind | Drp1 | In addition to *PINK1/Parkin*, mitophagy can also be initiated by *Drp1* [151]. |
| | ⊣ Ind | Hyperfused _OM | mitochondrial hyperfusion in the absence of *Drp1*-mediated fission protects mitochondria from autophagosomal degradation [156, 157]. |
| | ← Ind | AMPK | *AMPK* promotes mitophagy by enhancing mitochondrial fission (*MFF* phosphorylation) and autophagosomal engulfment (*TBK1* activation). This effect does not require *PINK1/Parkin* [152]. |
| | ← Ind | MP_Low | Loss of $\Delta\Psi_M$ rapidly recruits *Parkin* from the cytosol to mitochondria, boosting mitophagy [153]. |
| | ← Ind | FoxO1 | *FoxO1/3* regulate both ubiquitin-dependent and receptor-mediated mitophagy by upregulating *PINK1* and *Bnip3/Bnip3L*, respectively. In addition, *FoxO1/3* induce genes involved in each stage of autophagosome formation, including initiation (e.g. *Ulk1/Ulk2*), nucleation (e.g. *Becn1*, *Atg14*, and *Pi3kIII*), elongation (e.g. *Map1lc3b*, *Gabarapl,Atg4/5/12*), and fusion (e.g. *Tfeb* and *Rab7*) [154]. |
| | ← Ind | FoxO3 | *FoxO1/3* regulate both ubiquitin-dependent and receptor-mediated mitophagy, as well as each stage of autophagosome formation [154]. |
| | ⊣ Ind | mTORC1 | Constitutively active *mTORC1* in *TSC2*-/- cells impairs mitophagy, while its inhibition reestablishes it [46]. |
| | ⊣ Ind | Casp8 | *Caspase 8* cleaves and inhibits *Parkin* during apoptosis, a key player in the initiation of mitophagy [158]. Decreased *Parkin* levels block mitophagy and enhance apoptosis [155]. |
| Drp1 | **Drp1** = (**BIK** and (**BAK** or **BAX**)) or (not **p53** and ((**CyclinB** and **Cdk1** and not **Cdh1**) or (**AMPK** and not **MFN1_2**) or (**MP_Low** and **PINK1** and **Mitophagy_High**))) | | |
| | GTPa | | Dynamin-related protein 1 *Drp1* is required for mitochondrial fission [151]. *Drp1* is recruited to the mitochondrial outer membrane by several stimuli, including apoptosis (via *BIK*-mediated recruitment [159] and *BAK/BAX*-mediated retention [160]), *Cdk1/Cyclin B*-induced mitotic phosphorylation [161] counteracted by $APC/C^{Cdh1}$ [162, 163], or low $\Delta\Psi_M$ [164] coupled to mitophagy [165, 166, 167] and aided by *AMPK* [168]. In contrast, *p53* inhibits *Drp1* activity [169]. |

**Table S1g: Mitochondria module**

| | | |
|---|---|---|
| ← Ind | BIK | *BIK* activates the transmission of Ca(2+) to the mitochondria from the ER, which in turn recruits *Drp1* to the mitochondrial surface [159]. |
| ← Ind | BAK | *BAK* translocation to the mitochondria increases mitochondria-bound *Drp1* by sumoylation of *Drp1*, which subsequently accumulates on mitochondrial membranes after the completion of fission, but before the initiation of cytochrome C release [160]. |
| ← Loc | BAX | *BAX* translocation to the mitochondria increases mitochondria-bound *Drp1* by sumoylation of *Drp1*, which subsequently accumulates on mitochondrial membranes after the completion of fission, but before the initiation of cytochrome C release [160]. |
| ⊢ Ind | p53 | *p53* increased inhibitory phosphorylation of *Drp1* at Ser637, an effect mediated by Protein kinase A but did not require increases *p21* [169]. |
| ← P | CyclinB | *Drp1* is phosphorylated on Ser585 by *Cdk1/Cyclin B* complexes on during mitosis, which is required for effective mitochondrial fragmentation during mitosis [161]. |
| ← P | Cdk1 | *Drp1* is phosphorylated on Ser585 by *Cdk1/Cyclin B* complexes on during mitosis, which is required for effective mitochondrial fragmentation during mitosis [161]. |
| ⊢ Deg | Cdh1 | At the end of mitosis, $APC/C^{Cdh1}$ ubiquitylates *Drp1*, leading to its degradation [162, 163]. |
| ← Ind | AMPK | Under energy stress, *AMPK* phosphorylates *DRP1*'s mitochondrial outer-membrane receptor, fission factor *MFF*, aiding the translocation of *DRP1* from the cytosol to the mitochondrial membrane [163]. That said, the long-term indirect effects of sustained *AMPK* activation likely counteract the rapid fragmentation caused by this event, as *AMPK* activation was also linked to a shift in *Drp1* phosphorylation from Ser616, which accelerated its recruitment to the mitochondrial membrane, to Ser637, which hinders it [168]. |
| ← Ind | MP_Low | *Drp1* recruitment to damaged mitochondria is mediated by a $\Delta\Psi_M$ 'tipping point', below which mitochondria with functional *Drp1* undergo rapid fission [164]. |
| ← P | PINK1 | *PINK1* directly phosphorylates *Drp1* on S616, which accelerated its recruitment to the mitochondrial membrane [165]. |
| ← Ind | Mitophagy_High | Mitophagy results in the translocation of mitochondrial E3 ubiquitin ligase *MITOL/March5* from depolarized mitochondria to peroxisomes [166]. This potentiates *Drp1* function, as *MITOL/March5* mediates *Drp1* clearance from the mitochondrial membrane via by outer mitochondrial membrane-associated degradation (OMMAD) [167]. |

**Table S1g: Mitochondria module**

| | | | |
|---|---|---|---|
| | ⊢ Ind | MFN1_2 | We assume that in the absence of additional influences beyond *AMPK*-mediated *Drp1* activation, high levels of *MDN1/2* balance out increased *Drp1* activity. |
| Unfused | **Unfused** = not **Hyperfused_OM** and ((**Drp1** and not(**MFN1_2** and **Mitophagy_High**)) or **BAK** or **BAX**) | | |
| | MSt | | Our model captures overall (cell-wide) mitochondrial morphology in three stages. First, the fragmented state is represented by the *Unfused* = ON node state. Second, the "normal" morphology of a quiescent cell is represented by *Unfused* = OFF and *Hyperfused_OM* = OFF, while the third, hyperfused state is marked by *Hyperfused_OM* = ON. (We further distinguish between outer membrane vs. outer and inner membrane fusion). |
| | ⊢ Per | Hyperfused _OM | As we model three distinct overall mitochondrial morphology states with two Boolean nodes, we set *Unfused* = ON and *Hyperfused_OM* = ON to be mutually exclusive. Thus, a cell with hyperfused mitochondria must pass through a normal (*Unfused* = OFF and *Hyperfused_OM* = OFF) state before fully fragmenting. |
| | ← ComplProc | Drp1 | *Drp1* is required for mitochondrial fission [151]. |
| | ⊢ ComplProc | MFN1_2 | *MFN1/2* promote mitochondrial fusion, thus reducing the fraction of unfused mitochondria [113]. |
| | ⊢ Deg | Mitophagy _High | As mitophagy targets fragmented and damaged mitochondria, we assume that high levels of mitophagy act to reduce the fraction of unfused, damaged mitochondria. |
| | ← Ind | BAK | During *BAK/BAX*-dependent apoptosis, the mitochondrial network undergoes extensive fragmentation even in *DRP1*-null cells [170]. Thus, we assume that *BAK/BAX* alone can drive excessive fragmentation. |
| | ← Ind | BAX | During *BAK/BAX*-dependent apoptosis, the mitochondrial network undergoes extensive fragmentation even in *DRP1*-null cells [170]. Thus, we assume that *BAK/BAX* alone can drive excessive fragmentation. |
| mROS | **mROS** = **cROS_H** or (**Unfused** and (not(**MnSOD** or **SIRT3**) or **MP_Low** or **BAK** or **BAX**)) or (**MP_Low** and not(**PGC1** and (**Mitophagy_High** or **MnSOD** or **SIRT3**))) or (**Hyperfused_OM** and ((**MP_High** or **mROS**) and not(**MnSOD** and **SIRT3**))) | | |

**Table S1g: Mitochondria module**

| | | |
|---|---|---|
| Met | | Mitochondria are the main site of cellular ROS production [171]. In our model, we assume that this production exceeds the normal parameters of healthy mitochondrial function (including healthy proliferation) if one of the following conditions hold: i) cytosolic ROS levels are very high [172], ii) the mitochondrial network is fragmented (cell-wide) and is either missing both *MnSOD* and *SIRT3*, or has undergone *BAK/BAX*-mediated mitochondrial outer membrane permeabilization (MOMP) [173, 174, 126, 40, 175, 176]; iii) $\Delta\Psi_M$ is low, not counteracted by ongoing high mitophagy coupled with mitochondrial biogenesis (requiring *PGC-1*$\alpha$ activity), or by increased antioxidant *MnSOD* or *SIRT3* activity [177, 178, 179, 180]; iv) the mitochondrial network is hyperfused, with either high $\Delta\Psi_M$ (higher ROS production by the ETC) [181] or dysnfunctionally low $\Delta\Psi_M$ [182, 177]; both in the absence of the joint mitigating effects of increased *MnSOD* and *SIRT3* activity. |
| ← ComplProc | cROS_H | High levels of cytosolic ROS (e.g. hydrogen peroxide) can damage the ETC by triggering a mitochondrial permeability transition, which leads to mitochondrial complex I deficiency. As a result, the damaged ETC produces more superoxide [172]. |
| ← ComplProc | Unfused | Mitochondrial fragmentation mediated by fission increases ROS production [173, 174]. |
| ⊢ Cat | MnSOD | *MnSOD* transforms the toxic superoxide generated by the mitochondrial ETC into hydrogen peroxide (further transformed to water by catalase) and diatomic oxygen [126], and thus it is required for lowering mitochondrial ROS. |
| ⊢ Ind | SIRT3 | *SIRT3* boosts mitochondrial antioxidant responses by deacetylating components of the ETC to reduce ROS generation [127], as well as indirectly via deacetylaiton of *MnSOD*, *NRF2* and *PGC-1*$\alpha$ (which also promotes mitochondrial biogenesis) [40]. |
| ← ComplProc | BAK | *SIRT3* Mitochondrial outer membrane permeabilization (MOMP) by oligomerization of *BAK/BAX* was shown to increase ROS (superoxide) release along with Cytochrome C [175, 176]. |
| ← ComplProc | BAX | *SIRT3* Mitochondrial outer membrane permeabilization (MOMP) by oligomerization of *BAK/BAX* was shown to increase ROS (superoxide) release along with Cytochrome C [175, 176]. |
| ← ComplProc | MP_Low | Mitochondrial dysfunction, or the inability to maintain normal $\Delta\Psi_M$, is associated with increased mitochondrial ROS production [182, 177]. (Note that *transient* $\Delta\Psi_M$ reduction can shift the electron transport chain (ETC) into a more oxidized state and can temporarily reduce ROS production, but sustained cell-wide low $\Delta\Psi_M$ appears to increase mROS [177].) |
| ⊢ ComplProc | Mitophagy _High | As mitophagy targets damaged and dysfunctional mitochondria, high levels of mitophagy can lower overall mROS generation by selectively clearing the most prominent ROS producers [178, 179]. |

**Table S1g: Mitochondria module**

| | | | |
|---|---|---|---|
| | $\vdash$ ComplProc | PGC1 | *PGC-1$\alpha$* is a master transcriptional regulator of mitochondrial biogenesis, oxidative phosphorylation and reactive oxygen species detoxification [180]. *PGC-1$\alpha$* regulates the expression of mitochondrial antioxidant genes (*MnSOD*, catalase, peroxiredoxin 3 and 5, uncoupling protein 2, thioredoxin 2, and thioredoxin reductase) and lowers mROS production [128, 141]. |
| | $\leftarrow$ ComplProc | Hyperfused_OM | Hyperfused mitochondria can produce excess ROS under two conditions. First, if $\Delta\Psi_M$ is very high (G1/S transition), the ETC produces excess ROS that must be counterbalanced by ROS scavenging enzymes (*MnSOD*, catalase, peroxiredoxins, thioredoxins, and thioredoxin reductases) [181]. Alternately, mROS is maintained if $\Delta\Psi_M$ is very low [182, 177], the mitochondrial network is already producing excess mROS, and the inner membrane cannot fuse to boost $\Delta\Psi_M$ [118]. |
| | $\leftarrow$ ComplProc | MP_High | High $\Delta\Psi_M$ above a certain threshold ($\sim$ 140 mV) dramatically increases increase $H_2O_2$ production [181]. |
| | $\leftarrow$ Per | mROS | mROS damages ETC components, leading to dysfunction and further mROS generation. Moreover, exposing mitochondria to excessive oxidative stress can push them past a threshold that triggers the opening of the mitochondrial permeability transition (MPT) pore and/or the inner membrane anion channel (IMAC). This results in the simultaneous collapse of the $\Delta\Psi_M$, and a transient increased mitochondrial ROS generation by the ETC. This mitochondria-derived ROS can be released into cytosol, where it can trigger "Reactive Oxygen Species (ROS)-induced ROS release" (RIRR) in neighboring mitochondria [121]. |
| NADp_m | **NADp_m** = (not(**TCA_cycle** and **MP_Low**) and (**MP_High** or (**NADp_c** and not **mROS**))) or (**Pyruvate_Ext** and **NADp_c**) | | |
| | Met | | Mitochodtial $NAD^+$ is consumed by the TCA cycle and generated by the ETC [183]. Thus, mitochondrial $NAD^+$ in our model turns off if the TCA cycle is functional but the ETC is not (represented by *MP_Low* = ON). Along these lines, maintaining it requires no mROS damage to the ETC [184], and either high $\Delta\Psi_M$ [183] or normal cytosolic $NAD^+$. Alternatively, we assume that the presence of excess pyruvate can raise the cytosolic pool of $NAD^+$ to the point where the malate-aspartate shuttle slows to a halt [88], allowing mitochondria to recover their $NAD^+$ levels. |
| | $\vdash$ Ind | TCA_cycle | The TCA cycle consumes $NAD^+$ to generate NADH, which in turn carries eletrons to the ETC [183]. |
| | $\vdash$ Ind | MP_Low | The ETC's Complex I regenerates mitochondrial $NAD^+$ as it pumps an electron into the intermembranespace to increase the $\Delta\Psi_M$ [183]. We assume that the ETC is not working effectively in cells with low $\Delta\Psi_M$. |

**Table S1g: Mitochondria module**

| | | | |
|---|---|---|---|
| | ← Loc | MP_High | The ETC's Complex I regenerates mitochondrial $NAD^{+}$ as it pumps an electron into the intermembranespace to increase the $\Delta\Psi_M$ [183]. Thus, high $\Delta\Psi_M$ generation is coupled with mitochondrial $NAD^{+}$ production. |
| | ← Loc | NADp_c | As the net effect of the malate-aspartate shuttle is to oxydise cytosolic NADH to $NAD^{+}$ at the expence of reducing mitochondrial $NAD^{+}$ to NADH, this shuttle runs one-way: it replenishes cytosolic $NAD^{+}$ from the mitochondrial pool. Here we assume that if the cytosolic levels are high, the shuttle works slowly and allows mitochondrial $NAD^{+}$ levels to build up by de novo production of $NAD^{+}$ [185]. |
| | ⊣ Ind | mROS | mROS damages ETC compontents as well as mitochodrial DNA, blocking their ability to regenerate $NAD^{+}$ used up by the TCA cycle. Damage to Complex I in particular lowers mitochondrial $NAD^{+}$, which in turn blocks deacetlylation of *MnSOD* and further increases mROS [184]. |
| | ← Ind | Pyruvate _Ext | We assume that the increased pyruvate to lactate conversion triggered by access to external pyruvate increases cytosolic $NAD^{+}$ levels, allowign the malate-aspartate shuttle to slow and help mitochondrial $NAD^{+}$ levels to build up by de novo production of $NAD^{+}$ [185]. |

**Table S1h: Restriction_SW module**

| Target Node | Node Gate<br>Node Type<br>Link Type | <br>Node Description<br>Input Node | <br><br>Link Description |
|---|---|---|---|
| p21 | **p21** = **p21_mRNA** and (not **CyclinE** or **p53**) and not **Casp3** | | |
| | Prot | In this model, the *p21* node corresponds to nuclear p21 in cells with relatively high basal *p21* activity. *p21* activity and or localization can be lowered by loss of *FoxO* mediated transcription (see $p21_{mRNA}$ node) and via feedback from *Cyclin E/Cdk2* [186]. | |
| | ← TL | p21 _mRNA | *p21* protein activity requires the presence of *p21* transcription. |
| | ← TR | p53 | *p53*-mediated transcription of *p21* raises its levels sufficiently to override its repression by *Cdk2/Cyclin E*, resetting the switch to a high *p21*/low *Cdk2* state [187]. |
| | ⊣ Deg | CyclinE | *p21* and *Cyclin E/Cdk2* form a positive (double-negative) feedback loop in which *Cyclin E/Cdk2* activates the *SCF/Skp2* complex responsible for the degradation of *Cyclin E/Cdk2*-bound, phosphorylated *p21* [188]. *p21*, in turn, not only blocks *Cyclin E/Cdk2* activity, but it also inhibits *Cyclin D1*. Thus, *p21* interferes with the mitogen signal that turns on *Cyclin E* in the first place. In quiescent cells with high basal *p21* levels, this positive feedback renders cell cycle entry stochastic [186]. |

**Table S1h: Restriction_SW module**

| | | | |
|---|---|---|---|
| | $\dashv$ Lysis | Casp3 | *Caspase 3* cleaves and deactivates *p21* [189]. |
| pRB | **pRB** = (**p27Kip1** or not **CyclinE**) and not(**CyclinD1** or **CyclinA** or **Casp3**) | | |
| | TF | *pRB* is active in the absence of *Caspase 3*, *Cyclin D1*, *Cyclin A*, and *Cyclin E*. In addition, *pRB* maintains its activity when active $p27^{Kip1}$ counteracts the effects of *Cyclin E* [190, 191, 192, 193]. | |
| | $\leftarrow$ ComplProc | p27Kip1 | Active $p27^{Kip1}$ can counteract the inhibitory effects of active *CyclinE/Cdk2* complexes [190]. |
| | $\dashv$ P | CyclinE | *Cyclin E/Cdk2* complexes bind and phosphorylate *RB*, inhibiting its activity [194, 192]. |
| | $\dashv$ P | CyclinD1 | *Cyclin D1/Cdk4,6* complexes bind and phosphorylate *RB*, inhibiting its activity [195, 196, 192]. |
| | $\dashv$ P | CyclinA | *Cyclin A/Cdk1,2* complexes phosphorylate and deactivate *RB* [191]. |
| | $\dashv$ Lysis | Casp3 | *Caspase 3* cleaves *RB*, generating fragments that do not associate with *E2F1*, rendering *RB* inactive [197]. |
| p27Kip1 | **p27Kip1** = not **CyclinD1** and ((not **CyclinE** and **FoxO3** and **FoxO1**) or (not **CyclinA** and (**FoxO3** or **FoxO1**)) or not(**CyclinE** or **CyclinA**) or **AMPK**) and (not(**Cdk1** and **CyclinB**) or **AMPK**) and not **Casp3** | | |
| | Prot | Active $p27^{Kip1}$ is cleaved by *Caspase 3* and inhibited (sequestered) by *Cyclin D1/Cdk4,6* [190] or *Cyclin B/Cdk1* [198]. In addition, maintenance of $p27^{Kip1}$ requires one or both *FoxO* factors when sequestered by *Cyclin E/Cdk2* (one *FoxO* factor) or *Cyclin A/Cdk2* (both *FoxO* factors), but it cannot keep pace with the simultaneous activity of *Cyclin E/Cdk2* and *Cyclin A/Cdk2* [199] unless stabilized by *AMPK* [200]. | |
| | $\dashv$ IBind | CyclinD1 | Active *Cyclin D/Cdk4,6* complexes competitively bind to $p27^{Kip1}$ and progressively inhibit its ability to keep *Cyclin-E/Cdk2* inactive, thereby inducing cdk2 activity and cell-cycle progression [190]. |
| | $\dashv$ Deg | CyclinE | Active *Cyclin-E/Cdk2* phosphorylate $p27^{Kip1}$ at threonine 187 (Thr187) [201], which marks it for degradation by the $SCF^{SKP2}$ complex at the onset of S-phase [202]. (*Cyclin-E/Cdk2* complexes remain active in the presence of $p27^{Kip1}$ and promote its degradation when *Cyclin-A* is also active.) |
| | $\leftarrow$ TR | FoxO3 | *FoxO* factors are direct inducers of $p27^{Kip1}$ expression [203]. |
| | $\leftarrow$ TR | FoxO1 | *FoxO* factors are direct inducers of $p27^{Kip1}$ expression [203]. |
| | $\dashv$ Deg | CyclinA | *Cyclin A/Cdk2* complexes bind and inactivate $p27^{Kip1}$ by sequestration, phosphorylate it, and promote its degradation [204]. |
| | $\dashv$ P | AMPK | *AMPK*-mediated $p27^{Kip1}$ phosphorylation at Thr-198 stabilizes $p27^{Kip1}$ [200]. |

**Table S1h: Restriction_SW module**

| | | | |
|---|---|---|---|
| | $\vdash$ PLoc | CyclinB | *Cyclin B/Cdk1* complexes phosphorylate $p27^{Kip1}$ [204], and although they do not promote its degradation, phosphorylated $p27^{Kip1}$ is exported from the nuclear compartment and looses its ability to inhibit *Cdk* activity [198]. |
| | $\vdash$ PLoc | Cdk1 | *Cyclin B/Cdk1* complexes phosphorylate $p27^{Kip1}$ [204], and although they do not promote its degradation, phosphorylated $p27^{Kip1}$ is exported from the nuclear compartment and looses its ability to inhibit *Cdk* activity [198]. |
| | $\vdash$ Lysis | Casp3 | *Caspase 3* cleaves $p27^{Kip1}$ [205]; the cleaved fragments can no longer associate with *Cdk2 / Cyclin* complexes [206]. |
| Myc | **Myc** = not **p53** and ((**ERK** and (**eIF4E** or not **GSK3**)) or (**E2F1** and not **pRB** and (**eIF4E** or **ERK** or not **GSK3**))) | | |
| | TF | | *Myc* activity is turned on by stabilization of the protein via *ERK* phosphorylation, aided by either an increase in translation initiated by *eIF4E* or loss of degradation-promoting phosphorylation by *GSK3β* [207]. Alternatively, increased transcription by *E2F1* can also promote *Myc* accumulation in the absence of active *pRB* [208], provided that the protein is stabilized by *ERK*, *eIF4E*, or the absence of *GSK3β*. *p53* blocks transcription of *c-Myc*, enforcing G1 arrest independently of *p21* [209]. |
| | $\vdash$ TR | p53 | *p53* is a direct transcriptional repressor of the *c-Myc* promoter [209]. |
| | $\leftarrow$ P | ERK | Ser-62 phosphorylation by *ERK* increases its half life, leading to *Myc* accumulation [207, 210]. |
| | $\leftarrow$ TL | eIF4E | Increased translational initiation in the presence of activated *eIF4E* leads to an increase in *Myc* protein levels [211]. |
| | $\vdash$ P | GSK3 | Thr-58 phosphorylation by *GSK-3* promotes *Myc* degradation [207, 212]. |
| | $\vdash$ TR | pRB | *E2F1*'s ability to induce *Myc* is blocked by active (hypo-phosphorylated *pRB*) [213]. |
| | $\leftarrow$ TR | E2F1 | *E2F1* binds and activates the *c-Myc* promoter [214, 215]. |
| CyclinD1 | **CyclinD1** = not(**ATR** or **ATM** or **p21_H** or (**p21** and **p53**)) and ((not **p21** and ((not **GSK3** and **eIF4E** and (**Myc** or **E2F1**)) or (**Myc** and **CyclinD1**) or (**Myc** and **E2F1**) or (**E2F1** and **CyclinD1**))) or ((not **pRB** and **E2F1**) and ((**Myc** and **CyclinD1**) or (**Myc** and (not **GSK3** or **eIF4E**)) or (**CyclinD1** and (not **GSK3** or **eIF4E**))))) | | |

**Table S1h: Restriction_SW module**

| | | |
|---|---|---|
| PC | | Ongoing DNA synthesis keeps the *ATR* kinase active, which inhibits *Cyclin D1*. Similarly, double-strand DNA damage activates *ATM*, which also inhibits *Cyclin D1*. The precise regulatory logic of *Cyclin D1* as a function of transcriptional control by *Myc* and *E2F1*, combined with the regulation of its protein stability / activity by *GSK3β* / *p21* / *p53* is not known. Here, we assume that in the absence of *p21* (once *p21* levels drop due to growth factor signals and/or *Cdk2* activation), *Cyclin D1* can be activated by either *Myc* or *E2F1* in the absence of *GSK3β*. In the presence of *GSK3β*, we assume that *Cyclin D1* can be induced by the combined action of both *Myc* and *E2F1* [216], but sustained in an ON state by either. In the presence of basal (normal quiescent) levels of *p21*, we assume that *Cyclin D1* transcription requires *E2F1* unencumbered by *pRB*, as well as any two of the following: *Myc*, already active *Cyclin D1*, and no *GSK3β*. |
| ← TL | eIF4E | Increased translational initiation in the presence of activated *eIF4E* leads to an increase in *CyclinD1* protein levels [217]. |
| ⊢ P | GSK3 | *GSK-3β* phosphorylates *Cyclin D1* on Thr-286, promoting its ubiquitination and degradation [218]. |
| ⊢ IBind | p21 | *p21*$^{Cip1}$ is a Cyclin Dependent kinase inhibitor which binds to and blocks the activity of *Cdk2*, *Cdk3*, *Cdk4* and *Cdk6* kinases [219] and thus inhibits *CyclinD1/Cdk4,6* [220]. |
| ⊢ TR | pRB | *E2F1*'s ability to induce *Cyclin D1* is blocked by active (hypo-phosphorylated) *RB* protein [221]. |
| ← TR | Myc | Extracellular growth signals activate the MAPK pathway, leading to transcriptional activation of *Cyclin D1* by *Myc* [222, 223]. *Myc* overexpression leads to rapid *Cyclin D1* induction and subsequent cell cycle entry [224], while its absence halves *Cyclin D1* levels [225]. In addition, *Myc* induces *Cdk4*, aiding the assembly of active *Cyclin D1* / *Cdk4,6* complexes [225, 226]. |
| ← Per | CyclinD1 | In order to take into account both production and stability of *Cyclin D1*, we assumed that the presence of active *CyclinD/Cdk2,4* complexes renders transcriptional maintenance of their levels easier. |
| ← TR | E2F1 | The *Cyclin D1* promoter is bound by *E2F* factors including *E2F1* [227], and *E2F1* overexpression can increase *Cyclin D1* (though its effects are context-dependent, as *E2F1* overexpression can also lead to apoptosis) [227]. Dominant negative *E2F1* overexpression results in a 2-3 fold decrease in *Cyclin D* expression and *Cyclin D/Cdk4,6* activity [228]. |
| ⊢ P | ATR | During replication, checkpoint kinases such as *ATR* (active during normal DNA synthesis) suppress *Cyclin D1* by promoting it's Thr286 phosphorylation, leading to its degradation [229]. |
| ⊢ P | ATM | ATM to promotes cyclin D1 Thr286 phosphorylation, leading to its degradation [229]. |

**Table S1h: Restriction_SW module**

| | | | |
|---|---|---|---|
| | ⊣ Ind | p53 | *p53* represses *Cyclin D1* transcription by facilitating the recruitment of *HDAC1* to its promoter and thus promoting its deacetylation [230]. |
| | ⊣ IBind | p21_H | *p21*$^{Cip1}$ is a Cyclin Dependent kinase inhibitor which binds to and blocks the activity of *Cdk2*, *Cdk3*, *Cdk4* and *Cdk6* kinases [219] and thus inhibits *CyclinD1/Cdk4,6* [220]. |
| E2F1 | **E2F1** = (**Myc** or **E2F1**) and not(**pRB** or **CyclinA** or **CAD**) | | |
| | TF | In the absence of both *CyclinA* and *pRB*, *E2F1* transcription can be induced by *Myc* or maintained by active *E2F1*. *CAD* deactivates *E2F1* as it destroys the cell's DNA. | |
| | ← TR | Myc | *Myc* is required for growth-factor mediated induction of *E2F1* [231, 225]. It binds to and remodels the *E2F1* promoter, facilitating *E2F1* transcription [216]. In addition, *Myc* augments protein expression of *E2F1* [232]. Single-cell experiments show that *Myc* is a critical modulator of the amplitude of *E2F* activation [233]. |
| | ← TR | E2F1 | *E2F1* binds to its own promoter and up regulates transcription (as long as *Cyclin D/E* activity blocks *RB-E2F1* binding) [234]. |
| | ⊣ TR | pRB | *RB* binds to *E2F/DP1* complexes and switches their DNA binding activity from activation to repression [221, 235]. |
| | ⊣ P | CyclinA | The phosphorylation of the *E2F1*-binding *DP-1* protein by *Cyclin A*, which binds directly to *E2F-1* (as well as *E2F-2,3*) downregulates *E2F1* transcriptional activity in S phase [236, 237, 238]. |
| | ⊣ Deg | CAD | This link from Caspase-activated DNase (*CAD*) to *E2F1* ensures that apoptotic cells settle into an *E2F1*-negative attractor regardless of their initial state. The rationale for this is that *E2F1* cannot maintain its activity if DNA is fragmented. |
| CyclinE | **CyclinE** = (**ATP_H** or **CyclinE**) and **ATP** and **E2F1** and **Pre_RC** and **Cdc6** and not(**p21_H** or **pRB** or **p27Kip1** or **CHK1** or **CHK2** or **Casp3**) | | |
| | PC | In our model, the ON state of *Cyclin E* represents active *Cyclin E/Cdk2* complexes. Thus, its full activation requires high levels of ATP, transcription via *E2F1* not blocked by active *pRB*, binding to *Cdc6* and *pre-RC* complexes, and the absence of its inhibitors *p27*$^{Kip1}$, *CHK1*, *CHK2*, and *Caspase 3*. | |
| | ← Ind | ATP_H | The mechanistic link between normal-to high ATP production by hyperfused mitochondria is unclear, and may indeed be a largely indirect effect via AMPK/p53 activation [120]. That said, there is evidence that low ATP levels turn the balance between *p27*$^{Kip1}$ and *Cdk2/Cyclin E* in *p27*$^{Kip1}$'s favor as a CDKI, rather than a *Cdk2* substrate [239]. Thus, we assume that the hyperfused network only allows active *Cyclin E* buildup if mitochondrial ATP production is high; an assumpiton supported by observations that forced hyperfusion without increased ATP prodiction did not force *Cyclin E* levels to remain elevated after S-phase completion [120]. |

**Table S1h: Restriction_SW module**

| | | | |
|---|---|---|---|
| | ← Ind | ATP | As high ATP levels are required for *Cyclin E* buildup, we assume that at least nominal ATP levels remain required to maintain *Cyclin E* activity [120]. |
| | ← Per | CyclinE | We assume that once *Cyclin E* is activated with the aid of high ATP levels, it can remain active to continue S-phase progression without ongoing high ATP output from mitochondria. |
| | ← TR | E2F1 | *E2F1* is a potent transcriptional activator of *Cyclin E* [240]. |
| | ← Compl | Pre_RC | At the G1/S transition, *Cyclin E* is loaded onto chromatin by *pre-RC* complexes (*Cdc6* and *Cdt1* binding), where it is required for *MCM2* loading, origin firing and the start of DNA synthesis [241]. In addition, activation of its partner *Cdk2* by *Cdc6* is contingent on this localization [242]. |
| | ← Compl | Cdc6 | Chromatin association and full activation of *Cyclin E/Cdk2* requires *Cdc6* [242]. |
| | ⊢ IBind | p21_H | Above-basal $p21^{\mathrm{CIP1}}$ is a potent inhibitor of *Cyclin E/Cdk2* activity [243, 186]. |
| | ⊢ TR | pRB | *Cyclin E* transcription by *E2F1* requires the absence of active, un-phosphorylated *RB* [238]. |
| | ⊢ IBind | p27Kip1 | $p27^{Kip1}$ binds to and prevents the activation of *Cyclin E/Cdk2* complexes [190]. |
| | ⊢ P | CHK1 | *Chk1* activation during normal S-phase progression keeps *Cdk2* activity in a physiological range by binding to both *Cdk2* and *Cdc25A*, aiding the loss of *Cyclin E/Cdk2* activity [244]. |
| | ⊢ Ind | CHK2 | *CHK2* phosphorylates *Cdc25A* on serine 123, facilitating its destruction [245] and subsequently blocking *Cyclin E/Cdk2* activity [245]. |
| | ⊢ Lysis | Casp3 | *Caspase 3* cleaves and deactivates *Cyclin E*, which is then rapidly degraded [246]. |

**Table S1i: Origin_Licensing module**

| Target Node | Node Gate<br>Node Type<br>Link Type | Node Description<br>Input Node | Link Description |
|---|---|---|---|
| ORC | **ORC** = **E2F1** or (**Pre_RC** and **Cdt1** and **Cdc6**) | | |
| | PC | *ORC* proteins can bind at origins of replication when transcribed by *E2F1* or as part of a fully assembled and licensed *Pre-RC* complex (including active *Cdc6* and *Cdt1*). | |
| | ← TR | E2F1 | Expression of the *ORC1* gene is regulated by *E2F1* [247]. |

**Table S1i: Origin_Licensing module**

| | | | |
|---|---|---|---|
| | ← Compl | Pre_RC | Licensed but not yet fired replication complexes (*Pre-RC*s containing *ORC*, *Cdc6*, *Cdt1* and inactive *MCM*s) remain stable at sites of replication origin until fired by the activation of the *MCM* helicase [248]. |
| | ← Compl | Cdc6 | Availability of stable (unphosphorylated) *Cdc6* in the *Pre-RC* is necessary for the maintenance of licensed origins [248]. |
| | ← Compl | Cdt1 | Active (unphosphorylated and not geminin-bound) *Cdt1* bound to the *Pre-RC* is necessary for the maintenance of licensed origins [248]. |
| Cdc6 | **Cdc6** = not(**f4N_DNA** and **CyclinA**) and (((**E2F1** and **ORC**) and not **Plk1** and not **CyclinA**) or (**Pre_RC** and **ORC** and **Cdc6** and **Cdt1**)) and not **Casp3** | | |
| | Prot | | In our model the *Cdc6* node represents nuclear, chromatin-bound *Cdc6*. Thus, the node is only active during the assembly of pre-replication complexes, or their ongoing presence during DNA replication. *Cdc6* is ON in the absence of *Caspase 3* or *CyclinA / Cdk2* phosphorylation of *Cdc6* in all origins required for the completion of DNA replication (thus, its inhibition by *Cyclin A* also requires 4N DNA). In addition, active *Cdc6* requires either transcription by *E2F1* and recruitment by origin-bound *ORC* proteins in the absence of mitotic *Plk1* or maintenance of *Pre-RCs* by the presence of all of its components. |
| | ⊣ Ind | f4N_DNA | In our model, full deactivation of Cdc6 represents the firing of all ORCs as DNA replication is completed. Thus *Cyclin A*'s inhibitory action takes full effect once the cell reaches 4N DNA content [249]. |
| | ⊣ P | CyclinA | Phosphorylation of *CDC6* by *Cyclin A/Cdk2* during DNA replication leads to its re-localization to the cytoplasm [249]. |
| | ← TR | E2F1 | Transcription of Cdc6 is directly induced by E2F1 [250]. |
| | ← Compl | ORC | *ORC* recruits *Cdc6* to origins of replication [248]. |
| | ⊣ P | Plk1 | *Plk1* binds, phosphorylated and strongly recruits Cdc6 to the spindle pole during metaphase, then to the central spindle in anaphase, leading to its exclusion from chromosomes until telophase, when the majority of *Plk1* is degraded by $APC/C^{Cdh1}$ [251]. |
| | ← Compl | Pre_RC | Licensed but not yet fired replication complexes (*Pre-RC*s containing *ORC*, *Cdc6*, *Cdt1* and inactive *MCM*s) remain stable and *Cdc6*-bound until fired by the activation of the *MCM* helicase [248]. |
| | ← Per | Cdc6 | Stable (unphosphorylated) *Cdc6* in the *Pre-RC* is necessary for the maintenance of licensed origins [248]. |
| | ← Compl | Cdt1 | Active (unphosphorylated and not geminin-bound) *Cdt1* bound to the *Pre-RC* is necessary for the maintenance of licensed origins [248]. |

**Table S1i: Origin_Licensing module**

| | | | |
|---|---|---|---|
| | $\vdash$ Lysis | Casp3 | *Caspase 3* cleaves and deactivates *Cdc6* [252]. |
| Cdt1 | **Cdt1** = not **geminin** and **ORC** and **Cdc6** and not(**CyclinE** and **CyclinA** and **Cdc25A**) and ((**Pre_RC** and (**E2F1** or **Myc**)) or (**E2F1** and (**Myc** or not **pRB**))) | | |
| | Prot | Replication-origin bound *Cdt1* requires the absence of *geminin*, the presence of origin-bound *ORC* and *Cdc6*, and the absence of sustained *Cdk2* activity responsible for the firing of all origins during DNA synthesis (modeled as simultaneous *Cyclin E*, *Cyclin A* and *Cdc25A* activity). Bound into a licensed pre-replication complex (*Pre-RC*), *Cdt1* remains stable as long as it is transcribed by *E2F1* [253] or *Myc* [254] (this guarantees that *Pre-RC* complexes cannot persist indefinitely in the absence of de novo transcription). Alternatively, it can be turned on by *E2F1*, aided by *Myc* or the absence of *RB*, and *FoxO3* in cells with 4N DNA. | |
| | $\vdash$ IBind | geminin | *Geminin* binds to *Cdt1* at pre-replication complexes, where it blocks *Cdt1* binding to DNA, sequestering it away from *Pre-RC*s [255]. |
| | $\leftarrow$ Compl | ORC | *ORC*-bound origin of replication sites are the point of pre-replication complex assembly, where *Cdt1* is recruited by *ORC*-bound *Cdc6* [248]. |
| | $\leftarrow$ Compl | Cdc6 | *ORC*-bound *Cdc6* recruits *Cdt1* to *pre-RC* complexes [248]. |
| | $\vdash$ P | CyclinE | Sustained *Cdk2* activity during S-phase (modeled as simultaneous *Cyclin E*, *Cyclin A* and *Cdc25A* activity) is responsible for the firing of all origins required to complete DNA synthesis; it also leads to the phosphorylation and proteasomal degradation of *Cdt1* [248]. |
| | $\vdash$ P | CyclinA | Sustained *Cdk2* activity leads to phosphorylation and degradation of *Cdt1* [248]. |
| | $\vdash$ P | Cdc25A | Sustained *Cdk2* activity leads to phosphorylation and degradation of *Cdt1* [248]. |
| | $\leftarrow$ Compl | Pre_RC | Licensed but not yet fired replication complexes (*Pre-RC*) remain stable until fired during DNA replication [248]. |
| | $\leftarrow$ TR | Myc | *Cdt1* is a direct transcriptional target of the *Myc-Max* complex [254], ensuring its availability for *Pre-RC* formation and maintenance. |
| | $\leftarrow$ TR | E2F1 | *Cdt1* is a direct transcriptional target of *E2F1* [253], ensuring its availability for *Pre-RC* formation and maintenance. |
| | $\vdash$ TR | pRB | *E2F1*-mediated transcription of *Cdt1* is blocked by hypophosphorylated (active) *pRB* [253]. |
| Pre_RC | **Pre_RC** = **ORC** and **Cdc6** and **Cdt1** and not(**Replication** and **f4N_DNA**) | | |
| | PC | *Pre-RC* complexes assemble when *ORC*, *Cdc6*, and *Cdt1* are all bound to sites of replication origin along the DNA. The node denoting their licensing turns OFF at the moment of transition from ongoing *Replication* to *f4N_DNA* (it is blocked in the one time-point when both of these nodes are ON). | |

**Table S1i: Origin_Licensing module**

| | | | |
|---|---|---|---|
| | ← Compl | ORC | *Pre-RC* complexes assemble when *ORC*, *Cdc6*, and *Cdt1* are all bound to sites of replication origin along the DNA [248]. |
| | ← Compl | Cdc6 | *Pre-RC* complexes assemble when *ORC*, *Cdc6*, and *Cdt1* are all bound to sites of replication origin along the DNA [248]. |
| | ← Compl | Cdt1 | *Pre-RC* complexes assemble when *ORC*, *Cdc6*, and *Cdt1* are all bound to sites of replication origin along the DNA, leading to the recruitment of the *MCM* helicase [248]. |
| | ⊢ Unbind | Replication | *Pre-RC*s fire and fall apart during DNA replication [248]. |
| | ⊢ Ind | f4N_DNA | In our model the *Pre-RC* node turns OFF when *Replication* is completed, marked by the time-point when both *Replication* and *f4N_DNA* are ON. |
| geminin | **geminin** = **E2F1** and not(**pAPC** and **Cdc20**) and not **Cdh1** | | |
| | Prot | *Geminin* is present when transcribed by *E2F1* and not targeted for degradation by $APC/C^{Cdh1}$ or $APC/C^{Cdc20}$ [256]. | |
| | ← TR | E2F1 | *Geminin* is a direct transcriptional target of *E2F1* [253]. |
| | ⊢ Ubiq | pAPC | *Geminin* is a target of $APC/C^{Cdc20}$ at the metaphase/anaphase transition [256]. |
| | ⊢ Ubiq | Cdc20 | *Geminin* is a target of $APC/C^{Cdc20}$ at the metaphase/anaphase transition [256]. |
| | ⊢ Ubiq | Cdh1 | *Geminin* is a target of $APC/C^{Cdh1}$ ubiquitin ligase [257]. |

**Table S1j: Phase_SW module**

| Target Node | Node Gate<br>Node Type<br>Link Type | Node Description<br>Input Node | Link Description |
|---|---|---|---|
| CyclinA_mRNA | **CyclinA_mRNA** = ((**E2F1** and (not **pRB** or **Myc**)) or **FoxM1**) and not(**p53_4** or **CAD**) | | |
| | mRNA | In non-apoptotic cells (no *CAD*), *Cyclin A* is transcribed by *E2F1* in the absence of active *RB* or by *FoxM1*. In addition, high and/or sustained *p53* activity, represented in our model by the *p53_4* node, can also repress *Cyclin A* transcription [258, 259]. | |
| | ← TR | E2F1 | *Cyclin A* is transcriptionally activated by *E2F* factors [238]. |
| | ⊢ TR | pRB | Active *RB* blocks *E2F1*'s ability to transcribe *Cyclin A* [260]. |
| | ← TR | Myc | *Cyclin A* is transcriptionally activated in *Myc* overexpressing cells [261]. |

**Table S1j: Phase_SW module**

| | | | |
|---|---|---|---|
| | ← TR | FoxM1 | Depletion of *FoxM1* results in reduced *Cyclin A2* expression (it is not clear whether *FoxM1* is a direct transcriptional inducer of *Cyclin A*) [262, 263, 264, 265]. |
| | ⊢ TR | p53_4 | *p53* overexpression can repress *Cyclin A2*[259], an effect that likely requires high or sustained *p53* levels so that cells able to recover from G2 arrest without excessive damage do not loose their G2 cyclin profile and can continue to mitosis [258]. |
| | ⊢ Ind | CAD | This link from Caspase-activated DNase (*CAD*) to *Cyclin A* mRNA ensures that apoptotic cells settle into a G0-like attractor regardless of their initial state. The rationale for this is that no mRNA synthesis can be maintained if DNA is fragmented (we only use these links from *CAD* if needed). |
| Emi1 | **Emi1** = (**E2F1** or not **pRB** or not **p21**) and not(**Plk1** and **CyclinB** and **Cdk1** and (**U_Kinetochores** or **A_Kinetochores**)) | | |
| | Prot | | Our model allows the sustained presence of *Emi1* protein when it is either actively transcribed by *E2F1* [266, 267] or lacks joint inhibition by *pRB* [267] and *p21* [268]. Degradation of *Emi1* is mediated by *Plk1* and *CyclinB/Cdk1* complexes; initiation of this degradation requires at least temporary co-localization of *Emi1* with *Plk1* at mitotic spindle poles [269]. |
| | ← TR | E2F1 | *Emi1* is a direct transactional target of *E2F1* [266, 267]. |
| | ⊢ TR | pRB | Active retinoblastoma protein can block *Emi1* transcription mediated by *E2F1* [267]. |
| | ⊢ Ind | p21 | *p21* activation during DNA damage lead to a substantial decrease of *Emi1* levels, not observed in *p21*-null cells [268]. |
| | ⊢ P | Plk1 | *Plk1* phosphorylates *Emi1* at mitotic spindle poles, stimulating its *βTrCP* binding and ubiquitination [269]. |
| | ⊢ Ind | CyclinB | *Cyclin B/Cdk1* enhances the ability of *Plk1* to mediate *Emi1* destruction [269]. |
| | ⊢ Ind | Cdk1 | *Cyclin B/Cdk1* enhances the ability of *Plk1* to mediate *Emi1* destruction [269]. |
| | ⊢ Ind | U _Kinetochores | As *Plk1*-mediated phosphorylation of *Emi1* occurs at mitotic spindle poles, our model requires ongoing mitosis for this interaction [269]. |
| | ⊢ Ind | A _Kinetochores | *Plk1* phosphorylates *Emi1* at mitotic spindle poles, stimulating its *βTrCP* binding and ubiquitination [269]. |
| FoxM1 | **FoxM1** = (**Myc** and **CyclinE**) or (**CyclinA** and **Cdc25A** and **Cdc25B**) or (**Plk1** and **CyclinB** and **Cdk1**) | | |
| | TF | | In our model, *FoxM1* activity requires increased expression by *Myc* [270] and activating phosphorylation by *Cyclin E/Cdk2*. Alternatively, *FoxM1* activity can be sustained by potent *Cdk2 / Cdk1* activity in G2 (supported by *Cdc25A* or *Cdc25B*), or a serial phosphorylation by *Cyclin B/Cdk1* and *Plk1* during mitosis. |

**Table S1j: Phase_SW module**

| | | | |
|---|---|---|---|
| | ← TR | Myc | *FoxM1* is a direct transcriptional target of *c-Myc* [270]. |
| | ← P | CyclinE | *Cyclin E/Cdk2* complexes bind and phosphorylate FoxM1, potently inducing its transcriptional activity, which starts during S-phase [271]. |
| | ← P | CyclinA | In addition to *Cyclin E/Cdk2*, *Cyclin A/Cdk2* complexes can also keep *FoxM1* transcriptionally active by phosphorylating its autoinhibitory N-terminal region [272]. |
| | ← Compl | Cdc25A | Active *Cdc25A* binds to and enhances the transcriptional activity of *FoxM1*, potentually by bridging FoxM1 and active cyclin-*Cdk2* complexes [273]. |
| | ← Ind | Cdc25B | *Cdc25B* overexpression can increase *FoxM1*-dependent transcription, likely via aiding *Cdk1* activity [274]. |
| | ← P | Plk1 | *Plk1* binds and phosphorylates *FoxM1*, which activates *FoxM1*-mediated transcription in early mitosis [275]. |
| | ← P | CyclinB | *FoxM1* binds *Plk1*, and phosphorylation of two key residues at this binding domain by *Cyclin B/Cdk1* primes it for *Plk1* binding [275]. |
| | ← P | Cdk1 | *FoxM1* binds *Plk1*, and phosphorylation of two key residues at this binding domain by *Cyclin B/Cdk1* primes it for *Plk1* binding [275]. |
| Cdc25A | **Cdc25A** = not **CHK2** and ((**FoxM1** and **E2F1** and not **pRB**) or (not **Cdh1** and (**FoxM1** or (**E2F1** and not **pRB**)))) and (not(**GSK3** or **CHK1**) or **CyclinE** or **CyclinA** or (**CyclinB** and **Cdk1**)) | | |
| | Ph | | As the precise combinatorial regulation of *Cdc25A* throughput the cell cycle is unknown, our model assumes that accumulation of the *Cdc25A* protein requires transcriptional activation by both *E2F1* in the absence of *pRB*, and *FoxM1* to override destruction by $APC/C^{Cdh1}$. Alternatively, one of the two transcription factors can drive *Cdc25A* accumulation in the absence of $APC/C^{Cdh1}$. In addition, stabilization of *Cdc25A* either requires the absence of *GSK3β* and *CHK1* (both of which promote its degradation), or stabilization by *Cdk* activity. |
| | ← TR | FoxM1 | *FoxM1* is a direct transcriptional inducer of *Cdc25A* [273]. |
| | ← TR | E2F1 | *E2F1* is a direct transcriptional inducer of *Cdc25A* [276]. |
| | ⊣ TR | pRB | Active (hypo-phosphorylated) *pRB* blocks *E2F1*'s ability to drive *Cdc25A* transcription [276, 277]. |
| | ⊣ Deg | Cdh1 | The $APC/C^{Cdh1}$ complex degrades *Cdc25A* at mitotic exit [278, 279]. |
| | ⊣ P | GSK3 | *GSK3β* phosphorylates *Cdc25A*, promoting its proteolysis [280]. |

**Table S1j: Phase_SW module**

| | | | |
|---|---|---|---|
| | ⊣ P | CHK1 | *CHK1* phosphorylates *Cdc25A*, promoting its proteolysis and inhibiting its interaction with *Cyclin A/Cdk1/2* and *Cyclin B/Cdk1* [281, 282]. |
| | ⊣ P | CHK2 | *CHK2* phosphorylates *Cdc25A*, promoting its proteolysis and blocking its ability to keep *Cyclin A/Cdk1/2* active [245]. |
| | ← P | CyclinE | *Cdc25A* protein levels are stabilized during S-phase by *CyclinE/Cdk2*-dependent phosphorylation [283]. |
| | ← P | CyclinA | *Cdc25A* protein levels are stabilized during S and G2 by *Cdk2*-dependent phosphorylation. *Cdk2* first partners with *Cyclin E* [283], then continues to stabilize *Cdc25A* past the point of *Cyclin E* expression by partnering with *Cyclin A* [284]. |
| | ← P | CyclinB | During mitosis, *Cdc25A* is stabilizes by *Cyclin B/Cdk1* phosphorylation, which protects it from the proteasome [285]. |
| | ← P | Cdk1 | During mitosis, *Cdc25A* is stabilizes by *Cyclin B/Cdk1* phosphorylation, which protects it from the proteasome [285]. |
| CyclinA | **CyclinA** = **CyclinA_mRNA** and not **pAPC** and ((**Cdc25A** and (not **Cdh1** or **Emi1**)) or (**CyclinA** and ((not **Cdh1** and (**Emi1** or not **UbcH10**)) or (**Emi1** and not **UbcH10**)))) | | |
| | PC | | *Cyclin A* activity requires transcription (*Cyclin A* mRNA) and the absence of degradation by phosphorylated (mitotic) *pAPC*. In addition, turning ON inactive *Cyclin A* requires activation of *Cdk2* by *Cdc25A* [286] and the absence / *Emi1*-mediated inhibition of $APC/C^{Cdh1}$. Once active, *Cyclin A* maintains its activity in the absence of overpowering influences driving its degradation. Namely, *Cyclin A* relies on either *Emi1* or the absence of *UbcH10* for its ability to keep inactive $APC/C^{Cdh1}$ in check. To overpower active $APC/C^{Cdh1}$, *Cyclin A* requires both *Emi1* and no *UbcH10*. The precise combinatorial regulation of *Cyclin A* is not known; the above logic is consistent with *Cyclin A* activity pattern during cell cycle progression. |
| | ← TL | CyclinA_mRNA | Sustained availability of *Cyclin A* requires ongoing translation from *CyclinA* mRNA. |
| | ⊣ Deg | pAPC | *Cyclin A* is degraded by the $APC/C^{Cdc20}$ in prometaphase (as soon as the APC/C components are phosphorylated by Cdk1) [287, 288], before the full activation of the complex at SAC passage [289]. In our model, this stage of mitotic $APC/C^{Cdc20}$ activation is represented by *Cdk1*-phosphorylated $APC/C$ ($pAPC$). |
| | ← DP | Cdc25A | *Cdc25A* promotes active *Cyclin A/Cdk2* complex formation by removing inhibitory phosphorylation of *Cdk2* [286, 290]. |
| | ⊣ Deg | Cdh1 | *Cyclin A* is degraded by $APC/C^{Cdh1}$ in the presence of the *UbcH10* protein [291, 292, 199]. |
| | ← Compl | Emi1 | *Emi1* binding to *Cdh1* is required to stabilize *Cyclin A* levels at the G1/S transition, allowing *Cyclin A/Cdk2* to block $APC/C^{Cdh1}$ [293, 294, 295]. |
| | ← Per | CyclinA | We assume that once activated, *Cyclin A/Cdk2,1* complexes can sustain their activity until *Cyclin A* is degraded. |

**Table S1j: Phase_SW module**

| | | | |
|---|---|---|---|
| | $\underset{\text{Deg}}{\vdash}$ | UbcH10 | *Cyclin A* degradation by $APC/C^{Cdh1}$ requires *UbcH10* [292]. |
| Wee1 | **Wee1** = (**Replication** or **CHK1**) and not (**Cdk1** and **CyclinB**) and (**CHK1** or not (**Cdk1** and **CyclinA** and **Plk1**)) and not **Casp3** | | |
| | K | | *Wee1* is active during *Replication*, unless its activity is blocked by *CyclinA/Cdk1* OR *CyclinB/Cdk1* [296]. |
| | $\underset{\text{ComplProc}}{\leftarrow}$ | Replication | To model the sensitivity of *Wee1* activation to ongoing DNA synthesis even in the absence of damage, our model turns on *Wee1* immediately upon the start of DNA replication and maintains it until both Replication and the checkpoint kinase *Chk1* is OFF [297]. In addition, *Wee1* activity has been implicated in maintaining normal replication fork procession, linking its activity directly to ongoing replication [298]. |
| | $\underset{\text{P}}{\leftarrow}$ | CHK1 | During DNA replication *Wee1* is activated by the checkpoint kinase *CHK1* [297]. |
| | $\underset{\text{P}}{\vdash}$ | CyclinB | *Cyclin B/Cdk1* is a strong inducer of *Wee1* phosphorylation and deactivation [296]. |
| | $\underset{\text{P}}{\vdash}$ | Cdk1 | The somatic *Wee1* protein is an order of magnitude more sensitive to *Cdk1* activity than *Cdc25C*. Thus, both *Cyclin A/Cdk1* and *Cyclin B/Cdk1* strongly induce *Wee1* phosphorylation and deactivation [296]. |
| | $\underset{\text{P}}{\vdash}$ | CyclinA | *Cyclin A/Cdk1* is a strong inducer of *Wee1* phosphorylation and deactivation [296]. |
| | $\underset{\text{P}}{\vdash}$ | Plk1 | *Plk1* phosphorylation at S53 promotes *Wee1* degradation [299]. This event is primed by *Cdk1* phosphorylation of *Wee1* at S123 [299]. As the main partner of *Cdk1* in mitosis is *Cyclin B*, we assume that assistance from *Plk1* to block *Wee1* is more relevant when paired with *Cyclin A/Cdk1* complexes. |
| | $\underset{\text{Lysis}}{\vdash}$ | Casp3 | *Caspase 3* cleaves and deactivates *Wee1* [300]. |
| UbcH10 | **UbcH10** = not **Cdh1** or (**UbcH10** and (**Cdc20** or **CyclinA** or **CyclinB**)) | | |
| | UbL | | The ubiquitin-conjugating enzyme (E2) *UbcH10* is active in the absence of *Cdh1*. Alternatively, active *UbcH10* is maintained in the presence of *Cdh1* when some of its targets are present: *Cdc20* OR *CyclinA* OR *CyclinB* [292]. |
| | $\underset{\text{Deg}}{\vdash}$ | Cdh1 | *UbcH10* is degraded by $APC/C^{Cdh1}$. |
| | $\underset{\text{Per}}{\leftarrow}$ | UbcH10 | Active *UbcH10* cannot be autoubiquitinated in the presence of $APC/C^{Cdh1}$ substrates and thus remains active [292]. |
| | $\underset{\text{PBind}}{\leftarrow}$ | Cdc20 | The presence of $APC/C^{Cdh1}$ substrates, including *Cdc20*, inhibit the autoubiquitination of *UbcH10* but not its function, thus preserving APC activity [292]. |
| | $\underset{\text{PBind}}{\leftarrow}$ | CyclinA | The presence of $APC/C^{Cdh1}$ substrates, including *Cyclin A*, inhibit the autoubiquitination of *UbcH10* but not its function, thus preserving APC activity [292]. |

**Table S1j: Phase_SW module**

| | | | |
|---|---|---|---|
| | ← PBind | CyclinB | The presence of $APC/C^{Cdh1}$ substrates, including *CyclinB*, inhibit the autoubiquitination of *UbcH10* but not its function, thus preserving APC activity [292]. |
| CyclinB | **CyclinB** = (**FoxM1** or (**FoxO3** and **CyclinB**)) and not(**Cdh1** or (**pAPC** and **Cdc20**)) and not **p53_4** | | |
| | PC | *Cyclin B* node is ON when the concentration of *Cyclin B* proteins is high (does not represent the activity of *CyclinB/Cdk1* complexes). This occurs when *Cyclin B* is transcribed by *FoxM1*, maintained by *FoxO3* transcription, and not undergoing *APC*-mediated degradation by $APC/C^{Cdc20}$ or $APC/C^{Cdh1}$. In addition, high and/or sustained *p53* activity, represented in our model by the *p53_4* node, can also repress *Cyclin B* transcription [258, 301]. | |
| | ← TR | FoxO3 | *FoxO3* is a direct transcriptional regulator of *Cyclin B*; its activation in G2 helps increase/maintain *Cyclin B* levels [302]. |
| | ← TR | FoxM1 | *FoxM1* is a direct transcriptional regulator of *Cyclin B1* [265, 303]. |
| | ← Per | CyclinB | Here we assume that FoxO3 alone can only maintain, but not independently induce *Cyclin B1* expression. |
| | ⊢ Deg | pAPC | *Cyclin B* is degraded by $APC/C^{Cdc20}$ [291]. |
| | ⊢ Deg | Cdc20 | *Cyclin B* is degraded by $APC/C^{Cdc20}$ [291]. |
| | ⊢ Deg | Cdh1 | *Cyclin B* is degraded by $APC/C^{Cdh1}$ [291]. |
| | ⊢ TR | p53_4 | *p53* can repress *Cyclin B* transcription [301], an effect that likely requires high or sustained *p53* levels so that cells able to recover from G2 arrest without excessive damage do not loose their G2 cyclin profile and can continue to mitosis [258]. |
| Cdc25B | **Cdc25B** = **FoxM1** and **f4N_DNA** | | |
| | Ph | *Cdc25B* activation requires transcription by *FoxM1*, centrosomal localization, and activation by *Aurora A* kinase on replicated centrosomes. | |
| | ← TR | FoxM1 | *FoxM1* is an essential inducer or *Cdc25B* [304]. |
| | ← Loc | f4N_DNA | *Cdc25B* is localized at centrosomes, where it is activated by *Aurora A* kinase [305]. As *Aurora A* itself is only recruited to duplicated, centrosomes before their separation [306], *Cdc25B* activation requires duplicated centrosomes. As our model does not directly account for centrosome dynamics, we account for this by requiring the completion of S-phase (4N DNA). |
| Plk1 | **Plk1** = not **Cdh1** and (**FoxM1** or **Plk1_H**) and ((**CyclinB** and **Cdk1**) or (**CyclinA** and **Cdc25A** and not **Wee1**)) | | |

**Table S1j: Phase_SW module**

| | | | |
|---|---|---|---|
| | K | *Plk1* activity requires the absence of $APC/C^{Cdh1}$, transcription by *FoxM1*, or high *Plk1* levels transcribed earlier by both *FoxM1* and *FoxO3* (see *Plk1_H* below) [302]. In addition, *Plk1* activation requires phosphorylation by either *CyclinB/Cdk1* during mitosis or *Cyclin A/Cdk2* (aided by lack of *Wee1* and *Cdc25A*) at the G2/M boundary [307]. | |
| | ⊣ Ubiq | Cdh1 | The majority of *Plk1* is degraded in anaphase by the $APC/C^{Cdh1}$ complex [308]. |
| | ← TR | FoxM1 | *Plk1* is a direct transcriptional target of *FoxM1* [275]. |
| | ← Per | Plk1_H | Our model tracks the accumulauton of high-enough levels of *Plk1* to survive $APC/C^{Cdh1}$ mediated destruction into telophase via the *Plk1_H* node. Its ON state represents strong prior *Plk1* activation. Thus, it sustains the *Plk1* node in the absence of *FoxM1*-mediated transcription until *Plk1_H* itself is lost as *Plk1* levels fall. |
| | ← P | CyclinB | *Plk1* is activated by *Cyclin B/Cdk1* phosphorylation [309, 310, 311]. |
| | ← P | Cdk1 | *Plk1* is activated by *Cyclin B/Cdk1* phosphorylation [309, 310, 311]. |
| | ← P | CyclinA | *Plk1* activation at the G2/M boundary, before *Cdk1/Cuclin B* complexes are activated, requires active *Cyclin A/Cdk* [307]. |
| | ← Ind | Cdc25A | As we do not include a separate *Cdk2* node in our model, strong *Cyclin A/Cdk2* activity requires ongoing dephosphorylation of *Cdk2* by *Cdc25A* [312]. |
| | ⊣ Ind | Wee1 | *Cyclin A*-mediated induction of *Plk1* is blocked by *Wee1* kinase, which specifically inhibits *Cdk2* activity [307]. |
| Cdc25C | **Cdc25C** = **f4N_DNA** and **Plk1** and ((**Cdc25B** and not **CHK1** and (not **CHK2** or **Bipolar_Spindle**)) or (**CyclinB** and **Cdk1**)) | | |
| | Ph | In our model, *Cdc25C* is active in cells with replicated DNA (see *f4N_DNA* → *Cdc25C* link). Its activation is initiated by a small, initially cytoplasmic pool of *Cyclin B/Cdk1* activated by *Cdc25B* (not directly represented in our model) and further increased by *Cdc25B* itself, which translocates to the nucleus with the aid of *Plk1*. During mitosis, *Plk1* potentiates the ability of *Cyclin B/Cdk1* to maintain *Cdc25C* activity. In contrast, *CHK1/2* phosphorylate *Cdc25C*, leading to its nuclear exclusion and degradation [313, 314, 315, 316]. | |
| | ← Ind | f4N_DNA | The nature and localization of the signals responsible for the onset and maintenance of *Cdc25C* activity require replicated DNA (*f4N_DNA*) [317, 318]. Namely, *Cdc25C* is initially activated by a small pool of *Cyclin B/Cdk1* (below the ON-threshold of *Cdk1* in our model) which starts our at the replicated centrosome. Moreover, the pool of mitotic *Cdc25C* co-localized with active *Chk1/Cyclin B* is found on condensed chromosomes, again requiring the presence of *f4N_DNA* [319]. |

**Table S1j: Phase_SW module**

| | | | |
|---|---|---|---|
| | ← P | Plk1 | In addition, *Plk1* induces nuclear transport of *CDC25B*, where it contributes to the initiation of *Cdk1* activity [317]. During mitosis, *Plk1* helps maintain strong *Cdc25C* activation by phosphorylating it on the same site as *Cyclin B/Cdk1* [320], as indicated by the profound decrease of Cdc25C activity in Plk1-inhibited mitotic cells [310, 321]. |
| | ← Ind | Cdc25B | *CDC25B* starts the cascade leading to mitotic entry by activating a small centrosomal pool of *Cyclin B/Cdk1*, leading to their nuclear translocation where they trigger the activation of *Cdc25C* and eventually the larger nuclear *Cyclin B/Cdk1* pool [322, 323, 318]. |
| | ⊣ P | CHK1 | *CHK1* phosphorylates *Cdc25C*, leading to its nuclear exclusion, loss of access to its main target, *Cdk1* [319]. In addition, *CHK1* blocks the ability of *Cdc25B* to activate *Cdc25C* at the centrosomes by phosphorylating it and blocking its *Cdk1* activity [315, 316]. |
| | ⊣ P | CHK2 | *CHK2* phosphorylates *Cdc25C*, leading to its nuclear exclusion and degradation [313, 314]. |
| | ← Ind | Bipolar _Spindle | During mitosis, *CHK2* is localized to the centrosomes where it helps maintain SAC (not modeled). Thus, we assume that it cannot repress *Cdc25C* once a bipolar spindle has been assembled [324]. |
| | ← P | CyclinB | *Cyclin B/Cdk1* complexes are potent activators of *Cdc25C*, creating positive feedback that causes switch-like mitotic entry [325, 326]. |
| | ← P | Cdk1 | *Cyclin B/Cdk1* complexes are potent activators of *Cdc25C*, creating positive feedback that causes switch-like mitotic entry [325, 326]. |
| Cdk1 | **Cdk1** = (**CyclinB** and **Cdc25C**) and (not(**CHK1** or **CHK2**) or (not **Wee1** and **Cdk1**)) | | |
| | K | | Full *Cdk1* kinase activation requires its binding partner *Cyclin B* and the *Cdc25C* phosphatase, which maintains *Cdk1* in an active dephosphorylated state. *Cdk1* is inhibited by the checkpoint kinases *CHK1/2*, unless already full active and *Wee1* kinase is inhibited. |
| | ← Compl | CyclinB | Full kinase activation of *Cdk1* in our model requires it to complex with *Cyclin B* [327]. |
| | ← DP | Cdc25C | *Cdk1* is subject to inhibitory phosphorylation by *Wee1* or *Myt1*, and its dephosphorylation is carried out by activated *Cdc25C* [325, 327, 326]. |
| | ⊣ P | CHK1 | In the absence of *CHK1* kinase, a small cytosolic (centrosomal) pool of *Cyclin B/Cdk1* can be activated by *Cdc25B*, the nuclear translocation of which can trigger a positive feedback loop that activates the full *Cdk1* pool (assuming nuclear *Wee1* is also inactive). Thus, *CHK1* can maintain the OFF state of inactive *Cdk1* [307]. |

**Table S1j: Phase_SW module**

| | | | |
|---|---|---|---|
| | ⊣ P | CHK2 | The absence of *CHK2* activity is required to keep *Cyclin B/Cdk1* activate via *Cdc25B/C*-mediated dephosphorylation [328]. |
| | ⊣ P | Wee1 | *Wee1* is a nuclear protein that ensures the completion of DNA replication prior to mitosis by blocking nuclear *Cdk1* activation [329]. |
| | ← Per | Cdk1 | We assume that the presence of fully activated, nuclear *Cdk1* is able to overcome the effect of active *Wee1*, given that *Wee1* is very sensitive to *Cdk1*-mediated inhibitory phosphorylation [296]. |
| pAPC | **pAPC** = (**CyclinB** and **Cdk1** and **Plk1**) or (**CyclinB** and **Cdk1** and **pAPC**) or (**pAPC** and **Cdc20**) | | |
| | PC | In line with evidence that *Plk1* can aid full activation of *APC/C*, but *Cdk1* appears to be the more potent inducer, our model requires both *Cyclin B/Cdk1* and *Plk1* to activate *APC/C* from an OFF state, but only *Cdk1* activity to maintain it. In addition, ongoing phosphorylation of the functional $APC/C^{Cdc20}$ complex is no longer required. | |
| | ← P | CyclinB | *CyclinB/Cdk1* activation triggers mitotic entry and promotes $APC/C^{Cdc20}$ activity via APC/C subunit phosphorylation [330, 331]. |
| | ← P | Cdk1 | *CyclinB/Cdk1* activation triggers mitotic entry and promotes $APC/C^{Cdc20}$ activity via APC/C subunit phosphorylation [330, 331]. |
| | ← P | Plk1 | In addition to *Cyclin B/Cdk1* phosphorylation, full activation of the $APC/C^{Cdc20}$ complex also requires the kinase activity of *Plk1* [332]. |
| | ← Per | pAPC | Activated $APC/C^{Cdc20}$ initiates the Metaphase / Anaphase transition by degrading *Cyclin B* and securin [256]. Once active, $APC/C^{Cdc20}$ no longer requires sustained *CyclinB/Cdk1* or *Plk1* phosphorylation. |
| | ← Compl | Cdc20 | Once active, $APC/C^{Cdc20}$ no longer requires sustained *CyclinB/Cdk1* or *Plk1* phosphorylation. |
| Cdc20 | **Cdc20** = **pAPC** and not **Emi1** and not **Cdh1** and ((**Bipolar_Spindle** and not(**Mad2** and **Mps1**)) or (not **CyclinA** and not(**CyclinB** and **Cdk1**))) | | |
| | Prot | In our model, $APC/C^{Cdc20}$ complex formation is represented by the joint activity of *Cdc20* and phosphorylated *APC/C* (*pAPC*). *Cdc20* is thus ON in the presence of *pAPC* when both *Emi1* and *Cdh1* are absent ($APC/C^{Cdh1}$ is represented by the *Cdh1* node, see below). In addition, *Cdc20* activity requires a bipolar spindle, and either the absence of *Mps1* and *Mad2* at unattached kinetochores, or the absence of *Cdc20* phosphorylation by *Cyclin B/Cdk1* or by *Cyclin A/Cdk2* complexes to potentiate the interaction between *Mad2* and *Cdc20*, and *pAPC* (present and phosphorylated) [333]. | |
| | ← Compl | pAPC | *Cdc20* becomes active in early mitosis by binding to *APC/C*, an event that requires *Cyclin B/Cdk1*-mediated phosphorylation of several core *APC/C* subunits [279, 334]. |

**Table S1j: Phase_SW module**

| | | | |
|---|---|---|---|
| | ⊣ IBind | Emi1 | *Emi1* binds *Cdc20* and inhibits the ubiquitin ligase activity of $APC/C^{Cdc20}$ [294]. |
| | ⊣ Deg | Cdh1 | $APC/C^{Cdh1}$ complexes degrade *Cdc20*, leading to a complete switch from $APC/C^{Cdc20}$ to $APC/C^{Cdh1}$ during mitotic exit [279, 335]. |
| | ← Ind | Bipolar _Spindle | Completion of a bipolar spindle in which every chromosome is pulled in opposite directions by microtubule-attached kinetochodes is required for SAC passage and $APC/C^{Cdc20}$ activation [336]. |
| | ⊣ IBind | Mad2 | Eukaryotic cells do not separate their replicated genome until they pass the Spindle Assembly Checkpoint (SAC). Namely, all their chromosomes need to be are aligned with respect to the metaphase plane and the two copies of each chromosome need to be attached to opposite poles of the mitotic spindle [331]. This physical alignment is monitored via *Mad2*: kinetochores that remain unattached to microtubules catalyze the sequestration of *Cdc20* and thus inhibit $APC/C^{Cdc20}$ [337, 336]. |
| | ⊣ IBind | Mps1 | *Mps1* inhibits cytosolic *Cdc20*, as it aids *Cdc20* binding to *Mad2* and *BubR1* before kinetochores mature. Then it recruits factors that sustain the SAC to unattached kinetochores (*Mps1* inhibition in prometaphase removes all known SAC mediators from kinetochores) [338]. |
| | ⊣ P | CyclinA | *Cyclin A/Cdk2* complexes phosphorylate *Cdc20* and inactivate the $APC/C^{Cdc20}$ complex during S and G2 [339]. |
| | ⊣ P | CyclinB | *Cyclin B* partners with *Cdk1* to keep *Cdc20* phosphorylated, increasing its interaction with *Mad2* rather than *APC/C* [340]. |
| | ⊣ P | Cdk1 | *Cdk1*-phosphorylated *Cdc20* interacts with *Mad2* rather than *APC/C*, resulting in a block on $APC/C^{Cdc20}$ activation until completion of spindle assembly [333]. |
| Cdh1 | **Cdh1** = not(**CyclinB** and **Cdk1**) and not(**CyclinA** and (**Emi1** or **Cdc25A**)) | | |
| | PC | | $APC/C^{Cdh1}$ activity requires the absence of Cyclin Dependent kinase phosphorylation by *Cyclin B/Cdk1*, or *Cyclin A/Cdk2* aided by further inhibition of *Cdh1* by *Emi1*, or ongoing *Cdk2* activation by *Cdc25A* in the absence of *Emi1*. |
| | ⊣ P | CyclinB | *Cyclin B/Cdk1* phosphorylates *Cdh1* during mitosis, impairing its interaction with *APC/C* [279, 291]. |
| | ⊣ P | Cdk1 | *Cyclin B/Cdk1* phosphorylates *Cdh1* during mitosis, impairing its interaction with *APC/C* [279, 291]. |
| | ⊣ P | CyclinA | Active *Cyclin A/Cdk1,2* complexes phosphorylate *Cdh1* during S, G2 and early mitosis, impairing its interaction with *APC/C* until late stages of mitosis when *Cdk1/2* activity falls [279, 291]. |

**Table S1j: Phase_SW module**

| | | | |
|---|---|---|---|
| | ⊣ IBind | Emi1 | *Emi1* blocks $APC/C^{Cdh1}$ binding to its substrates [295], as well as its ability to add ubiquitin chains to them [341]. |
| | ⊣ Ind | Cdc25A | As we do not include a separate *Cdk2* node in our model, strong *Cyclin A/Cdk2* activity capable of overriding *Cdh1* activity even in the presence of *Emi1* requires ongoing dephosphorylation of *Cdk2* by *Cdc25A* [325]. |

**Table S1k: Cell_Cycle_Process module**

| Target Node | Node Gate<br>Node Type<br>Link Type | <br>Node Description<br>Input Node | <br><br>Link Description |
|---|---|---|---|
| Replication | **Replication** = **Pre_RC** and ((**E2F1** and **CyclinE** and **Cdc25A**) or (**Replication** and **CyclinA** and **Cdc25A** and (**E2F1** or not **f4N_DNA**))) and not **CAD** | | |
| | Proc | The *Replication* node represents ongoing DNA synthesis. This requires a non-apoptotic cell, licensed pre-replication complexes (*Pre_RC*). The start of DNA synthesis requires *E2F1*-meditated transcription of the genes that help execute it, as well as the firing of the first round of replication origins by *Cyclin E/Cdk2*. Once ongoing, replication is sustained by *Cyclin A/Cdk2* and *Cdc25A*, aided by *E2F1* and terminated by completion of a full round of synthesis (*4N_DNA*). | |
| | ← ComplProc | Pre_RC | Ongoing DNA replication requires licensed replication origins, which fire throughout DNA synthesis [342]. |
| | ← ComplProc | E2F1 | In addition to *E2F1* target genes directly included in our model, *E2F1* transcribes an array of critical S-phase genes responsible for carrying out DNA synthesis (e.g, *POLA1*, *POLA2*, *MCM3*, *MCM5*, *MCM6*, *PCNA*, *TOP2A*, *RFC2*, *TK1*) [343, 344]. |
| | ← ComplProc | CyclinE | DNA replication is initiated by fully active *Cyclin E/Cdk2* [345]. |
| | ← ComplProc | Cdc25A | Active *Cdc25A* is required for onset as well as progression through S-phase [346, 347]. |
| | ← Per | Replication | Once ongoing, DNA synthesis continues in the presence of active *Cyclin A/Cdk2*, only ending when DNA content is doubled. |
| | ← ComplProc | CyclinA | *Cyclin A/Cdk1* complexes regulate the origin firing program in mammalian cells and are required for the completion of DNA replication [347, 290]. |
| | ⊣ ComplProc | f4N_DNA | Complete duplication of a cell's DNA, represented in our model by *f4N_DNA* = ON, marks the end of active *Replication*. |
| | ⊣ ComplProc | CAD | Caspase-activated DNase (*CAD*) destroys DNA, preventing ongoing replication. |

##### Table S1k: Cell_Cycle_Process module

| | | | |
|---|---|---|---|
| ATR | **ATR** = **Replication** or **cROS_H** | | |
| | K | | *ATR* accumulates at replication forks during unperturbed DNA synthesis [348], as well as at sites of single-strand DNA damage induced by oxidative stress [349, 350]. |
| | ← Loc | Replication | *ATR* accumulates at replication forks during unperturbed DNA synthesis [348]. |
| | ← Ind | cROS_H | Elevated cytosolic ROS activates the *ATR/CHK1* pathway (likely by sustaining DNA damage) [350]. |
| CHK1 | **CHK1** = **ATR** and **Glucose** | | |
| | K | | *ATR* kinase activates *CHK1* at replication forks, which not only blocks premature mitosis but also regulates the rate of origin firing by keeping *Cdc25* protein levels from increasing above their physiological range [348]. |
| | ← P | ATR | *ATR* kinase activates *CHK1* at replication forks (by phosphorylation of serines 317 and 345), which not only blocks premature mitosis but also regulates the rate of origin firing by keeping *Cdc25* protein levels from increasing above their physiological range [351, 348]. |
| | ← Ind | Glucose | Glucose deprivation induces ubiquitin-mediated *CHK1* degradation [352]. |
| Nuclear _Membrane | **Nuclear_Membrane** = not(**CyclinB** and **Cdk1**) and not(**U_Kinetochores** or **A_Kinetochores**) | | |
| | MSt | | The nuclear membrane node represents a non-mitotic state where the cell's DNA is enclosed by the nuclear membrabe. Thus, this node is ON in the absence of *CyclinB/Cdk1*-mediated phosphorylations of the N-terminal head region of nuclear lamins, which form the meshwork structure near the inner nuclear membrane. This phosphorylations depolymerizes the lamin filament network and disolves the envelope [353]. Once disolved, the nuclear membrane does not form until the mitotic spindle is pulled apart in anaphase, and the chromosomes are concentrated near the two spindle poles. In our model this is represented by the deactivation of both unattached and attached kinetochore nodes. |
| | ⊢ P | CyclinB | *CyclinB/Cdk1* phosphorylate the N-terminal head region of nuclear lamins, causing their depolymerization and nuclear envelope breakdown [353]. |
| | ⊢ P | Cdk1 | *CyclinB/Cdk1* phosphorylate the N-terminal head region of nuclear lamins, causing their depolymerization and nuclear envelope breakdown [353]. |
| | ⊢ ComplProc | U _Kinetochores | Nuclear envelope reformation requires the removal of the Chromosome Passanger Complex (CPC) and Aurora B kinase from anaphase chromosomes. This occurs in anaphase, when the CPC relocalizes to the midline of the cell to aid cleavage furrow assembly [354]. In our model, *U_Kinetochores* = ON signals metaphase before the SAC is cleared, at which point the CPC is attached to centromeres and kinetochores. |

**Table S1k: Cell_Cycle_Process module**

| | | | |
|---|---|---|---|
| | ⊣ ComplProc | A _Kinetochores | Nuclear envelope reformation requires the removal of the Chromosome Passanger Complex (CPC) and Aurora B kinase from anaphase chromosomes. This occurs in anaphase, when the CPC relocalizes to the midline of the cell to aid cleavage furrow assembly [354]. In our model, *A_Kinetochores* = ON signals the end of metaphase and SAC clearance, at which point the CPC remains attached to centromeres and kinetochores. |
| f4N_DNA | **f4N_DNA** = ((**Replication** and ((**Pre_RC** and **CyclinA**) or **f4N_DNA**)) or (**f4N_DNA** and not **Cytokinesis**)) and not **CAD** | | |
| | MSt | 4N DNA content in our model is reached via the completion of *Replication* (via the firing of the last round of replication origins by *Cyclin A/Cdk* complexes) and maintained in non-apoptotic cells the absence of a contractile ring (marked by *Ect2*), driving cytokinesis. | |
| | ← ComplProc | Replication | DNA content is doubled by the process of *Replication*. |
| | ← ComplProc | Pre_RC | *Replication* can only complete DNA synthesis and produce double DNA content if the availability of licensed replication origins is not blocked [342]. |
| | ← ComplProc | CyclinA | *Cyclin A/Cdk1* complexes regulate the origin firing program in mammalian cells and are required for the completion of DNA replication [345, 290]. |
| | ⊣ Per | f4N_DNA | Once achieved, a cell's 4N DNA content is sustained up to the point of cytokinesis. |
| | ⊣ ComplProc | Cytokinesis | Cytokinesis resets the daughter cell DNA content to 2N. |
| | ⊣ Deg | CAD | Caspase-activated DNase (*CAD*) destroys DNA, preventing maintenance of a double DNA content. |
| U _Kinetochores | **U_Kinetochores** = **f4N_DNA** and not **Nuclear_Membrane** and not **Cdh1** and not **A_Kinetochores** and ((**CyclinB** and **Cdk1**) or **U_Kinetochores**) | | |
| | MSt | The *U_Kinetochores* node in our model is on from the moment the nuclear envelope is dissolved in prometaphase and the mitotic spindle starts to form, until all kinetochores are properly attached. In addition to the presence of unattached kinetochores, *U_Kinetochores* = ON requires attached sister chromatics, the absence of $APC/C^{Cdh1}$ activity. It is turned on my *Cyclin B/Cdk1* and remains on until the spindle is complete (or it is destroyed by $APC/C^{Cdh1}$). | |
| | ← ComplProc | f4N_DNA | Metaphase requires replicated sister chromatids (*f4N_DNA*), held together by their kinetochores, face in opposing directions and can be attached to opposite poles of the mitotic spindle. |
| | ⊣ Loc | Nuclear _Membrane | Dissolution of the nuclear membrane in prometaphase is required for unattached kinetochores to be accessible to microtublues and thus aid spindle formation [355]. |

**Table S1k: Cell_Cycle_Process module**

| | | | |
|---|---|---|---|
| | ⊣ ComplProc | Cdh1 | Premature activation of $APC/C^{Cdh1}$ destroys the incomplete spindle by triggering premature, aberrant anaphase. This occurs due to premature degradation of *APC/C* targets including *Securin* (responsible for keeping sister chromatids attached [356]), *Cyclin B*, *Cdc20*, and Aurora kinase A (*AURKA*) [357]. |
| | ⊣ ComplProc | A_Kinetochores | In our model, the transition from unattached to all attached kinetochores (*U_Kinetochores* → *A_Kinetochores*) marks the completion of the mitotic spindle and Spindle Assembly Checkpoint (SAC) passage. |
| | ← Per | U_Kinetochores | Once metaphase starts, the mitotic spindle remains incomplete as long as some of the kinetochores remain unattached. |
| | ← ComplProc | CyclinB | The start of mitotic spindle assembly is initiated by active *Cyclin B/Cdk1* [358]. |
| | ← ComplProc | Cdk1 | The start of mitotic spindle assembly is initiated by active *Cyclin B/Cdk1* [358]. |
| Bipolar_Spindle | **Bipolar_Spindle** = (**CyclinB** and **Cdk1** and **Unfused**) or (**Bipolar_Spindle** and **f4N_DNA** and not **Cytokinesis** and (**Ect2** or **Plk1_H** or ((**U_Kinetochores** or **A_Kinetochores**) and not **Cdh1**))) | | |
| | MSt | | The bipolar spindle node in our model represents a mitotic spindle attached to two centrosomes, moved to opposing poles of the dividing cell [355]. Thus it formed in response to *CyclinB/Cdk1* activation, aided by mitochondrial fragmentation. Once formed, it is maintained as long as the cell has 4N DNA (sister chromatids), has not undergone cytokinesis, and it is either still in metaphase (*U_Kinetochores* or *A_Kinetochores* are ON, $APC/C^{Cdh1}$ is OFF), or the cell is assembling a contractile ring (*Ect2* or *Plk1_H* are ON). |
| | ← Ind | CyclinB | *CyclinB/Cdk1* activation orchestrates the reorganization of the cell's microtubule network to drive mitotic spindle formation [359]. |
| | ← Ind | Cdk1 | *CyclinB/Cdk1* activation orchestrates the reorganization of the cell's microtubule network to drive mitotic spindle formation [359]. |
| | ← Ind | Unfused | Loss of *Drp1*-mediated mitochondrial fission induced mitochondrial aggregation around the microtubule organizing center (MTOC), centrosome over-duplication, severely misaligned metaphase chromosomes, and thus disruption of bipolar spindple formation [360, 120]. |
| | ⊣ Ind | Cdh1 | $APC/C^{Cdh1}$ activation leads to the separation of sister chromatids, and marks the start of anaphase [361]. |
| | ← Ind | f4N_DNA | Maintaining a bipolar spindle requires a cell with 4N DNA, either as attached sister chromatids or as separated chromosomes in telophase [355]. |
| | ← Per | Bipolar_Spindle | Once formed, the mitotic spindle persists until cytokinesis [355]. |

**Table S1k: Cell_Cycle_Process module**

| | | | |
|---|---|---|---|
| | $\leftarrow$ Ind | U _Kinetochores | Once formed, a bipolar mitotic spindle persists until through metaphase and the SAC [355]. |
| | $\leftarrow$ Ind | A _Kinetochores | Once formed, a bipolar mitotic spindle persists until through metaphase and the SAC [355]. |
| | $\leftarrow$ Ind | Plk1_H | As the *Plk1_H* node represents the active *Plk1* pool briefly protected from destruction by $APC/C^{Cdh1}$ [308] and responsible for recruiting *Ect2* to the central spindle, *Plk1_H* = ON marks ongoing anaphase before the cell assembles its contractile ring to trigger cytokinesis. Thus we use it as a logical condition for keeping the bipolar spindle node ON [355]. |
| | $\leftarrow$ Ind | Ect2 | As the *Ect2* node represents contractile ring assembly before cytokinesis, and as such we use it as a logical condition for keeping the bipolar spindle node ON [355]. |
| | $\dashv$ Ind | Cytokinesis | Cytokinesis separates the two poles of the bipolar spindple into two daughter cells, and thus turns the bipolar spindle node OFF [355]. |
| Mps1 | **Mps1** = **U_Kinetochores** or (**CyclinB** and **Cdk1**) | | |
| | K | | *Mps1* is a protein kinase critical to the spinde assembly checkpoint [362]. Once activated by *CyclinB/Cdk1* phosphorylation, *Mps1* inhibits cytosolic *Cdc20*, as it aids *Cdc20* binding to *Mad2* and *BubR1* before kinetochores mature. Then it recruits factors that sustain the SAC to unattached kinetochores (*Mps1* inhibition in prometaphase removes all known SAC mediators from kinetochores) [338]. |
| | $\leftarrow$ P | CyclinB | *CyclinB/Cdk1*-dependent phosphorylation of *Mps1* is required for its mitotic checkpoint activity [362]. |
| | $\leftarrow$ P | Cdk1 | *CyclinB/Cdk1*-dependent phosphorylation of *Mps1* is required for its mitotic checkpoint activity [362]. |
| | $\leftarrow$ Loc | U _Kinetochores | *Mps1* aids SAC protein assembly at unattached kinetochores [338]. |
| Mad2 | **Mad2** = **U_Kinetochores** and **Mps1** and not **A_Kinetochores** | | |
| | Prot | | Our model represents the SAC via the *Mad2* kinetochore-binding protein. *Mad2* is recruited to unattached kinetochores by *Mps1* and remains active as long as the cell has at least one unattached kinetochore and it is responsible for keeping *Cdc20* sequestered from *APC/C*. By keeping *APC* at bay until the spindle is complete, *Mad2* is required for the proper timing of anaphase [336]. |
| | $\leftarrow$ Compl | U _Kinetochores | The *Mad2* SAC protein is active and potent in the presence of even a single unattached kinetochore [336]. |
| | $\leftarrow$ Loc | Mps1 | Sustained *Mps1* activity is required for maintaining *Mad1/C-Mad2* complexes [363] and other SAC proteins at unattached kinetochores [338]. |
| | $\dashv$ ComplProc | A _Kinetochores | *Mad2* is inhibited by SAC passage, marked by the completion of the spindle and proper attachment of all kinetochore [336]. |

#### Table S1k: Cell_Cycle_Process module

| | | | |
|---|---|---|---|
| A _Kinetochores | **A_Kinetochores** = **f4N_DNA** and **Bipolar_Spindle** and **Plk1** and (**A_Kinetochores** or (**U_Kinetochores** and **CyclinB** and **Cdk1**)) and not **Cdh1** and not(**pAPC** and **Cdc20**) | | |
| | MSt | | The completed spindle, represented by the *A_Kinetochores* node, requires replicated and attached sister chromatids (*f4N_DNA*) and the absence of *APC/C* activity. It turns on when the process of spindle assembly (*U_Kinetochores*) is completed by active *Plk1* localized to unattached kinetochores in the presence of ongoing *Cyclin B/Cdk1* activity, and it remains on until anaphase (*APC/C* activation). |
| | ← ComplProc | f4N_DNA | Completion of the mitotic spindle requires replicated and attached sister chromatids (*f4N_DNA*). |
| | ← ComplProc | Bipolar _Spindle | Bipolar spindle formation is required for proper attachment and tension on all kinetochores [355]. |
| | ← ComplProc | Plk1 | *Plk1* activity at unattached kinetochores is required for promoting their attachment [364]. In its absence, kinetochores remain unattached and cells eventually undergo mitotic catastrophe and apoptosis [365]. |
| | ← Per | A _Kinetochores | Once assembled, separation of the mitotic spindle requires *APC/C* activity to promote the destruction of sister chromatid cohesion [356]. |
| | ← ComplProc | U _Kinetochores | The mitotic spindle is assembled gradually, as the number of unattached kinetochores gradually decreased by the formation of microtubule attachments. |
| | ← ComplProc | CyclinB | Ongoing *Cyclin B/Cdk1* at unattached kinetochores is necessary to keep *Plk1* active and allow the completion of mitosis [366]. |
| | ← ComplProc | Cdk1 | Ongoing *Cyclin B/Cdk1* at unattached kinetochores is necessary to keep *Plk1* active and allow the completion of mitosis [366]. |
| | ⊢ Deg | Cdh1 | $APC/C^{Cdh1}$ destroys the spindle by triggering anaphase via the degradation of *APC/C* targets, including *Securin* [357]. |
| | ⊢ Deg | pAPC | During normal mitosis, the completed spindle is pulled apart in response to $APC/C^{Cdc20}$-mediated degradation of *Securin*, which normally blocks *Separate* from severing the *Cohesin* rings keeping sister chromatids attached [356]. |
| | ⊢ Deg | Cdc20 | During normal mitosis, the completed spindle is pulled apart in response to $APC/C^{Cdc20}$-mediated degradation of *Securin*, which normally blocks *Separate* from severing the *Cohesin* rings keeping sister chromatids attached [356]. |
| Plk1_H | **Plk1_H** = **Plk1** and **FoxM1** and (**Plk1_H** or **FoxO3** or **FoxO1**) | | |
| | K | | The ON state of *Plk1_H* encodes the short-lived memory of a sufficiently large active *Plk1* pool to temporarily survive *Plk1* destruction by $APC/C^{Cdh1}$ [308], recruit *Ect2* to the central spindle, and thus aid the completion of cytokinesis [367]. Thus, *Plk1_H* requires ongoing *Plk1* activation and transcription by *FoxM1*, and either induction by *FoxO3* or *FoxO1*, or prior accumulation. |

**Table S1k: Cell_Cycle_Process module**

| | | | |
|---|---|---|---|
| | ← Per | Plk1 | Active mitotic *Plk1* is a prerequisite for the accumulation of the larger active *Plk1* pool denoted by *Plk1_H*. |
| | ← TR | FoxM1 | *Plk1* is a direct transcriptional target of *FoxM1*; loss of *FoxM1* severely reduces *Plk1* protein levels [264, 275]. |
| | ← Per | Plk1_H | Once accumulated, we assume that the *Plk1_H* pool of active *Plk1* remains stable in the absence of *FoxO*-mediated transcription. This is supported by negative feedback regulation of *FoxO* proteins by *Plk1* [21], indicating that ongoing high FoxO activity is likely not required for the maintenance of *Plk1_H*. |
| | ← TR | FoxO3 | *Plk1* is a direct transcriptional target of *FoxO3*, but *Plk1* appears to be sufficiently induced in the absence of *FoxO* preteens to aid its G2/M and mitotic functions. In contrast, accumulation if a large enough *Plk1* pool to briefly outlast $APC/C^{Cdh1}$ activation (modeled by the *Plk1_H* node), requires *FoxO* activity in G2 [302]. |
| | ← TR | FoxO1 | In addition to *FoxO3*, *FoxO1* also binds the *Plk1* promoter, potentially aiding its accumulation during G2 [368]. |
| Ect2 | **Ect2** = **f4N_DNA** and **Plk1_H** and **Cdh1** and not **U_Kinetochores** and not **A_Kinetochores** and **Bipolar_Spindle** | | |
| | GEF | *Ect2* activaiton at the spindle midzone represents the step of cytokinesis in our model. Thus, *Ect2* requires *f4N_DNA*, high *Plk1* activity, as well as *Cdh1* for the assembly of a normal spindle midzone. Finally, *Ect2* cannot be recruited to the mid zone before anaphase is completed. | |
| | ← Ind | f4N_DNA | Formation of a spindle midzone, where *Ect2* accumulates in preparation of cytokinesis requires recently separated sister chromatids (4N DNA content). |
| | ← Ind | Plk1_H | *Plk1* activity in telophase (*Plk1_H*) is required for the recruitment of *Ect2* to the central spindle [308, 369]. |
| | ← Ind | Cdh1 | $APC/C^{Cdh1}$-mediated destruction of Aurora kinase is required for the assembly of a robust spindle midzone at anaphase and for the normal timing of cytokinesis [370]. |
| | ⊣ ComplProc | U _Kinetochores | Formation of a spindle midzone requires the separation of sister chromatids; thus it cannot occur before anaphase. |
| | ⊣ ComplProc | A _Kinetochores | Formation of a spindle midzone requires the separation of sister chromatids; thus it cannot occur before anaphase. |
| | ← Loc | Bipolar _Spindle | The midzone of the bipolar spindle serves as source of signals that specify the location of contractile ring assembly and constriction, including recruitment of the *Ect2* RhoA GEF [371]. |
| Cytokinesis | **Cytokinesis** = **Ect2** and **Bipolar_Spindle** | | |

**Table S1k: Cell_Cycle_Process module**

| | | | |
|---|---|---|---|
| | Proc | In contrast to our previous model in [372] where *Ect2* recruitment to the central spindle marked the start of cytokinesis, in [373] we introduced a separate *Cytokinesis* node to mark cytokinesis and the subsequent resetting of daughter cell DNA content to 2N by a separate node. In addition to *Ect2* recruitment, completion of cytokinesis also requires a bipolar spindle [371]. | |
| | ← Ind | Ect2 | At the start of cytokinesis, the *Ect2* RhoGEF is recruited to the central spindle [367]. *Ect2* aids the accumulation of GTP-bound *RhoA* [374] and the formation of the contractile ring. In our model, *Ect2* recruitment to the central spindle marks the start of cytokinesis. |
| | ← Loc | Bipolar _Spindle | The midzone of the bipolar spindle specifes the location of contractile ring assembly and constriction [371]. |

**Table S1l: DNA_Damage_Signaling module**

| Target Node | Node Gate<br>Node Type<br>Link Type | Node Description<br>Input Node | Link Description |
|---|---|---|---|
| cROS | **cROS** = not **Glucose** or **ROS_Ext** or **cROS_H** | | |
| | Met | The *cROS* node represents a moderate rise in cytosolic ROS. This may be the result of glucose starvation [375] or external ROS exposure. The node is also ON by default if the cyrosolic levels are high (*cROS_H* = ON). | |
| | ⊢ ComplProc | Glucose | Glucose starvation blocks glycolysis and the pentose phosphate pathway, resulting in oxidative stress due to enhanced ROS production (e.g. $H_2O_2$) [375]. |
| | ← Loc | ROS_Ext | External ROS molecules such as the nonpolar $H_2O_2$ can cross the cell membrane and raise cytosolic ROS levels. |
| | ← Per | cROS_H | As the *cROS_H* node represents a higher cytosolic ROS burden with qualitatively different downstream consequences from the *cROS* node, *cROS_H* = ON is sufficient to keep the *cROS* node ON. |
| cROS_H | **cROS_H** = **cROS** and (**ROS_Ext** or (**mROS** and not **Glucose** and not **PGC1**)) | | |
| | Met | The *cROS_H* node represents high cytosolic ROS, above a threshold where its consequences are qualitatively distinct from those of the moderate *cROS* node (i.e., DNA damage leading to *ATM/ATR* activation and sustained high *p53* levels). In addition to moderate *cROS*, this requires external ROS or the combined effects of glucose starvation [375], mitochondrial ROS generation and a lack of *PGC-1*$\alpha$-driven antioxidant response [180]. | |
| | ← Per | cROS | *cROS_H* = ON is a step up from moderate ROS levels and thus requires *cROS* = ON. |
| | ← Loc | ROS_Ext | External ROS molecules such as the nonpolar $H_2O_2$ can cross the cell membrane and raise cytosolic ROS levels. |

**Table S1l: DNA_Damage_Signaling module**

| | | | |
|---|---|---|---|
| | ⊢ ComplProc | Glucose | Glucose starvation blocks glycolysis and the pentose phosphate pathway, resulting in oxidative stress due to enhanced ROS production (e.g. $H_2O_2$) [375]. |
| | ← Loc | mROS | As mitochondria are the primary sources of intracellular ROS, increased mROS production can raise cytosolic levels as well. |
| | ⊢ ComplProc | PGC1 | *PGC-1α* increases the expression of mitochondrial antioxidant genes such as *MnSOD*, catalase, peroxiredoxins, *UCP2*, thioredoxin 2, thioredoxin reductase [180]. |
| ATM | **ATM** = **cROS_H** or **CAD** | | |
| | K | Ataxia-telangiectasia mutated, or ATM serine/threonine kinase, is recruited to and activated by DNA double-strand breaks [349]. In addition, oxidative stress can directly activate *ATM* [376]. | |
| | ← Cat | cROS_H | Elevated cytosolic ROS directly activte *ATM* by creating active, disulfide-linked dimers [376]. In addition, double-strand damage caused by ROS can also activate ATM indirectly [349]. |
| | ← Ind | CAD | *CAD*-induced DNA fragmentation during apoptosis activates ATM [377]. |
| CHK2 | **CHK2** = **ATM** | | |
| | K | *ATM* kinase activates *CHK2* at DNA double strand breaks [351, 349]. | |
| | ← P | ATM | *ATM* kinase activates *CHK2* at DNA double-strand breaks, which blocks cell cycle progression [351, 349]. |
| p53 | **p53** = (**Nuclear_Membrane** and (((**AMPK** or (**ATR** and **CHK1**) or (**ATM** and **CHK2**)) and (not **Mdm2** or not **NF_kB** or **cROS** or (**mROS** and not **SIRT3**))))) or **p53_2** or **p53_3** or **p53_4** | | |
| | TF | *p53* is stabilized and its protein levels increase in response to stress. Here we use four Boolean nodes to track the sustained rise of *p53* and/or sustained oscillations that slowly boost/decrease mRNA expression in a subset of targets (others respond to the early rise of p53). We choose 4 steps because the time required for 4 consecutive pulses is a good qualitative match to the time it takes for *p53* to block *Cyclin B* expression during prolonged G2 arrest, *p21* mRNA expression saturates after 2-3 pulses [378, 258, 379]. The *p53* node represents the first level. Its activation requires a nuclear membrane (mitotic *p53* is sequestered to centrosomes) [380], phoshorylation by *AMPK* [381], *ATR/CHK1* or *ATM/CHK2* [351, 382], and either a lack of *Mdm2*-mediated degradation [383, 384, 385, 351], competition for co-activators with *NF-κB* [78], cytosolic ROS [386], or mitochondrial ROS in the absence of *SIRT3* [387]. In addition, the *p53* node is ON by default if any of the three nodes marking higher/prologed activation are also ON. | |
| | ← Loc | Nuclear _Membrane | During mitosis, *p53* is recruited to and kept at centrosomes (a localization process aided by basal *ATM*-mediated Ser-15 phosphorylation), where it is dephosphorylated and kept inactive to allow cell cycle progression in cells with normal mitotic spindles [380]. Thus, we assume that the lack of a nuclear membrane keeps *p53* transcription OFF. |

**Table S1l: DNA_Damage_Signaling module**

| | | |
|---|---|---|
| ← P | AMPK | *AMPK* induces Ser-15 phosphorylation of *p53* to initiate an *AMPK*-dependent cell-cycle arrest [381]. |
| ← P | ATR | *ATR* induces Ser-15 phosphorylation of *p53* in response to single-strand DNA damage, and activates it with the aid of *CHK1* [351]. |
| ← P | CHK1 | *CHK1/2* phosphorylate *p53* at Ser-20 [388] to prevent its ubiquitination by the ubiquitin E3 ligase *Mdm2* and subsequent proteosomal degradation [382]. |
| ← P | ATM | *ATR* induces Ser-15 phosphorylation of *p53* in response to double-strand DNA breaks, and activates it with the aid of *CHK2* [351]. |
| ← P | CHK2 | *CHK1/2* phosphorylate *p53* at Ser-20 [388] to prevent its ubiquitination by the ubiquitin E3 ligase *Mdm2* and subsequent proteosomal degradation [382]. |
| ⊢ Deg | Mdm2 | *Mdm2*, induced by *p53* [389], is part of a negative feedback responsible of shutting down *p53* accumulation when damage signals turn off. *Mdm2* poly-ubiquitinates *p53*, targeting it for degradation [383, 384, 385, 351]. |
| ⊢ BLoc | NF_kB | *NF-κB*-mediated inhibition of *p53* is likely the result of their competition for a limited pool of *p300/CBP complexes*, as they both require these transcriptional coactivators [78]. |
| ← Ind | cROS | ROS activate *p38 MAPK* (in addition to *ATM*), which in turn phosphorylates *p53* at Ser15, 33, 37 and 46, leading to its activation under increased ROS [386]. |
| ← Ind | mROS | ROS activate *p38 MAPK* (in addition to *ATM*), which in turn phosphorylates *p53* at Ser15, 33, 37 and 46, leading to its activation under increased ROS [386]. |
| ⊢ Deacet | SIRT3 | *SIRT3* deacetylates the mitochodrial pool of *p53*, reducing its transcriptional activity (*p53* localized to the nucleus as well as the mitocniodria when its levels rise above basal) [387]. |
| ← Per | p53_2 | We assume that the transcriptional influence of *p53*, either from the accumulation of its active (Ser15-phosphorylated and acetylated) form or from repeated pulses of *p53* oscillation (in response to double-strand breaks; not modeled here), accumulates and dissapears stepwise from level 1 (*p53* node representing accumulation rought equivalent to a single pulse) to level 4 (*p53_4* node, transcritional activity on par with the effect of 4 consecutive pulses). Thus, the *p53* node is kept ON by default when nodes representing higher activity levels are ON. |
| ← Per | p53_3 | The *p53* node is kept ON by default when all nodes representing higher activity levels are ON (see *p53_2* in-link for details). |
| ← Per | p53_4 | The *p53* node is kept ON by default when all nodes representing higher activity levels are ON (see *p53_2* in-link for details). |

**Table S1l: DNA_Damage_Signaling module**

p53_2 — **p53_2** = ((**p53** or **p53_2**) and ((**AMPK** or (**ATR** and **CHK1**) or (**ATM** and **CHK2**)) and **cROS_H** and **Nuclear_Membrane**)) or **p53_3** or **p53_4**

TF — The *p53_2* node represents the second level/pulse of *p53* activation. It's accumulation and maintenance requires *p53* or *p53_2*, a nuclear membrane [380], phoshorylation by *AMPK* [381], *ATR/CHK1* or *ATM/CHK2* [351, 382], and high cytosolic ROS [386]. In addition, the *p53_2* node is ON by default if any of the two nodes marking higher/prologed activation are also ON.

| | | |
|---|---|---|
| ← Per | p53 | We assume that the transcriptional influence of *p53*, either from the accumulation of its active (Ser15-phosphorylated and acetylated) form or from repeated pulses of *p53* oscillation, accumulates and dissapears stepwise from level 1 (*p53* node representing accumulation rought equivalent to a single pulse) to level 4 (*p53_4* node, transcritional activity on par with the effect of 4 consecutive pulses). Thus, the *p53_2* node turns ON/OFF when nodes representing the lower/higher activity levels are also ON/OFF. |
| ← Per | p53_2 | The *p53_2* node can be maintained ON by ongoing activating signals. |
| ← Loc | Nuclear _Membrane | During mitosis, *p53* is recruited to and kept at centrosomes (a localization process aided by basal *ATM*-mediated Ser-15 phosphorylation), where it is dephosphorylated and kept inactive to allow cell cycle progression in cells with normal mitotic spindles [380]. Thus, we assume that the lack of a nuclear membrane keeps *p53* transcription OFF. |
| ← P | AMPK | *AMPK* induces Ser-15 phosphorylation of *p53* to initiate an *AMPK*-dependent cell-cycle arrest [381]. |
| ← P | ATR | *ATR* induces Ser-15 phosphorylation of *p53* in response to single-strand DNA damage, and activates it with the aid of *CHK1* [351]. |
| ← P | CHK1 | *CHK1/2* phosphorylate *p53* at Ser-20 [388] to prevent its ubiquitination by the ubiquitin E3 ligase *Mdm2* and subsequent proteosomal degradation [382]. |
| ← P | ATM | *ATR* induces Ser-15 phosphorylation of *p53* in response to double-strand DNA breaks, and activates it with the aid of *CHK2* [351]. |
| ← P | CHK2 | *CHK1/2* phosphorylate *p53* at Ser-20 [388] to prevent its ubiquitination by the ubiquitin E3 ligase *Mdm2* and subsequent proteosomal degradation [382]. |
| ← Ind | cROS_H | ROS activate *p38 MAPK* (in addition to *ATM*), which in turn phosphorylates *p53* at Ser15, 33, 37 and 46, leading to its activation under increased ROS [386]. |
| ← Per | p53_3 | The *p53_2* node turns ON/OFF when all nodes representing the lower/higher activity levels are also ON/OFF. |
| ← Per | p53_4 | The *p53_2* node turns ON/OFF when all nodes representing the lower/higher activity levels are also ON/OFF. |

**Table S1l: DNA_Damage_Signaling module**

| | | | |
|---|---|---|---|
| p53_3 | **p53_3** = (((**p53_2** and **p53**) or **p53_3**) and ((**AMPK** or (**ATR** and **CHK1**) or (**ATM** and **CHK2**)) and **cROS_H** and **Nuclear_Membrane**)) or **p53_4** | | |
| | TF | | The *p53_3* node represents the third level/pulse of *p53* activation. It's accumulation and maintenance requires *p53* / *p53_2* or *p53_2*, a nuclear membrane [380], phoshorylation by *AMPK* [381], *ATR/CHK1* or *ATM/CHK2* [351, 382], and high cytosolic ROS [386]. In addition, the *p53_3* node is ON by default if *p53_4* is also ON. |
| | ← Per | p53 | We assume that the transcriptional influence of *p53*, either from the accumulation of its active (Ser15-phosphorylated and acetylated) form or from repeated pulses of *p53* oscillation, accumulates and dissapears stepwise from level 1 (*p53* node representing accumulation rought equivalent to a single pulse) to level 4 (*p53_4* node, transcritional activity on par with the effect of 4 consecutive pulses). Thus, the *p53_3* node turns ON/OFF when nodes representing the lower/higher activity levels are also ON/OFF. |
| | ← Per | p53_2 | The *p53_3* node turns ON/OFF when all nodes representing the lower/higher activity levels are also ON/OFF. |
| | ← Per | p53_3 | The *p53_3* node can be maintained ON by ongoing activating signals. |
| | ← Loc | Nuclear _Membrane | During mitosis, *p53* is recruited to and kept at centrosomes (a localization process aided by basal *ATM*-mediated Ser-15 phosphorylation), where it is dephosphorylated and kept inactive to allow cell cycle progression in cells with normal mitotic spindles [380]. Thus, we assume that the lack of a nuclear membrane keeps *p53* transcription OFF. |
| | ← P | AMPK | *AMPK* induces Ser-15 phosphorylation of *p53* to initiate an *AMPK*-dependent cell-cycle arrest [381]. |
| | ← P | ATR | *ATR* induces Ser-15 phosphorylation of *p53* in response to single-strand DNA damage, and activates it with the aid of *CHK1* [351]. |
| | ← P | CHK1 | *CHK1/2* phosphorylate *p53* at Ser-20 [388] to prevent its ubiquitination by the ubiquitin E3 ligase *Mdm2* and subsequent proteosomal degradation [382]. |
| | ← P | ATM | *ATR* induces Ser-15 phosphorylation of *p53* in response to double-strand DNA breaks, and activates it with the aid of *CHK2* [351]. |
| | ← P | CHK2 | *CHK1/2* phosphorylate *p53* at Ser-20 [388] to prevent its ubiquitination by the ubiquitin E3 ligase *Mdm2* and subsequent proteosomal degradation [382]. |
| | ← Ind | cROS_H | ROS activate *p38 MAPK* (in addition to *ATM*), which in turn phosphorylates *p53* at Ser15, 33, 37 and 46, leading to its activation under increased ROS [386]. |
| | ← Per | p53_4 | The *p53_3* node turns ON/OFF when all nodes representing the lower/higher activity levels are also ON/OFF. |

**Table S1l: DNA_Damage_Signaling module**

| | | | |
|---|---|---|---|
| p53_4 | **p53_4** = ((**p53** and **p53_2** and **p53_3**) or **p53_4**) and ((**AMPK** or (**ATR** and **CHK1**) or (**ATM** and **CHK2**)) and **cROS_H** and **Nuclear_Membrane**) | | |
| | TF | The *p53_4* node represents the fourth and highest level/pulse of *p53* activation. It's accumulation and maintenance requires *p53* / *p53_2* / *p53_3* or *p53_4*, a nuclear membrane [380], phoshorylation by *AMPK* [381], *ATR/CHK1* or *ATM/CHK2* [351, 382], and high cytosolic ROS [386]. | |
| | ← Per | p53 | We assume that the transcriptional influence of *p53*, either from the accumulation of its active (Ser15-phosphorylated and acetylated) form or from repeated pulses of *p53* oscillation, accumulates and dissapears stepwise from level 1 (*p53* node representing accumulation rought equivalent to a single pulse) to level 4 (*p53_4* node, transcritional activity on par with the effect of 4 consecutive pulses). Thus, the *p53_4* node turns ON/OFF when nodes representing the lower/higher activity levels are also ON/OFF. |
| | ← Per | p53_2 | The *p53_4* node turns ON/OFF when all nodes representing the lower/higher activity levels are also ON/OFF. |
| | ← Per | p53_3 | The *p53_4* node turns ON/OFF when all nodes representing the lower/higher activity levels are also ON/OFF. |
| | ← Per | p53_4 | The *p53_4* node can be maintained ON by ongoing activating signals. |
| | ← Loc | Nuclear _Membrane | During mitosis, *p53* is recruited to and kept at centrosomes (a localization process aided by basal *ATM*-mediated Ser-15 phosphorylation), where it is dephosphorylated and kept inactive to allow cell cycle progression in cells with normal mitotic spindles [380]. Thus, we assume that the lack of a nuclear membrane keeps *p53* transcription OFF. |
| | ← P | AMPK | *AMPK* induces Ser-15 phosphorylation of *p53* to initiate an *AMPK*-dependent cell-cycle arrest [381]. |
| | ← P | ATR | *ATR* induces Ser-15 phosphorylation of *p53* in response to single-strand DNA damage, and activates it with the aid of *CHK1* [351]. |
| | ← P | CHK1 | *CHK1/2* phosphorylate *p53* at Ser-20 [388] to prevent its ubiquitination by the ubiquitin E3 ligase *Mdm2* and subsequent proteosomal degradation [382]. |
| | ← P | ATM | *ATR* induces Ser-15 phosphorylation of *p53* in response to double-strand DNA breaks, and activates it with the aid of *CHK2* [351]. |
| | ← P | CHK2 | *CHK1/2* phosphorylate *p53* at Ser-20 [388] to prevent its ubiquitination by the ubiquitin E3 ligase *Mdm2* and subsequent proteosomal degradation [382]. |
| | ← Ind | cROS_H | ROS activate *p38 MAPK* (in addition to *ATM*), which in turn phosphorylates *p53* at Ser15, 33, 37 and 46, leading to its activation under increased ROS [386]. |

#### Table S1l: DNA_Damage_Signaling module

| | | | |
|---|---|---|---|
| Mdm2 | **Mdm2** = not(**ATM** or **ATR**) and (**AKT_H** or **p53** or **p53_2** or **p53_3** or **p53_4**) and not(**Casp2** or **Casp3**) | | |
| | UbL | The *Mdm2* E3 ubiquitin-protein ligase is a key inhibitor of *p53* accumulation [385, 384]. It's activity is inhibited by *ATM/ATR*-mediated phosphorylation [351], while its levels are increased by *AKT* phosphorylation (stabilization) or *p53*-mediated transcription. Finally, *Caspase2/3* cleave and deactivate *Mdm2*. | |
| | $\vdash$ P | ATM | *ATM* phosphorylates *Mdm2* at Ser-395, leading to its deactivation as well as rapid degradation [351]. |
| | $\vdash$ P | ATR | *ATR* phosphorylates *Mdm2* at Ser-407, inhibiting its ability to bind and block *p53* accumulation [351]. |
| | $\leftarrow$ P | AKT_H | *AKT* phosphorylates *Mdm2* at Ser-183, inreasing its nuclear stability and decreasing *p53* levels [390]. |
| | $\leftarrow$ TR | p53 | *p53* is a direct transcriptional inducer of *Mdm2*[383]. As its levels rise rapidly within the frist few hours of *p53* activation (see SM data in [378]), we model it as induced by any of the four *p53* nodes. |
| | $\leftarrow$ TR | p53_2 | *p53* is a direct transcriptional inducer of *Mdm2*[383]. |
| | $\leftarrow$ TR | p53_3 | *p53* is a direct transcriptional inducer of *Mdm2*[383]. |
| | $\leftarrow$ TR | p53_4 | *p53* is a direct transcriptional inducer of *Mdm2*[383]. |
| | $\vdash$ Lysis | Casp2 | *Caspase 2* cleaves *Mdm2* at Asp-367, removing its C-terminal RING domain required for *p53* ubiquitination [391]. |
| | $\vdash$ Lysis | Casp3 | *Caspase 2* cleaves and deactivates *Mdm2* [392]. |
| p21_H | **p21_H** = **p21_mRNA** and **p21** and **p53_3** and not **Casp3** | | |
| | CDKI | The *p21_H* node represents high, cell cycle arresting and senescence-promoting levels of *p21* accumulation in response to damage. Thus, this node requires *p21* mRNA expression, at least basal *p21*, and sustained increase / pulsing of *p53* (node *p53_3*) [393], as well as the absence of *Caspase 3*-mediated cleavage [189]. | |
| | $\leftarrow$ TL | p21 _mRNA | Increased mRNA levels are required to raise *p21* protein levels [378]. |
| | $\leftarrow$ Per | p21 | At least basal *p21* protein activity is required before the high *p21* node turns ON. |
| | $\leftarrow$ TR | p53_3 | *p21* mRNA levels increase with consecutive *p53* pulses, reaching saturation after 3-4 pulses on average [393]. Thus we chose to link our *p21_H* node to *p53_3*. |
| | $\vdash$ Lysis | Casp3 | *Caspase 3* cleaves and deactivates *p21* [189]. |

**Table S1m: TRAIL module**

| Target Node | Node Gate<br>Node Type | Node Description<br>Link Type | Input Node | Link Description |
|---|---|---|---|---|
| Trail | **Trail** = **Trail** | | | |
| | Env | The *Trail* node represents environmental availability of the *Trail* protein outside the cell. | | |
| | ← Env | Trail | The *Trail* input node remains on/off if set ON/OFF in the absence of *in silico* perturbation. | |

**Table S1n: Apoptotic_SW module**

| Target Node | Node Gate<br>Node Type | Node Description<br>Link Type | Input Node | Link Description |
|---|---|---|---|---|
| DR4_5 | **DR4_5** = **Trail** | | | |
| | Rec | The *DR4* and *DR5* death receptors, represented by the *DR4_5* node, are activated by extracellular *Trail* [394]. | | |
| | ← Ligand | Trail | *DR4* and *DR5* death receptors are activated by extracellular *Trail* [394]. | |
| Casp8 | **Casp8** = **DR4_5** or **Casp3** | | | |
| | PTase | *Pro-Caspase 8* may be cleaved independently by *DISC* (not directly reprsented; an adaptor protein for *DR4_5*) or *Caspase 3*. | | |
| | ← Compl | DR4_5 | *Trail*-bound (active) *DR4* and *DR5* receptors trigger the assembly of the pro-apoptotic death-inducing signaling complex (*DISC*), which binds a cluster of *pro-Caspase 8* proteins and initiates their cleavage into active *Caspase 8* [395]. | |
| | ← Ind | Casp3 | *Caspase 3* indirectly activates *Caspase 8* by cleaving *Caspase 6* [396], which, in turn, cleaves *Caspase 8* [397]. | |
| Casp2 | **Casp2** = (**U_Kinetochores** and **Mad2** and not(**CyclinB** and **Cdk1** and **Bipolar_Spindle**)) or **Casp3** | | | |
| | PTase | *Pro-caspase 2* is cleaved and activated by *Caspase 3*, or by failed cytokinesis marked by the presence of unattached kinetochores, an active SAC, and the absence of active *Cyclin B/Cdk1* complexes alongside a properly formed bipoar spindle to phosphorylate and inhibit *Caspase 2*. | | |

**Table S1n: Apoptotic_SW module**

| | | | |
|---|---|---|---|
| | ← Ind | U _Kinetochores | Although the precise molecular mechanism by which *Caspase 2* is activated during prolonged or stalled mitosis is unclear, its activation platform, the *PIDDosome*, has been localized to unattached kinetochores [398]. Even though a checkpoint protein keeps the *PIDDosome* unresponsive to DNA damage signals, the loss of protective *Cyclin B/Cdk1* phosphorylation only leads to *Caspase 2* activation in the presence of a partially assembled mitotic spindle, and requires active SAC. |
| | ← Ind | Mad2 | A functional spindle assembly checkpoint is required for mitotic cell death upon prolonged mitotic arrest [399] or spindle damage [400]. |
| | ⊣ P | CyclinB | *Cyclin B1/Cdk1* phosphorylate *caspase-2* at Ser 340, preventing its activation [401]. |
| | ⊣ P | Cdk1 | *Cyclin B1/Cdk1* phosphorylate *caspase-2* at Ser 340, preventing its activation [401]. |
| | ⊣ P | Bipolar _Spindle | Abberant mitotic spindle formation leads to *Caspase 2* activation followed by mitotic catastrophe [402, 403]. |
| | ← Lysis | Casp3 | *Caspase 2* is a target of *Caspase 3*, as its inhibition severely limits *Caspase 2* cleavage during apoptosis [404, 405]. |
| MCL_1 | **MCL_1** = not **Casp3** and not **Casp2** and (not **GSK3** or (**AKT_B** and (**ERK** or not **E2F1**))) and not(**Cdk1** and **CyclinB** and **U_Kinetochores**) | | |
| | Prot | | *Caspase 3* or *2*-mediated destruction of *MCL-1* must be absent for *MCL-1* to be ON. Avoiding degradation via textitGSK3 requires the *GSK3*-weakenjng presence of basal *AKT* activity (*AKT_B*) [406] and either *ERK*-mediated stabilization, or the absence of its repressor *E2F1*. Finally, during mitotic arrest (*U_Kinetochores*), *MCL-1* is deactivated by *Cyclin B/Cdk1* phosphorylation, which shields it from the *PPA2*-mediated dephosphorylation of its degradation-targeting sites [407]. |
| | ⊣ Lysis | Casp3 | *Caspase 3* cleaves and deactivated *MCL-1* [408]. |
| | ⊣ Lysis | Casp2 | *Caspase 2* activation destabilizes the *MCL-1* protein [409]. |
| | ⊣ P | GSK3 | *MCL-1* is phosphorylated by *GSK3*, leading to ubiquitinylation and degradation of Phosphorylation [406]. |
| | ← Ind | AKT_B | In order to account for the loss of *MCL-1* in the complete absence of growth factors versus its presence in low growth factor environments, we required basal *AKT* to modulate the strength of *GSK3* inhibition [406]. |
| | ← P | ERK | *ERK* phosphorylates *MCL-1*, promoting its interaction with *Pin1*, which stabilizes it [410, 411]. |
| | ⊣ TR | E2F1 | *E2F1* is a direct transcriptional repressor of *MCL-1* [412]. |
| | ⊣ P | CyclinB | In cells arrested in mitosis, phosphorylation by *Cyclin B/Cdk1* on T92 initiates *MCL-1* degradation [413]. |

**Table S1n: Apoptotic_SW module**

| | | | |
|---|---|---|---|
| | ⊣ P | Cdk1 | Phosphorylation by *Cyclin B/Cdk1* in cells arrested in mitosis initiates *MCL-1* degradation [413]. |
| | ⊣ Ind | U _Kinetochores | During prolonged mitotic arrest (*U_kinetochores*), *MCL-1* levels drop steadily due to phosphorylation by *JNK*, *p38* and/or *CKII* and its subsequent degradation by the E3 ubiquitin ligase *SCF* (*FBW7*) [407]. |
| BCLXL | **BCLXL** = not **Casp3** and (**BCL2** or not **BAD**) and (not **U_Kinetochores** or (**Plk1** and (not(**CyclinB** and **Cdk1**) or (**BCL2** and **MCL_1**) or **Bipolar_Spindle**)) or ((**BCL2** and **MCL_1**) and (not(**CyclinB** and **Cdk1**)))) | | |
| | Prot | | *Bcl-xL* activity requires the absence of *Caspase 3*. In addition, *BAD* can block *Bcl-xL* when *BCL2* is also OFF. Finally, mitotic *BCL-xL* can be inhibited by *Cdk1* activity if either *BCL2*, *MCL-1*, or *Plk1* are OFF, or bipolar spindle formation is compromised. In the absence of *Plk1*, loss of either *BCL2* or *MCL-1* can result in *Bcl-xL* inhibition (even without *Cdk1* phosphorylation), as we assume its targets are no longer competitively bound by its family members. |
| | ← Ind | Plk1 | In addition to other effects of prolonged mitotic arrest on *BCL-2* proteins, *Plk1* inhibition synergistically enhances the inhibitory phosphorylation of *BCL-2* and *BCL-xL*, as well as downregulation of *MCL-1* [414]. |
| | ⊣ P | CyclinB | During normal mitosis, *Cyclin B/Cdk1* only transiently phosphorylates part of the *BCL-xL* pool. Prolonged mitosis, however, results in high levels of *BCL-xL* (and *Bcl-2*) phosphorylation, priming the system for *Caspase 2*-mediated apoptosis [415, 416]. |
| | ⊣ P | Cdk1 | During normal mitosis, *Cyclin B/Cdk1* only transiently phosphorylates part of the *BCL-xL* pool. Prolonged mitosis, however, results in high levels of *BCL-xL* (and *Bcl-2*) phosphorylation, priming the system for *Caspase 2*-mediated apoptosis [415, 416]. |
| | ← Ind | Bipolar _Spindle | Prolonged mitosis, without proper bipolar spindle formation leading to SAC passage, is required for the accumulation of *BCL-xL* phosphorylation, which weaken its interaction with *Bax* [417]. |
| | ⊣ Ind | U _Kinetochores | Prolonged mitosis is required for the accumulation of *BCL-xL* phosphorylation, weakening its interaction with *Bax* [417]. |
| | ← Ind | MCL_1 | *MCL-1* competes with *BCL-xL* for *BAK* binding; the presence of *MCL-1* can keep part of the *BCL-xL* pool active [418]. |
| | ← Ind | BCL2 | *BCL2* competes with *BCL-xL* for *BAD* binding. Although *BCL-xL* is a stronger binding partner, we assume that *BAD* alone cannot fully block *BCL-xL* in the presence of *BCL2* [419]. |
| | ⊣ IBind | BAD | *Bad* can bind *BCL-xL* and displace it from *BAX*, thus deactivating it [419]. |

#### Table S1n: Apoptotic_SW module

| | | | |
|---|---|---|---|
| | ⊢ Lysis | Casp3 | *BCL2* is cleaved and deactivated by *Caspase 3* [420]. |
| BCL2 | **BCL2** = not(**Casp3** or **BAD** or **BIM** or **BIK** or (**BID** and **HIF1**)) and ((not **U_Kinetochores** or (**MCL_1** and **BCLXL**)) or (**Plk1** and (**BCLXL** or **MCL_1** or not(**Cdk1** and **CyclinB**)))) | | |
| | Prot | While the precise combinatorial logic governing *BCL2* activity is not clear from literature, we modeled *BCL2* as ON in the absence of *Caspase 3*, *BAD*, *BIM* or *BIK*. This choice makes *BCL2* the most sensitive of the three family members to activation of its three inhibitors. In addition, mitotic *BCL2* is blocked by *Cdk1* if both *BCL-xL* and *MCL-1* are OFF. In the absence of *Plk1*, loss of either *BCL2* or *MCL-1* can result in *BCL-2* inhibition (even without *Cdk1* phosphorylation), as we assume its targets are no longer competitively bound by its family members. In order to account for the apoptosis-sensitising role of *HIF-1α*, we assume that truncated *BID* can also repress *BCL2* when *HIF-1α* activation ups the ratio of *BAX* to *BCL2* [421]. | |
| | ⊢ Lysis | Casp3 | *BCL2* is cleaved and deactivated by *Caspase 3* [420]. |
| | ⊢ IBind | BAD | *BCL2* competes with *BCL-xL* for *BAD* binding. *BAD* displaces *BCL2* from its inhibitory binding of *Bax/Bak*. Although *BCL-xL* is a stronger binding partner, we assume that *BAD* alone cannot fully block *BCL-xL* in the presence of *BCL2* [419]. |
| | ⊢ IBind | BIM | *BIM* binds *BCL2* and they mutually inhibit each other's ability to activate further targets [422]. |
| | ⊢ IBind | BIK | *BIK* binds *BCL2* and they mutually inhibit each other's activity [423]. |
| | ⊢ IBind | BID | *BID* binds *BCL2* and they mutually inhibit each other's ability to activate further targets [424]. We assume that *tBID* can repress *BCL2* when *HIF-1α* is active, as it increased the ratio of *BAX* to *BCL2* proteins [421]. |
| | ⊢ Ind | HIF1 | *HIF-1α* was shown to increase the ratio of *Bax* to *BCL2* proteins, as well as upregulate the levels of extrinsic death signaing proteins (including *Trail* and *DR4/5*) while downregulating *c-FLIP* [421]. Thus, we assume that *HIF-1α* can boost *Trail* signaling to overpower *BCL2* even in the presence of potent survival signals. |
| | ⊢ Ind | U _Kinetochores | Prolonged mitosis is required for the accumulation of *BCL2* phosphorylation [415, 416, 420]. |
| | ← Ind | MCL_1 | *MCL-1* competes with *BCL-xL* for binding most of their apoptotic partners, including *BIK*, *BIM*, *BID*, *BAX* and *BAK*. |
| | ← Ind | BCLXL | *BCL2* competes with *BCL-xL* for binding most of their apoptotic partners, including *BIK*, *BIM*, *BID*, *BAX* and *BAK*. |

**Table S1n: Apoptotic_SW module**

| | | | |
|---|---|---|---|
| | ← Ind | Plk1 | In addition to other effects of prolonged mitotic arrest on *BCL2* proteins, *Plk1* inhibition synergistically enhances the inhibitory phosphorylation of *BCL2* and *BCL-xL*, as well as downregulation of *MCL-1* [414]. |
| | ⊢ P | CyclinB | *Cyclin B/Cdk1* phosphorylates *BCL2* (and *BCL-xL*) during mitosis [415, 416, 420]. |
| | ⊢ P | Cdk1 | Prolonged mitosis results in high levels of *BCL-xL* and *BCL2* phosphorylation, priming the system for *Caspase 2*-mediated apoptosis [415, 416, 420]. |
| BAD | **BAD** = **Casp3** or not(**AKT_H** or **AKT_B** or **ERK** or **S6K**) or (**Casp8** and not(**AKT_B** and **ERK** and **S6K**) and not(**AKT_H** and (**AKT_B** or **ERK**))) | | |
| | Prot | | *BAD* in our model is ON when cleaved by *Caspase 3*, or in the complete absence of survival signals (*AKT*, *ERK* or *S6K*). Alternatively, *BAD* can be cleaved and activated by *Caspase 8* in the absence of strong survival signaling. We modeled this inhibitory survival signal as either the combined activity of *ERK*, *S6K* and (at least) basal *AKT*, or high *AKT* in the joint presence of ERK and basal *AKT* (indicating that *AKT_H* will not drop by the next time-step). |
| | ← Lysis | Casp3 | *Caspase 3* cleaves *BAD*, generating a more potently apoptotic fragment [425]. |
| | ⊢ P | AKT_H | *Akt* phosphorylates *BAD* at Ser-136, inducing its sequestration away from the mitochondrial membrane where its BCL-2 family targets are located (e.g., *BCL2*, *BCL-xL*) [426]. |
| | ⊢ P | AKT_B | *Akt* phosphorylates *BAD* at Ser-136, inducing its sequestration away from the mitochondrial membrane where its BCL-2 family targets are located (e.g., *BCL2*, *BCL-xL*) [426]. |
| | ⊢ P | ERK | *ERK* phosphorylates *BAD* at Ser-112, inducing its sequestration away from the mitochondrial membrane where its BCL-2 family targets are located (e.g., *BCL-2*, *BCL-xL*)[427]. |
| | ⊢ P | S6K | *S6K1* phosphorylates *BAD* at Ser-155, directly blocking its binding to *BCL-xL* [428]. |
| | ← Lysis | Casp8 | *Caspase 8* is also able to cleave *BAD*, generating a more potently apoptotic fragment [425]. In addition, *TRAIL*-mediated apoptosis results in *BAD* cleavage by a Caspase upstream of MOMP, creating a potent apoptotic inducer before full *Caspase 3* activation [429]. |
| BIK | **BIK** = not(**MCL_1** or **BCLXL** or **BCL2**) | | |
| | Prot | | *BIK* is free to activate its target, *BAX*, only when it is not sequestered by any of the three BCL-2 family proteins in our model [418]. |
| | ⊢ IBind | MCL_1 | *MCL-1* binds *BIK*; they mutually inhibit each other [430]. |
| | ⊢ IBind | BCLXL | *BCL-xL* binds *BIK*; they mutually inhibit each other [431]. |

**Table S1n: Apoptotic_SW module**

| | | | |
|---|---|---|---|
| | ⊢ IBind | BCL2 | *BCL2* binds *BIK*; they mutually inhibit each other [423]. |
| BIM | **BIM** = **FoxO3** and **GSK3** and not(**ERK** or **MCL_1** or **BCLXL** or **BCL2**) | | |
| | Prot | | *BIM*'s pro-apoptotic activity requires expression driven by *FoxO3* and aided by *GSK3* and the absence of all three inhibitory *BCL2* family proteins. |
| | ← TR | FoxO3 | *FoxO3* is a transcriptional activator of *BIM* [432]. |
| | ← Ind | GSK3 | *GSK3* kinase is likely required for the *AP1*-dependent expression of *BIM* [433]. |
| | ⊢ Ind | ERK | The *MEK/ERK* pathway represses *BIM* protein levels, likely via transcriptional repression [434]. |
| | ⊢ IBind | MCL_1 | *MCL-1* binds *BIM* and inhibits its apoptotic activity [435]. |
| | ⊢ IBind | BCLXL | *BCL-xL* binds *BIM* and inhibits its apoptotic activity [422]. |
| | ⊢ IBind | BCL2 | *BLC2* binds *BIM* and inhibits its apoptotic activity [422]. |
| BID | **BID** = **Casp8** or (**Casp2** and (not(**BCL2** or **BCLXL** or **MCL_1**) or **ATM**)) | | |
| | Prot | | *BID* is truncated in response to *Caspase 8* activation. In addition, *Caspase 2* can also promote *BID* activation once all three pro-apoptotic *BCL2* family proteins are blocked. |
| | ← Lysis | Casp8 | In response to *TRAIL* (or *FAS* ligand), the initiator *Caspase 8* cleaves *BID* to its active truncated form [436, 437, 438]. |
| | ← Lysis | Casp2 | *Caspase 2* cleaves *BID* to its active truncated form [439]. |
| | ⊢ IBind | BCL2 | All three anti-apoptotic BCL2 proteins (*BCL2*, *BCL-xL* and *MCL-1*) sequesters *BID* into stable complexes, preventing them from activating *BAX* or *BAK* [440]. |
| | ⊢ IBind | BCLXL | All three anti-apoptotic BCL2 proteins (*BCL2*, *BCL-xL* and *MCL-1*) sequesters *BID* into stable complexes, preventing them from activating *BAX* or *BAK* [440]. |
| | ⊢ IBind | MCL_1 | All three anti-apoptotic BCL2 proteins (*BCL2*, *BCL-xL* and *MCL-1*) sequesters *BID* into stable complexes, preventing them from activating *BAX* or *BAK* [440]. |
| | ← P | ATM | *ATM* phosphorylates BID during mitosis, priming it for activation during mitotic catastrophe [441, 442]. Here we assume that this primed form of BID is more potentlty cleaved by *Caspase 2*, while it was shown not to alter its cleavage by *Caspase 8* [441]. |
| BAK | **BAK** = (**BID** and (**BIM** or **BIK** or not(**BCL2** and **BCLXL** and **MCL_1**))) or ((**BIM** or **BIK**) and not(**BCLXL** or **MCL_1**)) | | |

**Table S1n: Apoptotic_SW module**

| | | | |
|---|---|---|---|
| | Prot | Given that *BAK* is preferentially activated by *BID* compared to *BIM* [443] and that it is less responsive to sequestration by *BCL2* than the other two anti-apoptotic *BCL2* family proteins [444, 445], *BAK* in our model turns on when stimulated by *BID* if one or more *BCL2* family proteins are absent, or if *BIM* or *BIK* are also present. In contrast, *BIM* or *BIK* only activate *BAK* if *BCL-xL* and *MCL-1* are absent (*BCL-2* alone cannot block them). | |
| | ← Compl | BID | Activated (truncated) *BID* binds to mitochondrial *BAK*, resulting in its activation and oligomerization in the mitochondrial membrane, followed by *cytochrome c* release [446]. |
| | ← Compl | BIM | *BAK* is preferentially activated by *BID* compared to *BIM*, but *BIM* can also promote BAK oligomerization [443]. |
| | ← Compl | BIK | *BIK* can aid the activation of both *BAK* and *BAX* by triggering *BAK* oligomerization on the ER membrane and promoting a $Ca^{2+}$ efflux required for the fragmentation of hyper fused mitochondrial tubules, aiding *BAK* and *BAX* activation [447]. |
| | ⊢ IBind | BCL2 | *BCL2* can also bind *BAK* to prevent its oligomerization, but it does so less potently than the other two BCL-2 family members [444, 445, 448]. |
| | ⊢ IBind | BCLXL | *BCL-xL* binds *BAK* and prevent its oligomerization in the mitochondrial membrane [449, 444, 445]. |
| | ⊢ IBind | MCL_1 | *MCL-1* binds *BAK* and prevent its oligomerization in the mitochondrial membrane [444, 445]. |
| BAX | **BAX** = (**BIM** and (**BID** or **BIK** or not(**BCL2** and **BCLXL** and **MCL_1**))) or ((**BID** or **BIK**) and not(**BCL2** or **BCLXL**)) | | |
| | Prot | In contrast to *BAK*, *BAX* is preferentially activated by *BIM* compared to *BID* [443] and it is less responsive to sequestration by *MCL-1* than the other two anti-apoptotic BCL2 family proteins [444, 445]. *BAX* in our model turns on when stimulated by *BIM* if one or more *BCL2* family proteins are absent, or if *BID* or *BIK* are also present. In contrast, *BID* or *BIK* only activate *BAK* if *BCL2* and *BCL-xL* are both absent (*MCL-1* alone cannot block them). | |
| | ← Compl | BIM | Activated *BIM* binds to mitochondrial *BAX*, resulting in its allosteric activation and oligomerization in the mitochondrial membrane, leading to *cytochrome c* release [443]. |
| | ← Compl | BID | *BAX* is preferentially activated by *BIM* compared to *BID*, but *BID* can also promote *BAK* oligomerization [443]. |
| | ← Compl | BIK | *BIK* can aid the activation of both *BAK* and *BAX* by triggering *BAK* oligomerization on the ER membrane and promoting a $Ca^{2+}$ efflux required for the fragmentation of hyper fused mitochondrial tubules, aiding *BAK* and *BAX* activation [447]. |
| | ⊢ IBind | BCL2 | *BCL2* binds *BAX* and prevent its oligomerization in the mitochondrial membrane [444, 445]. |
| | ⊢ IBind | BCLXL | *BCL-xL* binds *BAX* and prevent its oligomerization in the mitochondrial membrane [444, 445]. |

**Table S1n: Apoptotic_SW module**

| | | | |
|---|---|---|---|
| | ⊣ IBind | MCL_1 | *MCL-1* can also bind *BAK* to prevent its oligomerization, but it does so less potently than the other two BCL-2 family members [444, 445, 450]. |
| Cyto_C | **Cyto_C** = **BAX** or **BAK** or (not **Mps1** and **Drp1** and **Unfused** and not **Mitophagy_High**) | | |
| | Prot | *Cytochrome C* release from mitochondria requires the oligomerization of either *BAK* or *BAX* [451]. In addition, *Drp1*-mediated mitochondrial fragmentation can aid cytochrome C release [452, 453], as long as the SAC protein *Mps1* [454] or high levels of mitophagy [455] do not counteract it. | |
| | ← Loc | BAK | *BAK* oligomerization at the mitochondrial membrane triggers MOMP, which results in the release of *cytochrome C* from mitochondria [451]. |
| | ← Loc | BAX | *BAX* oligomerization at the mitochondrial membrane triggers MOMP, which results in the release of *cytochrome C* from mitochondria [456, 457]. |
| | ⊣ Ind | Mps1 | During mitosis, fragmented mitochondria are blocked from leaking cytochrome C and killing the cell by mitochodrial recruitment of the SAC protein *Mps1* to the voltage-dependent anion channel 1 (*VDAC1*) [454]. |
| | ⊣ Ind | Drp1 | *Drp1* null cells show blunted and delayed cytochrome C release, due to a lack of *Drp1*-dependent remodelling and opening of cristae, where the majority of cytochrome c resides [453]. |
| | ← ComplProc | Unfused | Mitochondrial fission, which leads to loss of OPA1 function, opens the cristae junctions and releases cytochrome C [452]. |
| | ⊣ ComplProc | Mitophagy_High | High levels of mitophagy can engulf mitochondria debris and cytochrome C, and blunt the apoptosis signal [455]. |
| SMAC | **SMAC** = **BAX** or **BAK** | | |
| | Prot | *SMAC/Diablo* release from mitochondria requires the oligomerization of either *BAK* or *BAX* [451, 458]. | |
| | ← Loc | BAK | *BAK* oligomerization at the mitochondrial membrane triggers MOMP, which results in the release of *SMAC/Diablo* from mitochondria [451, 458]. |
| | ← Loc | BAX | *BAX* oligomerization at the mitochondrial membrane triggers MOMP, which results in the release of *SMAC/Diablo* from mitochondria [451, 458]. |
| IAPs | **IAPs** = **AKT_H** or not **SMAC** | | |
| | Prot | Inhibitor of Apoptosis Proteins (*IAP*s) are active in the absence of *SMAC* inhibition, or following *AKT_H* mediated upregulation (this protection from *SMAC* requires peak or oncogenic *AKT* activity). | |
| | ← Ind | AKT_H | *cIAP-2* and *XIAP* are both transcriptionally up-regulated in response to strong *PI3K/AKT1* activation [459]. |
| | ⊣ IBind | SMAC | *SMAC/Diablo* binds tightly to *IAP* proteins and blocks their ability to inhibit *Caspase 3* [460]. |

**Table S1n: Apoptotic_SW module**

| | | | |
|---|---|---|---|
| Casp9 | **Casp9** = (not **IAPs** and **Cyto_C**) or **Casp3** | | |
| | PTase | *Procaspase 9* is cleaved into active *Caspase 9* by *Caspase 3*, or by the apoptosome (which relies on *cytochrome C* for its assembly) in the absence of *IAP* proteins. | |
| | ⊣ IBind | IAPs | *XIAP*, *cIAP1* and *cIAP2* inhibit the *cytochrome c*-induced activation of *procaspase-9* [461]. |
| | ← Compl | Cyto_C | *Cytochrome c* binds to *APAF-1* proteins, promoting their assembly into the apoptosome, a platform for *procaspase 9* binding and cleavage into its active form [462]. |
| | ← Lysis | Casp3 | *Procaspase 9* is a direct cleavage target of *Caspase 3* [405]. |
| Casp3 | **Casp3** = (**Casp9** and **Casp8**) or (**Casp3** and (**Casp9** or **Casp8**)) or (not **IAPs** and (**Casp9** or **Casp8** or **Casp3**)) | | |
| | PTase | Activation of *Caspase 3* requires proteolytic cleavage of *procaspase-3* by initiator caspases such as *Caspase 9* or *Caspase 8*. In our model, cooperation of two of the three caspases (*Casp9*, *Casp8*, *Casp3*) is required in the presence of *IAP*s, which inhibit the proteolytic activity of *Caspase 3* by bind tightly to its active site. In the absence of *IAP*s, either of the three caspases can cleave and activate *Caspase 3*. | |
| | ← Lysis | Casp9 | Active *Caspase 9* cleaves *procaspase 3* [463]. |
| | ← Lysis | Casp8 | *Caspase 8* can cleave *Caspase 3* [464], but full *Caspase 3* activation also requires MOMP (potentially due to a need for *IAP* inhibition) [465]. |
| | ← Per | Casp3 | Once activated, *Caspase 3* helps sustain its own activation by cleaving *procaspase 8* and *6*. *Caspase 6*, in turn, generates additional active *caspase 8* and *9*. Together they all sustains a continuing active pool of Caspase 3. |
| | ⊣ IBind | IAPs | *IAP*s bind tightly to the active site of *Caspase 3*, keeping its activity in check [461, 466]. |

**Table S1o: DNA_Fragmentation module**

| Target Node | Node Gate<br>Node Type<br>Link Type | Node Description<br>Input Node | Link Description |
|---|---|---|---|
| CAD | **CAD** = **Casp3** and **Casp9** | | |

**Table S1o: DNA_Fragmentation module**

| | | |
|---|---|---|
| DNase | Caspase-activated DNase (*CAD*) is activated when its inhibition is released via the cleavage of *ICAD* (inhibitor of caspase-activated DNase). While *Capsase 3* and *7* (a direct target of *Caspase 9*) can inhibit *ICAD* [467], in our model they are both required, as *CAD* = ON is represents terminal, irreversible apoptotic commitment, which is fully locked in when both *Caspase 3* and *9* are on. | |
| ← Ind | Casp3 | *Caspase 3* relives *CAD* inhibition by cleaving its inhibitor *ICAD* [467]. |
| ← Ind | Casp9 | In addition of *Caspase 3*, *CAD* inhibition can also be relieved by *ICAD* cleavage by *Caspase 7*, which is a direct target of *Caspase 9* [467]. |

**Table S2a: Key to Node Type Symbols**

| Symbol | Node Type | Description |
|---|---|---|
| Cell | Cell | Node type used when a node's state is used to represent discrete phenotypes of an entire cell; appropriate for multi models. |
| DM | DM_Switch | Multi-stable regulatory module (Dynamically Modular Switch) in a higher-level model layer. |
| Conn | Connector | Grouping of nodes that are either mono-stable, or do not form a switch-like circuit that controls discrete phenotype transitions. For example, signaling input layers without feedback, or multi-step links between distinct switches can be represented as connector nodes in coarse-grained (switch-level) models. |
| Env | Environment | Nodes or modules that represent the extracellular environment of a single cell or cell collective. These are self-sustaining nodes or node groups that maintain their initial states and receive no feedback from the rest of the network (they act as inputs). |
| Proc | Process | Nodes or modules that stand in for complex cellular processes not modeled in detail (e.g., DNA replication or the process of aligning chromosomes at the metaphase plane during mitosis). |
| MSt | Macro_Structure | Nodes or modules that represent the state of large, complex cellular structures such as DNA content, cytoskeletal features, junctions or mitochondria. |
| Met | Metabolite | Regulatory node representing a metabolite (not protein, gene product or complex structure). |
| mRNA | MRNA | mRNA. |
| miR | MicroRNA | microRNA. |
| PC | Protein_Complex | Protein complex represented by a single node or via a key member of the complex. |
| Rec | Receptor | Cell surface receptor protein or complex. |
| Adap | Adaptor_Protein | Protein that helps scaffold a signaling complex or other large assembly of proteins. |
| Secr | Secreted_Protein | Protein secreted into the extracellular environment, such that the state of the node tagged with this type represents the availability fo this protein outside the cell. |
| TF | TF_Protein | Transcription factor. |
| K | Kinase | Kinase (enzyme that catalyzes the phosphorylation of its target). |
| Ph | Phosphatase | Phosphatase (enzyme that catalyzes the removal of phosphorylation from its target). |
| UbL | Ubiquitin_Ligase | Ubiquitin ligase (protein that recruits an ubiquitin-conjugating enzyme that has been loaded with ubiquitin to a target protein and assists or directly catalyzes the transfer of ubiquitin from the ubiquitin-conjugating enzyme to the target). |
| PTase | Protease | Protease (enzyme that catalyzes the breakdown of proteins into smaller fragments). |
| DNase | DNase | Deoxyribonuclease (DNase, for short); endonuclease that catalyzes the hydrolytic cleavage of the DNA backbone. |

**Table S2a: Key to Node Type Symbols**

| | | |
|---|---|---|
| CAM | CAM | Cell adhesion proteins located on the cell surface. |
| CDK | CDK | Cyclin-dependent kinase. |
| CDKI | CDKI | Cyclin-dependent kinase inhibitor. |
| GEF | GEF | Guanine nucleotide exchange factor. |
| GAP | GAP | GTPase-activating protein (also called GTPase-accelerating protein). |
| GTPa | GTPase | GTPase enzymes that hydrolyze ATP to ADP. |
| Enz | Enzyme | Enzyme that does not fit the more specific enzyme categories listed above. |
| Prot | Protein | Regulatory protein that does not fit any of the more specific classifications listed above. |
| MP | Membrane_Potential | A relative measure of membrane potential across a biological membrane, generally indicating whether this potential is within the normal range, or abnormally low / high in a way that affects other regulatory processes. |
| lncRNA | LncRNA | Long intervening noncoding RNA |
| SLig | Cell_Surgace_Ligand | Membrane-bound signaling molecule that serves as a ligand to receptors on neighboring cells. |

**Table S2b: Key to Link Type Symbols**

| Symbol | Link Type | Description |
|---|---|---|
| Env | Enforced_Env | This link type represents self-loops on Environment nodes, which guarantee that these nodes maintain their initial state throughout a time-course simulation unless they are explicitly altered by the simulation's settings. |
| Ind | Indirect | Regulatory influence that does not involve direct binding, processing, or enzyme activity. |
| ComplProc | Complex_Process | Regulatory influence that is not modeled in detail, but involves more than one molecule or a macrostructure. For example, the physical need for kinetochores on replicated sister chromatids for the assembly of certain protein complexes can be represented as a link from the node representing kinetochores to the regulatory proteins, with a Complex_Process link type. |
| Per | Persistence | This link type represents self-loops that alter the ability of a node to stay in a particular state depending on its own current state. For example, if transcription of a protein is easier to maintain than to induce de novo, this may be encoded by a logic gate that includes the node itself and creates a self-loop. The link type of this loop is "Persistence". |
| TR | Transcription | Action of a transcription factor to alter the expression of the target node (mRNA or protein). Link type should be used for induction as well as repression (the link effect contains this information). |

**Table S2b: Key to Link Type Symbols**

| | | |
|---|---|---|
| TL | Translation | Regulatory influence that controls the translation of mRNA into protein; should be used for induction as well as repression of translation. |
| Ligand | Ligand_Binding | Binding of extracellular ligand to its receptor. |
| Compl | Complex_Formation | Binding even that leads to a regulatory protein complex. |
| IBind | Inhibitory_Binding | Binding even that represses the target node's level or activity. |
| Loc | Localization | Regulatory influence that alters the localization of a molecule. |
| BLoc | Binding_Localization | Binding even that alters the localization of a molecule. |
| PBind | Protective_Binding | Binding even that increases / protects the target node's activity. |
| Unbind | Unbinding | A regulatory influence that causes the target node to be released from a protein complex and change its activity (increase or decrease) as a result. |
| P | Phosphorylation | Phosphorylation. |
| DP | Dephosphorylation | Dehosphorylation. |
| PLoc | Phosphorylation_Localization | Phosphorylation resulting in altered protein localization. |
| Ubiq | Ubiquitination | Ubiquitination, usually leading to protein degradation. |
| Deg | Degradation | Regulatory influence leading to the degradation of the target molecule (more general than Ubiquitination; the latter link type should be used when appropriate). |
| GEF | GEF_Activity | Action of a Guanine nucleotide exchange factor (GEF) leading to GTP loading onto (and usually the activation of) a GTPase. |
| GAP | GAP_Activity | Actions of a GTPase-activating protein (GAP) leading to the hydrolysis of GTP by (and usually de-activation of) a GTPase. |
| Lysis | Proteolysis | Protein cleavage. |
| Cat | Catalysis | Increasing the rate of metabolite production by an enzyme. |
| Epi | Epigenetic | Process that alters gene expression via modifying chromatin condensation or altering DNA methylation. |
| TrConf | Transcription_Conflict | Transcription from the same DNA locus using an alternate start site or proceeding in another direction, leading to a conflict between two transcriptional events. |
| Secr | Secretion | Secretion or shedding of a protein or other regulatory molecule to the extracellular environment. |
| RNAi | RNAi | This process represents inhibitory binding of cytoplasmic mRNAs by RISC-bound microRNAs that block translation and/or enhance mRNA degradation. |
| Acet | Acetylation | Acetylation |
| Deacet | Deacetylation | Deacetylation |
| -OH | Hydroxylation | Hydroxylation |

**Table S2c: Key to Link Effect Symbols**

**Table S2c: Key to Link Effect Symbols**

| Symbol | Link Effect | Description |
|---|---|---|
| $\leftarrow$ | Activation | Link in which the input node aids the expression, activity, persistence or localization of the target such that the target is easier to turn/keep in an ON state. It can be used for multi-level nodes as long as these levels represent increasing intervals of activity. |
| $\vdash$ | Repression | Link in which the input node hinders the expression, activity, persistence or localization of the target such that the target is easier to turn/keep in an OFF state. It can be used for multi-level nodes as long as these levels represent increasing intervals of activity. |
| $\bullet\!-$ | Context_Dependent | Link acting on a node in such a way that it activates it under certain conditions (e.g., when another input is OFF), but represses it when this condition is not met. The XOR gate is a good example of a context-dependent link effect. |
| $\perp$ | Inapt | This link-type refers to connections between complex regulatory switches (rather than molecules), where categorizing the effect of an input as Activation or Repression, or even context-depended activation or repression does not apply. This is usually the case with multi-state switches, where the 3 or more phenotypes represented by the discrete states of these switches do not have a meaningful ordering. Thus, stating that this switch is "activated" by another one is not appropriate. |

[89] Bruce Alberts, Alexander Johnson, Julian Lewis, David Morgan, Martin Raff, Keith Roberts, and Peter Walter. *Molecular Biology of the Cell.* W.W. Norton & Company, 6 edition, August 2017.

[140] Joshua C. Drake, Rebecca J. Wilson, Rhianna C. Laker, Yuntian Guan, Hannah R. Spaulding, Anna S. Nichenko, Wenqing Shen, Huayu Shang, Maya V. Dorn, Kian Huang, Mei Zhang, Aloka B. Bandara, Matthew H. Brisendine, Jennifer A. Kashatus, Poonam R. Sharma, Alexander Young, Jitendra Gautam, Ruofan Cao, Horst Wallrabe, Paul A. Chang, Michael Wong, Eric M. Desjardins, Simon A. Hawley, George J. Christ, David F. Kashatus, Clint L. Miller, Matthew J. Wolf, Ammasi Periasamy, Gregory R. Steinberg, D. Grahame Hardie, and Zhen Yan. Mitochondria-localized AMPK responds to local energetics and contributes to exercise and energetic stress-induced mitophagy. *Proceedings of the National Academy of Sciences of the United States of America*, 118(37):e2025932118, September 2021.
